# Binocular Mirror-Symmetric Microsaccadic Sampling Enables *Drosophila* Hyperacute 3D-Vision

**DOI:** 10.1101/2021.05.03.442473

**Authors:** Joni Kemppainen, Ben Scales, Keivan Razban Haghighi, Jouni Takalo, Neveen Mansour, James McManus, Gabor Leko, Paulus Saari, James Hurcomb, Andra Antohi, Jussi-Petteri Suuronen, Florence Blanchard, Roger C. Hardie, Zhuoyi Song, Mark Hampton, Marina Eckermann, Fabian Westermeier, Jasper Frohn, Hugo Hoekstra, Chi-Hon Lee, Marko Huttula, Rajmund Mokso, Mikko Juusola

## Abstract

Neural mechanisms behind stereopsis, which requires simultaneous disparity inputs from two eyes, have remained mysterious. Here we show how ultrafast mirror-symmetric photomechanical contractions in the frontal forward-facing left and right eye photoreceptors give *Drosophila* super-resolution 3D-vision. By interlinking multiscale *in vivo* assays with multiscale simulations, we reveal how these photoreceptor microsaccades - by verging, diverging and narrowing the eyes’ overlapping receptive fields - channel depth information, as phasic binocular image motion disparity signals in time. We further show how peripherally, outside stereopsis, microsaccadic sampling tracks a flying fly’s optic flow field to better resolve the world in motion. These results change our understanding of how insect compound eyes work and suggest a general dynamic stereo-information sampling strategy for animals, robots and sensors.

**Significance statement:** To move efficiently, animals must continuously work out their x,y,z-positions in respect to real-world objects, and many animals have a pair of eyes to achieve this. How photoreceptors actively sample the eyes’ optical image disparity is not understood because this fundamental information-limiting step has not been investigated *in vivo* over the eyes’ whole sampling matrix. This integrative multiscale study will advance our current understanding of stereopsis from static image disparity comparison to a new morphodynamic active sampling theory. It shows how photomechanical photoreceptor microsaccades enable *Drosophila* super-resolution 3D-vision and proposes neural computations for accurately predicting these flies’ depth-perception dynamics, limits, and visual behaviors.

## Introduction

Historically, stereo vision studies have focused on the disparity between the left and right eye images and how this is processed in the brain (1–7). Less attention has been paid to how the peripheral visual systems actively sample and encode depth information. This trend has been particularly notable with insect vision. Because the insect compound eyes are composed of rigid ommatidial lens systems, it was long thought that their static functional organization provides a pixelated low-resolution image of the world, often with little or no depth information (8, 9).

Remarkably, recent studies have revealed the morphodynamic, active nature of the fruit fly (*Drosophila melanogaster*) early vision in information capture (10, 11). Underneath the ommatidial lenses, light changes make photoreceptors rapidly contract (10, 11) and elongate in and out of their focal plane and sideways in a sophisticated piston motion (11). These microsaccades adjust the photoreceptors’ receptive field sizes and x,y-positions dynamically, sharpening light input in time to provide dynamic hyperacute vision beyond the compound eyes’ static optical resolution (11). With phototransduction reactions themselves - PIP_2_ cleavage from the cell membrane (10) - causing the microsaccades, a photoreceptor’s photon sampling itself initiates active vision (11). But it has remained unclear how these microsaccades happen globally, across the left and right eye, and whether and how they could contribute to visual behaviors and stereo vision.

Here, we study how the *Drosophila* photoreceptor microsaccades are organized (adapted) to the world order - its physical regularities - across the two eyes to sample information. We do this first *globally*, across the left and right eyes of living wild-type and mutant/transgenic fly strains, using ultrafast high-brilliance X-ray imaging (ESRF and DESY synchrotrons generating X-ray magnitudes >10^5^-times the conventional X-ray tubes) with electrophysiology. Combined with *local* high-speed photoreceptor and visual interneuron (LMC) recordings, these results show that photoreceptor microsaccade directions and dynamics are hardwired during development to match the optic flow of a locomoting fly, maximizing visual information capture. Because this active sampling is mirror-symmetric between the left and the right eye, it enables *Drosophila* hyperacute stereopsis. By implementing these experimental results into theoretical multiscale models, we simulate the adaptive *Drosophila* compound eye optics with photoreceptor microsaccades sampling light information across the eyes. Finally, we show how this new binocular active sampling theory accurately estimates object depth and predicts various visual behaviors.

## Results

To examine the global photoreceptor photomechanics in sub-micrometer spatial and ≤10 ms temporal resolution inside the compound eyes of intact living *Drosophila*, we performed *in vivo* X-ray imaging at the ESRF (beamline ID16b) and DESY (beamline P10) synchrotrons (Fig. 1*A*; Fig. S1).

**Fig. 1.**
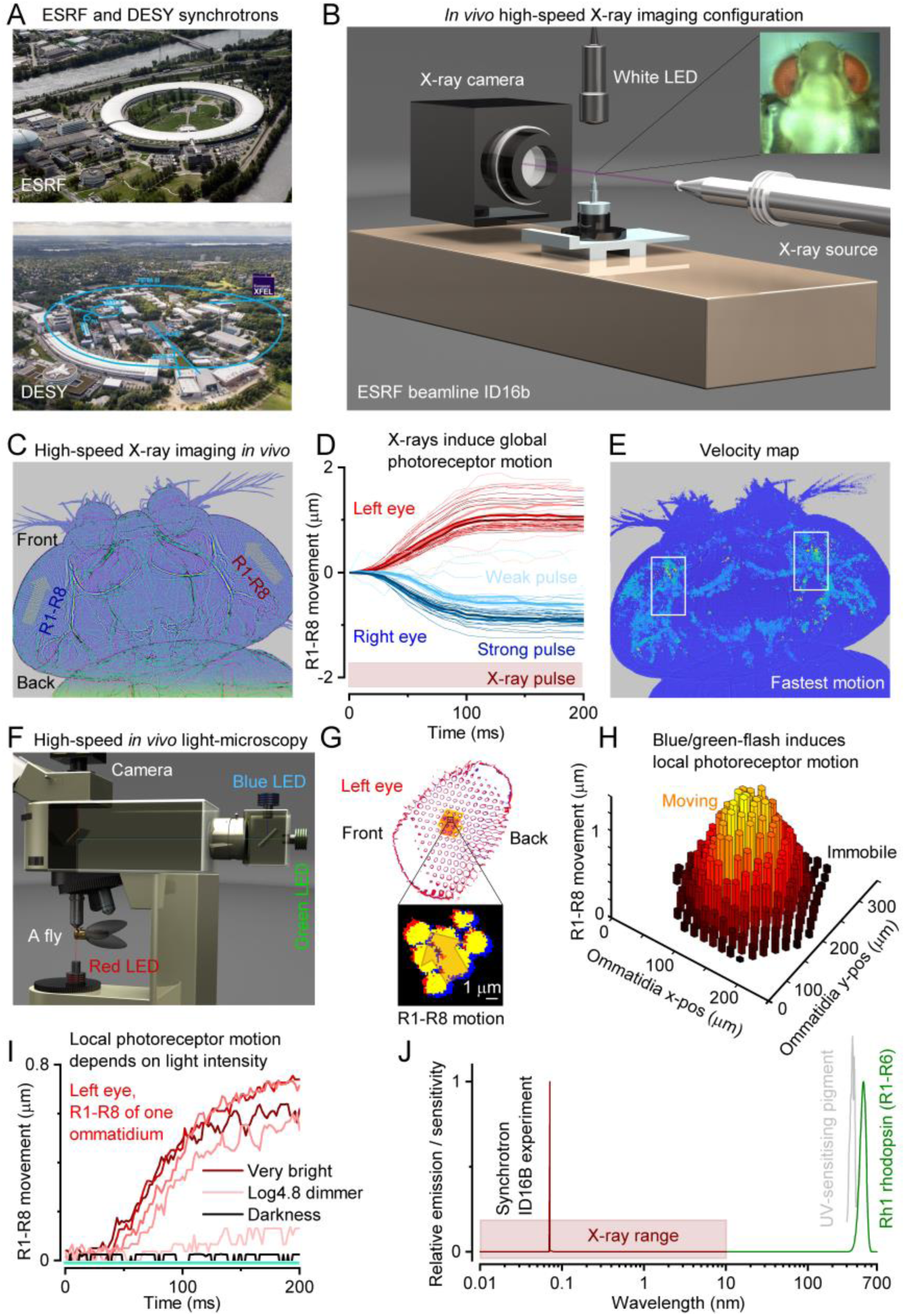
X-ray-imaging *Drosophila in vivo* reveals global mirror-symmetric right and left eye photoreceptor contraction dynamics that tie in with local photomechanical photoreceptor responses. (*A*) Experiments were performed using synchrotrons (see Fig. S1 to S3, S7 to S9). (*B*) ESRF beamline ID16b imaging configuration, using 100 nm resolution. (*C*) X-rays evoked fast synchronized mirror-symmetric photoreceptor (R1-R8) contractions inside the left and right eyes, causing the photoreceptors to sweep in global back-to-front vergence motion (arrows). (*D*) Photoreceptor movement began <10 ms from the X-ray onset, increasing with intensity until saturating. (*E*) The longest frontal forward-facing photoreceptors (12) moved the fastest, ∼15-20 µm/s. (*F*) High-speed light-microscopy of R1-R8 rhabdomere photomechanics to blue-green flashes under deep-red antidromic illumination (740 nm LED + 720 nm edge filter), with a fly held in a pipette tip. (*G*) A 200 ms blue/green-flash, delivered orthodromically (through the microscope optics) into the left fly eye (above), excited local photoreceptors (orange highlight) to twitch photomechanically in a back-to-front direction (arrow). (*H*) Rhabdomeres moved only in the ommatidia facing the incident blue/green-flash from above and remained still in the other ommatidia (11). Thus, R1-R8 motion did not involve intraocular muscles (each eye has a pair (13)), which otherwise would have moved the whole retina (14). (*I*) Local blue/green-light-induced photoreceptor movements’ early fast-phase depended upon the light intensity and closely resembled those evoked by X-rays (*D*). (*G* and *H*) R1-R8 of one ommatidium contracted together as a unit if any of their R1-R8 alone saw light changes, indicating intraommatidial mechanical photoreceptor coupling; see fig S32 and S33. (*J*) The experimental X-ray wavelength peak was ∼6,900-times shorter than R1-R6s’ peak sensitivity (∼480 nm).

### X-rays evoke mirror-symmetric photoreceptor motion in the left and right eye

We first imaged the compound eyes by brief (200-300 ms) high-intensity X-ray flashes (Fig. 1*B*; Fig. S2), which would limit radiation damage, while simultaneously activating local photoreceptors by a white LED flash, inducing their contraction. Unexpectedly, however, we found that the X-rays alone could rapidly (≤10 ms) activate every photoreceptor to contract in synchrony, causing them to sweep mirror-symmetrically inside the left and right eye in an opposing back-to-front vergence motion (Fig. 1, *C* and *D*; Fig. S3; Movie S1). This global motion’s size and speed increased broadly with X-ray intensity (Fig. 1*D*) and was large enough to conceal local photoreceptor contractions to the simultaneous LED test flashes. Velocity analyses further revealed that X-rays caused the strongest movements in the left and right eyes’ forward-facing photoreceptor pairs with the longest light-sensitive parts, the rhabdomeres (12), where the photomechanical transduction occurs (10, 11) (Fig. 1*E*; Fig. S3*E* and *F*; Movie S1).

These movements were not caused by radiation- or heat-induced tissue swelling or damage because immediately, as the X-ray stimulation was shut off in darkness, the photoreceptors stretched back to their original shapes within a second, enabling their contractions to be repeated for many minutes, sometimes ≥30 minutes. And crucially, the contractions stopped when the fly died and did not appear in freshly killed flies. Moreover, separate light-microscopy experiments through cornea-neutralized ommatidia (Fig. 1*F*; Fig. S31 to S33) revealed that 200 ms blue/green-flashes (presented within the photoreceptors’ receptive fields) made these cells contract with comparable motion directions (Fig. 1, *G* and *C*), time course and intensity-dependence (Fig. 1, *H* and *D*). These findings suggest that X-rays and visible light elicited the contractions through the same mechanism, requiring phototransduction activation. Interestingly, however, we further discovered that R1-R8 are mechanically coupled in an ommatidium. Activating a single photoreceptor out of R1-R8 within an ommatidium induced them all to contract simultaneously as a unit, without affecting photoreceptors in neighboring ommatidia (Fig 1, *G* to *I*; Fig. S32 and S33). Thus, the screening pigments around the ommatidia work to insulate the photoreceptors from non-incidental visible light contracting them, but this function fails with X-ray radiation.

We hypothesized that sufficiently high X-ray photon densities could either activate phototransduction directly through rhodopsin photo-isomerization (15, 16) or release visible photons through Compton scattering from the heavier atoms inside the eye (17), for example, from phosphorus in the membrane phospholipids, or radiation phosphene (18). Such low-energy photons would then photo-isomerize rhodopsin molecules or be absorbed by ommatidial screening pigments, preventing light from leaving the eye. The probability of an X-ray photon (λ_x_ ≈ 0.07 nm) activating a single rhodopsin-molecule (Rh1, λ_max_ ≈ 330 [UV-sensitizing pigment] and 480 nm [blue-green]) should be infinitesimal (Fig. 1*J*). Yet, each photoreceptor has millions of rhodopsin molecules and face ∼10^6-8^ X-ray photons in the synchrotron beam at each second. In these extreme conditions, rhodopsin photo-isomerizations – and the subsequent fast PIP_2_ cleavage from the photoreceptor membrane, as the plausible mechanism of photoreceptor contractions (10) – may become unavoidable.

### X-ray-activated phototransduction uncovers global R1-R8 microsaccade dynamics

We tested this hypothesis *in vivo* by recording wild-type and blind mutant (*hdc^JK910^*, *norpA^P24^* and *trp;trpl*) flies’ global electrical responses, so-called electroretinograms (ERGs), to 250 ms white-light and X-ray flashes (Fig. 2; Fig. S4 to S6; Movie S2) at DESY beamline P10 (Fig. 2*A*). These experiments, measuring retina-wide simultaneous photoreceptor activations, were performed by a remote-controlled LED stimulation/ERG recording system (Fig. 2*B*), synchronized with 100 fps high-speed X-ray imaging, after carefully positioning a recording microelectrode on the right eye and a reference electrode in the thorax and letting the flies dark-adapt for 1-2 minutes.

**Fig. 2.**
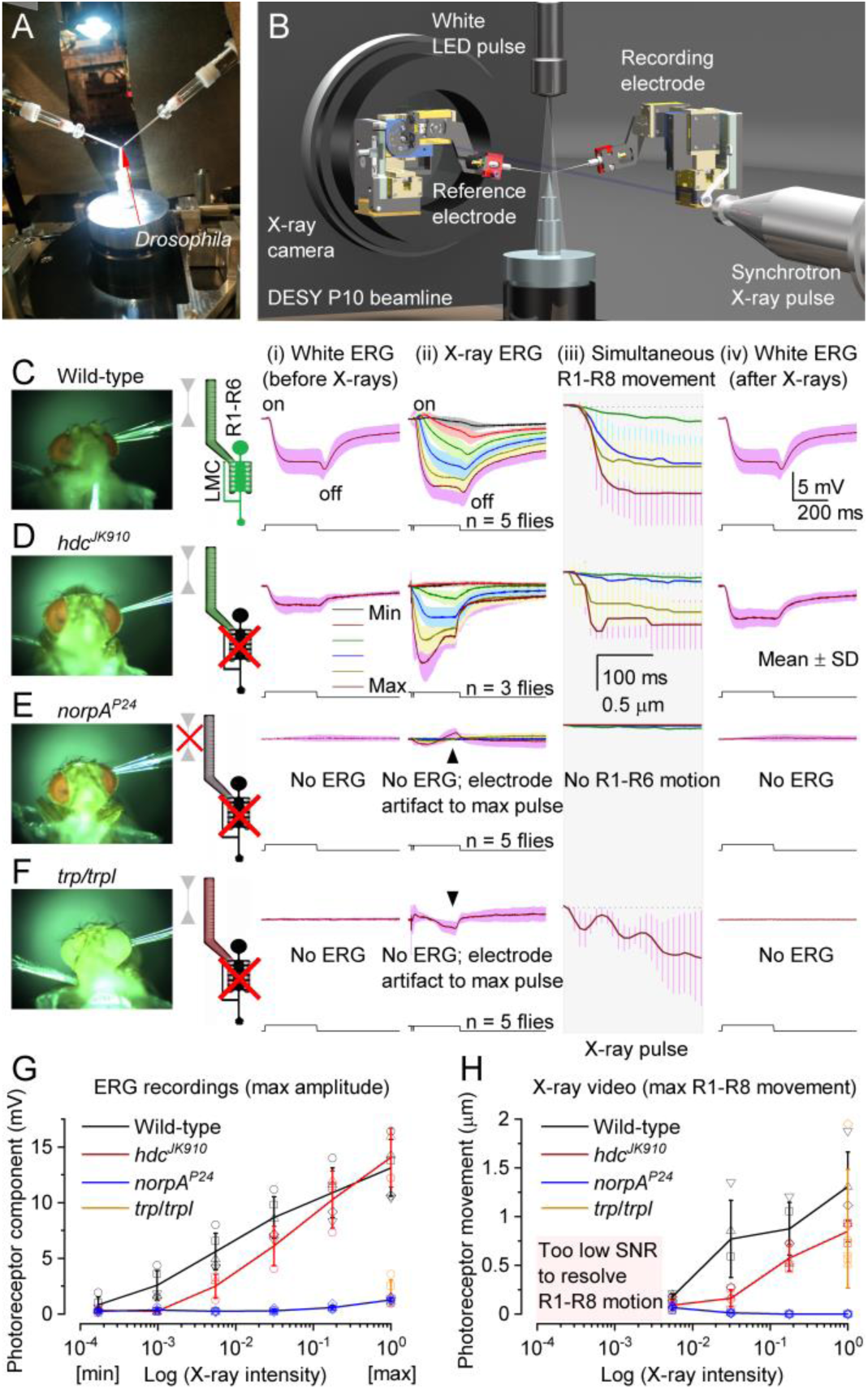
X-rays activate phototransduction. (*A*) Remote-controlled stimulation and recording system for head-fixed *Drosophila*, including two piezo-micromanipulators, an ERG amplifier, and a white LED, fitted in DESY P10 beamline. (*B*) Microelectrodes recorded the fly eyes’ combined response, electroretinogram (ERG), to white-light and X-ray pulses. (*C*) Wild-type ERGs to a white-light (i) and X-rays (ii) show on- and off-transients, indicating normal histaminergic synaptic transfer. Hyperpolarizing photoreceptor ERG component and (iii) R1-R8 photomechanical contraction increased with X-ray intensity. (iv) White-light ERG control recorded 20 s after the X-rays. (*D*) *hdc^JK910^* (i) white and (ii) X-ray ERGs lacked On- and Off-transients, indicating missing synaptic transfer. (ii) ERG photoreceptor component and (iii) R1-R8 photomechanics increased with X-ray intensity. (iv) *hdc^JK910^* white ERG control recorded 20 s later. (*E*) Blind *norpA^P24^* do not generate ERG responses or photomechanical photoreceptor contractions to white light or X-ray pulses. (*F*) Blind *trp;trpl* do not generate ERG responses while their photoreceptors contract photomechanically to white or X-ray pulses but in a less coordinated way. (*C* to *F*) In the R1-R6/LMC cartoons, green indicates the normal function, gray R1-R6 no contraction, and the black LMC no synaptic output. (*G*) Wild-type and *hdc^JK910^* ERG photoreceptor components increased sigmoidally with X-ray intensity, while those of *norpA^P24^* and *trp;trpl* did not respond. (*H*) Wild-type and *hdc^JK910^* photomechanical responses grew sigmoidally with X-ray intensity, while those of *norpA^P24^* did not respond. The maximal X-ray-induced photoreceptor contraction in *trp;trpl* (orange) was comparable to the wild-type and *hdc^JK910^*. (*G* and *H*) The normalized maximum intensity corresponds to 2.2 x 10^6^ photons/s/mm^2^.

Wild-type white-light control ERGs (Fig. 2*C*, i) showed a typical hyperpolarizing photoreceptor component between On- and Off-transients from the postsynaptic interneurons (19), LMCs. Remarkably, the test ERGs to progressively intensified X-ray flashes (ii), recorded 20 s after, showed comparable dynamics, suggesting that X-rays activated phototransduction, causing an electrical photoreceptor signal and its synaptic transmission. The photoreceptor component increased with the X-ray intensity, consistent with normal elementary response (quantum bump) integration (11). For the two brightest X-ray flashes, this component was larger than the white-flash one, presumably because the X-rays activated every photoreceptor in the eye (global activation). In contrast, the white-LED activated mostly the photoreceptors directly facing it (local activation). Importantly, high-speed imaging (iii) showed that the X-ray-evoked photoreceptor contractions closely followed their ERG dynamics (Movie S2), supporting the direct phototransduction-activation hypothesis. The robust control ERGs (iv), recorded after the X-rays, implied that the eyes worked normally with little (or no) radiation damage.

*hdc^JK910^*-mutant ERGs (Fig. 2*D*) gave further evidence that visible light (i) and X-rays (ii) activated phototransduction analogously. Both types of stimuli evoked photoreceptor components but no On- and Off-transients, consistent with *hdc^JK910^*-photoreceptors’ inability to synthesize neurotransmitter histamine and transmit visual information to LMCs and the brain (20). While the *hdc^JK910^*-phototransduction approximates wild-type (11, 20), histamine deficiency has been shown to cause an excitatory synaptic feedback overload from the lamina interneurons to R1-R6s, making *hdc^JK910^*-photoreceptors more depolarized with faster responses and reduced light-sensitivity in respect to the wild-type (20) (cf. Fig. 2*D*, i and iv to Fig. 2*C*, i and iv). Accordingly, and in further support of our hypothesis, we found both the *hdc^JK910^* X-ray ERG dynamics (Fig. 2*D*, ii) and photomechanical contractions (Fig. 2*D*, iii) faster and less sensitive than in the wild-type (Fig. 2*C*, ii-iii) over a broad intensity range (Fig. 2, *G* and *H*).

Conversely, *norpA^P24^*-mutants, in which faulty phospholipase-C molecules halt phototransduction PIP_2_ activation (10), showed (Fig. 2*E*) neither clear electrical responses to visible light (i) or X-rays (ii), producing effectively flat no-change ERGs (bar the small electrode charging artifacts), nor reacted photomechanically (iii) over the test intensity range (Fig. 2, *G* to *H*). Although similar “zero-response” controls were recorded from freshly killed flies (by freezing; Fig. S5*C*), concurrent X-ray imaging revealed that *norpA^P24^*-mutants were alive and active during the stimulation, seen by their antennal movements and intrinsic muscle activity. Thus, these results validated that the wild-type (Fig. 2*C*) and *hdc^JK910^* X-ray responses (Fig. 2*D*) were not caused by tissue shrinkage, damage, or movement artifacts but resulted from phototransduction activation.

Finally, we used *trp;trpl*-mutants (Fig. 2*F*), which can respond photomechanically to light flashes by cleaving PIP_2_’s bulky headgroup (InsP_3_) from the microvillar membrane (10) but not electrically because they lack the light-gated ion channels, which are required to open for generating electrical responses and synaptic signaling. Thus, these mutants provided a decisive test of whether the X-ray-induced photoreceptor movements (Fig. 1, Fig. 2, *A* to *E*) were photomechanical. However, owing to their minutes-long light recovery time (11), we used only one bright X-ray intensity. We found that *trp;trpl*-mutants neither responded electrically to white-light (i and iv) nor X-ray flashes (ii), but their photoreceptors contracted strongly both to X-rays (iii) and visible light (10, 11). Meaning, these movements were photomechanical, induced by phototransduction PIP_2_-cleavage. And whilst their dynamics showed characteristic oscillations after contracting ∼40-50 ms (11), these were unrelated to missing eye-muscle activation (each eye has a pair (13)). This is because, in the head-fixed wild-type flies, the local photoreceptor activation (Fig. 1F) did not trigger intraocular muscle contractions (Fig. 1*G*; Fig. S32 and S33), and yet their local and global photomechanics ensued alike (*cf.* Fig 2*C* to Fig. 1, *D* and *H*). Therefore, the *trp;trpl*-oscillations more likely reflected suboptimal Ca^2+^-dynamics [missing Ca^2+^-influx], mechanical damping/anchoring or both.

These results (Fig. 1 and 2) showed that a *Drosophila* photoreceptor responds to both X-rays and visible light but with different probabilities and that the synchrotron-based X-ray imaging activates all photoreceptors inside the left and right eye at once, revealing their photomechanical mirror-symmetric motion dynamics (Movie S1 and S2), hidden from the outside view. Interestingly, these global R1-R8 microsaccade dynamics suggest that when experiencing contrast variations in natural scenes, the two eyes’ frontal forward-facing photoreceptor pairs, which are ∼400 µm apart but should have overlapping receptive fields (RFs), would scan over the same small visual area in opposing but synchronized vergence motion. We, therefore, next asked whether the frontal photoreceptors sample the world in this way?

### Left and right eye photoreceptor receptive fields move mirror-symmetrically

To answer this question, we built a head-centered goniometric 2-axis rotation stage with an integrated microscope/high-speed camera system for targeted rhabdomere light stimulation and motion capture (Fig. 3*A*; Fig. S10). This device allowed us to measure a head-fixed *Drosophila*’s photoreceptor rhabdomeres’ x,y-positions *in situ* (Fig. 3, *B* to *D*; Fig. S11 to S14), as visualized by their virtual images, so-called deep-pseudopupils (DPPs) (21), to antidromic infra-red illumination (≥820 nm, propagating through its head/eyes), which the flies cannot see (11, 22). Moreover, to capture their photomechanical contractions (Fig. 3 *E* and *F*), the rhabdomeres could be stimulated orthodromically, through the ommatidial lens system, with light flashes presented at their RFs.

**Fig. 3.**
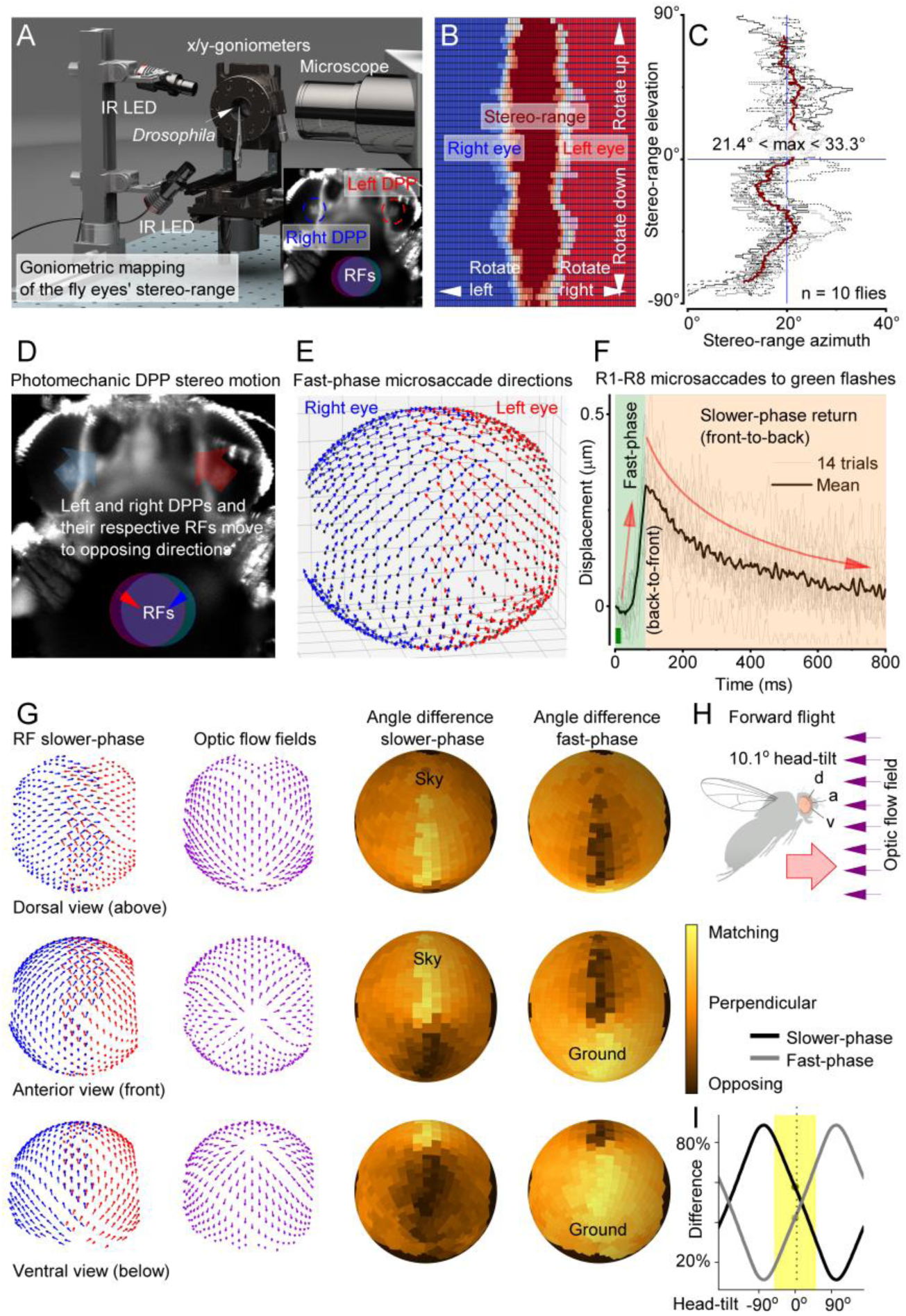
Left and right eye photoreceptor receptive fields (RFs) overlap frontally and move mirror-symmetrically, tracing forward translation induced optic flow. (*A*) A goniometric high-speed imaging system for mapping photoreceptors’ RFs. Inset: Infra-red (IR) back-lit R1-R7/8 photoreceptor rhabdomeres, forming the left and right eye deep-pseudopupils (DPPs) (21) (circled), ∼10x-magnified by the ommatidial lenses. Each eye’s DPP shows rhabdomeres from neighboring ommatidia that collect light from overlapping RFs (superposition). (*B*) Rotating the fly head through its central x,y-axes revealed its DPPs’ stereoscopic visual field (vine color); see Movie S3. (*C*) Because the frontal photoreceptors’ RFs overlap binocularly (∼23.5° azimuth, ∼180° elevation), these mirror-symmetric pairs could enable depth perception. (*D*) Ommatidial lenses invert the eyes’ fast up-medially-recoiling microsaccades (DPP fast-phase; big arrows), evoked by a 10-ms light flash within their overlapping RFs, to sweep their respective RFs down-laterally (small arrows). (*E*) Microsaccade fast-phase directions mapped across the left (red) and right (blue) eyes; slower-phase return in the opposite direction (cf. Movie S4; mean of 5 ♂ flies). (*F*) Brightening (10 ms light-flash) contracts R1-R8 front-to-back (fast-phase), and darkening returns them back-to-front (slower-phase); their RFs move in the opposite directions. The mean (black) and 14 consecutive R1-R7/8 contractions (light-grey), recorded through cornea-neutralized optics (cf. Fig. 1G); see Fig. S23 and S24 for fully light-adapted dynamics (Movie S5). (*G*) The corresponding slower-phase RF vector map (left) compared to the forward flying fly’s optic flow field (center), as experienced with the fly-head upright. Their difference (error) is shown for the slower- and fast-phases. The fast-phase matches the “ground-flow”, the slower-phase the “sky-flow”. (*H*) By adjusting microsaccadic sampling to optic flow through head-tilt, a fly can actively keep the passing world longer within its photoreceptors’ RFs, which theoretically should improve acuity to resolve the world in motion (see Fig. S56 to S61; Movie S6). d, dorsal; a, anterior; v, ventral viewpoints. (*I*) Upright (0°) head, and normal tilting around this position (yellow), keep RFs’ fast- and slower-phases in a balanced push-pull sampling state. Optimizing vision for specific behaviors, like object tracking, requires further self-adjustments in locomotion speed and head and body movements (Movie S7; Fig. S25 to S27).

We first identified those frontal photoreceptors in the left and right eye, which had overlapping RFs (Fig. 3*B*; Fig. S13 and S14) by systematically mapping their x,y-positions (Fig. 3*C*) with head-centric fine-rotations (0.35° step; Movie S3). These measurements revealed the eyes’ stereoscopic layout, where owing to the eyes’ optical superposition design (21, 23), a single point in space frontally is seen at least by 16 photoreceptors; the R1-R8 super-positioned in the left eye and the R1-R8 super-positioned in the right eye (Fig. 3 *B* and *C*). We further mapped how R1-R8 rhabdomeres, as revealed by the DPP images, were systematically rotated during ontogenic development for each eye location while retaining optical superposition with the changing eye curvature. This scanning revealed the left and right eyes’ highly-ordered mirror-symmetric R1-R8 angular orientation maps, with equatorial mirror-symmetricity (21) between the eyes’ upper and lower halves (Fig. S11 and S12).

Next, we analyzed the rhabdomeres’ photomechanical movement directions to UV- or green-light flashes (Fig. 3*D*; Fig. S15 to S30), as delivered at their RFs (Movie S4). The resulting deep-pseudopupil microsaccades were then translated into a 3D-vector map (Fig. 3*E*), covering the frontal stereo section and more peripheral parts of the eyes. Expectedly, the left (red) and right (blue) eye microsaccades were mirror-symmetric. But crucially, by comparing these movement maps to the deep pseudopupil angular orientation maps for each eye location (Fig. S12), we found that the local microsaccades occurred along their R1-R2-R3 photoreceptors’ rotation axis, implying that their sideways-movement directions were hardwired during development. Moreover, because DPPs are virtual images (24), which are magnified but not inverted by the ommatidial lens system (Movie S4; Fig. S15 to S19), the rhabdomeres inside the eyes recoiled accordingly (Fig. 3*F*); first bouncing along their location-specific back-to-front directions (fast-phase) before returning front-to-back (slower-phase), consistent with the X-ray-imaged photoreceptor movements (Fig. 1*C*). Therefore, during the light stimulation, the corresponding photoreceptor RFs - inverted by the ommatidial lenses (11) - scan the visual world with the same two phases but in the opposite directions (Fig. 3*D*).

Remarkably, the global 3D-vector-map of photoreceptors’ photomechanical RF-movement directions (Fig. 3*G*; red and blue arrows; Fig. S25) sweep along a forward flying/walking fly’s optic flow-field (purple arrows), which radiates from a focus at its apparent destination, curving around its left and right eyes. Their difference maps (yellow-matching; black-opposing) are shown for a characteristic upright head-position (Fig. 3*H*) for both the fast- and slower-phase. Generally, the fast-phase is in the flow-field direction and the slower-phase in the opposite direction (Movie S6). But keeping the head upright sets the RFs’ fast- and slower-phases in a balanced mid-state (Fig. 3*I*), where the fast-phase matches the “ground-flow” and the slower-phase the “sky-flow” (Fig. 3*G*). However, locomotion amongst real-world structures (25) would further burstify sampling (11) in a push-pull manner (Fig. 3*F*). Across the eyes, photoreceptors inside each ommatidium would uniquely and orderly ripple between the phases, as incident light-increments drive their RFs fast backward and light-decrement slower forwards, with some moving patterns thus staying longer than others within an RF; which should improve their neural resolvability/detection in time (11). Thus, the fast ventral components may improve resolving complex visual clutter, and the slow dorsal components the landscape and clouds in the skyline. Rotation (yaw) further enhances binocular contrasts (11), with one eye’s fast- and slower phases moving with and against their rotation, respectively, while simultaneously the other eye’s phases do the reverse (Movie S7; Fig. S25 to S27).

Control experiments confirmed the fast microsaccades purely photomechanical (Fig. S15, S20 to S23, and S28 to S36) and similar in both sexes (Fig. S16), reaffirming their phototransduction origin, and validated the X-ray data (**Fig. 2**). Accordingly, the synaptically-decoupled *hdc^JK910^* photoreceptor microsaccades (20) traced the wild-type-trajectories (Fig. S22), set by their matching rhabdomere orientations (Fig. S11 and S12). Moreover, the microsaccades adapted to light contrast changes much like voltage responses (Fig. S23 and S24), with different spectral photoreceptor classes’ microsaccades scaling with their ERGs (Fig. S28 to S30; Table S2 to S5). These results show that microsaccadic sampling along the local small-field motion axes initiates optic-flow processing (26) and suggest that such sampling and locomotion behaviors have jointly evolved to the physical world order to maximize visual information.

### L2-interneurons’ hyperacute motion sensitivity tracks microsaccade directions

To test directly whether the optic-flow-tuned microsaccadic sampling improved acuity of moving stimuli directionally, as suggested experimentally (Fig. 3 *E* to *G*) and predicted theoretically (11), we recorded neural responses of specific LMCs, L2-interneurons (Fig. 4; Fig. S38 to S46), to moving bars and panoramic black-and-white gratings, in which resolution, velocity and direction were changed systematically.

**Fig. 4.**
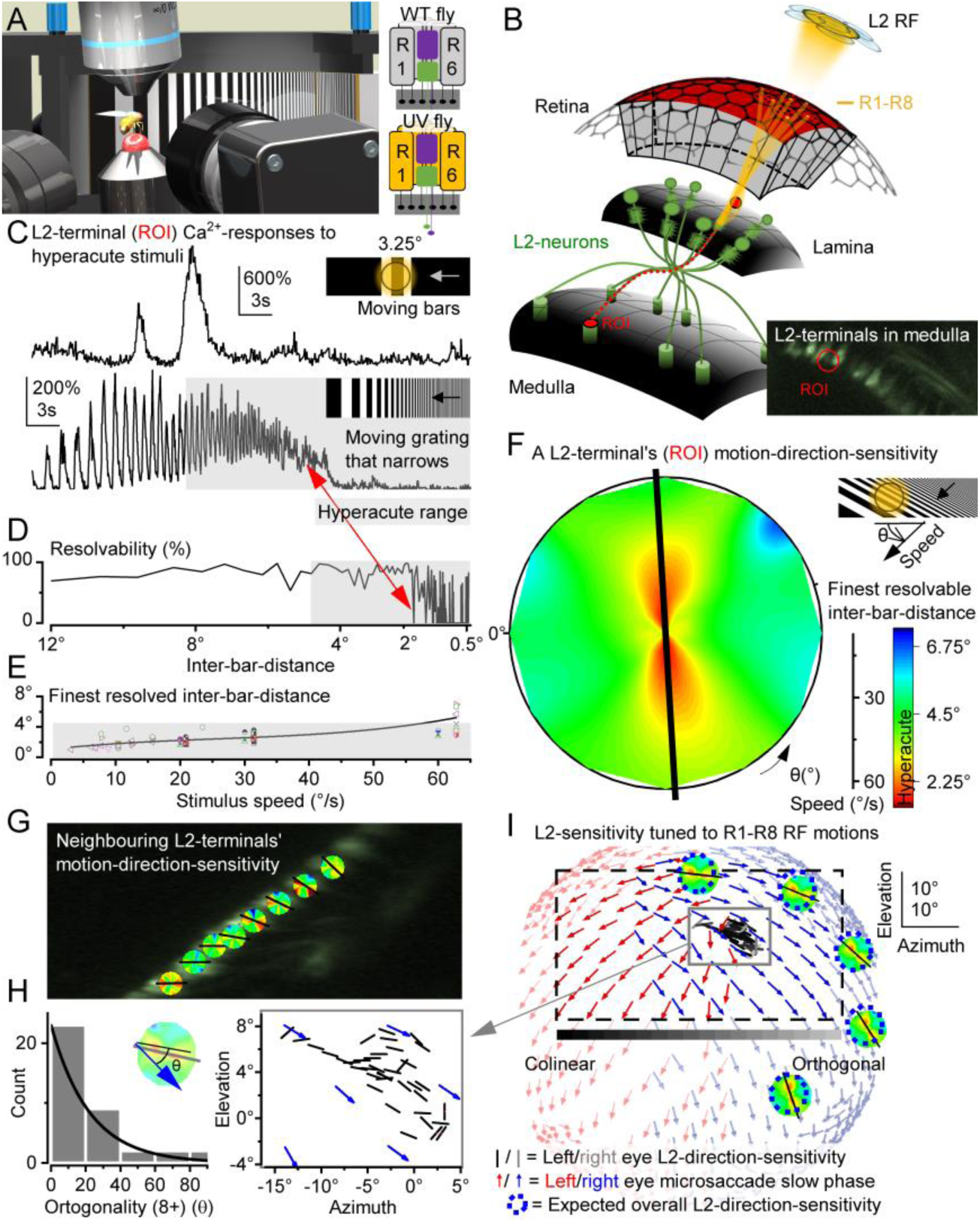
Hyperacute L2-terminal sensitivity follows microsaccade directions. (*A*) A UV-fly saw ultrafine (∼0.5°-pixel-resolution) UV-stimuli on a 150°x50° screen 38 mm away, while its L2-neurons’ GCaMP6f-fluorescence changes (Ca^2+^-responses) were recorded by high-speed 2-photon imaging. In UV-flies (22), UV-sensitive Rh3-opsin is expressed in R1-R6s, containing nonfunctional Rh1-opsin (*ninaE^8^*). (*B*) Each L2 receptive field (RF) samples information from 6 optically superimposed R1-R6 RFs. L2-retinotopy through axonal crossing: distal lamina L2s projects terminals to the frontal medulla. Inset: single L2-terminal Ca^2+^- fluorescence responses to UV-stimulation were analyzed as regions of interest, ROI (red). (*C*) L2-terminal responses resolve in time hyperacute moving bars (here, showing a larger 2^nd^-bar response) and black-and-white gratings (inter-bar-distance <4.5°, grey), crossing their RFs, over a broad range of orientations and velocities. (*D*) L2-resolvability for a dynamically narrowing grating, moving 20.9°/s. Red-arrow indicates the finest resolvable angle (inter-bar-distance; as a rounded-up conservative estimate). (*E*) Inter-bar-distance-resolvability depends on stimulus velocity. L2s’ GCaMP6f-readout resolved hyperacute patterns moving 60°/s. Note, the finest L2-resolvability, ∼1.09°, approaches the visual display’s 2-pixel limit (∼0.5° pixels) and that L2 voltage can encode even faster/finer inputs (22, 33–36). (*F*) An L2-terminal’s motion-direction sensitivity map is broadly hyperacute, here primarily along the vertical axis (the black line shows its fitted orientation-tuning). The map shows the finest resolvable inter-bar-distances to a dynamically narrowing moving grating stimulus (C-E), covering 360° directions at different speeds. (*G*) Neighboring L2- terminals show a gradual shift in their dominant motion-direction sensitivity (black arrows; see Supplement X for analytical details). (*H*) *Drosophila*’s combined L2-terminal motion-direction sensitivity map for the tested left eye region shows retinotopic organization (left, n = 4 flies), mainly co-linear to the corresponding left eye microsaccade directions (right, cf. Fig 3E). (*I*) Eye-location-specific L2-terminal direction-sensitivities map R1-R8 microsaccade directions. Thus, L2-terminals collectively generate a high-resolution neural representation of the moving world, enhancing visual information transfer during forward locomotion. The dotted rectangle specifies the visual area covered by the display screen.

These recordings were primarily done in so-called UV-flies (22), using a bespoke two-photon Ca^2+^-imaging system (Fig. 4 *A* and *B*), while presenting UV-stimuli in an ultra-fine spatiotemporal resolution to a fly walking on a track-ball (Fig. S38 and S39). R1-R6 photoreceptors of UV-flies express only Rh3 (UV-rhodopsin), and therefore see ultraviolet but not green (22), while their L2-neurons express the green-fluorescent Ca^2+^- reporter GCaMP6f. Critically, UV-flies show normal photomechanical microsaccades (Fig. S37) and, as their L2 green-fluorescence Ca^2+^-responses cannot activate the UV-sensitive R1-R6s through orthodromic green-light-transmission (22), they enable naturalistic low-noise conditions for recording high-precision neural signals (Fig. 4 *C* and *D*). Even so, the wild-type-eye L2-GCaMP6f-controls’ Ca^2+^-responses showed consistently similar general dynamics, and thus both results were pooled (Fig. 4*E*).

We found that L2-neurons robustly respond to hyperacute 1-4° moving gratings with location-specific velocity and motion direction sensitivities (Fig. 4 *C* to *E*; Fig. S40 and S41, Movie S8). Thus, by encoding spatial information in time, akin to photoreceptors (4), L2s can transmit finer image details than the compound eye’s optical limit, 4.5° interommatidial angle (12) (Fig. 4*F*; Fig. S41*C*), improving vision. Moreover, the angular maximum of L2 response acuity shifted systematically between neighboring medulla terminals (Fig. 4, *G* to *I*; Fig. S42 and S43), showing that directional motion information from microsaccadic photoreceptor sampling was retained at the medulla input layer. Crucially, the L2-terminals’ motion-sensitivity map was essentially co-linear to the photoreceptor microsaccade direction map (Fig. 4, *H* and *I*; Fig. S44), indicating angular conservation of synaptic information from R1-R6 to L2 (off-channel) LMCs, consistent with preserving the downstream optic flow processing (26). Future experiments need to test whether this is also true for L1 (on-channel) and L3 (27–29) LMCs, as asymmetric microanatomical adaptations (30–32) may further influence local motion computations.

These results demonstrate that L2s collectively convey a high-resolution neural representation of the moving world, maximizing visual information flow (Fig. 3*E*).

### Binocular microsaccades provide hyperacute depth information

By comparing two neural images generated by the left and right eye forward-facing photoreceptors, a fly may extract depth information from the corresponding left and right RF pairs’ (“pixels”) x,y-coordinate differences. This disparity, *d*, is inversely related to the scene depth, *z* (Fig. 5*A*; Movie S9). By applying ray tracing from the ommatidial lenses to the world (Fig. S47 to S61), with parameters taken from their rhabdomere Fourier-transform-beam-propagation simulations (37) and 100-nm-resolution X-ray-imaging (Fig. 1), we first estimated how the corresponding RFs at varying distances from the eyes, and their combined visual field, would look like if the photoreceptors were immobile (Fig. 5*B*).

**Fig. 5.**
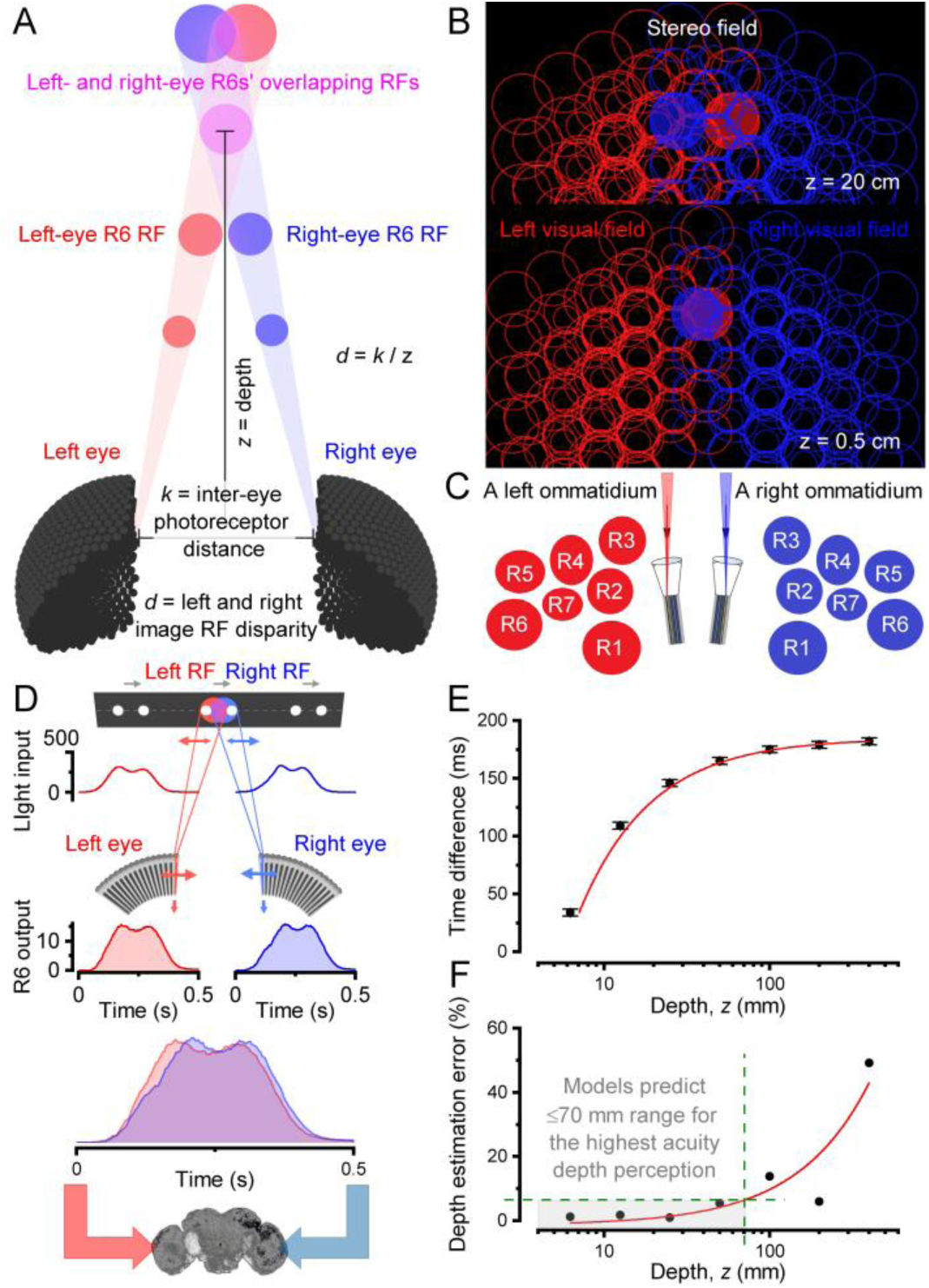
Forward-facing binocular photoreceptors’ biophysically realistic multiscale modeling predicts phasic motion disparity for hyperacute stereopsis. (*A*) With the corresponding left- and right-eye photoreceptors being a fixed distance, *k*, apart, their receptive field (RF) disparities inform about the object depth, *z*. (*B*) R1-R8’s beam-traced (37) RFs (half-width circular cuts of broadly bell-shaped functions; right-eye, blue; left-eye, red) tile the fly’s visual fields over-completely; shown at virtual planes 20 and 0.5 cm depths from the eyes. (*C*) R1-R7/8 rhabdomeres of each paired left and right ommatidia lay mirror-symmetrically (cf. Movie S3 and S9). Because rhabdomeres are of different sizes (11) and distances away from the ommatidium center, so too are their projected RFs (*B*). Therefore, in the neural superposition pooling, the resultant R1-R7/8 RFs do not overlay perfectly into one 4.5°-“pixel” (classic view) but instead tile over-completely each small area in the eyes’ visual fields. (*D*) Phasic voltage response differences of binocularly paired photoreceptors enhance object resolvability in time and carry information about the object depth, z, to the fly brain (see Fig. S55 to S61). Two dots, 3.5° apart moving left-to-right at 50°/s, cross binocular RFs of the corresponding left and right R6s 25 mm away. The resulting mirror-symmetric microsaccades make the RFs move along (right R6) and against (left R6) the passing dots, shaping their light inputs and voltage outputs (Movie S10). (*E*) The proposed binocular mirror-symmetric microsaccadic sampling model (Fig. S59) translates the depth of a moving object into the distance in neural time. The closer the object to the fly’s eyes, the shorter the time difference between the responses. Error bars indicate stochastic jitter. (*F*) The model predicts that *Drosophila* cannot estimate the depth of more distant objects accurately. The error is >10% when an object is >10 cm from the fly eyes, comparable with their distance discrimination estimate (0.2–20 cm (38)).

Static case: the mirror-symmetric sampling array of the paired left and right-eye ommatidia (Fig. 5*C*), in which each R1-R7/8 rhabdomere is a different size (11) and distance (23) from the ommatidium center (Fig. S50 and S54), leads to overlapping RF tiling over the frontal stereo field (Fig. 5*B*; Table S1 and S6). Each eye’s spatial sampling matrix is further densified by the neural superposition signal pooling between seven neighboring ommatidia, in which R1-R7/8s’ RFs of different sizes stack up unevenly (Fig. S57 and S58). This massively overcomplete sampling array greatly differs from the classically considered organization (8, 9), where each ommatidium was considered a sampling point, or a pixel, with a *Drosophila* seeing the world through ∼880 such “pixels”; giving poor spatial resolution with marginal stereopsis. In contrast, our simulations, using the real R1-R7/8 rhabdomere spacing and sizes (Fig. 1 to 3), imply that its left and right eyes’ RF overlap disparity could accentuate frontal resolvability and stereo vision.

But how would the frontal RFs and their neural responses change during photomechanical microsaccades? Furthermore, given that these are left-right mirror-symmetric (Fig. 1 to 4), could their phase differences to rotation be exploited for dynamic triangulation (Fig. 5*A*) to extract depth information in time about the real-world distances and relative positions?

Dynamic case: to simulate how the *Drosophila* left (red) and right (blue) eyes probably see left-to-right moving objects, we set their frontal photoreceptors in their respective model matrixes to contract mirror-symmetrically to light changes (Fig. 5*D*; two left-to-right moving dots) along with the measured dynamics (Fig. 3; Fig. S23 and S55). These caused their respective RFs (red and blue disks) to narrow and slide in and out of each other in opposing directions, phasically shaping their neural responses (Fig. S56 to S61; Movie S10), as calculated by biophysically realistic *Drosophila* photoreceptor models (Fig. S53 to S55) (11, 39, 40). The responses for the left RFs, which moved against the object motion, rose and fell earlier than the responses for the right RFs, which moved along the objects and so had more time to resolve their light changes. Such phase differences in time broadly correspond to the case where similar but not identical images are sequentially presented to each eye, allowing one to perceive the 3D space.

Importantly, R1-R8s’ size-differing, moving, narrowing, and partially overlapping RFs, with stochastic R7/R8 rhodopsin-choices (41) and R1-R6 microstructural/synaptic variations (11, 30), make the retinal sampling matrix stochastically heterogeneous (Fig. S57 and S61). This eliminates spatiotemporal aliasing in early neural images (11). Therefore, theoretically, this dynamic sampling can reliably feed the fly brain with 3D hyperacute information flow. In the centers interlinking the binocular inputs (42), such as the lobula complex (43–45) (Fig. S62), the distance of an object crossing the corresponding left and right eye photoreceptor RFs could then be represented as distance in time (Fig. 5*E*; Fig. S59). To velocity-normalize these distance estimates, their corresponding response waveforms could be correlated with those of their near neighbors (Fig. S59; Movie S10). These results imply that neural motion- and depth-computations innately mix, as they share the same input elements, being consistent with the neurons of the motion detection channels serving vision and behaviors more broadly (42, 46) than just specific reductionist ideals.

### Visual behavior confirms frontal hyperacute stereopsis

To test whether *Drosophila* possesses super-resolution stereo vision, as our theory (Fig. 5; Fig. S59) predicts, we performed visual salience (Fig. S69 to S71; Table S7 to S10) and learning experiments (Fig. S72 to S77; Table S11 to S19) with hyperacute 3D- and 2D-objects in a flight simulator system (Fig. 6). This apparatus was designed so that a tethered fly had no monocular cues to construct 3D representations of the objects neurally, without optically distorting its perception (Fig. S68). In nature, flying insects typically keep an object of interest in frontal view, fixating it by small side-to-side head/body rotations (47, 48). Such movements, by modulating light input and thus mirror-symmetric microsaccades at the binocular eye regions, should accentuate 3D-perception (Fig. 5). But conversely, given the photoreceptor RF dynamics and binocular separation, 3D-perception must diminish with increasing distance, as sampling uncertainties increase, predicting ∼3-70 mm hyperacute stereo-range (Fig. 5*E*; Fig. S59). Therefore, we presented stimuli 25 mm from the eyes, well within this range.

**Fig. 6.**
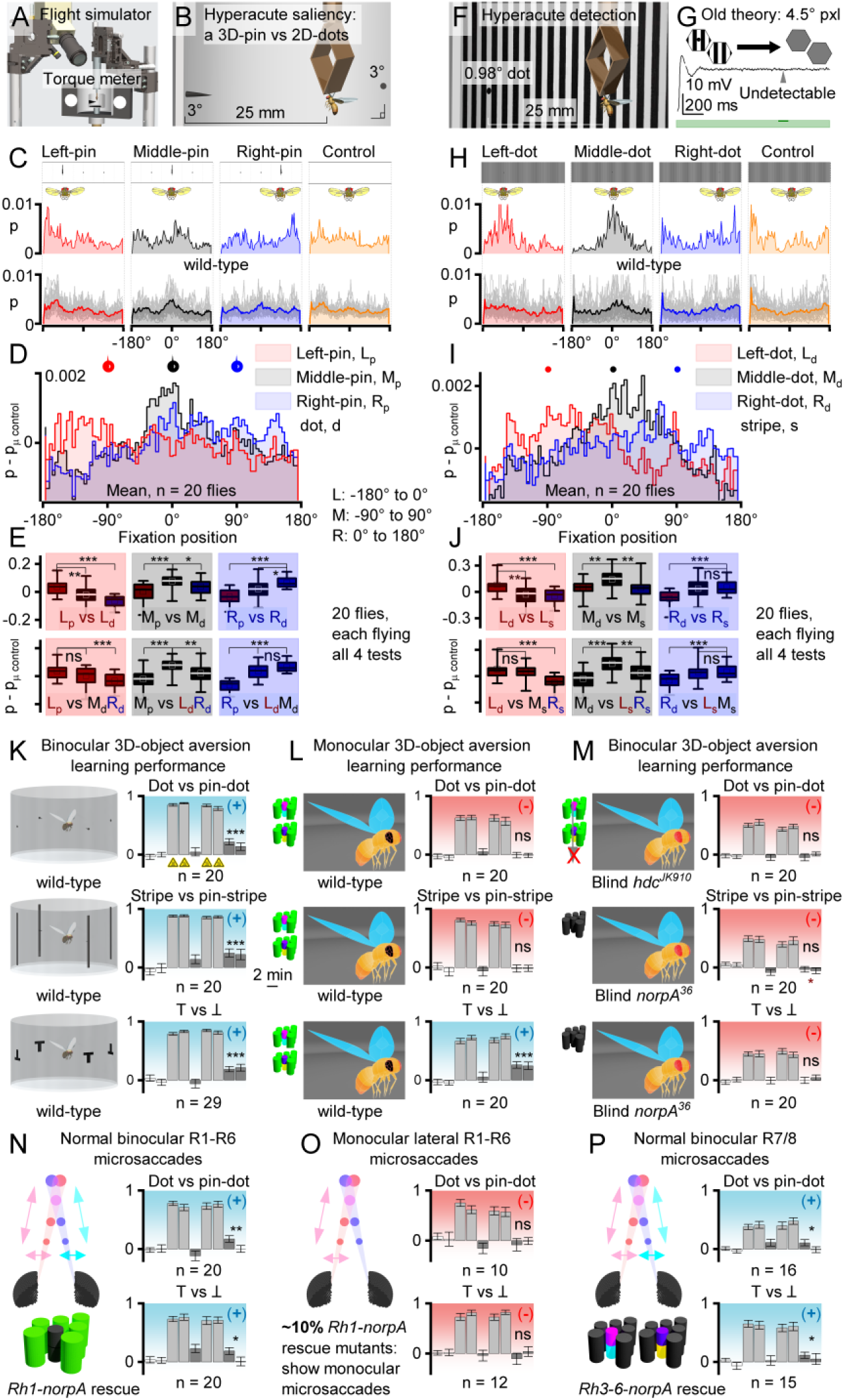
Hyperacute stereopsis requires two eyes with mirror-symmetric microsaccades. (*A* and *B*) In a flight simulator, a torque-meter-tethered flying *Drosophila* controls how a white cup rotates around it, showing three black dots (3.9° Ø), 90° apart, one with a black 4-mm-center-pin (1° Ø). Axially 25 mm away, the 3D-pin at -90° (left), 0° (middle), or 90° (right) dot is monocularly indistinguishable from the 2D-dots. (*C*) Yet, flies fixate more on the pins than on the competing dots, implying 3D-pin salience. A single fly’s (above) and population (below) frontal fixation probability to the left, middle and right pin/dot-positions, and during a blank-control. (*D*) Fixation probabilities for the 3 pin-positions; blank-control subtracted to minimize experimental bias. (*E*) Positional salience (e.g. left-pin vs left-dots, above) and competition (e.g. left-pin vs middle- and right-dots, below) statistics indicate that *Drosophila* see hyperacute 3D-pins amongst 2D-dots (super-resolution stereopsis). (*F*) Testing hyperacute 2D-object detection. (*G*) Old theory simulation: a fly with static 2D-vision and 4.5° ommatidial pixelation cannot detect a black 0.98° dot hidden amongst 1.2° stripes, as its optically-corrected contrast difference over a photoreceptor’s RF (5.4° half-width) is only ∼1.6% of that of the stripes alone, evoking response differences < voltage variation (noise) for such a contrast pulse (green, 100 ms). (*H*) Nevertheless, flies fixate on the hidden dot, irrespective of its position. A single fly’s (above) and population (below) frontal fixation probability to the left, middle and right dot positions, and during a stripe-control. (*I*) Fixation probabilities for the 3 dot-positions; stripe-control subtracted to minimize bias. (*J*) Positional detection (e.g., left-dot vs. left-stripes, above) and salience (e.g., left-dot vs. middle- and right-stripes, below) statistics/trends indicate that *Drosophila* find hyperacute dots visually interesting. (*K*) *Drosophila* learns to avoid hyperacute 3D-pins or 2D-lines/dots (above and middle), associated with IR-heat punishment (training, triangles), equally well to the classic T vs. Ʇ conditioning (below). (*L*) One-eye-painted *Drosophila* fails to learn hyperacute 3D- and 2D-object avoidance (above and middle), demonstrating that super-resolution stereo vision requires two eyes. Yet monocular *Drosophila* shows normal T vs. Ʇ conditioning (below), indicating that one eye is enough to learn large 2D-patterns, consistent with retinal-position-invariant pattern recognition (49). (*M*) Blind *hdc^JK910^* (above) and *norpA^36^* (middle and below), with no synaptic photoreceptor outputs but normal audition and olfaction, failed to learn the test stimuli, validating that the wild-type learning (K and L) was visual. (*N*) Rh1-rescue *norpA^36^* with functional R1-R6 photoreceptors and normal mirror-symmetric left and right eye microsaccades learned hyperacute 3D-stimuli (above) and large 2D-patterns (below), but less well than wild-type. Thus, R7/R8s also contribute to stereopsis. (*O*) ∼10% of Rh1-rescue *norpA^36^* flies showed only left or right eye lateral microsaccade components, leading to asymmetric and asynchronous binocular sampling. These flies neither learned hyperacute 3D-stimuli (above) nor large 2D-patterns (below). Meaning, mirror-symmetric microsaccadic sampling is necessary for hyperacute stereopsis. (*P*) *norpA^P24^* with rescued R7/R8- photoreceptors, showing normal microsaccades, learned to differentiate both coarse 2D- and hyperacute 3D-patterns. Thus, R7/R8s alone are sufficient for hyperacute stereopsis. Note, the exact microsaccadic movements during the experiments are unknown as it was too difficult to measure the DPP movement concomitantly.

In salience experiments, a tethered flying fly explored a white panoramic scene, which had a small (4-mm- long) black hyperacute (*i.e.*, <4.5° interommatidial pixelation (12)) 3D-pin, protruding from a small black dot (3.9° Ø), and two similar-sized black 2D-dots, each 90° apart (Fig. 6, *A* to *C*). The pin-position was varied for three trials, and the fourth (control) was a blank scene, presented in random order. For each trial, we measured a fly’s fixation behavior: how much time it kept each part of the scene at the fontal view, given as probability. The conventional compound eye acuity theory (8, 9) states that because all these three objects had the same contrast and were smaller than the eyes’ interommatidial pixelation, their differences would be invisible, giving them equal salience, and *Drosophila* should fixate all three of them with equal probability. Whereas, our *mirror-symmetric microsaccadic sampling theory* (Fig. 5; Fig. S59) predicts that for a fly with hyperacute 3D-vision, the 3D-pin would appear different from the 2D-dots, with its saliency increasing fixations. In supporting our theory, the results showed that *Drosophila* prefers to fixate hyperacute 3D-pins, irrespective of their positioning (Fig. 6, *C* to *E*). Equally, in separate experiments, the flies readily fixated on hyperacute 2D-dots (0.98°) hidden in a 1.0° hyperacute stripe-scene (Fig. 6, F to J), which by the con theory would be impossible (Fig. 6*G*). Moreover, the flies’ hyperacute optomotor responses (Fig. S63 to S65) followed the predictions of our theory (Fig. S66 and S67).

In learning experiments (Fig. 6*K*), *Drosophila* saw both hyperacute 2D-objects (black bars, above, or dots, middle) and hyperacute 3D-objects (black pins inside bars or dots) and were taught by associative heat punishment (Fig. S73) to avoid one or the other stimulus. Again, in support of our theory, the flies readily learned to avoid the punishment-associated stimulus, validating that they saw hyperacute 3D-objects different from their 2D-counterparts (of the same area/contrast). This learning was robust, matching the classic large-pattern T vs. Ʇ performance (49) (below). But importantly, it was abolished when either the left or the right eye was painted black (Fig. 6*L*, above and middle), indicating that hyperacute 3D-vision requires inputs from both eyes. In contrast, the large-pattern T- vs. Ʇ-learning still occurred with one eye only (below), consistent with the reported retinal-position-invariance in visual pattern recognition (49). Whereas, blind *hdc^JK910^* (Fig. 6*M*, above), *norpA^36^* (middle) and *norpA* (below) mutants, having no synaptic photoreceptor outputs but functioning auditory and olfactory senses, failed to learn the test stimuli, corroborating that wild-type *Drosophila* see the nearby world and learn its objects in hyperacute stereo.

Finally, we tested whether learning hyperacute 3D-stimuli requires either R1-R6 or R7/R8 photoreceptors or both with intact microsaccadic sampling. Here, we exploited our serendipitous finding that rescuing R1-R6 or R7/R8 photoreceptors in blind *norpA^P24^*-mutants make their microsaccades’ lateral component more fragile to mechanical stress or developmentally imperfect, with not every tethered fly showing them (Fig. S74). Therefore, after the learning experiments, we recorded each fly’s light-induced deep pseudopupil movement (Fig. 3) and ERG, quantifying their microsaccades and phototransduction function, respectively. We found that whilst most *norpA^P24^* Rh1-rescue flies (R1-R6s are sampling, R7/R8s not) showed normal binocular microsaccades (Fig. 6*N*), ∼10% showed microsaccades only monocularly (Fig. 6*O*). Importantly, however, each fly eye (both left and right) showed a characteristic ERG, indicating that its phototransduction, and thus axial microsaccade movement from PIP_2_ cleavage (10, 11) was unspoiled. The flies with normal lateral microsaccades (Fig. 6*N*) learned the difference between hyperacute pins and dots (above) and large T- vs. Ʇ-patterns (below), but less well than wild-type flies (Fig. 6*K*), establishing that R1- R6 input is sufficient for hyperacute stereo vision but that R7/R8s must also contribute. Conversely, the flies that showed monocular lateral microsaccades (Fig. 6*O*) neither learned hyperacute 3D objects (above) nor large 2D-patterns (below), indicating that misaligned binocular sampling corrupts 3D-perception and learning. Whereas R7/R8 rescued *norpA^P24^*- (Fig. 6*P*) and *ninaE^8^*-mutants confirmed that the inner photoreceptors also contribute to hyperacute stereopsis.

These findings concur with our simulation results, which predicted that asynchronous binocular sampling should break stereopsis (Fig. S60). Collectively, these results demonstrate that binocular mirror-symmetric microsaccadic sampling is necessary for super-resolution stereo vision and that both R1-R6 and R7/R8 photoreceptor classes contribute to it.

## Discussion

We showed how the *Drosophila* compound eyes’ binocular mirror-symmetric photoreceptor microsaccades (Fig. 1 to 3) generate phasic disparity signals in much finer resolution than ommatidial pixelation, suggested by their interommatidial angle (Fig. 4 and 5). The fly brain could use these signals to triangulate object distance to a neural distance signal in time (Fig. 5), enabling stereopsis (Fig. 6). We also revealed how the microsaccades across the eyes track a flying fly’s optic flow field to enhance information from the world in motion (Fig. 3 and 4). Visual behavior matched the modeling predictions (Fig. 5 and 6), demonstrating that the neural image generated by mirror-symmetric microsaccadic sampling must result in a higher quality perceptual representation of the stimulus as compared to the neural image generated by immobile photoreceptors (8, 9), or asymmetric or asynchronous binocular sampling (Fig. S60). By integrating *in vivo* assays from subcellular to whole animal 3D-perception with multiscale modeling from adaptive optics to depth computations (Fig. S49 to S61), these results establish a new morphodynamic light information sampling and processing theory for compound eyes, to better understand insect vision and behaviors (11, 50). To further demonstrate its explanatory power, we also verified its predictions of *Drosophila* seeing nearby objects in higher resolution (Fig. S66) and “optomotor behavior reversal” (51) not resulting from spatial aliasing (Fig. S67).

It has been long thought that because the eye and head movements are dominated by axial rotation, they should provide little distance information as objects, near and far, would move across the retina with the same speed (52). In contrast, our study highlights how the visual systems can use microsaccades, and eye/head rotations, to both contrast-enhance (Fig. S26; Movie S7) and extract depth information (Movie S10). Rapid mirror-symmetric inward-rotating photomechanical photoreceptor microsaccades in the left and right eyes cause phase-difference signals, which inform the *Drosophila* brain in time how far an object is from its eyes. But when the world is still, a fly can further contract its intraocular muscles (13), rotate or move its head from side-to-side, as insects with compound eyes commonly do during fixation, to generate both binocular and motion parallax (53) signals to resolve object depth.

With mirror-symmetric microsaccadic sampling, flies and possibly other insects with binocular compound eyes can have an intrinsic sense of size. For two objects with equal angular size and velocity as projected on the eyes, the closer one, and thus physically smaller (a mate), generates a brief and precise binocular disparity in time. While the other object, further away and thus bigger (a predator), generates longer-lasting but more blurred disparity.

This encoding strategy applies to machine vision. Super-resolution depth-information about a nearby object (moving or still) can be extracted in time, for example, by piezo-resonating synchronously and mirror-symmetrically two horizontally separated sampling matrixes (left and right) with overlapping views and then correlating their phasic differences for each corresponding pixel; equating to a two-matrix extension of the VODKA sensor principle (54)). In more sophisticated optic flow-optimized 3D systems, binocular photomechanical pixel-sensors could move along their specific concentric rotation axes as in the *Drosophila* eyes.

We note that recent work has shown that human cones (55) and vertebrate rod-photoreceptors (56) contract photomechanically, comparable to *Drosophila* photoreceptor microsaccades (10, 11). It will be interesting to see whether these microsaccades increase visual acuity and participate in stereo vision and whether high-intensity X-rays also activate them (15–17).

## Materials and methods

The multiscale experimental and theoretical approaches used in this study are explained in detail in the SI Appendix, organized in Sections (I-VIII).

## Acknowledgments

We thank T. Salditt, B. Hartmann, and M. Sprung for help and support with DESY experiments; E. Chiappe, R. Strauss, B. Brembs, A. Zelhof, and E. Buchner for flies; E. Chiappe for help with 2-photon-imaging preparation optimization; M. Reiser for sharing unpublished *Drosophila* eye structural data; T. Salditt, G. de Polavieja, G. Belušič, A. Straw, L. Fenk, A. Nikolaev, and A. Lin for discussions.

## Funding

This work was supported by Jane and Aatos Erkko Foundation Fellowships (M.J. and J.T.), The Leverhulme Trust (RPG-2012–567: M.J.), the Biotechnology and Biological Sciences Research Council (BBSRC: BB/F012071/1, BB/D001900/1, and BB/H013849/1: M.J.), the Engineering and Physical Sciences Research Council (EP/P006094/1: M.J.), the White Rose BBSRC Doctoral Training Program (BB/M011151/1: M.J. and B.S.), the Open Research Fund of the State Key Laboratory of Cognitive Neuroscience and Learning (M.J.), High-End Foreign Expert Grant by Chinese Government (GDT20151100004: M.J.), DESY-synchrotron (I-20190808 EC, I-20180674 EC and I-20170823: M.J. and R.M.) and ESRF-synchrotron (LS-2780: M.J. and R.M.) beam-time grants.

## Supplementary Information

### Materials and methods

#### I Measuring X-ray-induced global photoreceptor movements and ERG

**Overview**

This section describes ESRF and DESY synchrotron experiments to measure the *Drosophila* eyes’ global photomechanical photoreceptor movements (synchronous left and right eye microsaccades) to high-brilliance X-ray stimuli with simultaneous electrophysiological (electroretinogram, ERG) responses. It gives central background information and additional supporting evidence for the results presented in the main paper, including:

- X-rays activate phototransduction similar to visible light, causing photoreceptors to contract photomechanically while generating a normal electrical response.
- The left and right eye microsaccades are mirror-symmetric.
- Microsaccades are photomechanical – independent of intraocular muscle activity.

##### I.1 *In vivo Drosophila* preparation

Under a stereomicroscope, 3-4 day old (12∶12 light-dark-cycle reared) *Drosophila* were gently attached inside a size-adjusted pipette tip by puffing air so that their head and upper thorax protruded from its small end. Using a low melting point (60-64 °C) beeswax (Fig. S1 *A* and *B*), a fly was swiftly fixed to the tip end without touching its eyes and leaving the abdomen intact for respiration. Next, its head was waxed to the thorax, and the proboscis was stretched and waxed to the pipette wall to minimize muscle-induced head and vergence eye movements. We took special care not accidentally dent the eyes during the preparation, as this can damage the photoreceptor microsaccades’ sideways component (see Section II.4., below). In some preparations, such as the one shown in Fig. S1B, we also fixed the antennae with a beeswax blob to minimize muscle activity. This procedure did not change the experimental results.

The large end of the pipette tip was super-glued on a standard preparation holder metal pin (Fig. S1A, inset). The fly was then transported to the X-ray beamline’s tomographic rotation stage and connected from the pin in a desired orientation and position -for either one or two eye imaging (Fig. S1 *C* and *D*). Once the fly was aligned correctly for the X-ray imaging/stimulation experiments with the selected magnification, we took a photograph of its eyes for the records.

**Fig. S1.**
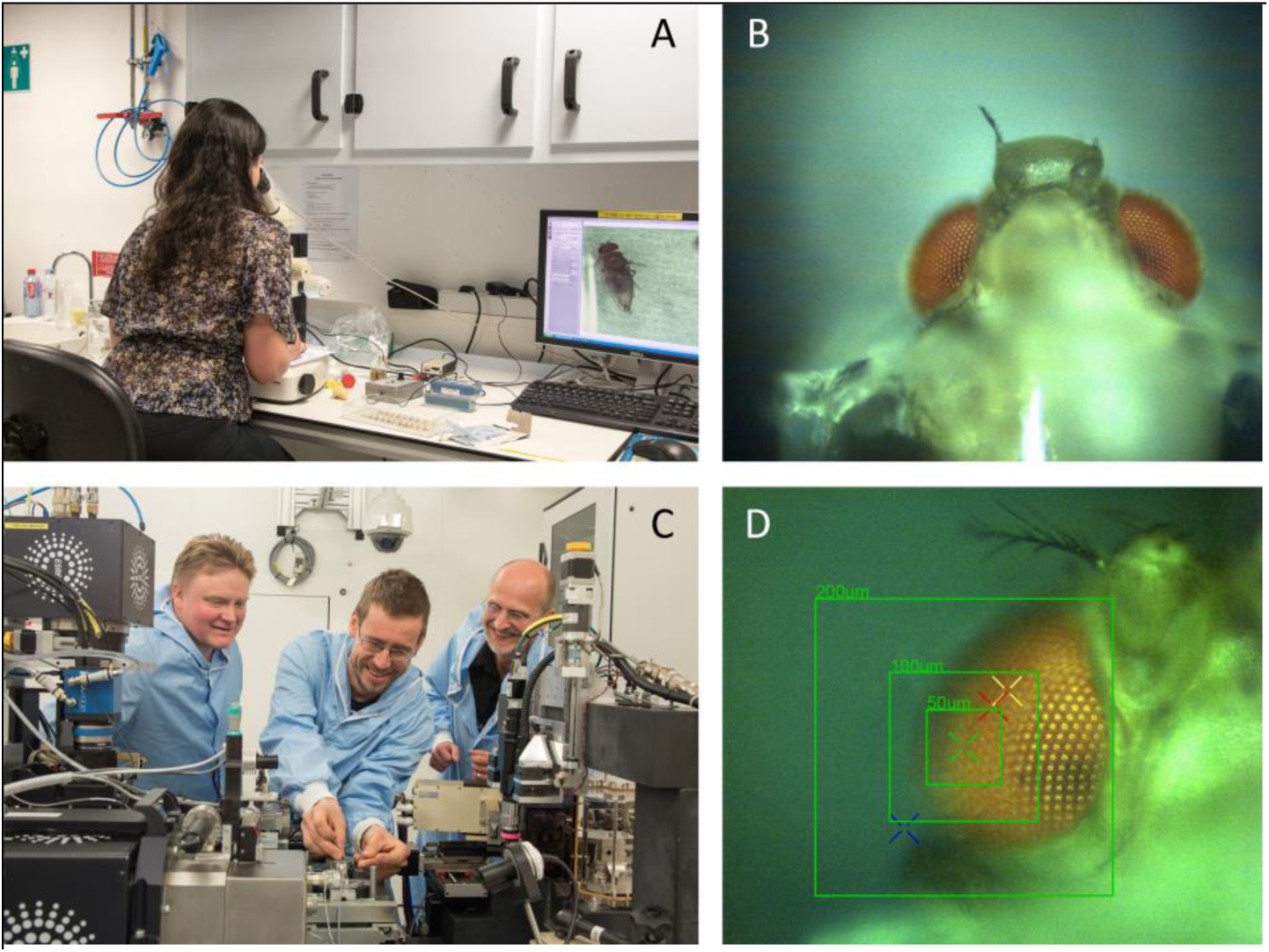
Preparing *Drosophila* for *in vivo* X-ray imaging experiments. (*A*) Flies were freshly prepared under a stereomicroscope by fixing them inside a pipette tip while avoiding any physical stress to their eyes at the ESRF or DESY biological sample preparation room, to be ready only minutes before the experiments. (*B*) Each *Drosophila* was fastened to a pipette tip by beeswax. Waxing the mouthparts and back of the head to the tip-rim minimized any head muscle including movements, those by intraocular muscles, while leaving the eyes intact and unobscured. Inside the pipette, the fly could move its legs and abdomen, with spiracles free for respiration. In some preparations, as shown here, the antennae were also wax-immobilized. Inset (between *A* and *B*): the pipette tip was super-glued to a metallic connector bin and transported to the beamline. (*C*) *Drosophila* preparation was clamped from its connector pin to the tomographic rotation-stage; here shown at ESRF ID16b beamline. (*D*) At the radiation-protected observation hut, the fly’s orientation and positioning were remotely set for X-ray imaging using the live video feed from the beamline cameras. Each fly head was photographed at its imaging position for the records.

##### I.2 *In vivo* X-ray imaging

In the initial ESRF beamline experiments (Fig. S2*A*), we generated X-ray pulses of pre-set intensities and durations (typically 100-300 ms) to record photomechanical photoreceptor microsaccades (100 frames/s) to a 10 ms bright white LED flash, as synchronized by TTL-pulses. The high-intensity LED was positioned ∼5 cm above the fly head to generate locally - in the upper-section of the eye - photomechanical photoreceptor contractions. Their speed, size, and direction would be then revealed by high-resolution (200 nm pixel) X-ray imaging. However, surprisingly, we found X-rays themselves made all the photoreceptors in the two eyes rapidly contract mirror-symmetrically in synchrony (Fig. S2*B*). The size and speed of these contractions directly depended on them upon the X-ray intensity. Meaning, the white LED flash was not needed to activate photoreceptor contractions, as X-ray seemed to activate them directly. Moreover, during X-ray imaging, the beamline lights were either on or off, but this had little or no effect on the photoreceptor contraction amplitudes. This observation is consistent with the findings that photomechanical photoreceptor microsaccades occur equally well in the dark- and light-adapted eyes (11) (see Section II.6., below).

**Fig. S2.**
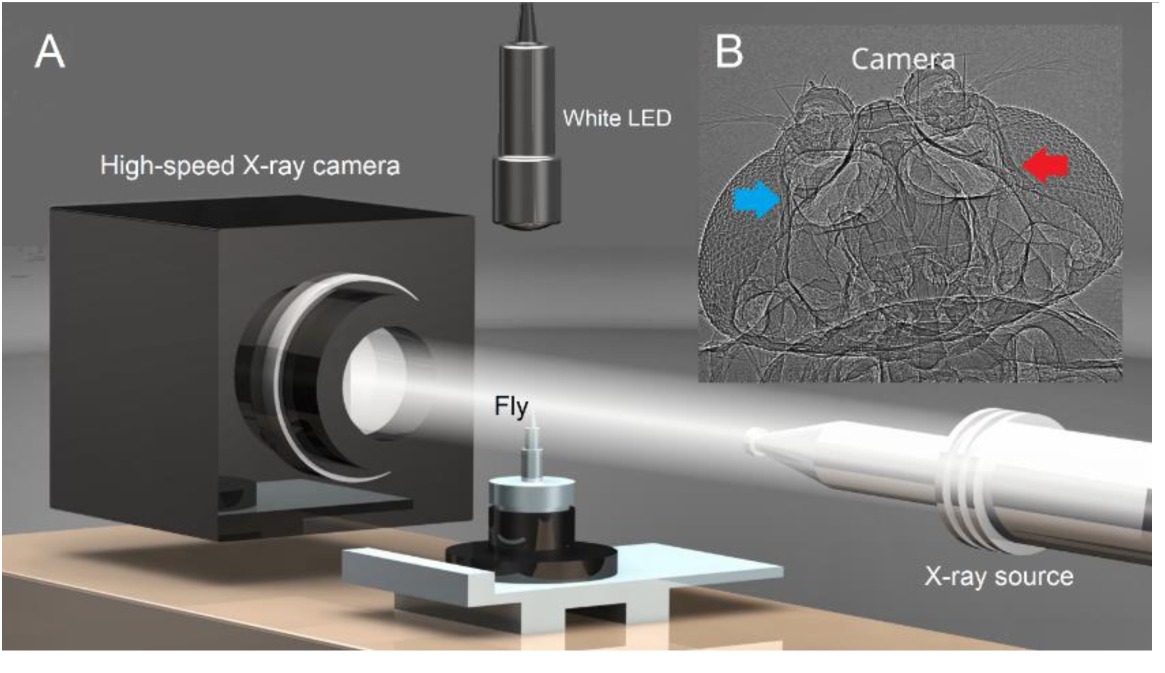
Schematic of the initial X-ray imaging configuration. (*A*) White LED light stimulation was not needed for imaging photomechanical photoreceptor contractions because brief high-intensity X-ray pulses, used for the *Drosophila* eyes, also simultaneously activated photoreceptors photomechanically. Dead control flies (killed by freezing and thawing), which displayed structurally intact eyes, never showed X-ray-induced photoreceptor contractions. (*B*) *In vivo Drosophila* head high-speed X-ray video reveals its internal structure with global photoreceptor contraction dynamics. Ommatidial lenses are on the eye surfaces and underneath them the radially arranged string-like photoreceptors. X-ray-activation made photoreceptors in the right (blue arrow) and left eyes (red) contract rapidly and mirror-symmetrically in the back-to-front direction.

During a typical test protocol that consisted of six 300-ms-long intensifying X-ray pulses, the flies remained alive as we often saw spontaneous antennae movements, which made us fix the antennae with beeswax in some later preparations (Fig. S1*B*). After the experiments, we checked that the flies were still alive by observing their leg movements inside the pipette tip. Sometimes, we even let a fly out of the pipette tip to see it walk. Because the photoreceptor contractions to a given X-ray pulse (i) could be reliably repeated without extensive changes in their dynamics (Fig. S3), (ii) these dynamics (their speed and size) were intensity-dependent. Moreover, (iii) these dynamics matched those of the visible-light-induced photoreceptor microsaccades, first measured within a single ommatidium (11), and in the current study, across the eyes (see Section II., below). Therefore, it seemed plausible that X-rays were directly activating phototransduction. Besides, if the photoreceptor microsaccades were a part of intraocular-muscle-induced retinal movements - driven by clock-spikes (57), fast gaze-stabilization reflexes, or visuomotor feedbacks (13, 58) -, we would not expect them to show adaptive intensity-dependent dynamics but instead be of similar size and speed at all tested X-ray-intensities. Such dynamics we never saw.

Detailed top speed and total displacement depth profiles by cross-correlation analysis show that photoreceptors’ proximal ends near the basement membrane moved more vigorously than their distal ends during the X-ray pulses, while the lenses remained still (Fig. S3*E*). However, the cells deeper in the brain likely move more than indicated here. We suspect this because (i) the brain processes appear utterly transparent in the X-ray images (possibly due to their size, organization, and X-ray optical properties), and (ii) contracting receptors could be seen pulling the whole basement membrane while contracting.

We also calculated similar speed and displacement profiles along the top-bottom axis, showing that the photoreceptors near the eye’s medial edges moved the most (Fig.S3*F*). Since the medial photoreceptors have binocular overlap (see Section II.1.ii., below) and participate in the proposed dynamic stereo vision (see Section V.10., below), this specialization may provide better depth perception. For example, shifting the receptive fields fast over a larger area than the more lateral-inferior receptors. Interestingly, while the top-bottom displacement profile shows a somewhat monotonically decreasing trend, the speed profile has a visually distinct bump between 20° and 60° rotations from the top. This bump is possibly a specialization to the optic flow a fly experiences during its forward locomotion. Visual objects appear to move in general fastest during forward locomotion when located perpendicularly to the locomoted direction.

In Fig. S3, we show one of the most successful experiments of the granted beamtimes. The slight variations in the rotation of the fly with respect to the X-ray beam and (ii) the head’s tilt caused the photoreceptor contractions to occur more out-of-plane in some specimens than in others, almost as if twisting. Moreover, the increasing angle between the camera-image and the microsaccade planes decreased the observed motion sinusoidally.

**Fig. S3.**
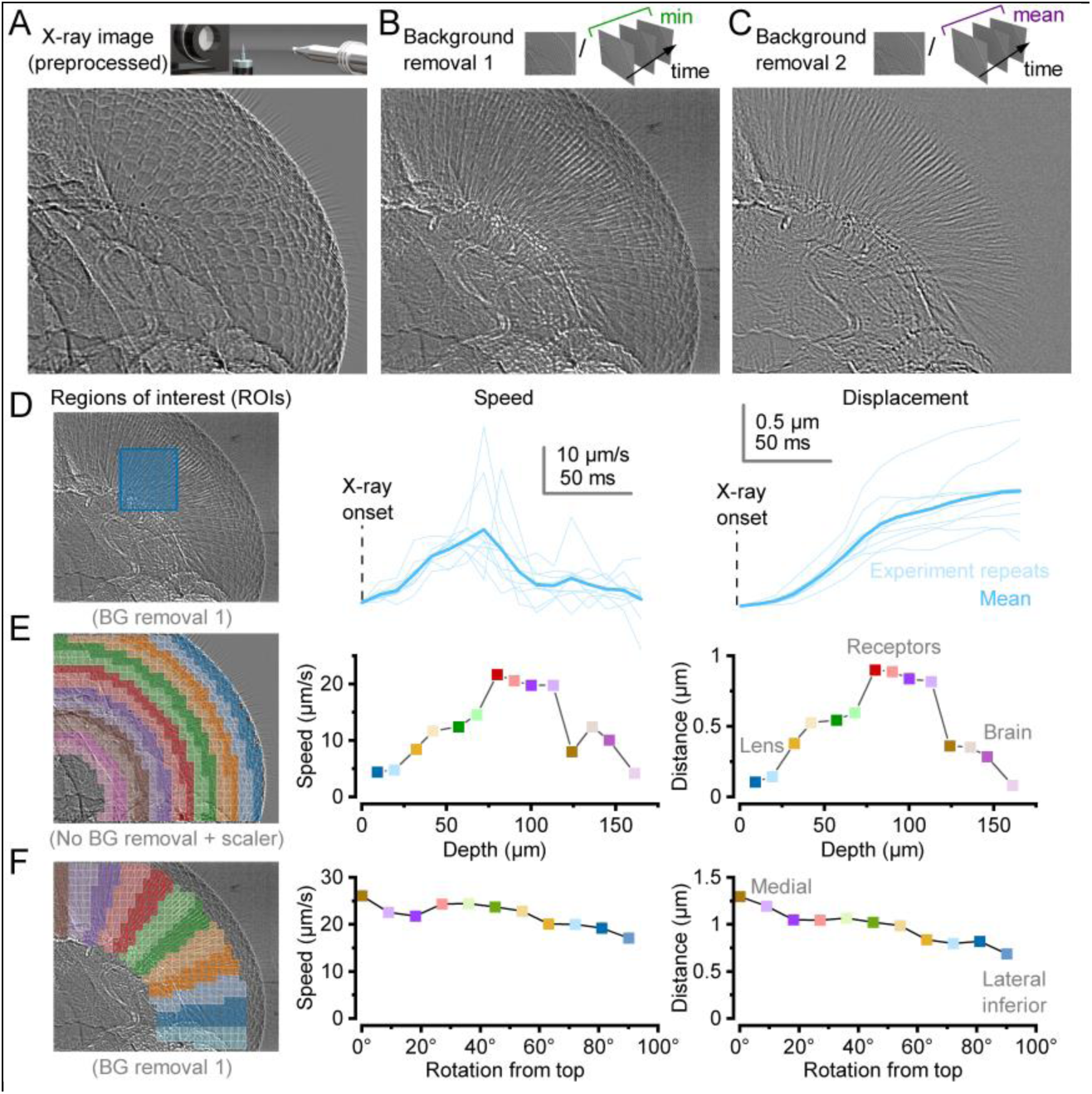
Photomechanical photoreceptor contractions to bright 200 ms X-ray pulses were repeatable for many minutes. (*A*) A preprocessed X-ray image of a wild-type *Drosophila* eye. (*B*) Partial background subtraction using the recording minimum frame division enhances moving features during the X-ray pulse. (*C*) Full background subtraction using the mean frame division preserves only the moving features. (*D*) Localized motion analysis shows speed and displacement kinematics comparable to photoreceptor voltage responses when ∼30,000 microvilli (refractory phototransduction units) repeatedly sample bright (high-photon-count) pulse stimulation (11, 39, 40, 59). Each repeat here is followed by 2 s of darkness. (Movie S1) (*E*) Total displacement and top speed profiles in the tissue depth suggest that the photoreceptor layer, especially photoreceptors’ proximal ends, move the most together with the basement membrane. (*F*) Radial or top-bottom total displacement and top speed profiles indicate that the frontal photoreceptors, which are the longest and contain more microvilli (12), move the most. Such larger movements may be a beneficial adaptation for the proposed dynamic depth estimation.

To further test that the photoreceptor movements during X-ray imaging were not caused by heat-induced tissue shrinkage or expansion, we freshly killed some flies by placing them in a freezer for >30 min and repeated the recordings. None of the freshly killed flies showed photoreceptor contractions or other intra-cutaneous movements, although these were seen when the flies were alive, suggesting that direct X-ray phototransduction activation caused the photoreceptor contractions.

##### I.3 ERG-recording at X-ray beamlines

To test whether (i) X-rays activate photoreceptors and (ii) photoreceptors contract photomechanically, we combined *in vivo* X-ray source imaging with electrophysiology. The wild-type and blind mutant eyes’ global electrical responses to high-brilliance X-ray pulses were recorded using the conventional electroretinogram (ERG) method with extracellular microelectrodes (19, 20, 57) (Fig. S4).

**Fig. S4.**
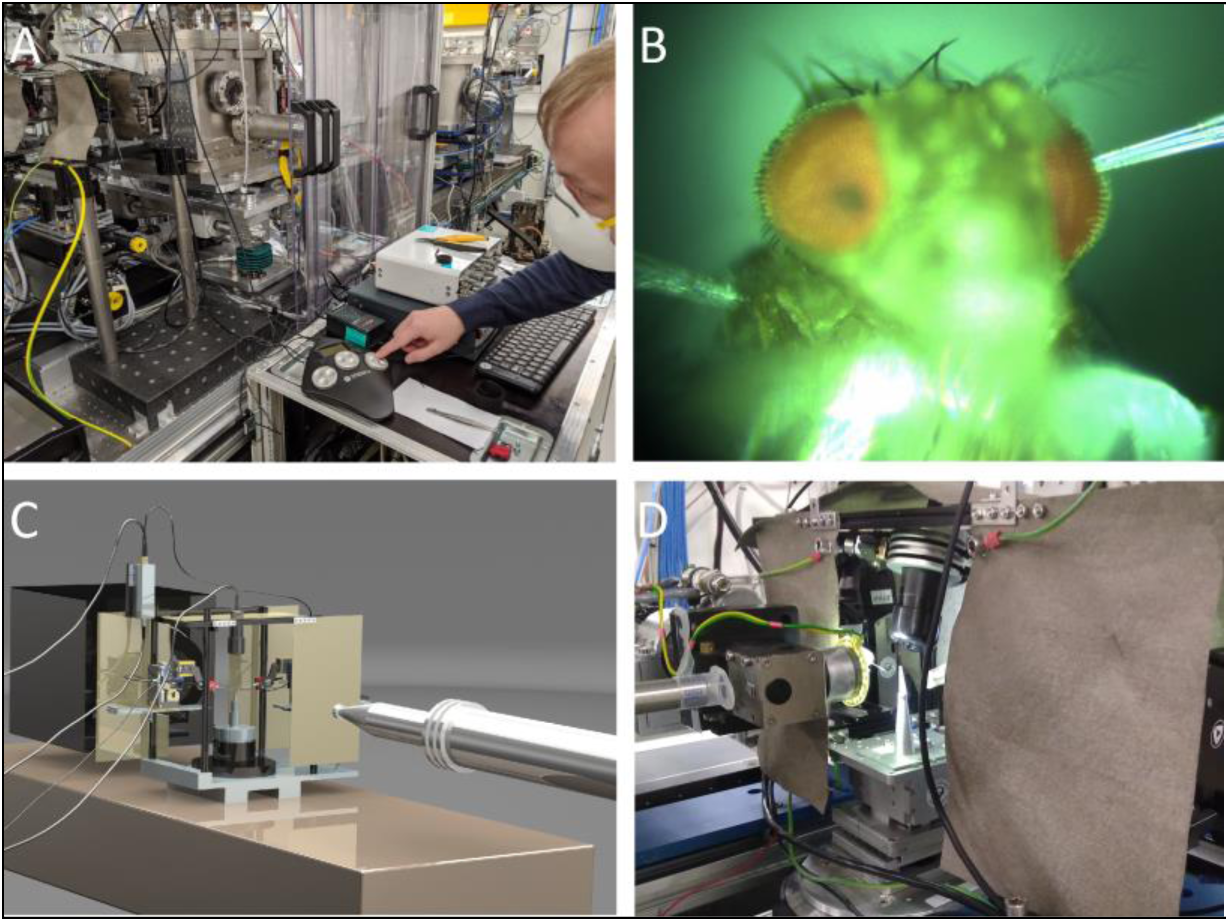
X-ray imaging with simultaneous electrophysiology. (*A*) The recording and reference microelectrodes were gently positioned to touch the fly using two remote-controlled piezo-micromanipulators. (*B*) The recording and reference microelectrodes touched on one eye’s corneal surface and the torso, respectively. (*C*) Schematic of the X-ray imaging configuration with simultaneous ERG recording. (*D*) X-ray imaging and electrophysiology configuration at GINIX endstation at P10 beamline, DESY.

A fly was affixed by beeswax to a size-adjusted pipette tip to ensure its head remained stationary (see Section.I.1., above). The pipette was super-glued on a standard preparation holder pin, used to transport and connect the fly – in a desired orientation and position - to the X-ray tomographic rotation stage. Blunt (low-resistance) filamented borosilicate glass capillary microelectrodes (0.7 mm inner and 1.0 mm outer diameters) filled with fly Ringer (containing in mM: 120 NaCl, 5 KCl, 10 TES, 1.5 CaCl_2_, 4 MgCl_2_, and 30 sucrose) were attached to electrode holders (containing a chloridized silver wire) and connected to a microelectrode amplifier (EXT-02 B; npi Electronic, Germany) (Fig. S4*A*). We carefully positioned the electrodes with two remote-controlled piezo-micromanipulators (uMp, Sensapex, Finland) while getting continuous visual feedback from the live video stream and electrophysiological laptop-computer display (Biosyst-software (11, 60)). The recording electrode was placed to touch one eye’s corneal surface and the reference electrode the fly’s torso (Fig. S4*B*). The electrode positioning was further helped by the microelectrode amplifier’s simultaneous auditory feedback, in which pitch-change signaled the closing of the circuit when both the electrodes touched the fly.

**Fig. S5.**
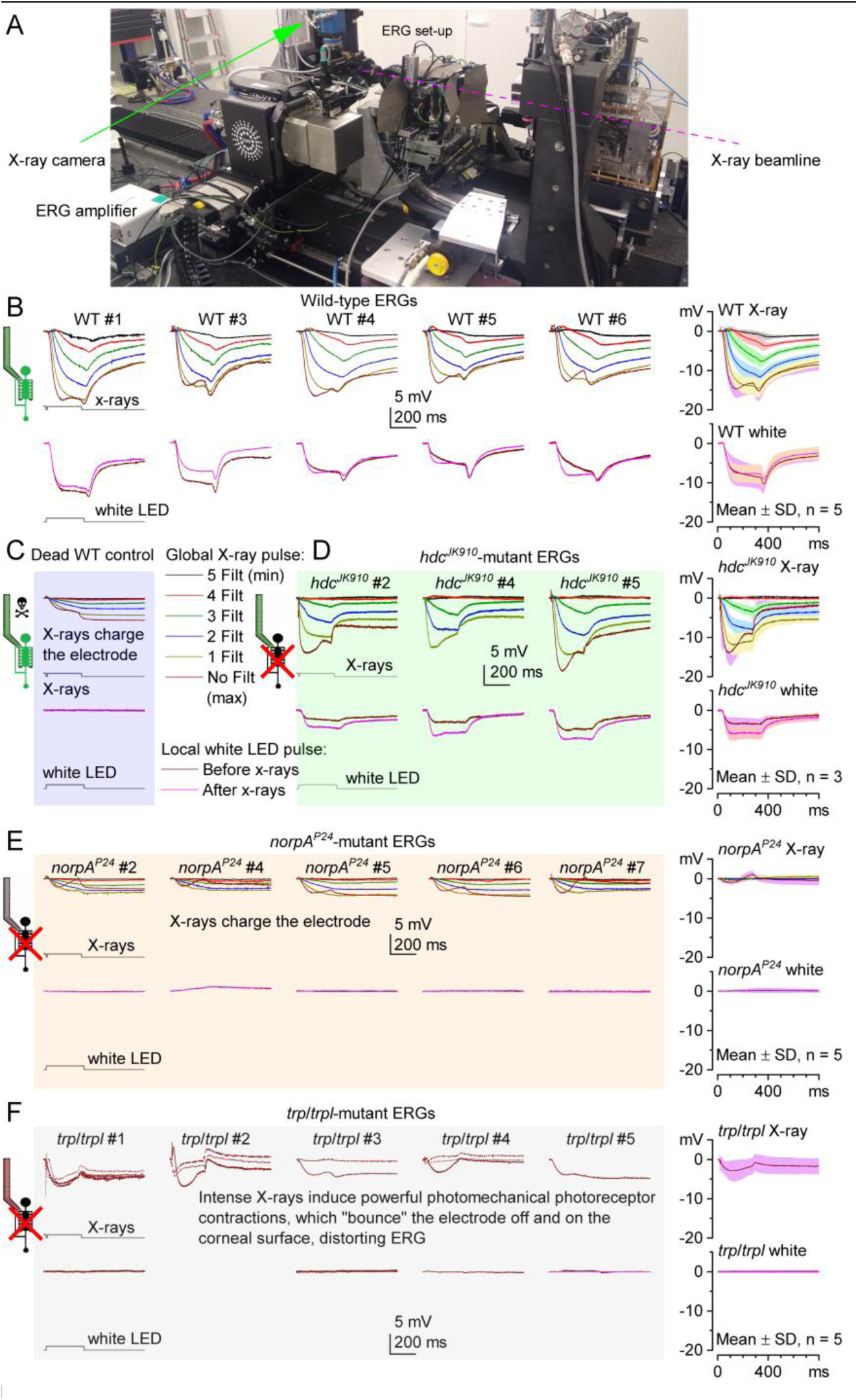
*in vivo Drosophila* ERG responses to X-ray pulses show normal visible-light-like phototransduction and synaptic transmission. (*A*) Remote-controlled portable ERG recording system, shown here as set in ESRF beamline ID16b. (*B*-*F*) ERG recordings from DESY P10 beamline. (*B*) ERGs of five wild-type (WT) flies (Canton-S genotype) and their mean ± SD (right; after subtracting the capacitive artifacts (*C*) to intensifying test X-ray pulses (above) and white-light controls (below) before and after X-ray stimulation. (*C*) ERGs of a dead fly (killed by freezing) show microelectrodes’ charging artifacts, increase with intensifying with X-ray pulses (above), and no responses to white control pulses (below). (*D*) ERGs of three mutants *_hdcJK910_*_-_ normal phototransduction but no synaptic transmission (missing on- and off-transients) to both X-ray and white stimuli. (*E*) Five blind *norpA^P24^*-mutant X-rays ERGs show similar capacitive artifacts as the dead fly (*C*) and no phototransduction or synaptic transfer. (*F*) X-ray ERGs of five blind *trp;trpl*-mutants (no phototransduction channels) show capacitive artifacts (*C*) with additional complexities. These combined artifacts were almost certainly caused by X-ray-induced strong photomechanical photoreceptor contractions “kicking” the recording electrode off the cornea (as seen in the corresponding X-ray videos). Predictably, ERGs to control white-light pulses were flat, indicating neither photoreceptor voltage response nor synaptic transmission.

To minimize electrical noise during the experiments, we electrically grounded the recording system. First, the two micromanipulators were fastened to a bespoke rectangular cuboid metal frame (Fig. S4 *C* and *D*). This structure had metal-mesh curtains that could be closed so that the fly was shielded inside a Faraday cage while leaving a narrow slit between its front and back curtains for the X-ray beam (Fig. S5*A*). Then, by connecting this Faraday cage and the micromanipulators to the microelectrode amplifier’s central ground, we obtained low-noise ERG recording conditions with very little or no 50 Hz mains hum.

As the initial control stimulus, and to test that each fly was healthy, we recorded its dark-adapted eyes’ global voltage response (ERG) to a 200 ms while-light flash (Fig. S5*B*). This stimulus was delivered from a white-LED, positioned ∼2 cm above the fly head, with the beamline lights off. About 30 s later, we recorded the same fly’s ERG (low-pass filtered at 500 Hz and sampled at 1 kHz) and photomechanical responses (100 frames/s) to X-ray pulses, in which intensities and durations (100-300 ms) were set by remotely operating the beamline’s neutral-density filters and high-speed shutter. To record the eyes’ photoreceptor movement video and ERG responses simultaneously, we used TTL-pulses to synchronize the shutter, the high-speed X-ray-imaging camera, and the microelectrode recording system.

The highest intensity X-ray pulses could partly taint the recorded ERG signal by capacitively charging the ringer-filled borosilicate microelectrodes. These electrode artifacts were most apparent in the ERGs of the dead flies (freshly-killed by freezing), which otherwise generated no electrical response (Fig. S5*C*), and their waveforms were microelectrode-dependent, varying slightly between the preparations and the exact electrode positioning in the beamline. For example, the charging artifact was reduced if only one electrode were within the X-ray beam instead of both. We utilized this observation by keeping the reference electrode outside the X-ray view, where it touched the torso (Fig. S4*B*) rather than the fly head, which would have been the conventional configuration. With this new arrangement, we could subtract the average dead-fly ERG from the ERGs of the living flies (Fig. S5 *B* to *F*). However, this procedure was not perfect as it left a small erroneous capacitive artifact that varied from fly to fly (Fig. S5 *B* to *F*, right subfigures). But since the tested phototransduction phenotype ERGs were unambiguous to both white-light and X-rays, showing their predicted waveforms, these minor artifacts made no real difference in the analyses.

The wild-type ERGs to X-ray pulses showed the intensity-dependent hyperpolarizing photoreceptor response component and the light On- and Off-transients (19, 36), caused by histaminergic synaptic transmission (20, 61, 62) (Fig. S5*B*). These ERG transients were missing from all the tested blind mutant fly recordings (Fig. S5 *D* to *F*). In further tests, by using longer (900-1,000 ms) X-ray pulses to evoke larger responses, the synaptic transient became more prominent (Fig. S6), consistent with the reported intracellular recordings (35, 36, 63). These dynamics were robust and repeatable. They were seen in every successfully-prepared living sighted *Drosophila* (n = 5), verifying that X-ray-induced phototransduction response and its synaptic transmission to the visual interneurons (Large Monopolar cells, LMCs (20, 61, 62)) happened normally in wild-type flies.

**Fig. S6.**
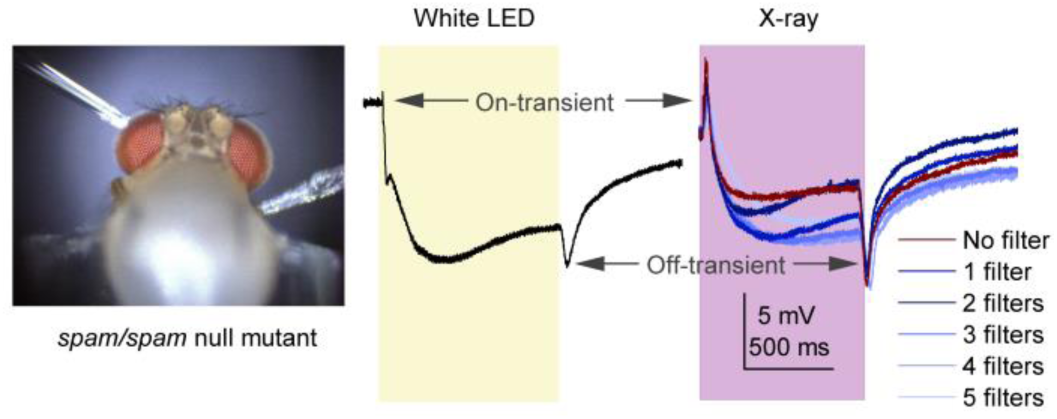
Longer X-ray pulses induce visible-light-like ERG responses with large transients. (see also Fig. S20, S25, and S26 in Sections SII.4 and SII.8, below). These exemplary ERG responses to 900 ms white-light and X-ray pulses (of different intensity) were recorded one after another from the same *spam*/*spam* null mutant *Drosophila* in DESY P10 beamline. Although *spam* mutants have reverted to an ancestral fused rhabdom state (64), their photoreceptors still respond to X-rays. The on- and off-transients, which indicate synaptic transmission from photoreceptors to large monopolar cells, are characteristically prominent in prolonged ERG responses (*cf.* Fig. S5*B*).

##### I.4 X-ray-imaging methods (general)

The X-ray imaging experiments were performed at two large-scale facilities: ESRF (beamline ID16b) and DESY (beamline P10). Both instruments are based on the same concept of focusing the X-ray beam to create a fine focal spot of below 100 nm using two mirrors in the so-called Kirkpatrick-Baez arrangement (one focusing vertically the other horizontally) (Fig. S7). By placing the sample at a small distance downstream of the focal spot and the detector further downstream, geometrical magnification is achieved. The effective resolution of the acquired radiographic projections is further limited only by the dimensions of the focal spot. In this experiment, we did not strive for the best spatial resolution. Rather, we optimized the setup to enable *in vivo* imaging by balancing the X-ray dose, exposure time, image contrast, and resolution. The optimization process is complex as the deposited X-ray dose scales with the 4^th^ power of the spatial resolution; furthermore, the temporal resolution is equally important to avoid blurring caused by the photoreceptor contraction.

Both instruments work with near monochromatic X-rays (on ID16B, ΔE/E was ∼1%). At ID16b at ESRF, we selected 17.5 keV photons corresponding to 0.07 nm wavelength; at P10 in DESY, the energy was set to or 13.8 keV, corresponding to 0.12 or 0.089 nm. These instruments’ approximated maximal used photon fluxes were 3 x 10^5^ photons/s/µm^2^ and 6 x 10^6^ photons/s/µm^2^, respectively. The estimated skin dose on the insect eye is 100 Gy per projection for the ID16b experiment. The detector’s image formation is governed by near-field diffraction of the partially coherent wavefront as transmitted by the sample. This is due to the partially coherent nature of the X-ray beam in both setups. The effective pixel size was set to 70 nm at ID16b and 167 nm at P10 with exposure times down to 10 ms controlled by a fast shutter upstream the sample (ESRF) or the camera frame rate (DESY). In the current study, we performed 2D radiography, for which the sample rotation allowed us to select the best viewing angles. For a typical experimental regime, consisting of seconds apart 200-300 ms X-ray pulse series (e.g., Fig. 1D), the flies survived multiple repetitions, with some lasting up to 40-50 min before dying with the photoreceptor movements ceasing.

**Fig. S7.**
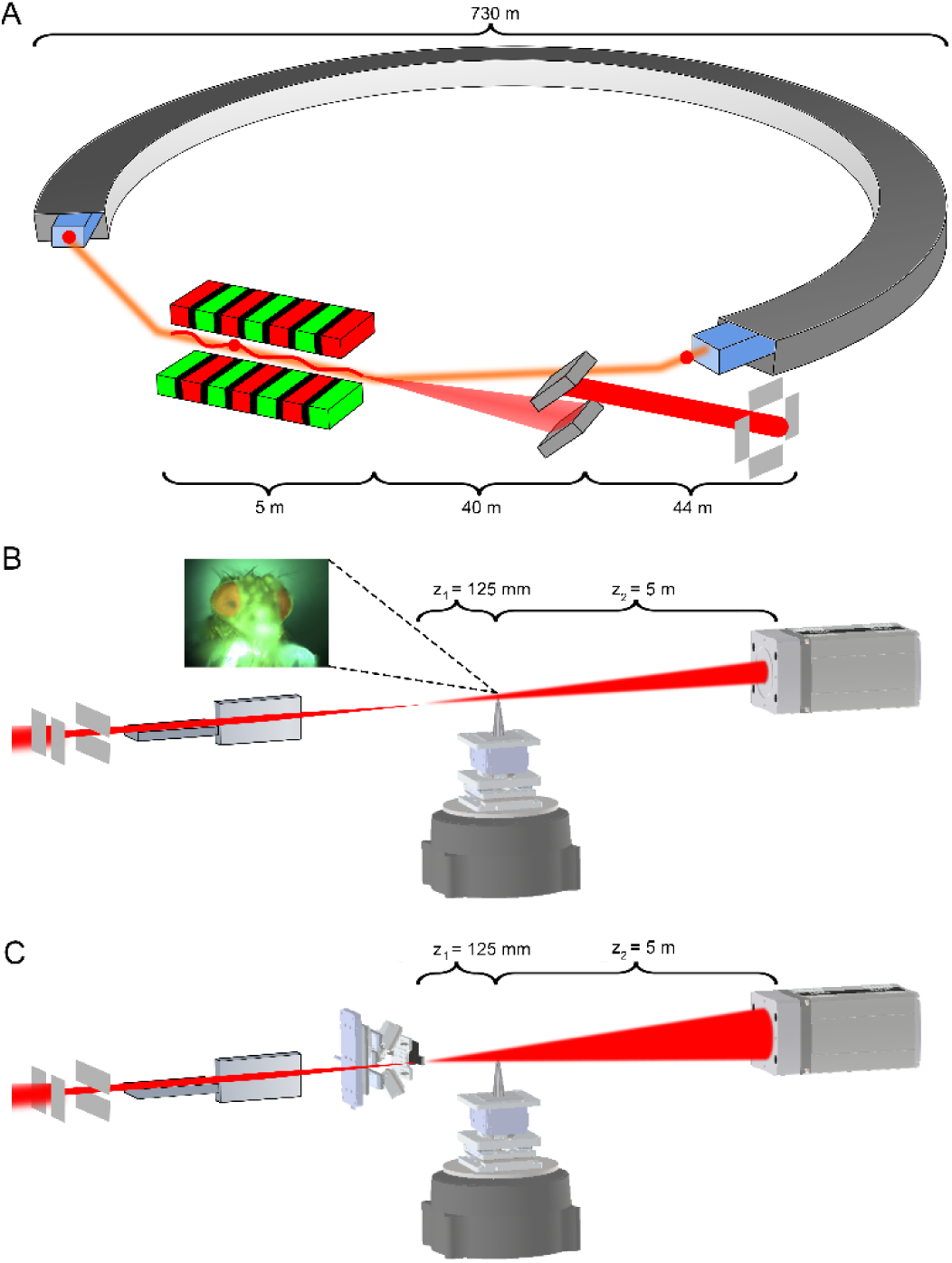
High-brilliance X-ray imaging of *Drosophila* photoreceptor microsaccades using synchrotron radiation setup at the GINIX instrument (P10/PETRA III, DESY) (*A*) Schematic of the synchrotron source and beamline. 2.3 km long DESY storage ring. An electron beam (red dot) travels into a 5-m- long undulator (red/green; the beam path is shown in orange and oscillation within the undulator in red). From the undulator, the synchrotron radiation coil (red cone) is directed 40 m to the double-crystal monochromator SI(111) (two gray squares). The monochromator exports 13.8 keV monochromatized X-ray beam (red) 50 m to the vertical and horizontal slit system of GINIX (four gray rectangles ). (*B*) KB-beam configuration. X-ray beam (red) from the GINIX slit system (four gray rectangles) travels through the head of *in vivo Drosophila*, positioned at motorized sample stage (dark gray), 5 m to the X-ray camera (gray square at the end). (*C*) KB-beam configuration with additional waveguide (WG) filter. X-ray beam (red) from the GINIX slit system (four gray rectangles); the two pairs of gray 3D squares are Kirkpatrick–Baez mirrors, focusing the X-ray beam through the sample (the *Drosophila* head), positioned at the motorized stage (dark gray). There is a 5 m distance between the specimen and the detector; the gray square at the end is the X-ray camera.

***The different X-ray imaging configurations used at GINIX endstation, P10, DESY* Fig. S7** shows a schematic of the different synchrotron beam configurations at beamline P10 of PETRA III (DESY, Hamburg), powered by a low-emittance E = 6 GeV,∼730 diameter storage ring (Fig. S7*A*). The source of the P10 beamline is a 5 m U29 undulator, operated in the third harmonic. The X-ray beam was monochromatized by a double-crystal SI(111) monochromator, installed at ∼40 m behind the source, to a photon energy of 10 keV. The entrance slits in the second experimental hutch (eh2), where the “GINIX” endstation (65, 66) is installed, received the beam at about 44 m behind the monochromator. For the *Drosophila* experiments, we used two different beam setups and imaging configurations at the GINIX station:

1. In the KB configuration, a pair of Kirkpatrick–Baez (KB) mirrors focused the X-ray beam to a size of 300 nm x 300 nm (Fig. S7*B*). With the respective focal distances of 300 mm and 200 mm, for the sequentially arranged vertically and the horizontally focusing mirrors, the setup achieved about 125 mm a working distance (in the air); once the beam leaves the diamond window of the evacuated KB mirror tank. Holographic projection images were recorded by an sCMOS sensor with a pixel size of 6.5 µm, coupled with a 1.1 fiber-optic to a 15 µm Gadox scintillator. The detector was placed 5 m behind the KB focus to achieve sufficient geometric magnification M. For the chosen M = (z1 + z2)/z1, an effective pixel size of 170 nm and an illuminated field of view in the object plane of 275 µm x 165 µm were reached.
2. In the waveguide configuration, a 1D X-ray waveguide (formed by a thin film Mo/C[35nm]/Mo sandwich structure (67), was placed into the X-ray focus of the KB-mirrors (Fig. S7*C*). This configuration yielded a smoother, Gaussian-shaped illumination, increased coherence, and a higher numerical aperture. The illuminated field of view in the object plane was 435 µm x 165 µm.

For both the configurations, the sample (specimen) was mounted on the same fully-motorized stage, with an additional dedicated optical table at the side for the microelectrode manipulator (Fig. S4). The sample could be inspected by a motorized on-axis video camera during the experiment.

##### I.5 Analyzing X-ray-induced global photoreceptor microsaccades

***Preprocessing*** The raw X-ray images were preprocessed using a custom computer script (68); first, to crop out any unused camera sensor area and only include those images in which the X-ray beam shutter was fully open. Next, a flat-field correction was performed by dividing each image by the corresponding mean flat image, based on the animal and the used X-ray attenuator setting. This pixel-wise division of the sample image by the non-sample image (the flat image) removed most of the non-sample features, caused, for example, by dust on the X-ray optics (vacuum windows) or imperfections of the KB surfaces, from the final images (Fig. S8). Each mean flat image was averaged from 20 to 200 frames. This procedure helped to estimate the non-sample features more precisely in the presence of photon shot noise and small image fluctuations. To further reduce the noise and fluctuations, especially in higher attenuator settings, we ran all flat-field corrected images through a Gaussian filter using spatial and temporal kernels of 7 and 3 pixels, respectively.

In the X-ray images, global photoreceptor activation appeared as a faint twist of rhabdomeres against a stationary background. To improve the detection of moving features, we added a further preprocessing step of band-pass filtering. The normalized spatial wavelengths outside the 0.03 to 0.1 range were set to zero in the Fourier space. This experimental preprocessing method resulted in a seemingly random mesh of strong-featured edges (Fig. S8*B*), in which motion visually corresponded to the unfiltered X-ray images. The band-pass range was selected to best contain the rhabdomeric motion component. While lower spatial wavelengths (higher frequencies) presumably contained more noise and higher wavelengths (lower frequencies) of more extensive stationary features such as facet lenses. Overall, this frequency filtering seemed to provide a better target for the cross-correlation-based motion analysis. However, it came with the expense of slightly reduced spatial specificity and the need for an additional scaling factor due to the stationary edges parallel to the motion.

In Fig. S3*E* and *F*, we used the background subtraction method by dividing each frame (i) by the minimum-value frame over the X-ray recording or (ii) by the mean-value frame to enhance any moving features while fainting or completely removing the stationary. We found that the minimum frame subtraction leads to better motion analysis results, although the images are noisier than the mean frame subtraction. Understandably, the background subtraction methods are not reliable when analyzing the motion of stationary features, which is why in Fig. S3*E*, we did not use them. Instead, we scaled the speed and displacement values to match their maximums with the maximums given by the minimum-frame background subtraction.

**Fig. S8.**
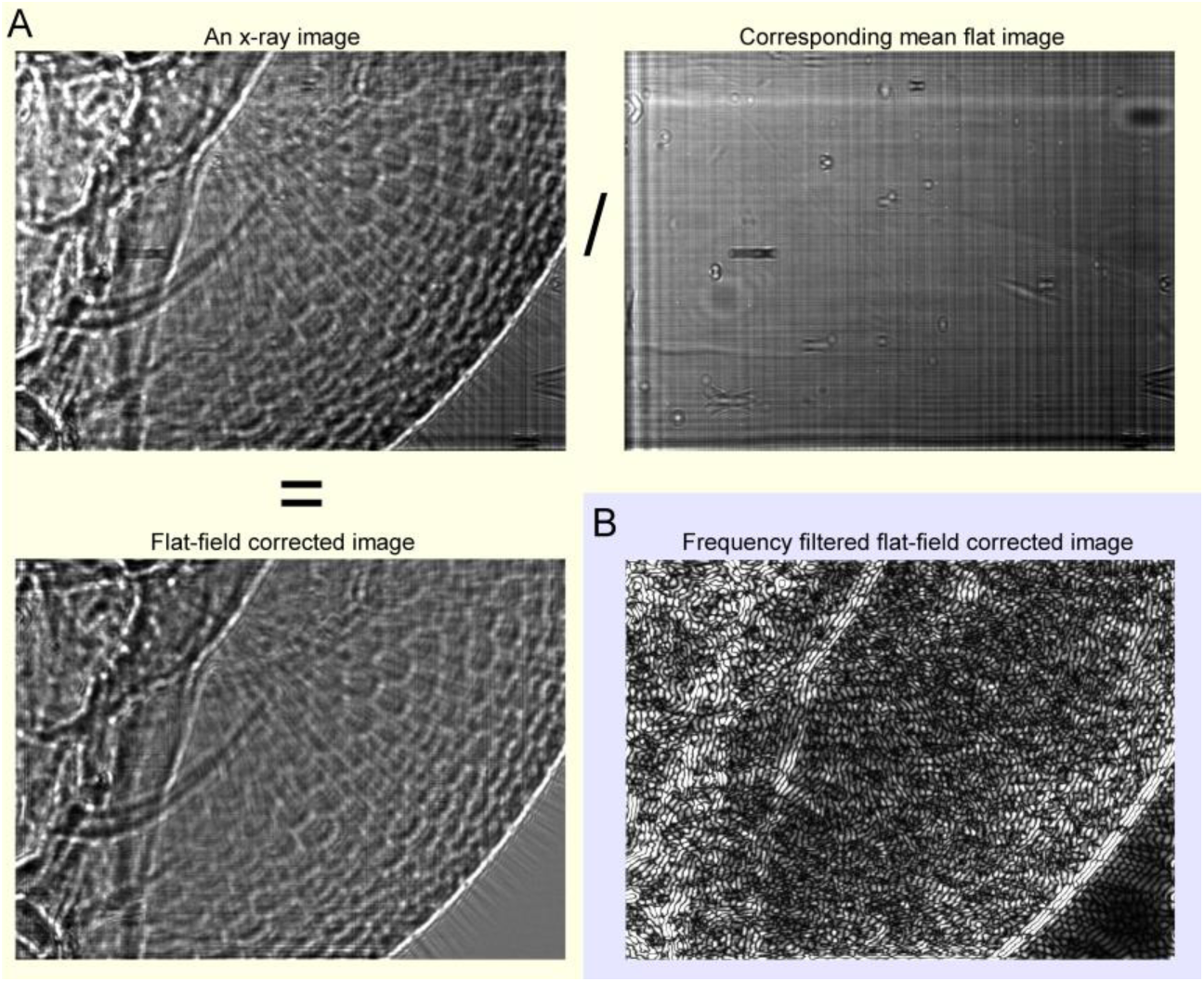
Preprocessing X-ray images. (*A*) Flat field correction by a pixel-wise division of a raw detector image and a flat (no-sample) image removes nearly all non-sample features. (*B*) The experimental frequency filtering of spatial frequencies creates a mesh-like structure of seemingly random edges. These robust features lead to fewer erroneous matches made by the cross-correlation-based motion analysis with few drawbacks.

***Motion analysis by cross-correlation*** To quantify the rhabdomeric motion from the preprocessed time-series of X-ray images, we created a custom Python script to perform template matching using the open-source computer vision library OpenCV. This script later refined and packaged under the name *Movemeter* calls the *cv2.matchTemplate* function to perform the following normalized cross-correlation between source and template images (Fig. S9)

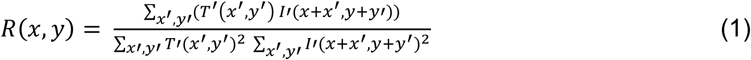

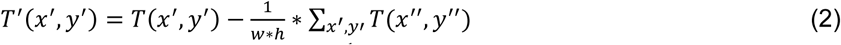

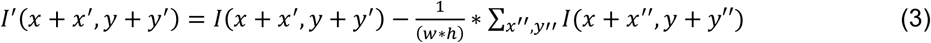

Here, *R* is the 2-dimensional cross-correlation image. *R*(*x,y*) is the value of a pixel at some x, y coordinates, *x’*, *x”* and *y’*, *y”* are summation indices limited by the cross-correlation window width *w* and height *h* within the ranges [0,1,2…,*w*-1] and [0,1,2,…,*h*-1], *I* is the source image, and *T* is the template image. We used a frame *k* as the source image and the subsequent frame *k+1* cropped by the cross-correlation window as the template image. *k* denotes the image frame index from 0 to N-1, while *N* is the count of frames acquired during an X-ray flash. In the cross-correlation image *R*, pixel values measure the similarity between the source and template images at each x, y location. Therefore, by taking the x, y location of R’s maximum value for each frame pair by *argmax* operation and calculating the cumulative sum, one can quantify the inter-frame displacement within a window in pixels. We restricted the inter-frame displacement to 10 pixels in maximum to reduce erroneous matches where a sudden displacement of a hundred or more pixels could happen between two subsequent frames.

The cross-correlation window size was set to 32 x 32 pixels, and a rectangular region of interest (ROI) was filled with windows every 32 pixels in *x* and *y*. The ROIs were placed on image areas where rhabdomeric motion was visually apparent while simultaneously avoiding non-rhabdomeric movement sources from antennae or tracheal tubes. In the absence of rhabdomeric motion, as it was for some blind mutants, we used ROIs similar to the wild-type flies.

The displacement values we report are mean results from all windows and characterize the mean motion within the ROI. The values were calculated as the directionless mean square root displacements using the x and y motion components as

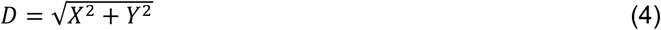

where *D*, *X*, and *Y* are displacement value arrays of length *N*-1, and *N* is the count of frames acquired during an X-ray flash. These values were transformed from the pixel units into micrometers using the pixel size unique for each detector and configuration. Where the frequency filtering preprocessing step was used, we scaled up all values by a factor of 4 to have perfect correspondence to the total displacement estimates made manually with a ruler in Fiji (ImageJ 1.53c). The need for this additional scaling was likely because the frequency filter preprocessing step also produced stationary edges parallel to the rhabdomeric motion, leading to the rhabdomeric motion underestimation (when calculating the mean displacement over many windows within an ROI).

**Fig. S9.**
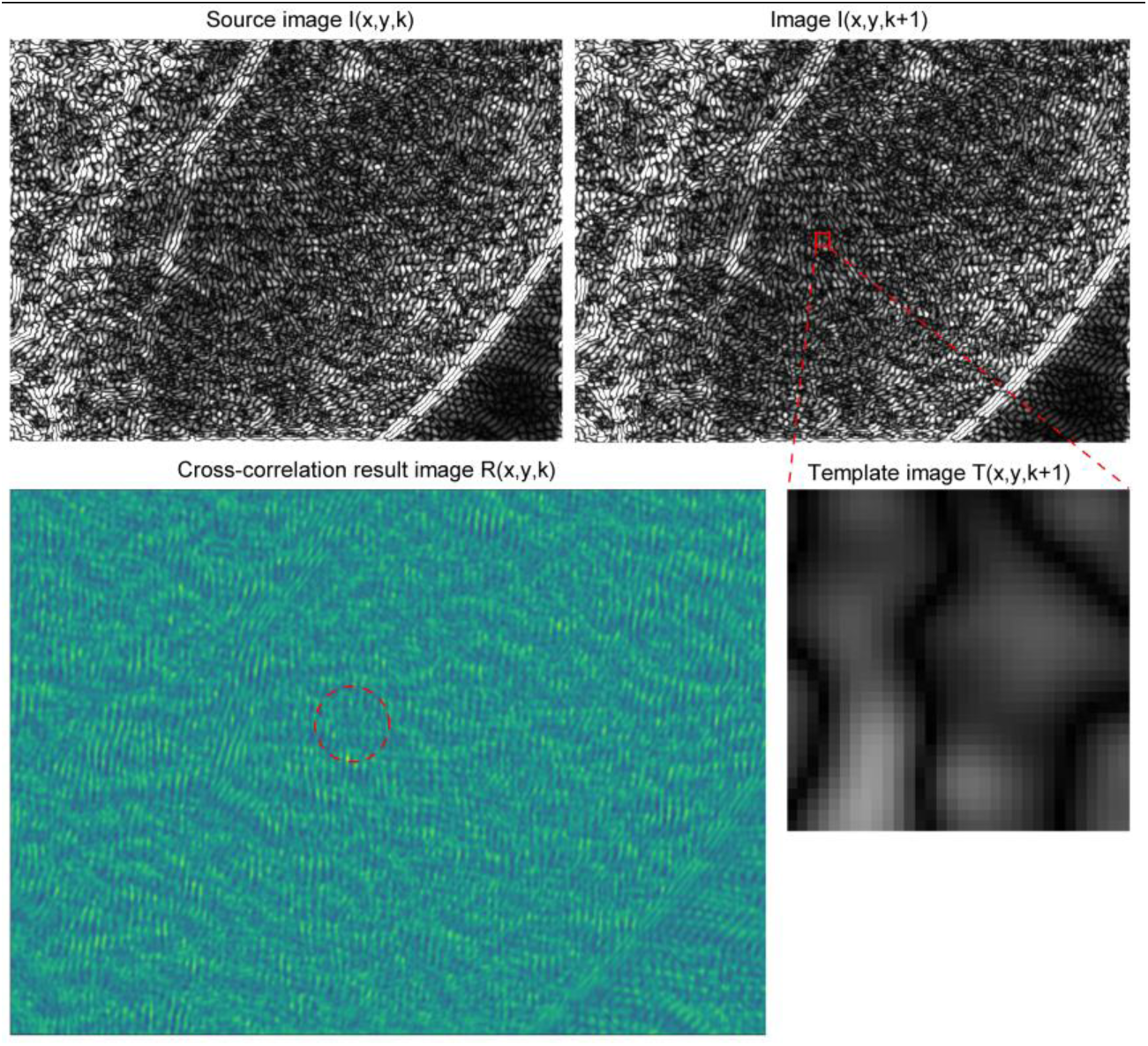
Motion analysis using cross-correlation-based template matching. The cross-correlation image’s maximum value tells the template image’s best-matched location on the source image highlighted by the red dashed circle. Comparing these locations between many frame *k* and *k+1* pairs reveals kinematics occurring within a window over time.

***Heat-map analysis*** To calculate the rhabdomere motion heat maps during an X-ray flash, we used our MATLAB-based implementation that was also used for some of the rhabdomere displacement graphs. It uses the *imregtform* function from MATLAB’s Image Processing Toolbox to estimate geometric transformation that aligns the source image *k* with the template image *k+1* cropped by the window to arbitrary numerical accuracy set by the optimizer parameters. We configured the *imregtform* optimizer as *monomodal* (images having similar brightness and contrast, captured with the same sensor) and used the following tuning of the optimizer parameters

optimizer.GradientMagnitudeTolerance = 10^-7^
optimizer.MaximumIterations = 1000
optimizer.MaximumStepLength = 0.1
optimizer.RelaxationFactor = 0.99

However, instead of selecting an image subsection for the heat maps, the motion analysis windows were set to span the whole image. We used the window size of 32 x 32 pixels and windows, laid out every 4 pixels in x- and y-coordinates, filling the original image in a grid-like manner. Much smaller window sizes led to noisier heat maps, whereas larger window sizes resulted in blurrier heat maps as the heat-map image pixels became more correlated with their neighbors. The 4-pixel inter-window-distance was the lowest value that still gave reasonable computational times on University’s computing cluster.

By estimating all the translations between *k* and *k+1* frames for k=0,1,2…*n*-1, where *n* is the number of images taken during an X-ray flash, we obtained the *X* and *Y* displacement arrays over time for each window in pixels. Then the data was converted to directionless mean-square movement values, and finally, these mean square values were presented as N-1 heat-map images using MATLAB’s *imshow* function. Clearly erroneous pixels, in which a sudden inter-frame change of tens of pixels or more occurred, were set to zero. We only present the final heat-map frame (in this paper, excluding the video) since it characterizes the overall displacement within the complete 200 ms time duration.

The scripts to process and analyze the X-ray images are downloadable from the repository: https://github.com/JuusolaLab/Hyperacute_Stereopsis_paper/tree/master/AnalyzeMovementData

#### II *In vivo* high-speed optical imaging of photoreceptor microsaccades

**Overview**

This section describes the experimental and theoretical approaches to measure photomechanical photoreceptor movements (microsaccades) (10, 11) across the left and right *Drosophila* eye using *in vivo* high-speed imaging. It gives central background information and additional supporting evidence for the results presented in the main paper, including:

- mmatidial R1-R7/8 rhabdomere patterns across the left and right eyes are mirror-symmetric and aligned so that their R2-R5 axis is largely collinear with frontally expanding optic flow field.
- left and right eye microsaccades are mirror-symmetric but generally move along the R1-R2-R3 rhabdomere tips’ orientation axis. Therefore, the microsaccade movement directions are determined primarily by the eyes’ mirror-symmetric ommatidial ultrastructure that rotates concentrically, as organized developmentally during the eyes’ morphogenesis.
- mirror-symmetric photoreceptor microsaccade directions extend the two eyes’ binocular (frontally overlapping) sampling range for stereopsis to about 30°.
- microsaccades, R1-R7/8 rhabdomeres move simultaneously laterally and axially, away and closer to the ommatidium lens. These fast morphodynamics shift and narrow the photoreceptors’ receptive fields optically.
- are robust in the dark- and light-adapted eyes, with light adaptation accelerating their dynamics while retaining contrast sensitivity. At room temperature (∼20-22 °C), the microsaccade frequency response can follow contrast modulation at least until ∼27-32 Hz.
- a microsaccade, ommatidial R1-R7/8 move as a single unit. Inside an ommatidium, rhabdomeres are mechanically coupled so that even a single photoreceptor’s photomechanical activation alone moves all R1-R7/8 sideways, generating the microsaccade’s lateral component.
- R1-R7/8 photoreceptors in an ommatidium contribute to the microsaccade; the more photoreceptors are light-activated, the larger the microsaccade. Therefore, microsaccades can be used as a metric to quantify the light-activated phototransduction state.
- *Drosophila* eye is somewhat sensitive to mechanical stress, with accidental denting during *in vivo* preparation making, especially for some mutants and transgenic flies, reducing functional integrity to generate the lateral microsaccade components.
- findings are consistent with the hypothesis that the eye-location-specific microsaccade movement directions require well-organized interommatidial rhabdomere pivoting (angled anchoring) and mechanical coupling (possibly by inter-rhabdomeric tip-links).
- Finally, during and between experiments, the microsaccades’ dynamic variability suggests that a fly’s intrinsic activity state – in the form of synaptic feedback signals to the photoreceptors - might further modulate them. Note, this variability was not caused by the flies deteriorating due to IR illumination (for visualizing the photoreceptors), as the average microsaccade size rarely diminished during the experiment.

##### II.1 Deep pseudopupil imaging (of optically superpositioned photoreceptors)

Photomechanical *Drosophila* photoreceptor movements (10, 11), named *photoreceptor microsaccades* (11), can be viewed non-invasively *in vivo* by observing the eyes’ deep pseudopupil (Fig. S10) (24). Deep pseudopupil (DPP) is a virtual image of multiple distal R1-R7/8 rhabdomere endings, which align with the angle the eye is observed at while being ∼10x-magnified by the ommatidial lens system (Fig. S10 *A* and *B*; see also Movie S4 that illustrates its optical principle) (24). Here we describe how to map such optically superpositioned R1-R7/8 rhabdomeres’:

i. **Angular orientation**
ii. **Stereo vision range -** the central binocular visual field, which is viewed simultaneously by the corresponding left and right eye photoreceptors
iii. **Photomechanical microsaccade movement components**
iv. **Microsaccade movement directions** to light flashes, as delivered at their receptive fields (RFs) across the *Drosophila* eyes.

Because R8 rhabdomere lies directly underneath R7 in optical superposition with neighboring ommatidia’ R1-R6 and contributes to photoreceptor microsaccades (see Section SII.8, below), we consider and call the DPP rhabdomere pattern as R1-R7/8.

**Fig. S10.**
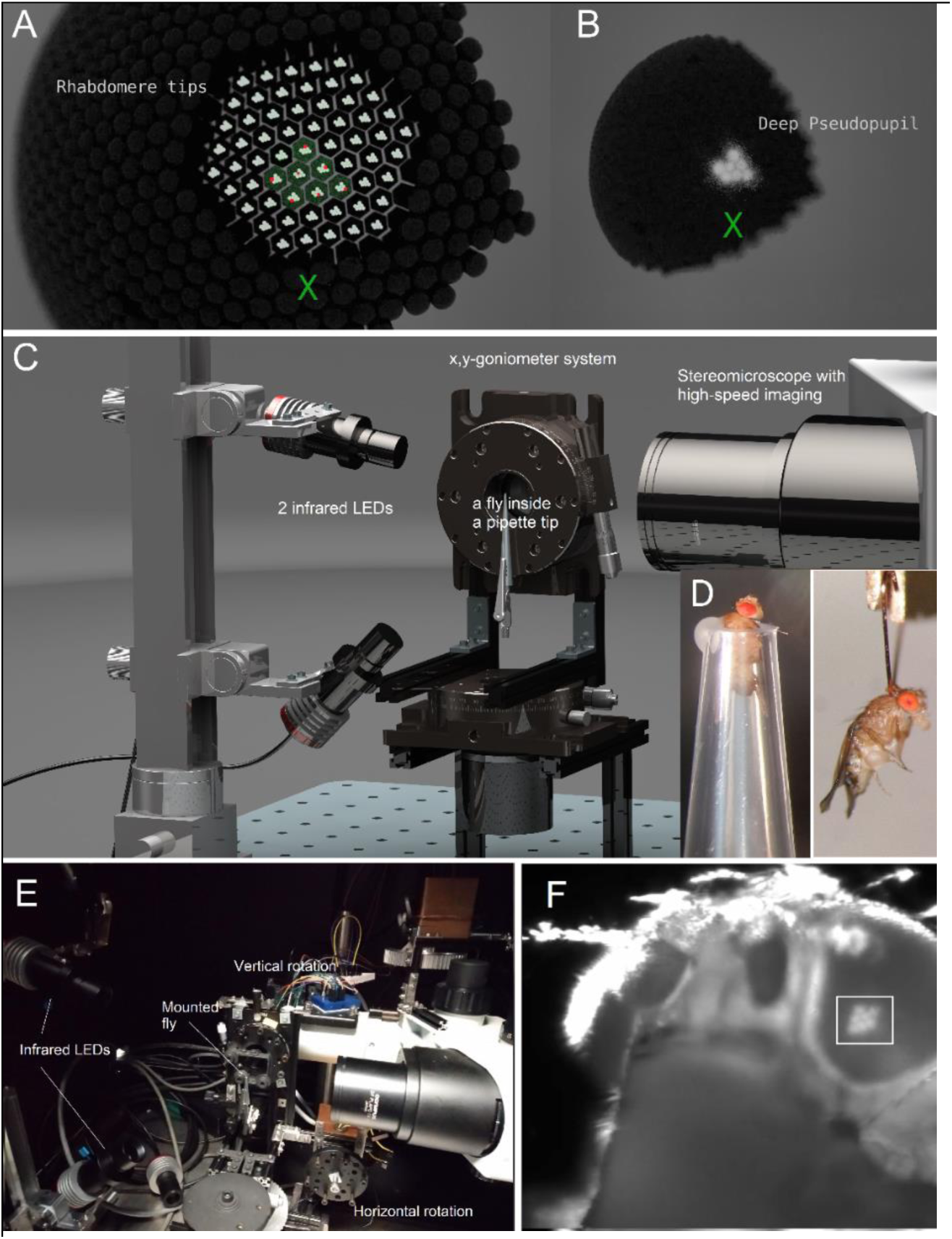
Imaging deep pseudopupil and photoreceptor microsaccades with the goniometric system. (*A*) R1-R7/8 photoreceptor rhabdomere tips (white dots) centered in hexagonal ommatidia that tile the right *Drosophila* eye. The red R1-R7/8 rhabdomeres inside the green-tinted ommatidia are in optical superposition (with R7 on top of R8). These rhabdomeres point to the same small area in the visual space and respond only to incident light changes (green X) at that visual area. (*B*) The optically superimposed rhabdomeres form a deep pseudopupil (DPP) virtual image (24) (∼10x-magnified by the ommatidial lens system), and their light-activation generates a DPP photoreceptor microsaccade. See Movie S3 and S4. (*C*) The DPPs of local photoreceptor rhabdomeres were observed and recorded across the fly eyes *in vivo*. This method combined transcutaneous infrared back-illumination (invisible to flies) with their goniometric x,y-rotation under a long-working-distance microscope imaging, using a high-speed camera system. In the same experiments, the DPP photoreceptors could also be light-activated by delivering UV- or green-stimulation through the microscope optics at the center of their receptive fields (RFs), evoking photomechanical DDP photoreceptor microsaccades. (*D*) Goniometric DPP imaging was performed both from pipette-tip-held and tethered *Drosophila* preparations (with legs and wings either wax-restrained or not), in which the fly head was fixed immobile. (*E*) A side-view of the goniometric high-speed imaging system, which enabled us to systematically map the DPPs, stereoscopic range, and microsaccade dynamics across the fly eyes. (*F*) We used IMSOFT software to log the exact angular camera position in respect to the recorded DPP images; needed for mapping the DPP orientation, stereoscopic range, and microsaccade movement directions across the fly eyes.

Head-fixed living intact *Drosophila*, either held inside a “cut-to-fit” pipette-tip or tethered to a small hook (see Section II.4., below), were connected to the center of a custom-made goniometric stage (Fig. S10 *C* to *E*) for precise x,y,z-positioning and rotation in both the horizontal and vertical axes. A fly’s fine positioning could be set either by remote-controlled stepping motors or manually. During the positioning and later high-speed imaging, each fly was monitored under antidromic infrared (IR) light, which *Drosophila* cannot see (22) but high-sensitivity CMOS camera sensors detect readily. This back-illumination through the fly head was delivered by two 850 nm LEDs (powered by a Cairn Research optoLED driver, UK), mounted on a separate x,y,z-adjustable positioning arm (Fig. S10 *C* and *E*).

The IR light was turned on for each DPP recording only briefly (210 ms in most experiments). We measured its heat production during the system calibration, deliberately keeping its intensity low to minimize tissue damage while still obtaining a sufficiently high DPP image signal-to-noise ratio for the high-speed recording. With this arrangement, we could perform hours-long DPP recordings in living *Drosophila* without noticeable deterioration in the observed dynamics.

###### II.1.i Mapping deep pseudopupil angular orientation across the eyes and comparing it to the optic flow fields

A fly’s exact position was recorded using two 1,024-step rotary encoders (E6B2-CWZ3E, YUMO, China) connected to the open-source electronics platform Arduino microcontroller (Italy) and fed into a computer running the IMSOFT software (Joni Kemppainen, 2019-21). Each fly was centered by its eyes at both 0° and -90° vertical rotation. The vertical rotation reference point was where the left and right eye pseudopupils aligned with the antennae pedicels. We first examined its DPP microsaccades to UV and/or green flashes. If these occurred, indicating that the preparation had no apparent structural eye damage (see Section II.4., below), it was rotated through the horizontal x-axis at 0° y (vertical), with imaging – either with or without light-stimulation - being triggered every 10° from −50° to +40° (x-range). Further horizontal-range imaging was carried on for every 10° of vertical rotation, covering −110° to +110° (y-range), until it was impossible to see the DPP. During imaging, the flies were shielded from ambient light by a black-painted Faraday cage and lightproof curtains. The experiments were conducted at room temperature (∼20-22 °C).

We imaged the DPPs across the fly eyes using an Olympus SZX12 stereomicroscope with a long-working-distance DF PLAPO 1x objective (Fig. S10 *C* and *E*) of 0.11 numerical aperture (NA). IR images were recorded using an Orca Flash 4.0 CMOS V3 video camera (Hamamatsu, Japan) at 100 frames/s, outputting 1024 x 1024 pixels at 2 x 2 binning. The camera was computer-controlled by the IMSOFT software (Fig. S10F), allowing for experimental parameter modifications.

Ultrastructurally, the R1-R7/8 rhabdomere patterning inside ommatidia, and thus their DPPs (with R7 endings concealing R8s below), are mirror-symmetric both vertically (between the left and right eye) and horizontally (along the equator, diving the dorsal and ventral eye halves (21, 69)) (**Fig. S11**). Using goniometric imaging, we further quantified how the ommatidial R1-R7/8 rhabdomere patterns align across the eyes. The local rhabdomere orientation, which was defined as the rotation of the angle between R3-R2-R1 (yellow line) and R3-R4-R5 (green), shifts gradually and systematically, generating the characteristic mirror-symmetric global map for the left and right eyes. The map reveals that rhabdomeres align locally to follow a global concentrically-expanding diamond-shaped pattern, suggesting that their orientation at each eye position is fixed developmentally to the corresponding frontally expanding optic flow field. Note that in female blowflies (*Calliphora*), the local ommatidial row orientation, determined by the deep pseudopupil method, also changes with the eye location (70). There, the local ommatidial row orientation was found to be aligned to the receptive fields of specific lobula plate tangential cells, thought to signal translations along the longitudinal body axis and roll rotations (71).

**Fig. S11.**
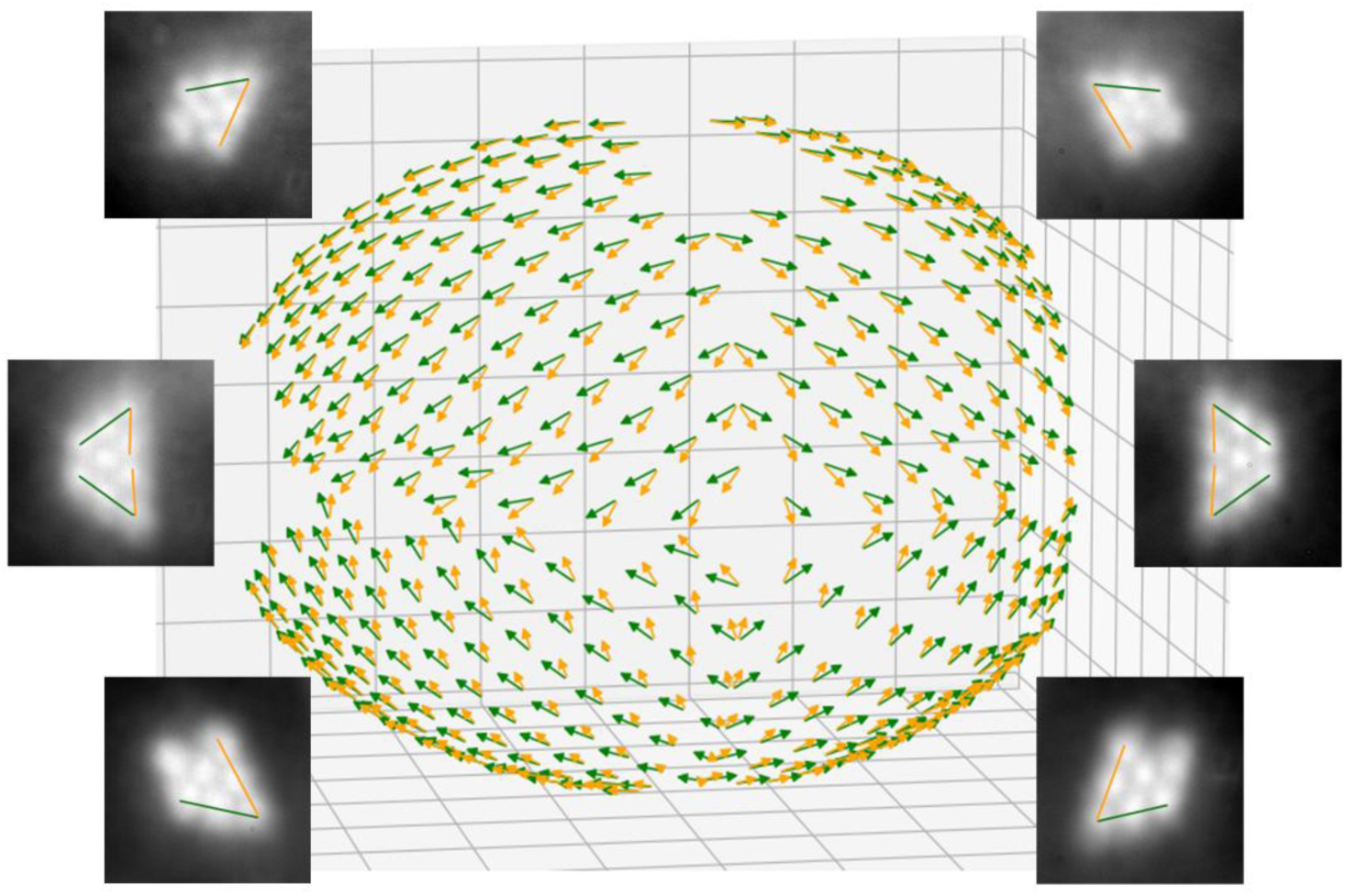
Developmental R1-R7/8 rhabdomere orientation map across the *Drosophila* eyes. Local DPP images – *i.e.,* optically superpositioned photoreceptor rhabdomeres - were recorded at different eye locations systematically (by x,y-rotating a fly in the goniometric system; Fig. S10). The ommatidial left and right eye rhabdomere patterns (inset images) are horizontally and ventrally mirror-symmetric, aligned in a concentrically expanding diamond-shape. Notice that at the equator, the DPP images of the corresponding just-above and just-below rhabdomeres fuse. The map shows the average global rhabdomere orientations of 5 flies. The yellow arrows indicate the ommatidial orientations of R3-R2-R1 rhabdomeres and the green arrows of R3-R4-R5 rhabdomeres across the left and right eyes.

Therefore, we further computed how accurately the local ommatidial rhabdomere orientations across the two eyes align with the concentrically expanding optic flow field, which they would face in a forward flight (Fig. S12). Characteristically, when flying forward, the fly head is at an upright posture, having a 10.1° backward tilt (Fig. S12*D*). The calculations included this slight tilt.

***Optic flow field calculations and field error*** The directions of the optic flow field as experienced by a forward flying fly were calculated using a simple sphere-tangent algorithm. The source code is available from: https://github.com/JuusolaLab/Hyperacute_Stereopsis_paper/tree/master/AnalyzeMovementData

The flow directions depend on the head’s rotation with respect to the locomotion direction. Three rotation axes (yaw, pitch, roll) unambiguously express the head rotation, and they were fixed as follows. First, the pitch axis is naturally set by the left-right symmetry. Second, the roll axis is defined here through the zero rotation (0, 0, 0) to match the GHS-DPP coordinate system. At zero rotation, the antennae point towards the positive y-axis so that an observer located on the positive y-axis would see the DPPs aligning with the antennae pedicels. Finally, the yaw axis is perpendicular to the two other axes.

In the sphere-tangent algorithm used for the optic flow simulation, a vector −*j*^, pointing towards the negative y-axis, was forced to a sphere’s tangent plane in DPP-microsaccade data interpolation points as

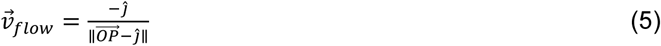

where 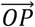 is the vector from the origin (sphere center) to a point *P* on the sphere’s surface and *j*^ is the Cartesian unit vector in the direction of the y-axis. After normalizing the vectors to unit length, these vectors gave the optic flow field directions at (0, 0, 0) head rotation, assuming that the fly is flying towards the positive y-axel. Notably, the optic flow direction is undefined at the field’s source and sink points. Finally, the optic flow at other fly rotations (with the fly still flying towards the positive y-axis) was calculated by rotating the optic flow field along the pitch, yaw and roll axes and using a matrix multiplication with the appropriate rotation matrices.

The optic flow field at different head rotations, the rhabdomere orientations, and the DPP-microsaccade directions were compared against each other. The difference between any two of these unit vector fields, here referred to as A and B, was calculated point-wise as:

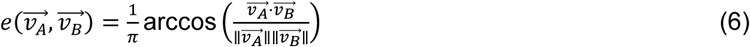

where 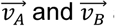 are vectors located on the same point, and the operators ⋅ and ∥ ∥ denote the inner product and the vector norm (length), respectively. The *e* error is directly proportional to the angle between the two vectors, and its values are limited on the closed interval [0,1]. Here *e* = 0 means that the vectors are parallel (no error), *e* = 0.5 means that they are perpendicular (50% error), and *e* = 1 means that the vectors are antiparallel (maximal error). Finally, the average mean error between the fields was calculated as the geometric mean of individual vector errors.

The receptive field (RF) fast movement phase was calculated as an inverted DPP microsaccade vector. This procedure was done because the convex lens system inverts the image on the retina, making the receptive fields move in the opposite direction compared to the rhabdomeres. However, as a virtual image (24), the DPP is non-inverted and moves in the same direction as the rhabdomeres. Finally, the RF slow-phase was calculated as the inverse of the RF fast-phase. This assumption was needed because the much slower (seconds-long) relaxation phase of the DPP microsaccades was not imaged during the GHS-DPP experiments.

For each recorded eye position, we first compared the corresponding optic flow field direction (Fig. S12*A*, purple arrows) to the measured deep-pseudopupil R1-R7/R8 rhabdomere pattern orientation (Fig. S12*B*). Then, we performed a global search for the fixed angle, in which the optic flow lines cut the R1-R7/R8 rhabdomere pattern orientation across all recorded eye positions with minimum error (Fig. S12*C*). In nearly every position, the optic flow lines - as these curve around the two eyes - cut their rhabdomeres primarily along the R2-to-R5 axis (Fig. S12*D*), with only 15.6% mean error over the entire global map. This analysis established that ommatidial rhabdomeres rotate during development so that their R2-to-R5 axes align collinearly to follow the directions of local parallax vectors within an optic flow field axes generated during forward translation. Video-file showing the analyses can be downloaded from: https://github.com/JuusolaLab/Hyperacute_Stereopsis_paper/tree/master/AnalyzeMovementData

Notably, in *Calliphora*, the orientation of ommatidial rows within the hexagonal eye lattice and the preferred local directions of some lobula plate tangential cells are aligned (71).

**Fig. S12.**
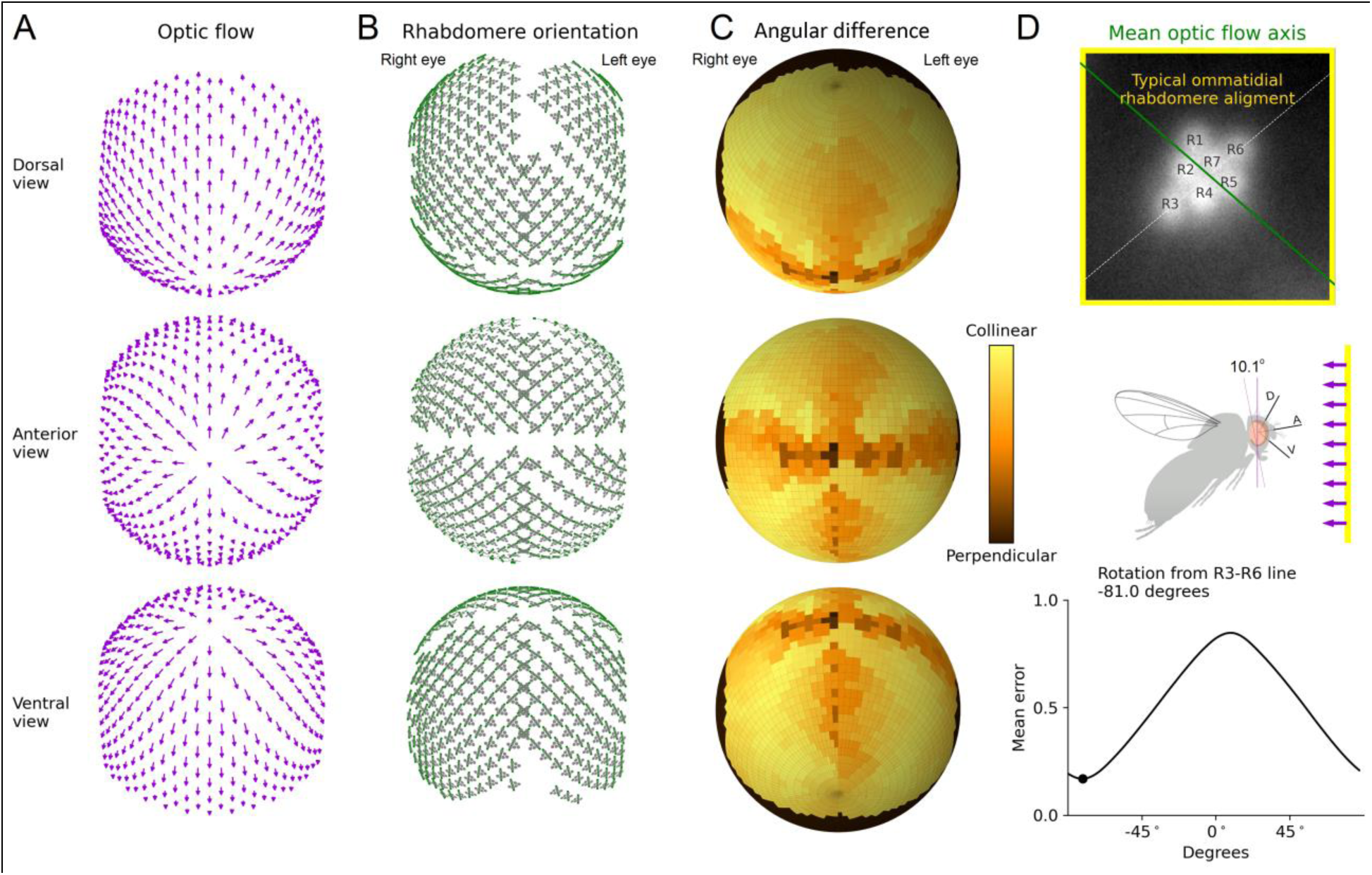
Ommatidial R1-R7/8 rhabdomeres across the *Drosophila* eyes align with the forward flight optic flow field. (*A*) Optic flow field facing the *Drosophila* eyes in the normal forward flight position with the fly head in a slight 10.1° backward tilt (*cf*. the fly schematic in *D*) (*B*) Local rhabdomere orientation patterns (gray) across the left and right eyes; plotted with their R2-R5 axis (green), which best aligns them to the optic flow (*cf*. the DPP image in *D*). The rhabdomere orientation map shows the mean of 5 wild-type flies. (*C*) The minute differences between the local rhabdomere orientation R2-R5 axes and optic flow axis over most of the eyes confirm their global collinear alignment. Notice that at the focal point, from which the flow field radiates outwards, this comparison becomes less reliable, resulting in a slightly darker central region in the difference map. (*D*) Ommatidial rhabdomeres are aligned across the eyes so that optic flow crosses them along the R2-R5 axis (-81° rotation against the longest R3-R6 axis). The mean error between rhabdomere orientation and optic flow was calculated for the characteristic upright head position (with a slight 10.1° backward tilt) as seen in free flight. Because of the biased focal point values in *C*, the minimum mean error is a slight overestimate, meaning that the ommatidial rhabdomeres’ R2-R5 axis optic flow alignment is ≥85% accurate globally (<15.6% error).

###### II.1.ii Mapping *Drosophila*’s stereo vision range

By knowing the exact angular camera position regarding the left and right eye DPP images, we could further use the scanned images across the eyes to generate a map of the field of view shared by both eyes (Fig. S13). This optically measured frontal binocular range gives the angular x,y-limits of a fly’s potential stereo vision. Overall, these binocular overlap measurements concur with the earlier results using epifluorescence deep pseudopupil mapping (72, 73). Movie S3 shows how the goniometric DPP imaging was used to map the eyes’ binocular stereo range in relative darkness; *i.e.,* having no visible light stimulation. Notice also in the Movie how the rhabdomeres’ angular orientation shifts systematically with eye location, following the developmental R1-R7/8 rhabdomere orientation map (*cf*. Fig. S11 and Fig. S12*B*).

**Fig. S13.**
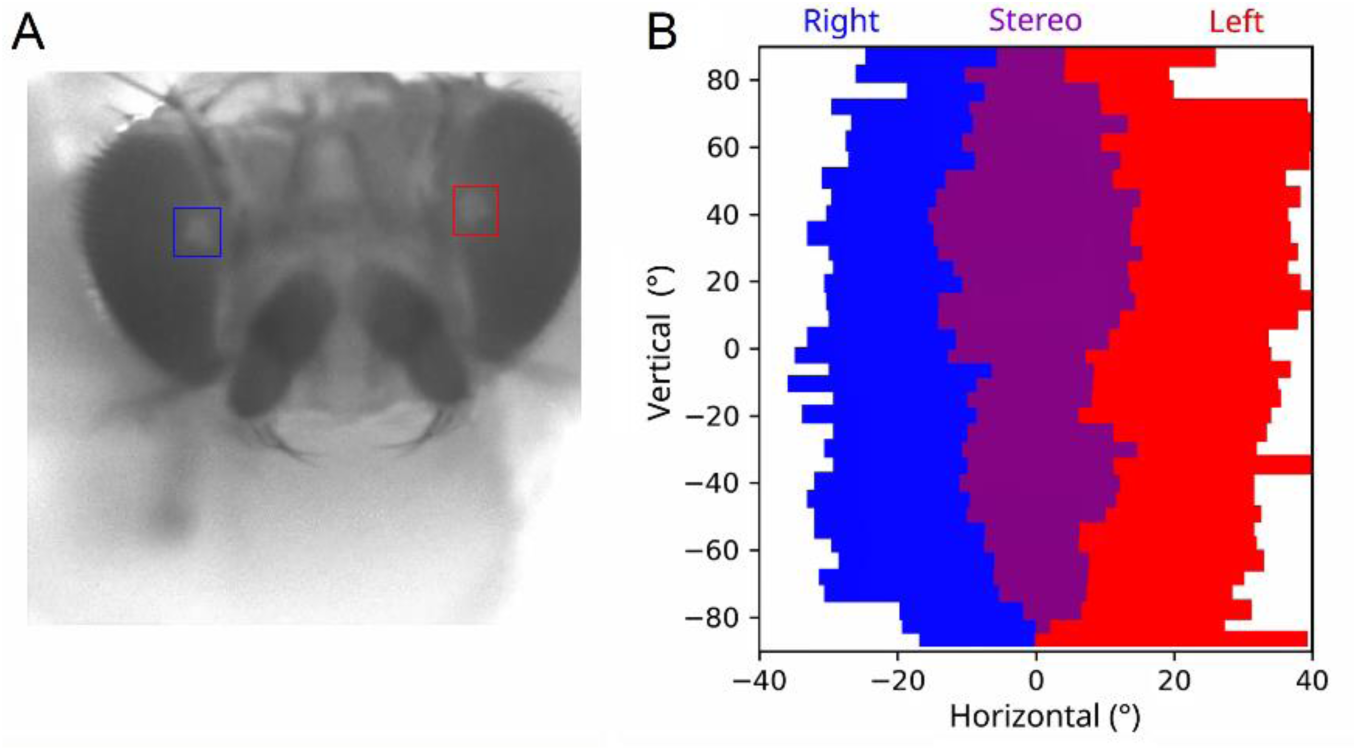
Mapping *Drosophila*’s stereo vision range by goniometric deep pseudopupil imaging. (*A*) A wild-type female fly shows normal right (inside the blue box) and left (red box) eye DPPs frontally, indicating that its two eyes collect simultaneously light information from the same point in space. (*B*) The same fly’s optically estimated stereoscopic visual field (purple), extending about 180° vertically and 10-35° horizontally. The blue and red bars show the frontally measured right and left eyes horizontal visual fields at each tested vertical position. Their purple overlap demarcates the predicted stereo field. Movie S3 shows how the goniometric system was used to measure individual flies’ stereo vision range.

We tested experimentally (Fig. S14) and through computer simulations (Section II.2) the possibility that the numerical aperture (NA) of the used microscope biases the estimated stereo range. In the former, one fly’s stereo range was measured repeatedly under three different NA configurations of 0.11, 0.054, and 0.015 at a fixed vertical rotation (Fig. S14*C*). Only the 0.015 NA resulted in smaller range estimates (Fig. S14D; t-test p = 1 and p = 2.09 x 10^-3^), but this is probably a side effect caused by the reduced image quality that makes it harder to separate the DPP edge from the eye edge visually.

**Fig. S14.**
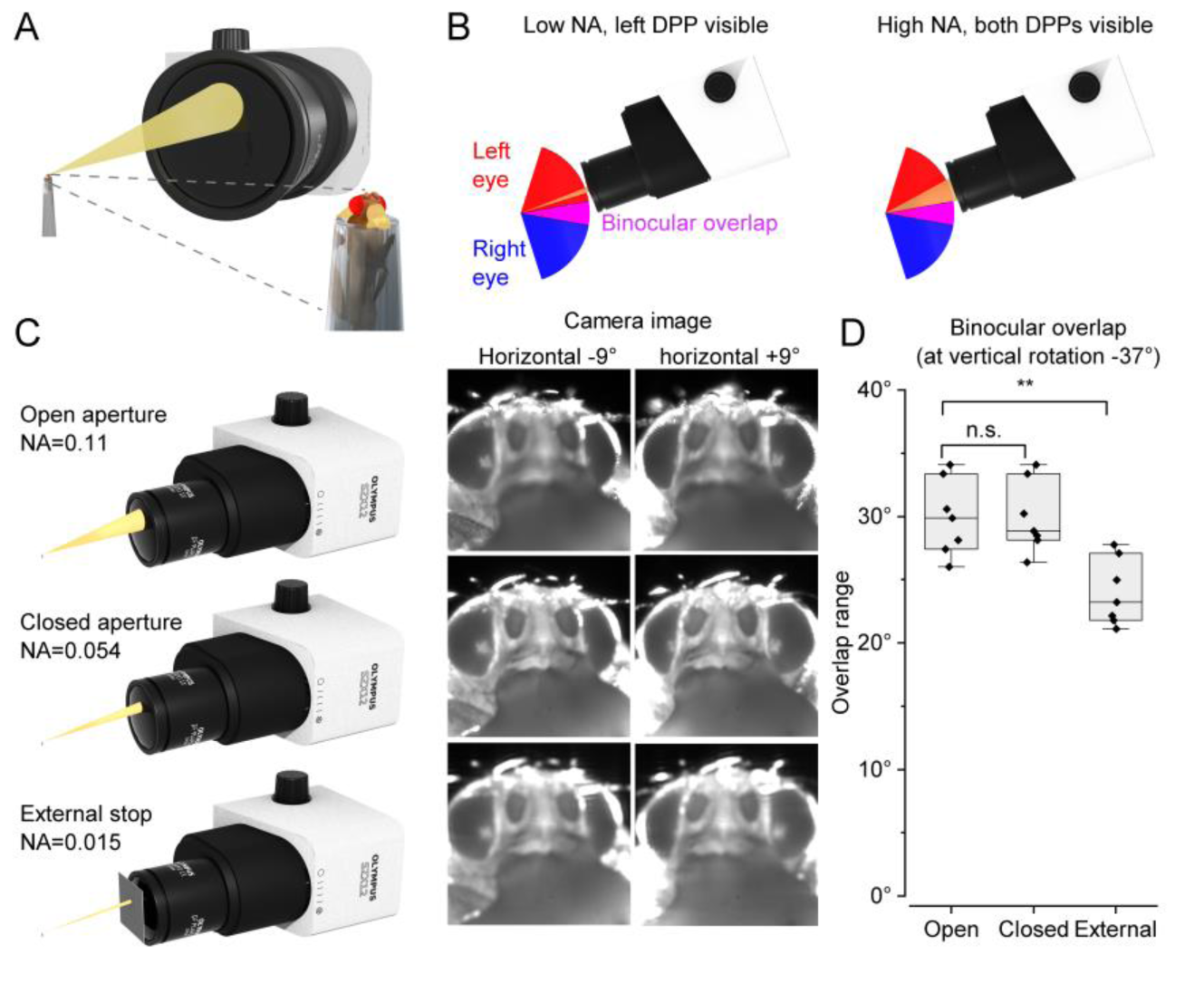
Testing how the microscope aperture affects the stereoscopic visual field estimation. (*A*) Only part of the light originating from a point in the fly eye enters the microscope to form an image, as illustrated here with a yellow cone. (*B*) In a high NA microscope, estimation broadening can occur because the light is collected over a range of horizontal angles. (*C*) The binocular overlap of a single fly was measured at vertical rotation -37° seven times for each of the three aperture configurations: (i) the microscope’s aperture control fully open, (ii) the control closed, and (iii) using an external aperture stop with 2.5 mm wide square opening. Images taken at ±9° horizontal rotations show no systematic broadening under our experimental settings. (*D*) Quantified binocular overlaps show no difference between the open and closed aperture controls. The 6° reduction using the external stop is likely a result of the reduced image quality, making it difficult to distinguish the R5 DPP from the eye edge.

###### II.1.ii Quantifying photoreceptor microsaccades’ lateral and axial components

We recorded *Drosophila* photoreceptors’ DPP microsaccades to 200-ms-long 365-385 nm UV- and 546 nm green-LED flashes. The LEDs were mounted and centered in the microscope’s eyepiece socket, which through the microscope head’s dichroic mirror (image splitter) shared the same “best-focused” DPP image with the camera. At this point, all ommatidia’s optical axes converge to the eye’s center of curvature (21). Therefore, the axially centered light stimulation was delivered through the microscope optics at the receptive fields (RF) of those R1-R7/8 rhabdomeres in optical superposition. During stimulation, 20 images (at 100 frames/s) were taken by IMSOFT and saved in the TIFF format.

***Image analysis*** Imaging data were analyzed using a custom-made DPP analyzer program (Joni Kemppainen, 2019-21). This program performed image cross-correlation analyses (11) to quantify the photomechanical microsaccade sizes, temporal dynamics, and moving directions. These data could then be extracted and plotted in other software packages.

***Cross-correlation analysis*** Photomechanical microsaccades were analyzed from high-speed videos using cross-correlation analysis as described earlier (11). 2D cross-correlation was calculated between each frame and the reference frame, typically the frame before the stimulus. Weighted means in x- and y-direction were calculated from each 2D cross-correlation result, which was ≥95% of the maximum (peak) value. Lastly, the reference frame cross-correlation x- and y-positions were subtracted from each frame, giving their difference to the reference frame.

The scripts to process and analyze the images are downloadable from the repository: https://github.com/JuusolaLab/Hyperacute_Stereopsis_paper/tree/master/AnalyzeMovementData

When photoreceptors contracts photomechanically, generating a microsaccade (Fig. S15*A*), their rhabdomeres are expected to move both *laterally* and *axially* in respect to the ommatidium lens (11). Using a ray-traceable 3D computer graphics (CG) model (see Section II.2. Computer simulations of deep pseudopupil imaging, below), we simulated how these movement components should affect the optical DPP images and, thus, the actual DPP recordings *in vivo*. Simulations for rhabdomeres moving *laterally* predict that their virtual DPP images (10x-magnified by the ommatidial lens system) should also move laterally in proportion (Fig. S15 *B*, left and *C*). Similarly, simulations for rhabdomeres moving *axially*, away from the ommatidium lens, predict that in most cases their DPP image should darken (Fig. S15 *B*, right and *C*). But this depends on the rhabdomere tips’ starting position. Correspondingly in most cases, when rhabdomeres approach the ommatidium lens, their DPP image should brighten.

***Photoreceptor microsaccade lateral component*** The frame-by-frame DPP image series analyses of the actual *in vivo* high-speed recordings revealed how photoreceptors move laterally, quantifying their time-course and directions during microsaccades. Characteristically, in wild-type dark-adapted eyes, a bright flash evoked a maximal photoreceptor rhabdomeres displacement within ∼100 ms (the movement fast-phase) before returning more slowly to their original positions (the slower-phase) (Fig. S15*D*, left). These lateral movements were robust in individual eyes yet varied from fly to fly, ranging from ∼0.5 to ∼2.1 µm. Correspondingly, as the ommatidium optics project these movements to the visual space, the microsaccades shifted the photoreceptors’ receptive fields (RFs) ∼1.5 to ∼6.3°. Predictably, the control experiments using blind *norpA^36^*-mutants, in which faulty phototransduction (faulty Phospholipase-C) prevents photomechanical contractions from happening, showed no DPP microsaccades (Fig. S15*D*, right).

***Photoreceptor microsaccade axial component*** From the same DPP recordings, we estimated the microsaccades’ simultaneous axial component (Fig. S15 *D* and *E*), as a proportional photomechanical rhabdomere movement away and back toward the ommatidium lens. To eliminate motion artifacts, we measured this dynamic axial displacement in the DPP image pixels’ dynamic intensity change, tracking frame-by-frame only the pixels within the rhabdomere tips. We found the measured fast darkening and brightening dynamics in wild-type flies time-locking with their corresponding lateral DPP movement (Fig. S15 *D* and *E*, left). Because these two microsaccade components are synchronous, they should have the same photomechanical phototransduction origin.

As expected, the blind *norpA^36^*-mutants lacked the rhabdomere darkening/lightening dynamic completely (Fig. S15 *D* and *E*, right). Notably, these control flies also served a second purpose of eliminating the role of light-induced rhodopsin concentration changes affecting these dynamics. Because *norpA*- photoreceptors possess normally functioning rhodopsin-photopigments (Rh1-Rh5), having wild-type-like light-activation properties (74), the observed wild-type DPP darkening/brightening dynamics (Fig. S15 *D* and *E*, left) cannot result from rhodopsin-metarhodopsin photoisomerizations. Moreover, interestingly, the fast DPP intensity dynamics (as a prospective sign of axial rhabdomere motion), which co-occur with the fast lateral DPP jumps, are not seen during eye-muscle-induced whole retina movements (see Fig. S34 in Section III., below). Hence, collectively, this evidence maintains that the recorded DPP microsaccades were almost certainly photomechanical (i.e., generated by phototransduction alone).

**Fig. S15.**
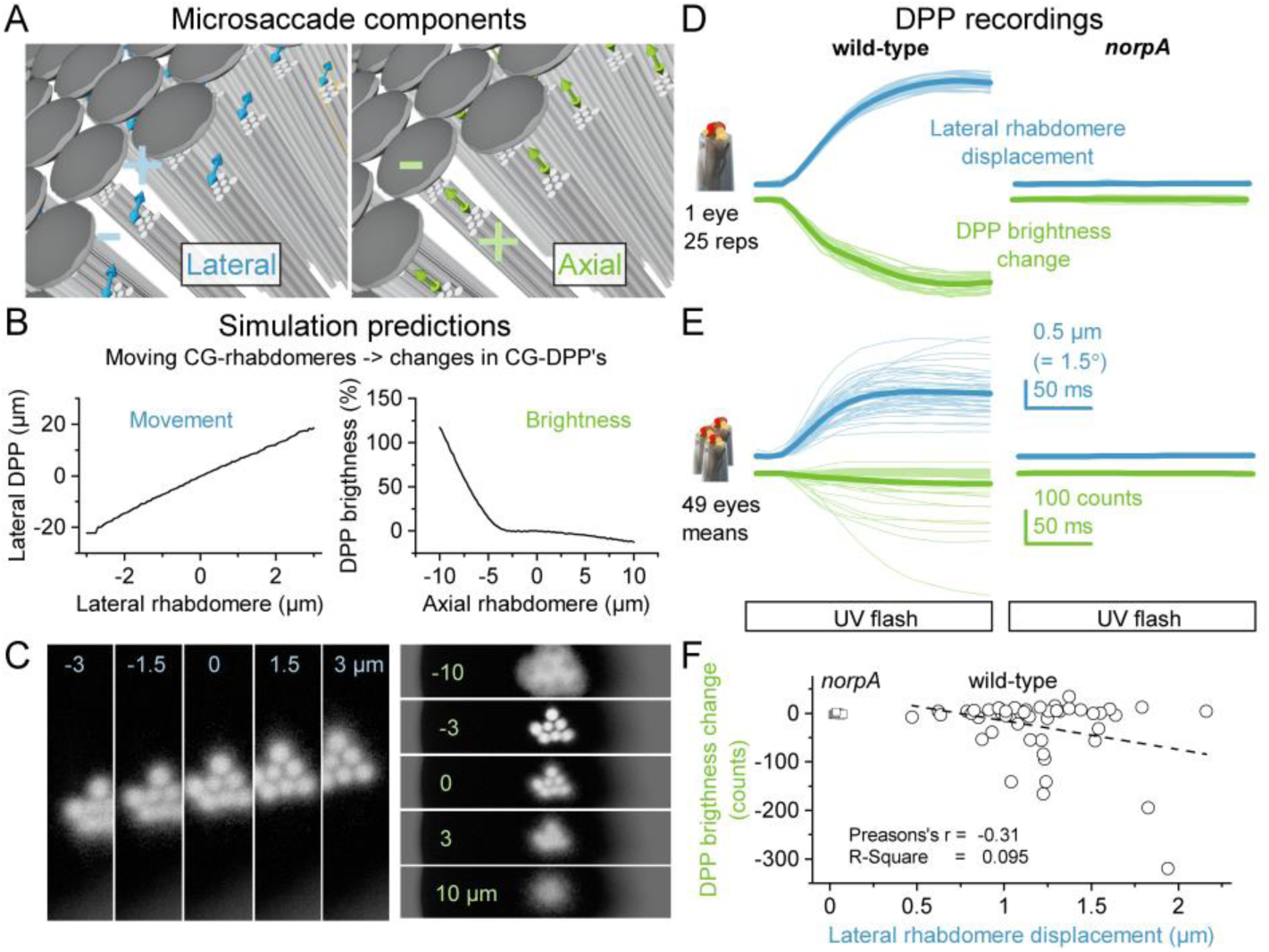
Axial and lateral photoreceptor microsaccade components. (*A*) Schematic of the photomechanical lateral (left) and axial (right) photoreceptor microsaccade components inside ommatidia. (*B*) Ray-traceable 3D CG *Drosophila* compound eye model (Section II.2, below) predicts that the ommatidium optics linearly translate lateral rhabdomere movements to deep pseudopupil (DPP) movements (left). Rhabdomere axial movements change the DPP brightness and size. However, this intensity-distance relationship is complex and depends on the initial rhabdomere resting position (distance) to the ommatidium lens. (*C*) Left: predicted DPP images when R1-R7/8 rhabdomeres are at different lateral positions inside ommatidia (zero is the center-position in *B* and *C*). Right: predicted DPP images when R1-R7/8 rhabdomeres are at different axial positions inside ommatidia (zero is the rhabdomere tip position at 21 µm from the ommatidial lens inner surface in *B* and *C*; see also Fig. S50 in Section V). (*D*) Examples of DPP microsaccades’ corresponding lateral (blue) and axial (green) components as recorded from an individual wild-type fly (left) and *norpA*-mutant. Thick lines give their means. (*E*) Averaged DPP microsaccades’ corresponding lateral (blue) and axial (green) components of many wild-type flies and *norpA*-mutants and their population means (thick lines). (*F*) DPP brightness changes plotted against the corresponding lateral R1-R7/8 rhabdomere displacement, as seen during the recorded DPP photoreceptor microsaccades. On average, the observed DPP darkens during a microsaccade.

The mean *axial* fast-phase photoreceptor microsaccade component - as averaged over the tested wild-type population - shows DPP darkening. This finding implies that the light-activation makes dark-adapted rhabdomeres, on average, move away from the ommatidium lens, which would typically make them collect light from a narrower angle (narrowing their receptive fields, RF). However, the data shows significant variations between individual flies (Fig. S15 *E* and *F*), with some recordings also indicating brightening, i.e., the rhabdomeres approaching the lens. Notably, this analysis is only suggestive, as we do not know the rhabdomeres’ actual axial resting position (at the start of the experiment) in any recording. Intriguingly, the physics dictate (see Fig. S50, Section V, below) that if the rhabdomere were “too far” (>22 µm) from the lens, they would collect light from a wider angle (see also (75)). In this somewhat counterintuitively case, to narrow their RF, the rhabdomeres should move toward the lens. Therefore, a plausible explanation is that the rhabdomeres’ axial resting position varies from one experiment to another, from fly to fly. The rhabdomere resting position, for example, could depend on the photoreceptors’ light/dark-adaptation state. Alternatively, it could be actively set by the flies’ internal (intrinsic) activity state, using the central synaptic feedbacks to the retina/lamina (30, 32, 36), or slow eye-muscle-induced axial drift (see Fig. S34F, below).

###### II.1.iv Mapping lateral microsaccade movement directions across the eyes in ♂and ♀ flies

Scanning the DPP microsaccades across the eyes revealed that their lateral (sideways) movement components, as measured at each corresponding left and right eye location, are mirror-symmetric, confirming the X-ray imaging results (see Section I., above). To further analyze factors contributing to their local dynamics, we performed a minimum error search by comparing the global microsaccade movement direction map to the corresponding global photoreceptor orientation map (Fig. S16). This analysis established that R1-R7/8 photoreceptors in most ommatidia across the eyes move collinearly back-and-forth approximately along the R1-R2-R3 rhabdomere orientation axis (Fig. S16D).

**Fig. S16.**
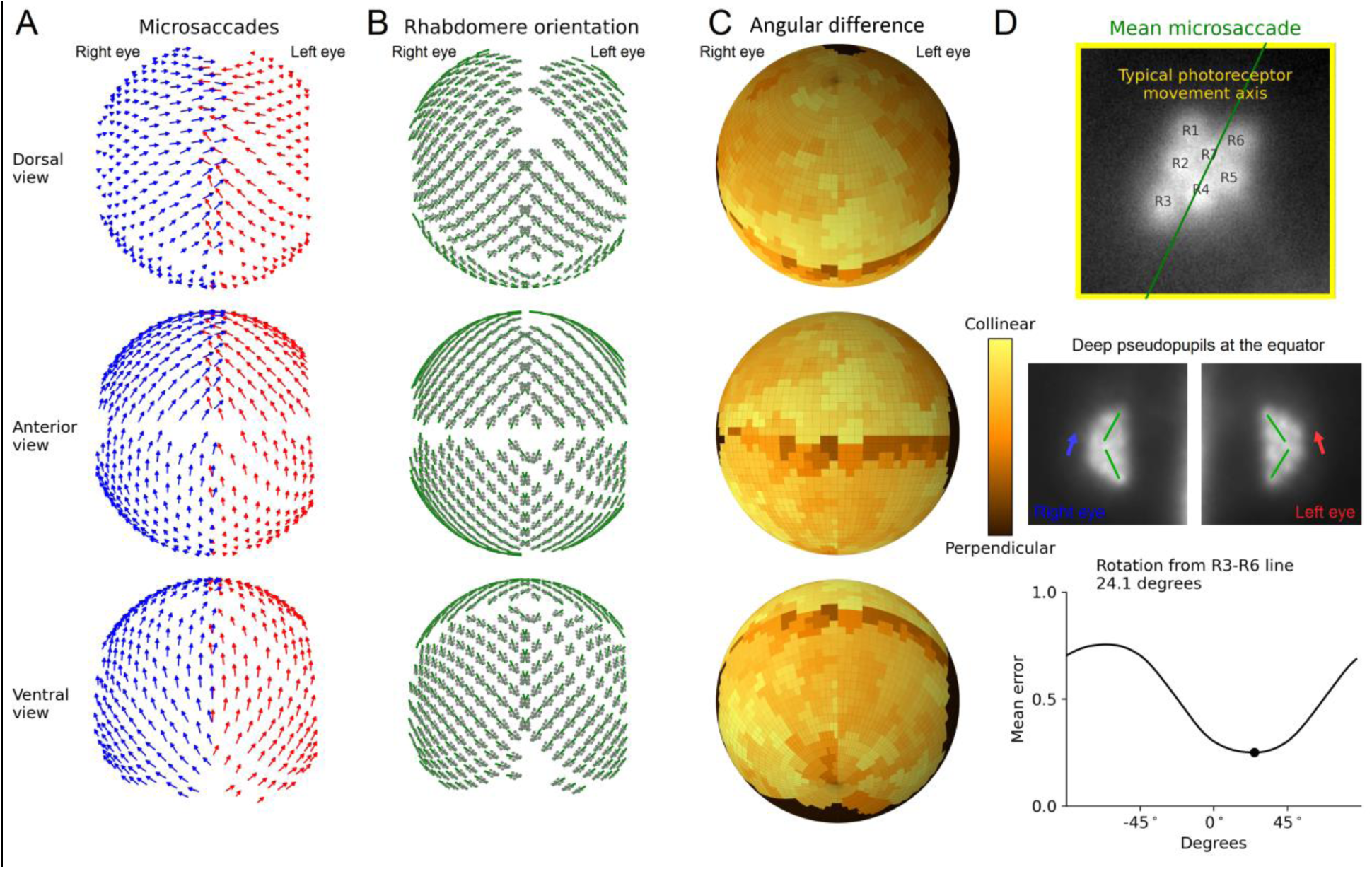
R1-R7/R8 microsaccades move along the R1-R2-R3 axis. The global microsaccade movement direction map is compared to the global rhabdomere orientation map. (*A*) R1-R7/R8 photoreceptor microsaccade movement directions of the right (blue) and left (red) eyes are mirror-symmetric. The microsaccade direction map shows the mean of 5 wild-type flies. (*B*) Local rhabdomere orientation patterns (gray) across the left and right eyes; plotted with the R1-R2-R3 axis (green), along which their microsaccades move (*cf*. the deep-pseudopupil image in *D*). The rhabdomere orientation map shows the mean of 5 wild-type flies. (*C*) The persistent match between the local rhabdomere orientation R1-R2-R3 axes and microsaccade movement axes verifies their approximately collinear alignment globally. In the equatorial deep-pseudopupil images, the upper and lower rhabdomere patterns fuse (inset), slightly obscuring their orientation calculation at the difference map’s equator. (*D*) During photoreceptor microsaccades, R1-R7/8 rhabdomeres move predominantly along the R1-R2-R3 axis (green; 28.6° rotation from the R3-R6 line, purple). This collinearity holds broadly irrespective of their eye position. Notice that the resulting mean error minimum (24.6%) is an overestimate because it also includes the slightly obscured equatorial comparisons in *C*. Dynamic representation of these calculations can be downloaded using the link above.

We further tested whether the male and female eyes’ microsaccades differ in movement dynamics and direction (Fig. S17). However, these analyses gave no clear evidence for visual sexual dimorphism.

**Fig. S17.**
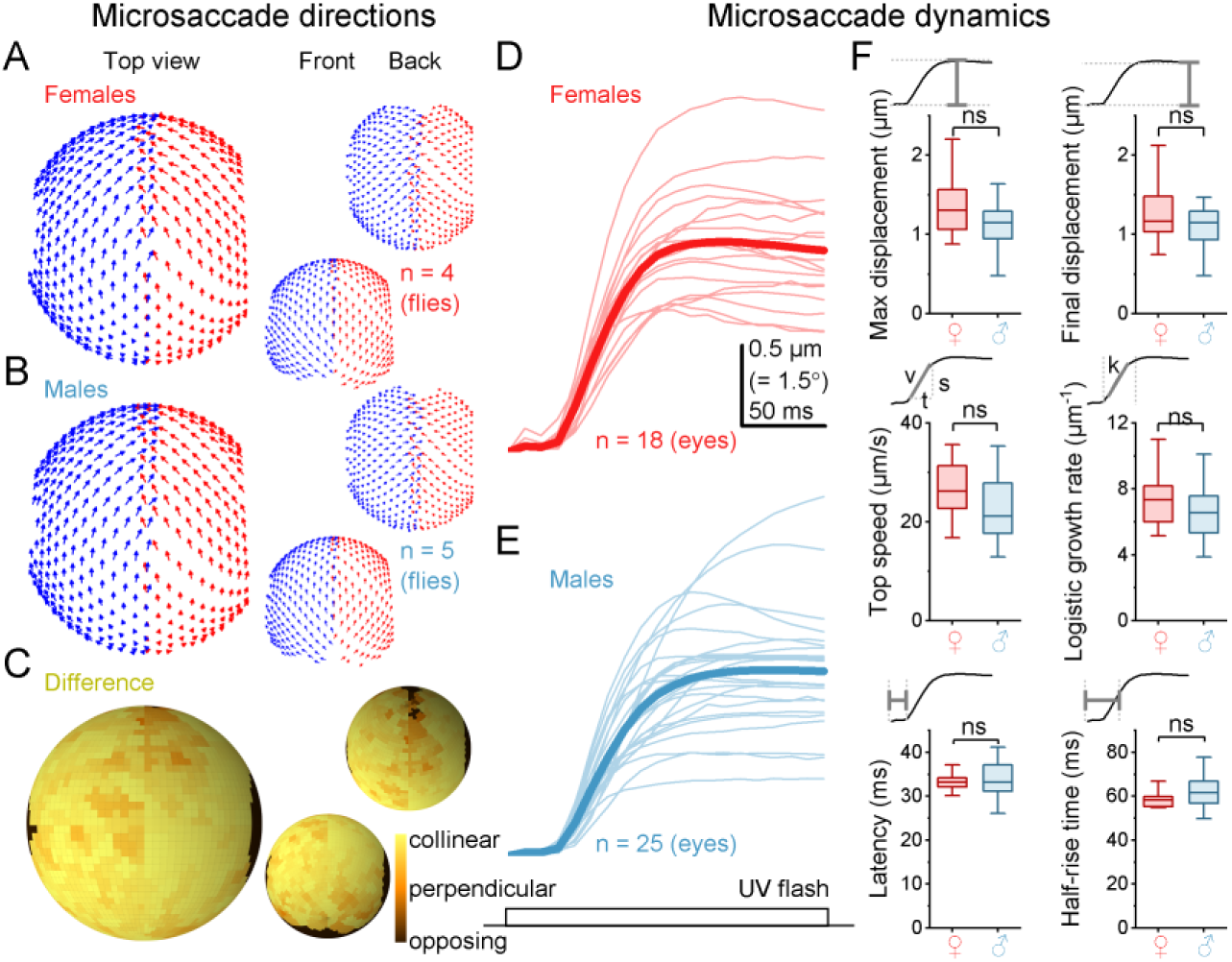
Male and female eyes have similar R1-R7/R8 microsaccade movement directions and dynamics. (*A*) ♀ microsaccade movement direction maps of the right (blue) and left (red) eyes, viewed from the top, front, and back. (*B*) ♂ microsaccade movement directions maps. (*C*) The female and male microsaccade movement directions are collinear, showing no apparent global or local differences (*D*) Microsaccade movement dynamics of 30 s dark-adapted female flies to a 200-ms-long bright UV flash. (*E*) The corresponding photoreceptor microsaccade movement dynamics of male flies. (*F*). The female and male eyes’ microsaccade dynamics are similar for the six tested metrics, indicating no apparent sexual dimorphism in visual information sampling.

Collectively, these results (Fig. S11 to S17) strongly suggest that:

- The lateral microsaccadic movement component inside each ommatidium happens along some structural (developmentally-set) lowest resistance (energy minimum) R1-R7/8 anchoring.
- The photoreceptor microsaccades are similar in ♂and ♀ flies.
- These movements were practically free of spontaneous intraocular muscle activity, which otherwise would have distorted their local and global mirror-symmetry.

A video showing the analyses is downloadable from: https://github.com/JuusolaLab/Hyperacute_Stereopsis_paper/tree/master/AnalyzeMovementData

##### II.2 Computer simulations of deep pseudopupil imaging

Computer simulations were used to test how the microscope system’s numerical aperture (NA) affects the infra-red DPP imaging; especially, how the NA influences the number of ommatidia contributing to the DPP image and how a high NA can lead to overestimation of the binocular overlap.

A microscope system’s numerical aperture (NA) is a dimensionless number that characterizes the range of angles it can accept light. For the infra-red DPP imaging, NA optically limits the ommatidial area, wherein the optically superimposed rhabdomeres can be pooled into the pseudopupil image. Most stereomicroscopes with their long-working distance objectives typically have relatively low NAs (≤∼0.2). The NA is defined as

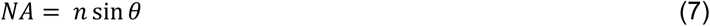

where *n* is the index of refraction (IOR) for the used immersion medium (n=1 in the air), and *θ* is the half-angle subtended by the microscope lens at the viewed object (76). In binocular overlap, a simple geometrical consideration suggests that the overlap can be theoretically overestimated by 2*θ*. In practice, however, the left-eye-right-eye symmetry during the horizontal rotation is such that the circular aperture collects less light from the horizontal extremes of the entrance pupil, and these extreme or high order light rays contribute relatively little to the formed image.

The f-number or the f-stop, *N*, is defined as:

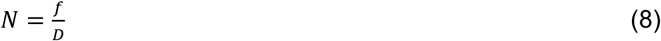

where *f* is the focal length, and *D* is the used objective’s effective aperture (entrance pupil diameter). The image depth of field increases with f-number. For a point-like object at distance *d* from the entrance pupil, it follows from Eq. 7 and Eq. 8 that

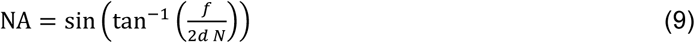

We used this equation to calculate the numerical apertures in the computer simulations.

The computer simulations were implemented as a ray-traceable 3D computer graphics (CG) model of the fly eyes capable of producing the DPP as an optically emergent feature the same way the real fly eyes do. The CG-eye model was fully parametric and script initialized, making it easy to translate the model for other insect species, for example. Another advantage of the CG approach is that because the 3D models are primarily collections of numerical data about the vertices and faces, they are naturally independent of the rendering engine or the modeling software. Therefore, it is relatively easy to import the CG-eye model into any other software.

The CG-eye model’s main building block was a simplified ommatidium with a facet lens, cylindrical R1-R7/8 rhabdomere tips, a basement membrane segment, and simplified screening pigments (Fig. S18*A*). The CG- ommatidium was generated using the open-source graphics software Blender 2.8 (https://www.blender.org/) and its built-in Python interface for scripting. The facet lens was modeled as a double convex lens with a lens diameter of 16 µm and a curvature radius of 11 µm, and a lens thickness of 8 µm, as described before (75). The rhabdomere tips were 3 µm long, simplified circular cylinders with a µm diameter for the R1-R6 and a 1.0 µm diameter for the central R7/8, placed on the retinal plane locations quantified from a retinal electron micrograph (Table S1). The rhabdomere tips were placed 21 µm apart from the facet lens center point. We modeled the screening pigments as a hollow, thin-walled hexagonal cylinder with a 16 µm radius, spanning from the lens to the basement membrane. Finally, the basement membrane segment was modeled as a thin hexagonal plate with a 16 µm radius.

**Fig. S18.**
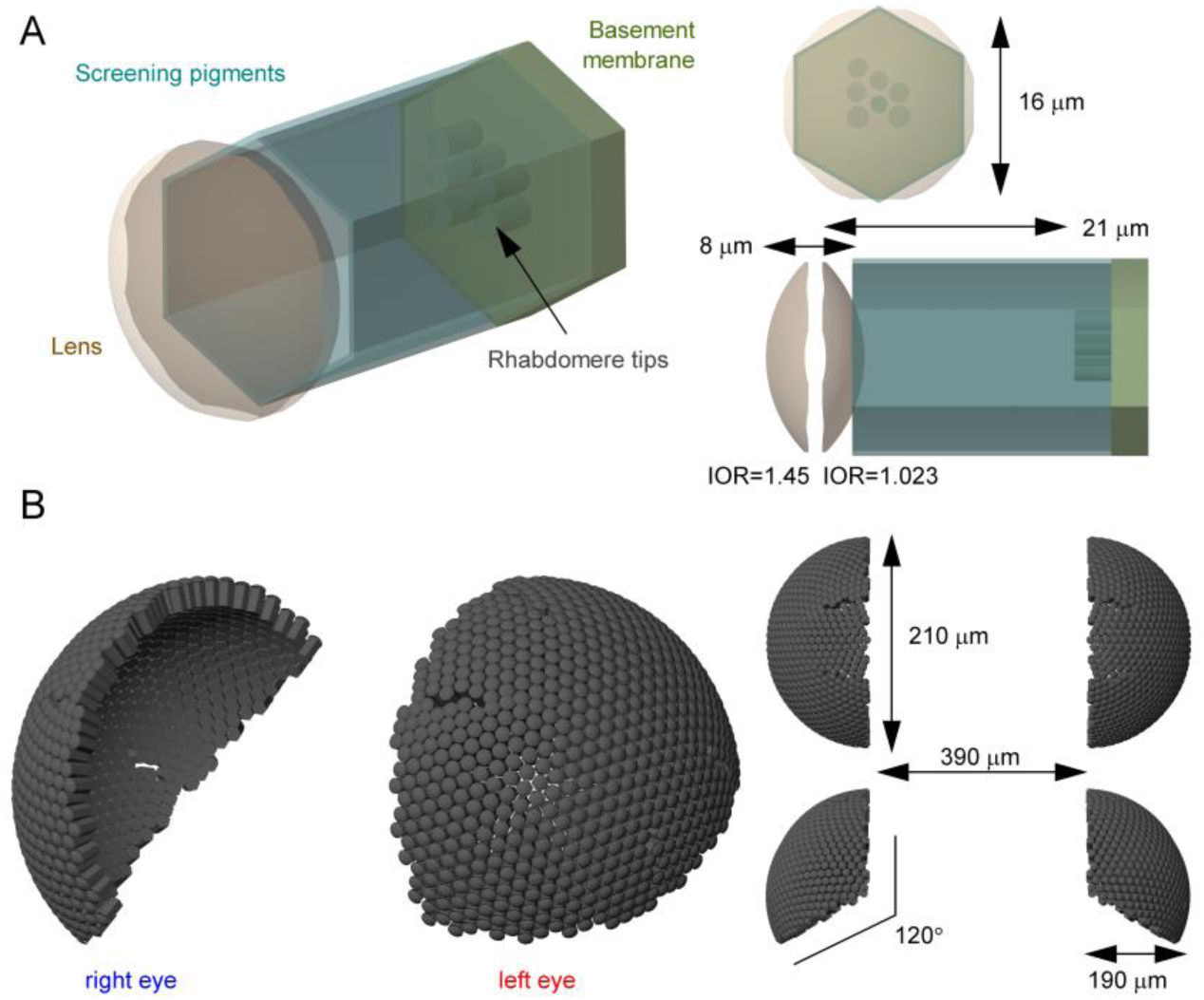
The *Drosophila* compound eye CG-model schematic orthographic view, as rendered by the Blender Workbench Engine. (*A*) The simplified ommatidium forms the basic building block of the CG eye model. The lens is configured as a glass material, the rhabdomere tips white light-emitting material, and the pigments and the basement membrane as black matte material. (*B*) The CG-eye model consists of 1,400 ommatidia evenly distributed on two semi-spherical surfaces so that their optical axes intersect at the center point of the respective eyes.

**Table S1.** Rhabdomere (x, y) locations in the CG-model’s retinal plane (see Fig. S47A).

| Rhabdomere | x (μm) | y (μm) |
| --- | --- | --- |
| R1 | -1.6881 | 1.0273 |
| R2 | -1.8046 | -0.9934 |
| R3 | -1.7111 | -2.9717 |
| R4 | -0.0025 | -1.9261 |
| R5 | 1.6690 | -0.9493 |
| R6 | 1.6567 | 0.9762 |
| R7/8 | 0.0045 | -0.0113 |

To proceed from one ommatidium to many, we evenly distributed a realistic amount of the CG-ommatidia across two skewed semi-spherical surfaces with a long radius of 210 µm and a short radius of 180 µm, that were 390 µm apart from each other’s center points (Fig. S18*B*; Fig. 1*C*). The ommatidia are the most parallel at the left and right eyes’ medial edge, where binocular overlap occurs. Therefore, here, the outwards projected ommatidial optical axes of the eyes never intersect but diverge. Furthermore, the inferior eye edge, adjacent to the thorax, was defined by the principle that no ommatidial axis should make an angle larger than 120° from the top in the coronal plane. Finally, we also considered the dorsal-ventral midline, where the rhabdomere pattern on the dorsal side appears as a mirror version of the ventral side and vice versa. Overall, this somewhat simplified eye assembly led to a quite realistic outcome.

To simulate light propagation in the model, we used the (physics-based, unbiased) ray-tracing render software LuxCoreRender 2.4 and its Blender plugin BlendLuxCore (https://luxcorerender.org/). The render engine successfully simulates light refraction on the facet lenses leading to the DPP virtual image formation under the right viewing conditions. We configured the material output node for the facet lenses as a glass material. We used the index of refraction (IOR) of 1.450 for the outer lens surface and 1.023 for the lens inner surface to match the real IOR values of 1, 1.45, and 1.34 for the air, lens, and crystalline cone volumes (75). The rhabdomere tips were configured as white matte material with white light emission to mimic the antidromic illumination. The screening pigments and the basement membrane hexagons were configured as matte material of absolute black to absorb any incident light. Bidirectional ray tracing with the Metropolis sampler and a 3-samples-per-frame halt condition was used for rendering. To observe the CG DPP, we enabled the camera’s depth of field option with a sufficiently small f-stop value and set the focus at the converging point of the ommatidial axes.

To illustrate the rhabdomere or DPP microsaccades, we used real microsaccade direction data acquired in the goniometric DPP light-flash experiments. For each CG-ommatidium, we used the nearest microsaccade direction available in the dataset to set the animation start and end locations using the programmable keyframe animations in Blender. In some of the images and videos, we also used blue and red beams projecting from the rhabdomere plane to illustrate how the contralateral receptive fields move and intersect during microsaccades. This effect was achieved by placing two spotlight sources, each with a 45° emission angle, in the rhabdomeric plane of two contralateral R6 rhabdomeres. Also, a light scattering volume was added outside the eyes to make the beams visible.

The CG eye model is publicly available at the git repository: https://github.com/JuusolaLab/Hyperacute_Stereopsis_paper/tree/main/CG-Compound-Eye

In the NA simulations, we systematically changed the virtual imaging system’s NA and f-number to survey how these parameters:

- Contribute to optical pooling the ommatidial rhabdomeres’ DPP images.
- Affect the eyes’ binocular range estimates.

In the NA binocular overlap simulation, the camera was placed 10 mm apart from the eyes’ center point and set to have a focal length of 10 mm. We used F-stop values of 3, 10, 30, 100 and 300 that correspond to NAs of 0.164, 0.0499, 0.0167, 0.00500 and 0.00167, respectively (Eq. 9). During video rendering, the camera was slowly rotated horizontally from -45° to +45° as in the binocular overlap estimation experiments. A video showing the analyses is downloadable from: https://github.com/JuusolaLab/Hyperacute_Stereopsis_paper/tree/main/CG-Compound-Eye

In the numerical aperture (NA) simulation (Fig. S19), we illuminated only selected CG-model ommatidia and observed the emerged DPP pattern when viewed with a high NA and a low NA microscope. In Blender, both cameras were configured to an f-stop value of 1.0, and they were 1 mm away from the eye’s center point, at which the camera focus was set. We varied the focal length parameters to change the NA. The high NA camera had a focal length of 0.5 mm, and the low NA camera had a focal length of 0.1 mm. These values correspond to NAs of 0.243 and 0.0499, respectively (Eq. 9).

**Fig. S19.**
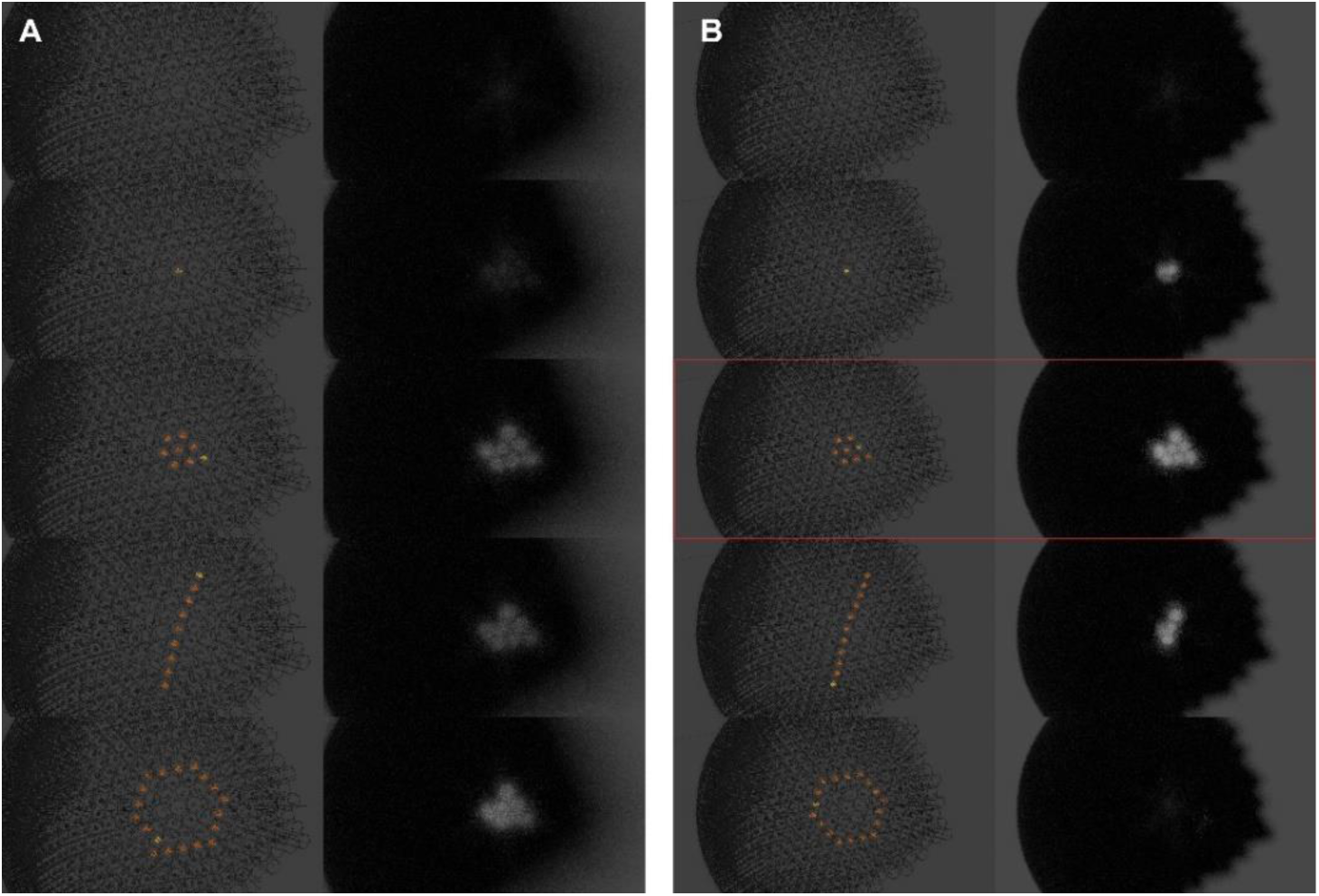
Simulating deep pseudopupil imaging in the *Drosophila* eye. (*A*) A microscope system with a high numerical aperture (NA) collects light from a wide angle. Therefore, it will form the best DPP image of optically superimposed R1-R7/8 rhabdomere endings in the seven neighboring ommatidia. But it can also generate lower quality images of R1-R7/8 rhabdomeres of a single ommatidium (orange dot) or those, which are optically pooled along with specific arrangements (*e.g.,* along an orange line or hexagon) of more distant ommatidia. (*B*) A microscope system with a low NA collects light from a narrow angle. Thus, it will only generate a complete DPP image from the optically superpositioned R1-R7/8 rhabdomere endings in the seven neighboring ommatidia. Red rectangle: a typical stereomicroscope with a relatively low NA (<0.2) would only collect a DPP image from neighboring seven ommatidia.

These simulations (Fig. S19) suggest our stereomicroscope (NA of 0.11) would have primarily pooled the DPP images of the optical superposition R1-R7/8 rhabdomeres from the seven nearest neighbor ommatidia (Fig. S19*B*, middle). Therefore, the estimated stereo vision range and rhabdomere orientation maps are likely to be accurate, not over- or underestimates biased by this new high-speed imaging method and its instrumentations’ physical limitations.

##### II.3 ERG recordings

Head-fixed *Drosophila*, either inside a pipette-tip or tethered to a small hook (see Section II.6., below), were connected to the center of a custom-made electrophysiological setup (60, 77). Blunt (low resistance) filamented borosilicate glass microelectrodes (0.5 mm inner and 1.0 mm outer diameters) filled with fly Ringer (containing in mM: 120 NaCl, 5 KCl, 10 TES, 1.5 CaCl_2_, 4 MgCl_2_, and 30 sucrose) were attached to electrode holders (containing a chloridized silver wire) and connected to a microelectrode amplifier (model SEC-10L; npi Electronic, Germany). (20). Using micromanipulators, we carefully placed the recording electrode on the eye and the reference electrode elsewhere on the fly head. Using the setup’s Cardan-arm system, we fixed the fiber-optic-end of the LED light source in a predefined x,y,z-position above the fly head, directly stimulating the eye’s anterior-dorsal part. The eye’s voltage responses were then recorded to 1-s-long bright Green (546 nm) and UV (365 nm) pulses separately.

##### II.4 Microsaccade and ERG recordings from the same flies

We tested whether the used fly head immobilization methods affect the fly eyes’ DPP microsaccades and ERG responses to the UV- and green test light flashes (Fig. S20). To ensure ocular recording stability, 3- to-10-days-old *Drosophila* were either:

- affixed inside a pipette-tip or a metal holder cone from the head cuticle and proboscis (20, 60)
- tethered to a small hook from the head/thorax’s dorsal side, similar to the flight simulator experiments (22), except that here their legs and wings were immobilized by waxing.

If performed correctly, the tethering method avoided any mechanical stress to the eyes resulting from pressure experienced while being pushed through the pipette/cone. Nevertheless, we used the pipette-tip fixation method for most experiments because it was easier and faster to perform and effectively reduced sporadic muscle-induced retinal movements (11).

Either way, practice improved the microsaccade recording success rates, which for the wild-type flies approached 100%. Yet, for specific transgenic flies and mutants, such as the UV-flies and *hdc^JK910^*, the rates were consistently lower for the pipette-restrained than tethered flies. Therefore, we conclude:

- The *in vivo Drosophila* preparation is structurally fragile to mechanical stress, with genetic manipulations/mutations reducing its eyes functional integrity to generate photomechanical microsaccades’ *lateral component* (sideways movement)
- The observed fly-to-fly amplitude variations in their microsaccades’ *lateral component* (Fig. S20*B* and Fig. S21*A*) must, in part, reflect the preparation quality. But it may also partly signify synaptic feedback strength (20, 33, 35, 36, 78) - top-down signaling from the brain (42), reflecting each fly’s intrinsic activity state or attentiveness during the experiments. Thus, for example, *dSK*-mutants’ intracellular R1-R6 voltage responses are faster and smaller than wild-type flies because they receive tonic feedback overload from visual interneurons (78, 79). Correspondingly, their photoreceptor microsaccades are also faster and smaller (see Section II.8.i. and Fig. S29*B*, below).
- The photomechanical microsaccades’ *axial component* is more robust against mechanical stress, as it is readily observed *ex vivo*, even in fully dissociated ommatidia (10, 11).

In contrast, the ERG responses of the same flies, as recorded separately from both their left and right eyes (Fig. S20*C*), showed invariably characteristic extracellular voltage responses to the test flashes (practically 100% success rate), irrespective of whether their eyes showed the microsaccadic sideways movement or not. This finding is consistent with the hypothesis that the microsaccades’ lateral component requires interommatidial rhabdomere pivoting and mechanical coupling (such as tip-links; see Section II.8., below).

Together the microsaccade and ERG recordings showed that the dark-adapted pipette-tip-fixed and tethered flies - having their legs and wings immobilized by beeswax - generated similar (equally strong) photoreceptor responses to temporal light pulses (Fig. S20 *B* and *C*, Fig. S21 *A* and *B*). Some suggestively larger ERGs were measured from a few individual tethered flies with mobile legs and wings. This finding is consistent with the earlier observations about extracellular neural activity differences (local field potentials and spiking) in the *Drosophila* visual system during resting and flying (42), but we did not investigate it further here.

Crucially, the microsaccade and ERG amplitudes scaled with the number of light-activated photoreceptors within an average ommatidium, being the largest in the wild-type eyes when all R1-R7/8 were activated (Fig. S20 *B* and *C*, Fig. S21 *A* and *B*). This strong correspondence means that both the microsaccade and ERG responses would directly signal the underlying photon sampling and phototransduction processes. These results made it very likely that photoreceptor microsaccades would have also happened at least equally well during the *in vivo* two-photon Ca^2+^-imaging (see Section III, below) and flight simulator experiments (see Section VI, below). In both of these approaches, we used tethered flies without waxing their wings and legs, thereby providing them with a higher degree of mobility.

**Fig. S20.**
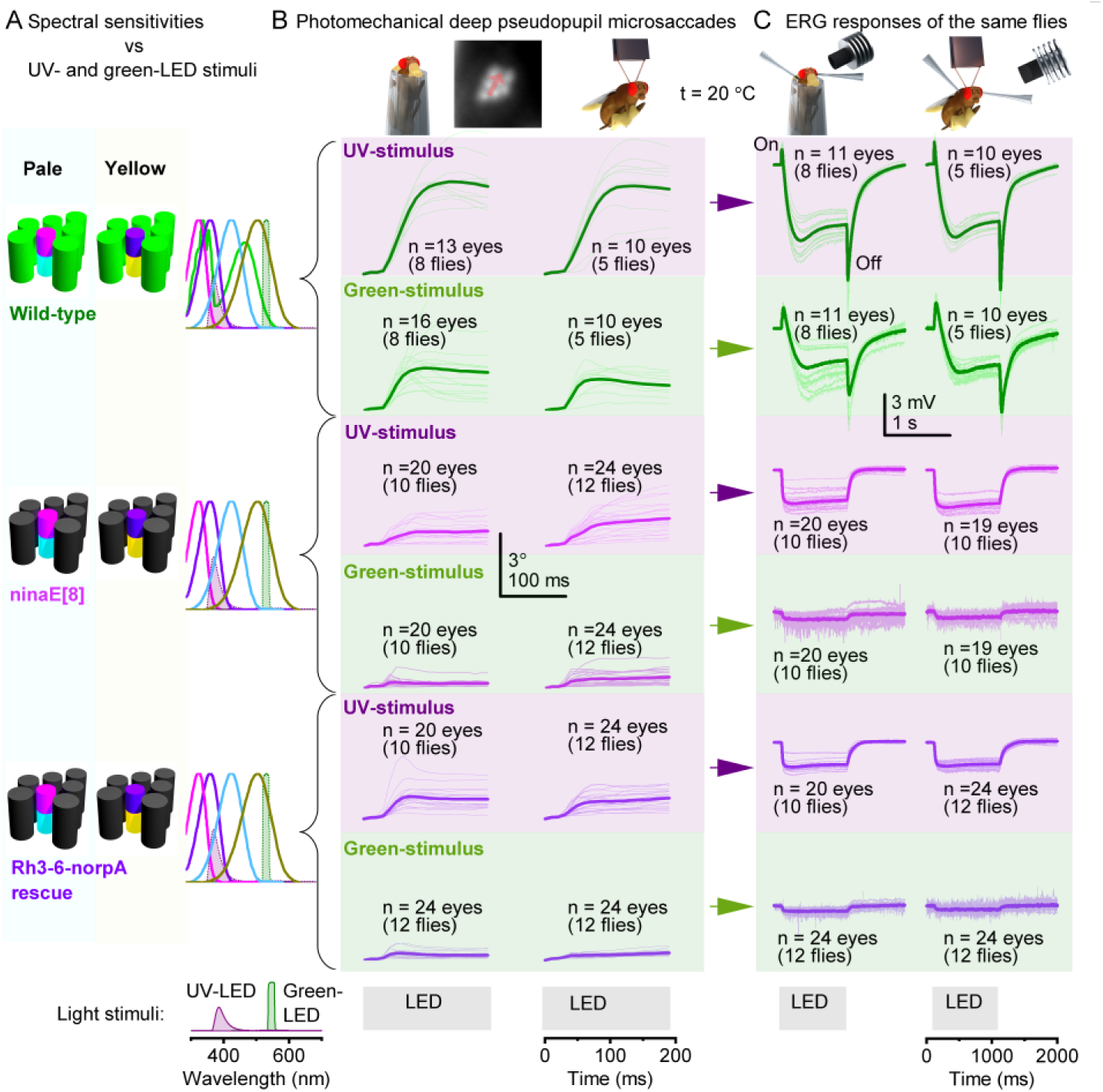
Dark-adapted pipette-tip-fixed and tethered *Drosophila* generate equally strong microsaccades and ERG responses to light pulses. (*A*) Wild-type flies’ ommatidia come with two different spectral compositions. The outer R1-R6 photoreceptors express blue-green rhodopsin Rh1, while the inner R7/R8 are either the pale or yellow type. Their respective rhabdomeres and spectral sensitivities (nomograms; co-colored) are shown against the test UV- and green LEDs’ spectral emission (filled curves). In *ninaE^8^* and *norpA* Rh[3, 4, 5, 6] rescue flies, the outer R1- R6 photoreceptors in the ommatidia (dark gray rhabdomeres) are blind while the inner R7/8 photoreceptors maintain their light-sensitivities. (*A*) Because the UV- or green-stimulation overlapped with the tested photoreceptor classes’ spectral sensitivities, it light-activated the imaged R1-R7/8 rhabdomeres, causing them to bounce sideways along their eye-location-specific movement axis. These microsaccades were larger in the wild-type flies - with all R1-R8 functioning - than in the mutant flies having only their R7/R8s functional, suggesting that the photoreceptor movements summed up photomechanically. Each fly’s microsaccade dynamics were calculated by cross-correlating the consecutive image frames in 10 ms resolution, shown for 5-12 flies (thin traces; from both left and right eye images) and their average (thick traces). Because R7/R8 light-activation alone also moved the blind R1-R6 in unison, R1-R8 rhabdomeres must be mechanically coupled/pivoted in each ommatidium, possibly by anchoring and ultrastructural links; see also Fig. S28 and Fig. S29, below. Overall, the photoreceptor microsaccades of pipette-tip-fixed and tethered dark-adapted *Drosophila* showed similar dynamics. (*C*) The same flies’ electroretinograms (ERGs) to the UV- and green-LED stimulation showed the predicted spectral sensitivities. The wild-type ERG verified the R1-R6 photoreceptors’ normal phototransduction/synaptic signaling (on- and off-transient (19, 22, 36) and the DPP (*B*) movements’ photomechanical origins. Predictably, the on- and off-transient of *ninaE^8^* mutants and *norpA* Rh[3, 4, 5, 6] rescue flies were greatly diminished (22). Overall, the ERGs of pipette-tip-fixed and tethered *Drosophila* showed similar dynamics. (*B* and *C*) Because the UV-light activated more photoreceptor types than the green light (*cf*. their nomograms in A), both the UV-microsaccades and UV-ERGs were larger for all tested flies than the green ones. The ERG light stimulation was ∼10-fold weaker than the stimuli in pseudopupil experiments, measured by a spectrometer. The tethered *Drosophila* had wax-restrained legs and wings.

**Fig. S21.**
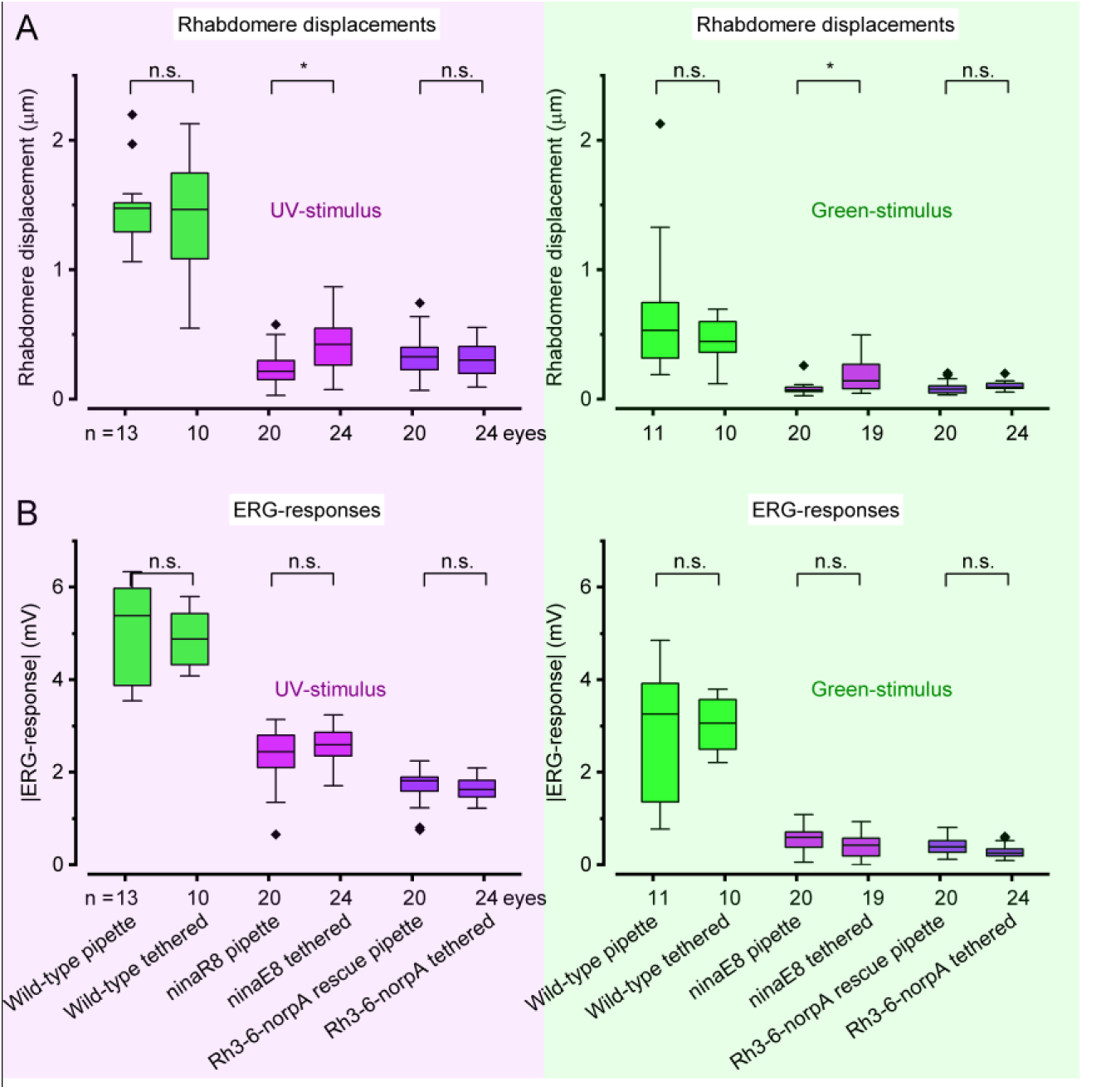
The deep pseudopupil (DPP) microsaccade and ERG response statistics for the pipette-tip-fixed and tethered wild-type (R1-R6 and R7/8 functional), *ninaE^8^* (R7/8 functional), and Rh3-6-*norpA* rescue flies (R7/8 functional). (*A*) DPP microsaccades are given as rhabdomere movements inside the ommatidia. Remarkably, for a bright UV-light pulse (left), an average wild-type R1-R6 rhabdomere moved photomechanically sideways about its average width (see Section V.8., below). In visual space, this corresponds to its receptive field jumping ∼5.3°. These displacements were smaller for the green-light pulses (right), matching the directly measured rhabdomere movements to blue-green light inside single ommatidia (see Section II.8ii. and Fig. S32*E* below). Because the UV-light activated (above) more photoreceptor types - and many of them (such as R1-R6) more intensely - than the green light (below; *cf*. their nomograms in Fig. S20*A*), the UV- microsaccades were larger for all tested flies than the green ones. (*B*) Correspondingly, the UV-ERG (left) photoreceptor components – *i.e.,* with the on- and off-transients excluded - were larger than those of the green-ERGs (right) for all the tested flies. (*A* and *B*) For each tested fly, its microsaccade amplitudes scale directly with its ERG amplitudes for both the UV- and green-stimulation: the larger the microsaccades, the larger the ERG responses. See also Section II.8 with Fig. S28 and Fig. S29, below. Only for *ninaE^8^*, both the UV- and green-microsaccades were larger in the tethered flies, but this was not seen in their ERG responses. For all other genotypes, the head-fixation methods made no difference in their photoreceptor responses. One-way ANOVA, comparing the pipette-tip-fixed and tethered flies for each genotype, using posthoc Tukey.

##### II.5 Separating photoreceptor microsaccades from eye-muscle activity

When monitoring the wild-type and mutant flies’ DPPs, one sees - from time to time - them shifting position or moving slightly, caused by intraocular muscles nudging the whole retina around (13) (see Section III. High-speed optical imaging of eye-muscle-induced whole retina movements and antennae castings, below). While this intrinsic activity (11, 13, 14) likely contributes to *Drosophila*’s active gazing strategy (13, 14) and spatial awareness, it is mechanistically separate from the local photomechanical photoreceptor microsaccades (11) and can interfere with the microsaccade recording. Fortuitously, immobilizing a fly - with the beeswax cross-bridging its head and stretched proboscis to the pipette/holder rim (11, 60, 77) - reduces intraocular muscle activity, in many cases keeping spontaneous retinal movements few and far apart. When carefully prepared, most pipette-restrained flies showed highly reliable and consistent photomechanical microsaccades. Moreover, similar to the tethered fly recordings, because the microsaccades were precisely timed to the light input, we could afterward (if needed) exclude any traces with spurious dynamics attributable to intermixing intraocular muscle activity. Thus, those odd (very few) recordings, which showed intraocular muscle activity parallel with photomechanical photoreceptor contractions, were disregarded from the analyzed data.

Importantly, since the microsaccades of the synaptically-decoupled (Fig. S2*D*), and thus behaviorally blind (see Section VII.6, below), *hdc^JK910^* control flies followed the wild-type-trajectories (Fig. S22), the observed dynamics (Movie S4) did not involve intraocular muscles. These results further concur with the corresponding wild-type and *hdc^JK910^* X-ray microsaccade imaging results (see Section I.3, above).

**Fig. S22.**
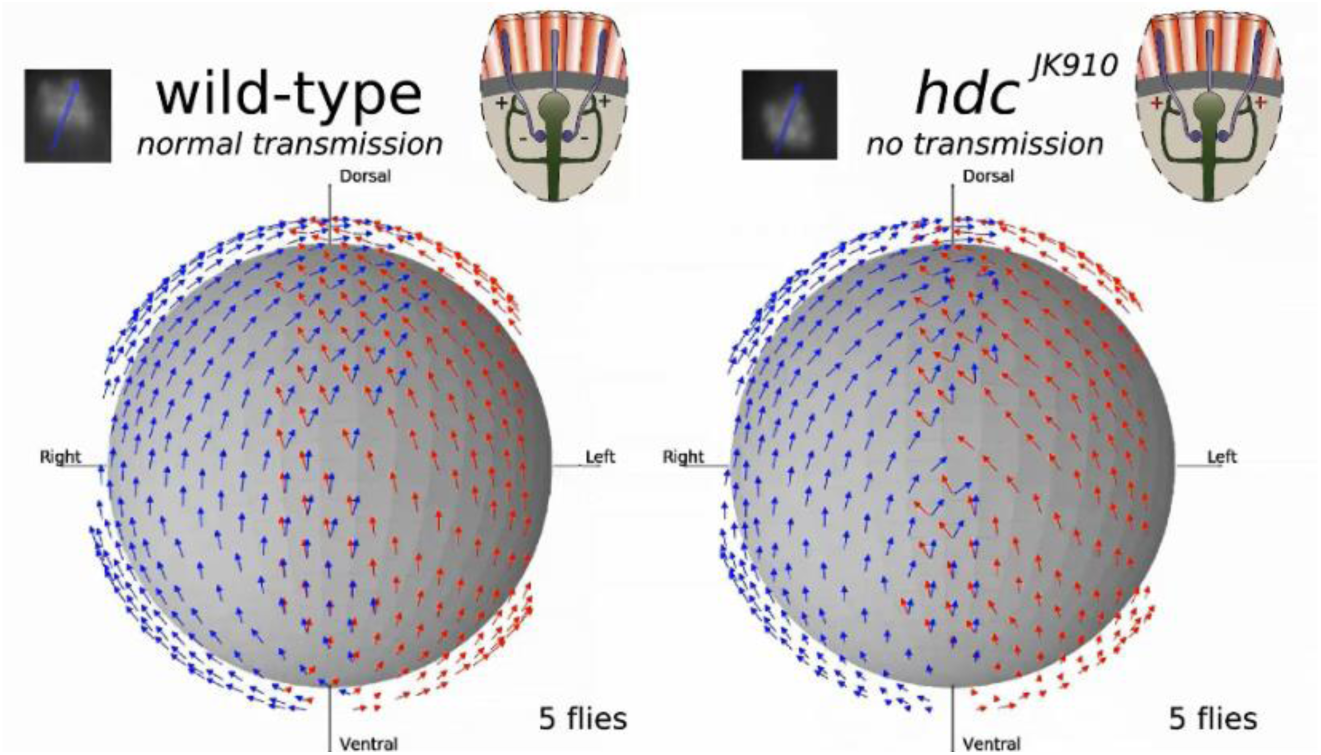
The left (red arrows) and right eye (blue) photoreceptor microsaccade movement trajectories of wild-type (left) and *hdc_JK910_* (right) flies match. Because *hdc_JK910_* photoreceptors lack neurotransmitter histamine, they cannot synaptically convey light information directly or indirectly to downstream motor neurons. Therefore, these movements must lack any intraocular muscle components. The movement trajectories were calculated through image cross-correlation from light-triggered high-speed DPP motion video recordings. Markedly, the left and right eye’s photoreceptor microsaccades are mirror-symmetric, reflecting the eyes’ mirror-symmetric gross anatomy and ommatidial ultrastructure. Each map shows the mean microsaccade directions of 5 pipette-tip fixed flies.

##### II.6 Measuring photoreceptor microsaccade frequency response

We have previously shown that bright pulses (of the same intensity increment) evoke equally large microsaccades in both dark-adapted and brightly light-adapted photoreceptors and that the microsaccades follow bursty light intensity changes reliably (11). These results established that photoreceptor adaptation enables microsaccadic light input modulation over a broad range of environmental lighting conditions (11). Here, we further assessed how fast stimulus contrast changes - light increments (positive contrasts) and decrements (negative contrasts) – the photoreceptor microsaccades could follow in light adaptation and whether their positive and negative contrast response dynamics differ *in vivo*.

Using pipette-tip-fixed wild-type *Drosophila* (see Section II.1., above), we first light-adapted those local photoreceptors, contributing to the deep-pseudopupil image, to a bright continuous UV-light background; estimated emission intensity >10^7^ photons/s/photoreceptor. Because an R1-R6 photoreceptor has ∼30,000 phototransduction units (microvilli), each of which samples incoming photons with refractory dynamics, this light background should result in ∼5 x 10^5^ quantum bumps/s, steady-state-depolarizing the photoreceptors ∼30-35 mV above their dark resting potential (11, 40). Then, using high-speed DPP imaging (200 fps), we recorded these optically superpositioned photoreceptors’ microsaccade responses to specific point-source stimuli (Movie S5), in which sinusoidal or pulsatile +/-1 contrast modulation frequency either accelerated in time (Fig. S23 *A* and *B*) or was constant (Fig. S23*C*). Thus, the microsaccades were evoked by temporal contrast changes at their RF center, delivered through the microscope optic. Finally, we established the microsaccades’ frequency response function by measuring and analyzing these photomechanical responses for the accelerated temporal contrast modulation frequency.

The temporal contrast modulation stimuli evoked strong photoreceptor microsaccades with explicit biphasic behavior (Fig. S23). The microsaccades’ activation phase to positive contrasts (light increments) was significantly faster than their recovery phase to negative contrasts (light decrements), generating characteristic “jump-and-recoil” responses, with the quick “jumps” dominating their waveforms. These dynamics were superimposed on a gradual ∼8-second-long photomechanical contraction creep-up until the responses became too small to be reliably cross-correlated from the high-speed video (as limited by the imaging systems’ signal-to-noise ratio). At that point, the photoreceptors’ contraction creep-up also began to wane. Overall, the microsaccades followed both sinusoidal and pulsatile contrast frequencies up to 27- 32 Hz, with the reliably detectable response amplitudes varying from one fly preparation to another, having an average 3 dB cut-off frequency of about 12.5 Hz (Fig. S23 *A* and *B*).

**Fig. S23.**
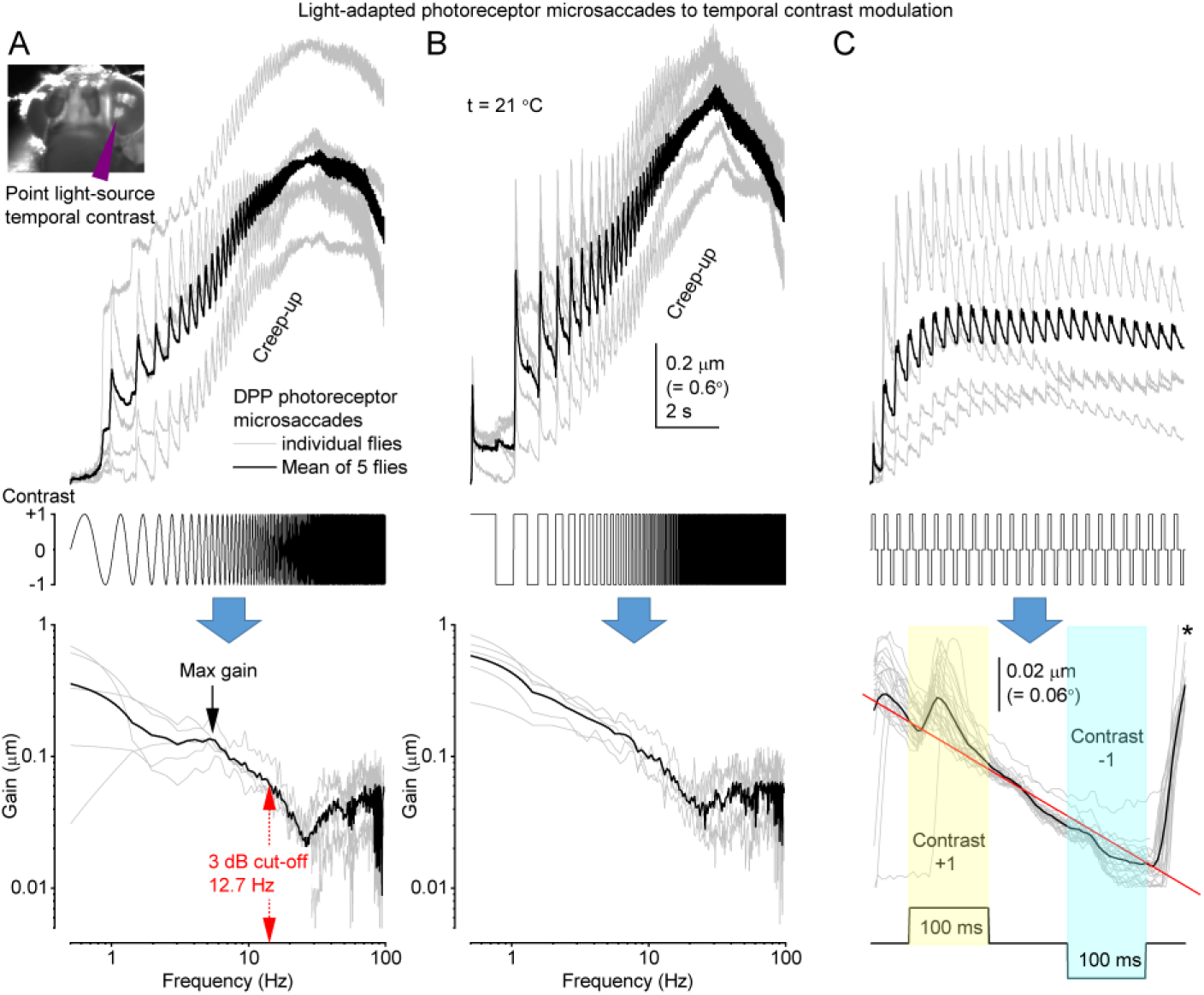
Light-adapted deep pseudopupil microsaccades reliably follow fast temporal luminance contrast changes of a stationary point source. (*A*) Microsaccades to frequency accelerated sinusoidal +/-1 contrast stimulus (above). Their frequency response (below) indicates that microsaccades can follow up to ∼30 Hz follow modulation (below), with the mean 3 dB cut-off at 12.7 Hz. (*B*) Microsaccades to frequency accelerated pulsatile +/-1 contrast stimulus. Microsaccades can follow up to ∼30 Hz follow modulation, showing accentuated (jump-like) dynamics even to very brief positive contrast changes. (*C*) Microsaccades to repeated 100-ms-long +/-1 contrast pulses. While the microsaccade amplitudes vary considerably from one fly to another, their temporal dynamics are very similar. Below: The mean microsaccade waveforms, sectioned from the black trace above. Their fast activation-phase (up-surge) and slower recovery-phase (down-surge) dynamics are superimposed on a longer adapting trend (downwards slope; red line). Notice how the fast-phases to the light increment, just after 100 ms of darkness (-1 contrast), are greatly amplified (*). Each subfigure (*A* to *C*; upper panels) shows microsaccades (responses) of five different flies (thin gray traces) and their mean (thicker black traces).

The accelerating temporal contrast frequency (Fig. S23 *A* and *B*) evoked progressively smaller photoreceptor microsaccades. In other words, the transient microsaccade phases to positive contrasts (light increments) were the more prominent, the longer the photoreceptors were exposed to negative contrasts (light decrements). These dynamics agree with the theory of refractory stochastic photon sampling by a photoreceptor’s ∼30,000 microvilli (11, 39, 40, 59). Each microvillus is a photon sampling unit capable of transducing a photon’s energy to a unitary response (quantum bump, QB); whilst, QBs from many microvilli integrate a photoreceptor’s macroscopic voltage response (11, 39, 40, 59, 60, 80, 81). Following each QB, the light-activated microvillus becomes refractory for ∼50–300 ms (11, 39, 40). Therefore, during a long positive contrast pulse, a photoreceptor’s sample rate gradually saturates, as fewer microvilli are available to generate QBs and participate in thrusting the microsaccade (10, 11). Whereas, during a long negative contrast pulse, the microvilli recovered from refractoriness so that for the next positive contrast, more microvilli contracted, accentuating the microsaccade’s fast phase (11); see also (10). Correspondingly, the microsaccade responses to the sinusoidal contrasts (Fig. S23*A*) were, on average, less transient than to the pulsatile contrasts (Fig. S23*B*).

The light-adapted photoreceptors’ microsaccades (Fig. S23*C*) to repeated very brief +1 (light-yellow) and - 1 (light-cyan) contrast pulses (100-ms-long) showed these differences in their respective fast- and slow-phase dynamics. Typically, these microsaccades retained ∼0.1-0.4 µm movement range at the rhabdomere level, meaning that a photoreceptor’s receptive field (RF) would repeatedly jump ∼0.3-1.2° in visual space. In other words, in the natural diurnal environment, even a fleeting contrast change could shift a photoreceptor’s RF in the world ≥1/3 of its acceptance angle (Δ*ρ*^*d*^); see Section IV below. Moreover, such microsaccades happen within ∼35 ms for contrast increments, which is 2-to-4-times faster than after prolonged dark-adaptation, ∼70-120 ms (11).

Although the slow-phase (recovery) amplitudes (to -1 contrast) were smaller than the fast-phase (activation) amplitudes (to +1 contrast), both phases were distinguishable, and when corrected for the sloping adapting trend (Fig. S24*A*), somewhat resembled a *Drosophila* R1-R6 photoreceptor’s voltage responses to similar contrast stimuli (82). Characteristically, in both sets of recordings, their fast-phases to the light increment, immediately after 100 ms of darkness (-1 contrast), were greatly accentuated, as predicted by the refractory stochastic photon sampling theory; see also (83). Nevertheless, it was also apparent that the microsaccades traced the voltage response dynamics, giving the impression of mechanically band-passed versions of the photoreceptor voltage output (11). We further quantified this notion by comparing their frequency response functions to dynamic stimulation at comparable light adaptation and temperature (20-22 °C) (Fig. S24*B*).

**Fig. S24.**
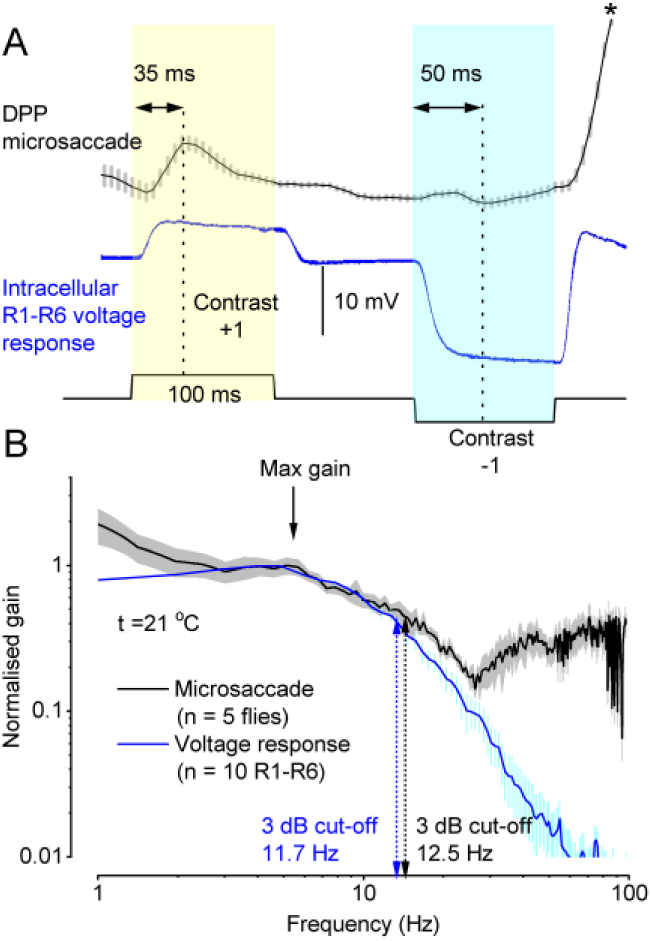
Light-adapted microsaccades to contrast steps show many similarities to the corresponding photoreceptor voltage responses. (*A*) Mean deep-pseudopupil (DPP) microsaccade with its SD (gray; data from Fig. S23C) after the adaptive trend removal and a corresponding intracellular R1-R6 photoreceptor voltage response (blue). Positive contrast steps evoke larger photoreceptor microsaccades than negative contrast steps. In particular, after a -1 contrast (darkness), light increments evoke substantial movements (*). Moreover, microsaccades to +1 contrast (activation) move faster than -1 contrast (recovery), peaking in ∼35 and ∼50 ms, respectively, from the stimulus onset. Photoreceptor voltage responses show comparable dynamics but occur faster. (*B*) The mean microsaccade frequency response to sinusoidal modulation with its SD (gray; data from Fig. S23A) and the mean - intracellularly recorded - photoreceptor voltage response transfer-function (blue) to Gaussian white-noise contrast stimulation. Microsaccades result from photomechanical phototransduction processes (10, 74) and involve mechanical coupling between the neighboring cells (see Sections II.8.i. and II.8.ii., below). Nevertheless, as judged by the corner-frequency, their overall dynamics appear equally fast to an individual photoreceptor’s intracellular voltage responses; see also (10).

To make these comparisons (Fig. S24), we recorded light-adapted *Drosophila* R1–R6 photoreceptors’ intracellular voltage responses *in vivo* (11, 60, 77) with filamented sharp quartz microelectrodes (120–220 MΩ; filled with 3 M KCl) pulled on a Sutter P2000 (USA) electrode puller. The photoreceptors were first light-adapted for 30 s to a bright background at the center of their receptive field. Then, their voltage responses to the luminance contrast pulses and the pseudorandomly modulated luminance changes (∼0.32 mean contrast with 1-500 Hz flat spectrum) were recorded. The data were pre-filtered at 500 Hz, sampled at 1 kHz, and analyzed offline with Biosyst software (Juusola, 1999-2020) as described formerly (39, 60, 77, 84). In brief, we calculated the transfer function *T*(*f*) between the average voltage response, or “signal” *s*(*t*), and the contrast stimuli *c*(*t*) using their 1,024-point-long spectral estimates, *S*(*f*) and *C*(*f*), respectively:

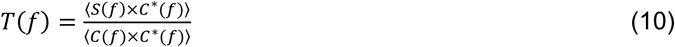

Here 〈 〉 denotes the average over the different stretches and * the complex conjugate. The transfer function’s gain part (blue trace) is shown in Fig. S24*B*. Its 3dB corner frequency was similar to that of the microsaccade frequency response function. This finding is in keeping with the previous voltage response and photomechanical movement comparison (10) and the signal-to-noise analyses of the equivalent voltage and microsaccade responses to 20 Hz bursty light intensity changes (11).

##### II.7 Simulating how pitch, yaw, and roll change optic flow to photoreceptor receptive fields (RFs)

In the natural environment, *Drosophila* perform complex flight maneuvers that involve rotations in three dimensions: *pitch*, head up or down about its wing-to-wing axis; *yaw*, turning left or right about its vertical center axis; and *roll*, rotation about its head-to-abdomen axis. All these axial rotations cause predictable changes in the optic flow the photoreceptors face.

Knowing how ommatidial lens inverts images (see Section V., below) and how local contrast changes evoke mirror-symmetric bidirectional microsaccades (see Section II.6., above), we calculated the pitch-, yaw- and roll-induced optic flow changes within photoreceptors’ receptive fields (RFs) across the left and right *Drosophila* eyes (Fig. S24 to S27). To better appreciate these simulations, one needs to consider that:

- The ommatidial lens system makes the photoreceptor RFs, projected in the visual space, move in the opposite direction to their microsaccades (11) (see Section V., below; Fig. S56*E*). Therefore, for bright objects, if a microsaccade’s fast phase moves back-to-front and the slower phase front-to-back, the photoreceptor’s RF moves first front-to-back and then returns back-to-front. This way, in a forward flight, the RF first seemingly “locks on” the optic flow of things and travels with them before returning to “lock on” the next things passing by. Moreover, when an RF moves with a moving object, the object stays longer within the RF, and its details can be better resolved in time than when the RF moves against the object motion (11). Dark objects will cause a similar effect in the retina due to cooperative local motion, only with a slightly longer lag.
- Microsaccade directions and polarity shift gradually across the eyes, aligned by the R1-R7/8 rhabdomeres’ developmental orientation map (see Section II.1., above; Fig. S11. to S13). For example, the fast microsaccade component shifts from front-to-back at the ventral eye (south hemisphere) to back-to-front at the anterior and dorsal eye (north hemisphere) (Fig. S16*A*). Therefore, attributable to microsaccades’ (i) north-south hemispheric shift in polarity (ii), left-right mirror-symmetricity across the two eyes, and (iii) opposing activation and relaxation phases, the two eyes subdivide into *four optic flow processing quarters*. Equally, how the photoreceptor RFs travel over the visual space shifts in direction and polarity along these quarters but in a reverse way.
- Contrast differences of visual objects further burstify sampling, making photoreceptors ripple between the phases (see Section II.6., above), with light increments driving RFs fast backward and light-decrement slower forwards; as happens in the eyes’ south hemisphere.

***Pitch*** Movie S6 shows the difference between the photoreceptors’ two RF movement phases and optic flow across the right and left eye when a fly rotates a complete circle about its wing-to-wing axis, viz. performs a backward “somersault.” During the “somersault,” its right and left eyes will always experience a centrally expanding flow field, irrespective of whether its proboscis (“nose”) points up, down, left, or right. Therefore, the right and left eye’s mirror-symmetric RF motions (Fig. S25*A*) match the right- and leftward curving optic flow equally well (Fig. S25*B*) at each given head rotation position (Movie S6; Fig. S25 *C* and *D*). Nonetheless, because (i) the fast and slower RF movement directions oppose each other and (ii) their polarities gradually shift along the eyes’ north-south-axis, how the RF phases trace the optic flow will be juxtaposed between the eyes’ north and south hemispheres (Fig. S25 *C* and *D*).

The simulations reveal that the backward-pitching partitions the eyes’ optic-flow-tracing with a north-to-south traveling wavefront. North of the wavefront, the slower RF movement phase matches and the fast phase opposes the optic flow. While south of it, the RF phases reverse (Movie S6). However, when the fly flips upside-down, so do the RF phases, as its eyes now face optic flow from behind. Right through the “somersault,” these dynamics make the eyes’ corresponding north and south differences (Fig. S25 *C* and oscillate with the RF movements’ 180° phase shift (Fig. S25*E*).

**Fig. S25.**
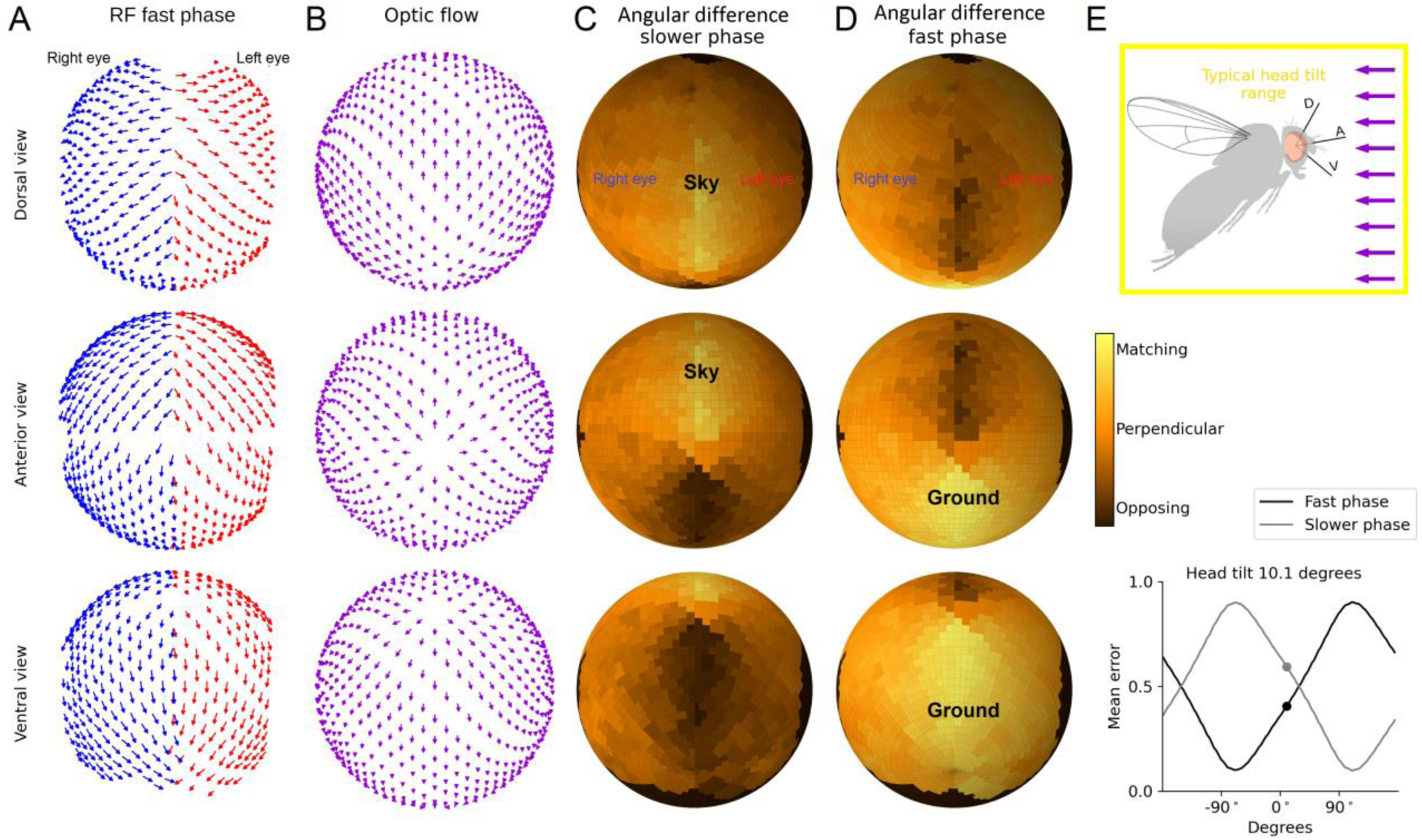
Pitch rotation optic flow juxtaposes the slower and fast photoreceptor receptive field (RF) movement phases along the fly eye’s north and south axis. (*A*) Photoreceptor RF movement fast phase directions across the right (blue) and left (red) eyes; Fig. 3G shows the corresponding slow phase directions. (*B*) Pitch-induced optic flow across the fly eyes is shown for the characteristic forward flight position, in which the two eyes face direct frontal flow. The fly head is upright with a slight 10.1° tilt, shifting the optic flow radiating focus slightly below the equator. (*C*) Difference between the RF slower (relaxation) movement phase and optic flow across the right and left eyes (including their frontal binocular stereo range). The RF slower phase broadly matches the optic flow at the eyes’ north hemisphere. Note the graded nature of the maps. For example, the flow fields are more orthogonal than opposite to the slow-phase microsaccade directions at the side of the eyes (facing the sky). Ultimately, this relationship varies with the flight posture, depending on the head tilt (see Movie S6). (*D*) Difference between the RF fast (activation) movement phase and optic flow across the right and left eyes (including their frontal binocular stereo range). The RF fast phase matches the optic flow at the eyes’ south hemisphere. (*E*) Upper inset: a fly’s characteristic forward flight position with the upright head’s slight tilt. Lower inset: when a fly pitches backward, the directional differences (mean error) between the optic flow and the photoreceptor RFs’ two movement phases, as calculated across the eyes, oscillate with the opposing 180° cycles.

In a fly’s normal forward flight posture (Fig. S25) - with its upright head having a slight 10.1° tilt (*cf*. Fig. S12)- the RFs’ fast- and slower-phases are set in a balanced mid-state, where the fast-phase broadly matches the “ground-flow” and the slower-phase the “sky-flow”. This visual field partitioning into a “slower-phase-matched north hemisphere” and a “fast-phase-matched south hemisphere” may help a fly to see better nearby fast-moving frontal and ventral world objects, such as other *Drosophila* and passing-by food items, and slow-moving more extensive features, such as landscape and clouds, further in the skyline.

***Yaw*** Movie S7 shows how a fly’s right or left turns accentuate phasic differences in binocular contrasts when holding its normal flight posture with the upright head (Fig. S26). Again, the simulations disclose how the optic flow processing differs between the eye quarters (Fig. S26 *A* and *B*), but this time the right and left eye is juxtaposed against each other, rather than the eyes’ north and south halves; as happens in *pitch*.

Explicitly, during a right or left turn, one eye’s RF fast and slower phases move with and against the optic flow (11), respectively, while simultaneously the other eye’s phases do the reverse (Fig. S26 *C* and *D*). Furthermore, since the mean errors between the opposing fast and slower RF movement phases and optic flow are for both eyes (Fig. S26*E*), these values approach 50% while oscillating with the RF movements’ 180° phase shift (*i.e.,* in opposing polarity).

Photoreceptors encode these opposing phases of moving objects in their voltage responses (11), and we later show how their binocular differences – as dynamic disparity signals - could be used by the fly brain to encode visual object depth (see Section IV, below).

**Fig. S26.**
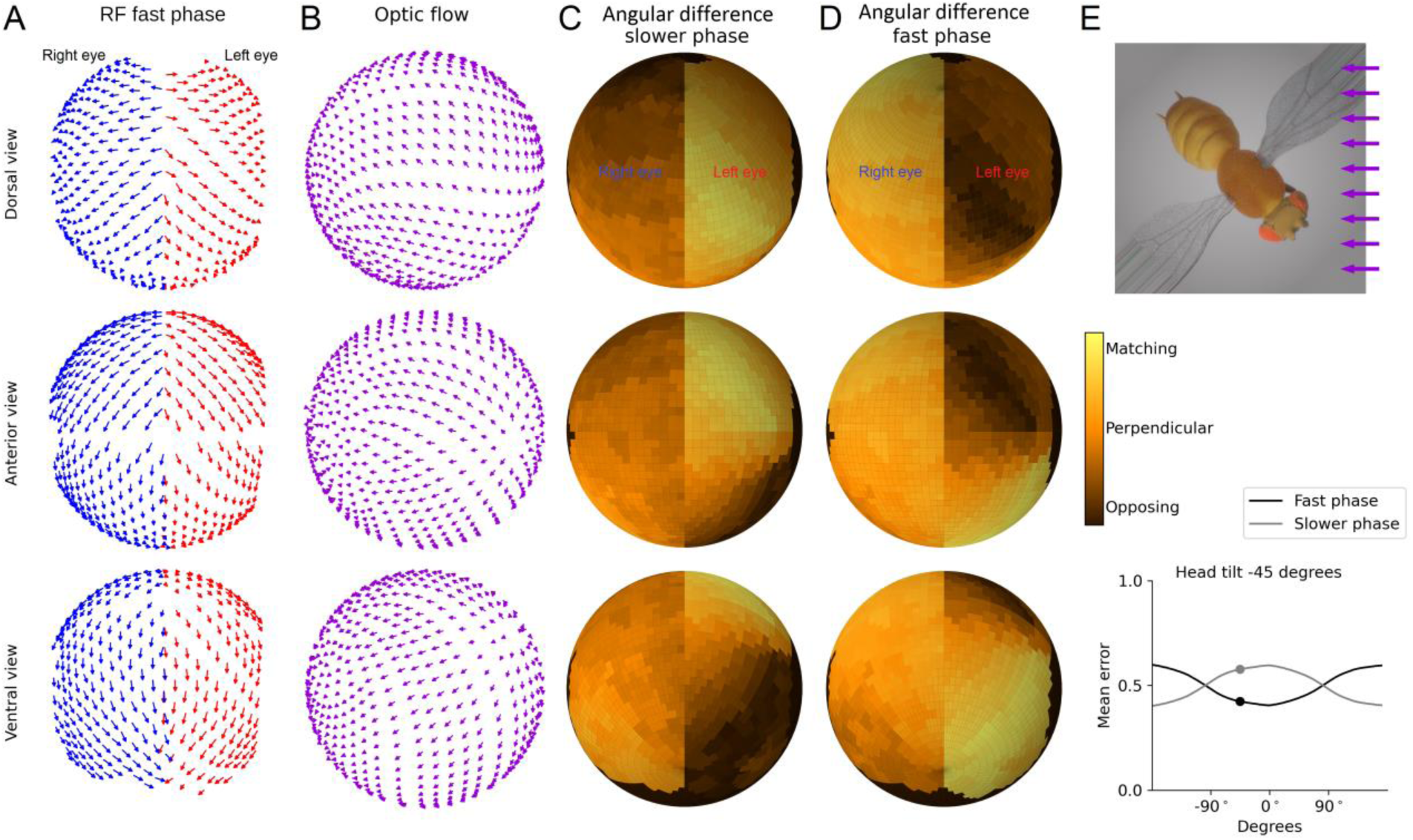
Yaw rotation optic flow juxtaposes the right and left eyes’ slower and fast photoreceptor receptive field (RF) movement phases. (*A*) Photoreceptor RF movement fast phase directions across the right (blue) and left (red) eyes. (*B*) Yaw induced optic flow across the fly eyes for the flight position, in which the eyes face the flow at a -45° angle. The fly head is upright with a slight 10.1° tilt, shifting the optic flow radiating focus slightly below the equator. (*C*) Difference between the RF slower (relaxation) movement phase and optic flow across the right and left eyes. The RF slower phase matches the optic flow at the right eye but opposes at the left eye. (*D*) Difference between the RF fast (activation) movement phase and optic flow across the right and left eyes. The RF fast phase matches the optic flow at the left eye but opposes at the right eye. (*E*) Upper inset: a fly turning against the optic low, snapshot show at -45° angle. Lower inset: when a fly yaw rotates, the directional differences (mean error) between the optic flow and the photoreceptor RFs’ two movement phases, as calculated across the eyes, oscillate with the opposing 180° cycles.

***Roll***. Fig. S27 shows the difference between the two RF phases and optic flow across the right and left eye when a fly rotates about its head-to-abdomen axis. Because a fly always faces frontal optic flow throughout this roll rotation, the RF movements’ fast- and slower-phases remain in a state of static opponency. Consequently, their local differences to optic flow across the eyes (Fig. S27 *C* and *D*) remain similar to that seen in the characteristic forward flight (Fig. S25 *C* and *D*), with the north-hemisphere RF movements’ slower-phase and the south-hemisphere RF movements’ fast-phase matching the optic flow.

**Fig. S27.**
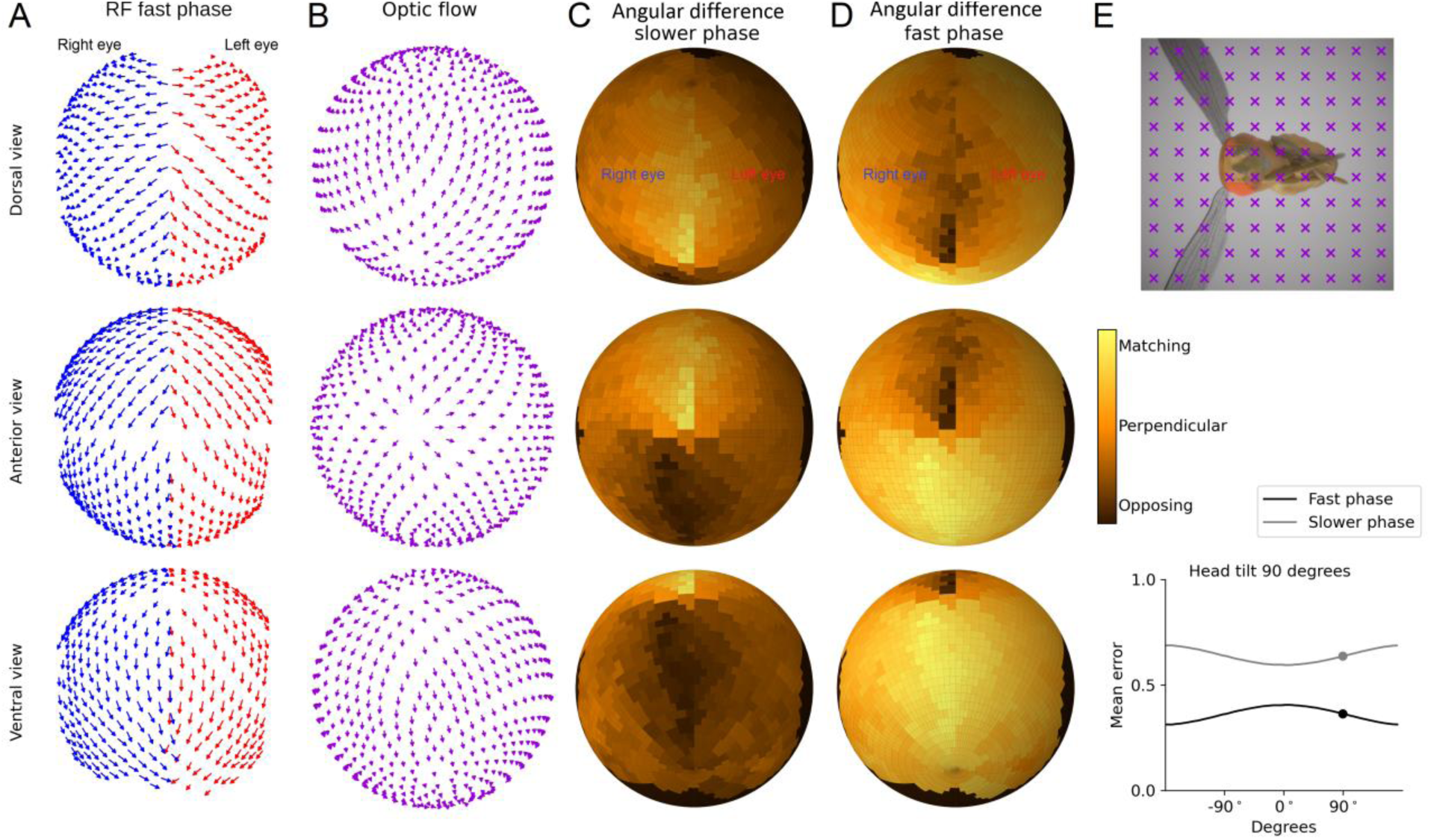
Roll rotation optic flow juxtaposes the north and south eye hemispheres while keeping their slower and fast photoreceptor receptive field (RF) movement phase differences constant. (*A*) Photoreceptor RF movement fast phase directions across the right (blue) and left (red) eyes. (*B*) Roll-induced optic flow across the fly eyes for the characteristic forward flight position, in which the two eyes face direct frontal flow. The fly head is upright with a slight 10.1° tilt, shifting the optic flow radiating focus slightly to the left. (*C*) Difference between the RF slower (relaxation) movement phase and optic flow across the right and left eyes (including their frontal binocular stereo range). The RF slower phase matches the optic flow at the eyes’ north hemisphere. (*D*) Difference between the RF fast (activation) movement phase and optic flow across the right and left eyes (including their frontal binocular stereo range). The RF fast phase matches the optic flow at the eyes’ south hemisphere. (*E*) Upper inset: a fly rolling, snapshot show at 95° angle, when the red Xs indicate frontal optic flow. Lower inset: theoretically, if the fly head faces the optic flow frontally with a fix (0°) angle, the roll will not change the directional difference (mean error) between the optic flow and the photoreceptor RFs’ two movement phases. Thus, the errors would be flat throughout the roll. However, here, the upright head’s slight 10.1° tilt made the error wobble a bit.

##### II.8 Testing mechanical coupling of intra-ommatidial photoreceptors

We examined whether light-activating a single R1-R8 causes it to contract alone or whether this induces ommatidial R1-R8s to move as a unit. Because of the underlying R1-R6 superposition and the left/right eye structural and microsaccadic mirror-symmetricities, both outcomes should sharpen phase differences in moving light input and its binocular R1-R6 outputs to capture stereo- and optic-flow-information better. However, different trade-offs (speed/accuracy) and costs (energy/robustness) might have resulted in selecting one or the other. The results from two separate assays established that *single photoreceptor activation moves all photoreceptors in the same ommatidium*:

i. Using the goniometric system and electrophysiology, we measured DPP microsaccades and ERG responses to UV- and green-light of otherwise blind flies, in which only one photoreceptor type (R7s, R8s, or R1-R6) or all R7/8s were rescued (transgenic Rhodopsin-specific *norpA* rescue flies) (Fig. S28). We also did such recordings in UV-flies, in which R1-R6 the green-sensitive Rh1 was replaced with the UV-sensitive Rh3, and in *ninaE^8^* mutants (22), in which inner photoreceptors (R7/8) functioned normally but outer photoreceptors (R1-R6) were blind (Fig. S29). The expectation was that if photoreceptors move independently, then instead of all 7 R-images moving together, only R7/8, or only R1-R6, would move in flies with R-specific rhodopsin rescue. Instead, we found that all R1-R8 rhabdomeres in optical superposition are dependent. They move together as a unit (Fig. S28 and S29), with the measured microsaccade and ERG responses matching the rescued photoreceptors’ spectral sensitivities (Fig. S30).
ii. Using the cornea neutralization method (Fig. S31) with a targeted single R1-R8 stimulation, we directly measured how R1-R8 rhabdomeres move as a unit inside an ommatidium. We found that activating only the photoreceptors (or just a few of them) in a single ommatidium with a light-spot (Fig. S33) evoked a collective R1-R7/8 microsaccade inside the ommatidium. Meanwhile, the other rhabdomeres outside this ommatidium remained still. Whereas for larger light stimulation areas (light-field stimulation; Fig. S32), the ommatidial rhabdomeres in the stimulus center, experiencing the highest photon rates, moved the most. In contrast, the ommatidial rhabdomeres at the stimulus edge, with the lowest photon rates, moved the least.

###### II.8.i Pseudopupil microsaccades and ERG responses to single photoreceptor class activation

The *norpA* Rh-rescue flies recordings showed that light-activating just a single spectral class of photoreceptors in the optically superpositioned R1-R7/8 rhabdomeres (forming the observed DPP image (24)) is enough to generate a sideways-moving microsaccade (Fig. S28). These data provided strong evidence that R1-R7/8 photoreceptors in each ommatidium do not move independently but are mechanically coupled.

**Fig. S28.**
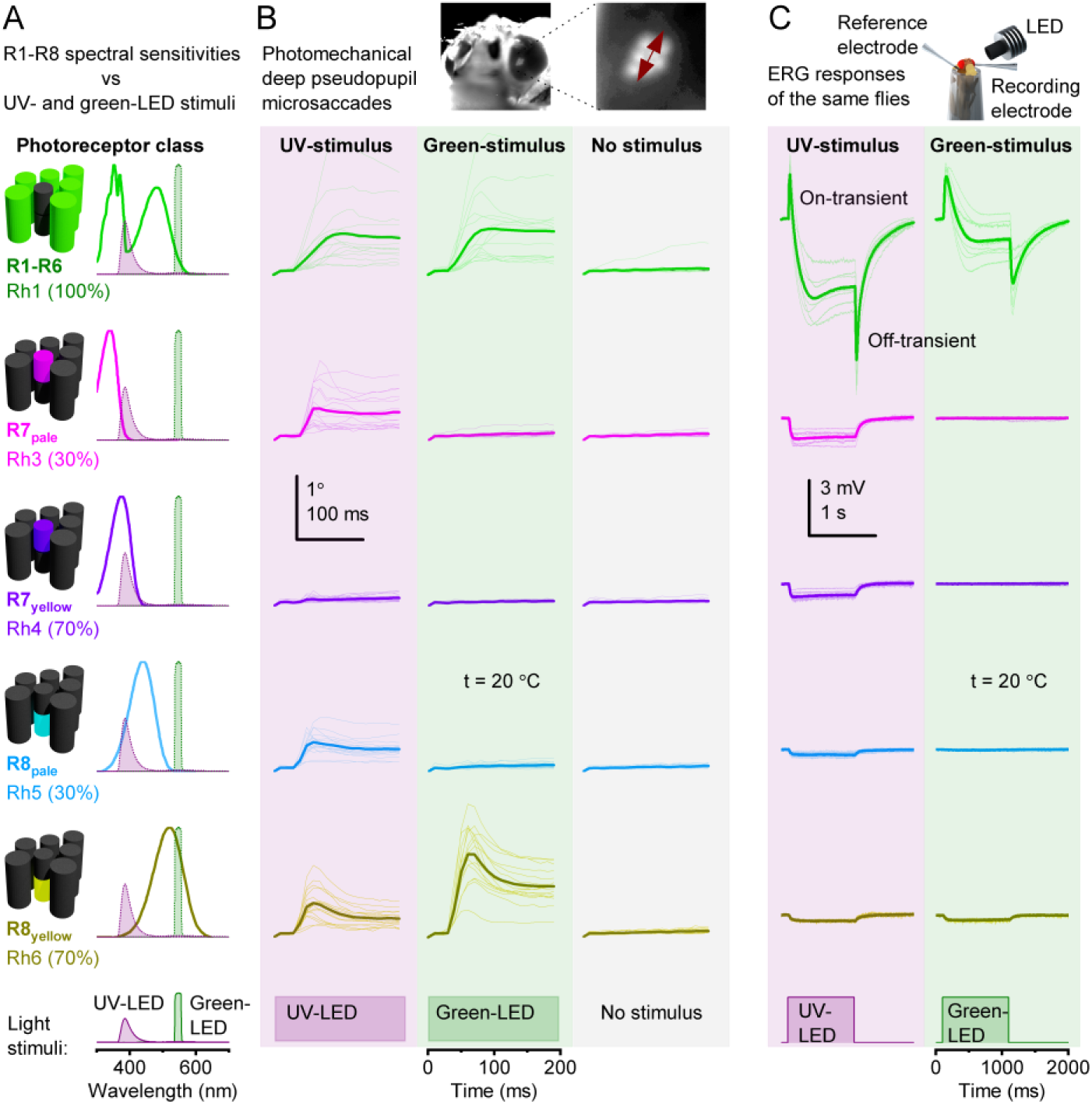
Rhabdomeres inside an ommatidium are mechanically coupled: single spectral-class photoreceptor activations move all R1-R8 rhabdomeres together. (*A*) *norpA* rhodopsin rescue flies have ordinary eye and ommatidial morphologies. However, they have only one functional spectral photoreceptor class: R1-R6, R7_pale_, R7_yellow_, R8_pale,_ or R8_yellow_, with specific prevalence (%) and stochastic ommatidial distribution across the eyes (41). Their respective rhabdomeres and nomograms (co-colored) are shown against the test UV- and green LEDs’ spectral emission (filled curves). The other photoreceptors in the ommatidia (dark gray rhabdomeres) are blind. (*B*) If the UV- or green-stimulus activated the tested photoreceptor spectral class, this caused a fast photomechanical DPP movement, with the observed R1-R7/8 rhabdomeres together bouncing sideways along an eye-location-specific movement axis. The movements were the largest in the Rh1- (R1-R6; green) and Rh6-rescue (R8_yellow_; dark yellow) flies and the smallest (barely distinguishable) in the Rh4-rescue (R7_yellow_; purple). For each spectral class, the movement responses were calculated by cross-correlating the consecutive image frames in 10 ms resolution (11); shown for ten flies (thin traces; from both left and right eye images) and their average (thick traces). Without stimulation, the pseudopupil remained still, showing rarely eye-muscle-induced retinal jitter ((11, 13), *e.g.,* one thin-trace in Rh1 “no stimulus”-subfigure). Therefore, in each ommatidium, R1-R8 rhabdomeres are mechanically coupled/pivoted, possibly by anchoring and ultrastructural links. And one photoreceptor’s light-activation is enough to move its intra-ommatidial neighbors in unison. (*C*) The same flies’ electroretinograms (ERGs) to the UV- and green-LED stimulation showed the predicted spectral sensitivities. ERG verified the rescued photoreceptors’ normal phototransduction/synaptic signaling (on- and off-transient (19, 22, 36)) and the DPP (*B*) movements’ photomechanical origins. The used fiber-optic bundle and its positioning made the LED-light stimulate the anterior-dorsal fraction of the two eyes’ photoreceptors -including the dorsal rim area, where R7 and R8 express the same UV-sensitive Rh3-rhodopsin (85) (pink). Thus, the stimulation location may explain why Rh3-rescue-flies’ UV-ERG was larger than Rh5-rescue-flies’ (assuming equal photon efficiencies of Rh3 and Rh5 - which probably are not identical). The ERG light stimulation was ∼10-fold weaker than the stimuli in pseudopupil experiments, measured by a spectrometer (*B*). All data from pipette-tip-fixed flies.

Using bright spectrally-distanced green- and UV-light stimuli (385 nm UV-LED and 547 nm green-LED peak-wavelengths; Fig. S28A), we further evaluated R1-R6, R7_yellow_, R7_pale_, R8_yellow,_ and R8_pale_ photoreceptors’ relative contributions in powering a microsaccade. For the tested stimuli, R1-R6 and R8_yellow_ activations caused the largest microsaccades and R7_yellow_ activation the smallest. However, because of the mechanical coupling, R1-R8s collective photomechanical sensitivity covers a broad color spectrum. With each rhodopsin having a wide spectral range that overlaps with the other rhodopsins, most monochromatic colors will simultaneously activate multiple photoreceptor spectral classes. Their photomechanics then add up the total microsaccade dynamics. Therefore, for example, an R7_yellow_ photoreceptor will always move along ommatidial R1-R8 microsaccades, irrespective of whether it was directly light-activated or not.

These results further substantiate that intraocular-muscle-activity rarely interferes with *an immobilized Drosophila*’s photomechanical photoreceptor microsaccade dynamics (*cf*. Sections II.4 and II.5., above). Had the microsaccades been or included fast light-triggered muscle-reflexes, their amplitudes to both the UV- and green-stimuli would have been similar, showing spectrally-independent dynamics. Whereas, had the microsaccades been spontaneous or driven by clock-spikes (57), they would have occurred regularly throughout the recordings. Instead, the results showed that individual flies’ microsaccade sensitivity followed their rescued photoreceptors’ spectral sensitivities (*e.g.,* R8_yellow_ in Fig. S28 *A* and *B*) and that the microsaccades never occurred in the “no-stimulus”-control recordings (Fig. S28*B*).

Summing up the rhodopsin rescue *norpA*-mutants (Rh1+Rh3+Rh4+Rh5+Rh6) R1-R6 microsaccades’ *average* lateral movements to the UV-flash gave a total movement of 0.728 µm. However, this movement range is, in fact, less than half of the wild-type flies’ *average* R1-R6 microsaccade movement of 1.538 µm (Fig. S28 to S30). On the other hand, summing up the rhodopsin rescue *norpA*-mutants (Rh1+Rh3+Rh4+Rh5+Rh6) R1-R6 microsaccades’ *maximum* lateral movements (of the best/healthiest preparations) gave a total movement range of 1.966 µm. This value fell comfortably within the wild-type microsaccade movements, ranging from 1.052 to 2.166 µm.

In comparison, the rhodopsin rescue *norpA*-mutants’ (Rh1+Rh3+Rh4+Rh5+Rh6) integrated *average* ERG photoreceptor component to the same UV-flash (6.318 mV) is similar to the wild-type flies’ *average* ERG photoreceptor component (5.034 mV). This finding strongly suggests that the different photoreceptor types’ ERGs sum up the total ERG photoreceptor component.

The discrepancy between the *average* lateral microsaccade amplitudes and *average* ERG responses suggests that the rhodopsin-rescued *norpA*-mutants’ lateral microsaccade component is not always fully rescued and may display sub-optimal structural integrity. This finding is consistent with our observations about the fragility of some mutant flies microsaccades to preparation-induced mechanical stress (Section II.4.) and expression variability (Section VII.6., Fig. S74)

*Other predictable observations* further indicate R1-R7/8 photoreceptors’ photomechanical contractions mechanic coupling to generate their collective microsaccades:

- UV-flies - constructed on *ninaE^8^* mutants (Fig. S29*A*) by rescuing R1-R6 function with UV-sensitive Rh3-rhodopsin expression - have also functional R7/8 photoreceptors (Fig. S29*B*). These R7/8 photoreceptors are sufficient to evoke the UV-flies’ ommatidial R1-R7/8 microsaccades to the green flash, comparable to *ninaE^8^* microsaccades (Fig. S29*B*).
- Rh3-6-*norpA* rescue flies with functioning R7/8 photoreceptors showed similar microsaccade and ERG dynamics to *ninaE^8^*-mutants.

*Other predictable observations* indicate synaptic feedback modulating R1-R7/8 microsaccades:

- *dSK* mutants’ microsaccades (Fig. S29*B*) were faster and smaller than those of the wild-type flies, consistent with their accelerated photoreceptor voltage responses (78, 79). *dSK* mutant R1-R6 photoreceptors have been shown to experience a tonic synaptic feedback overload from the lamina visual interneurons, which continuously depolarize them, making their voltage responses smaller and faster.

**Fig. S29.**
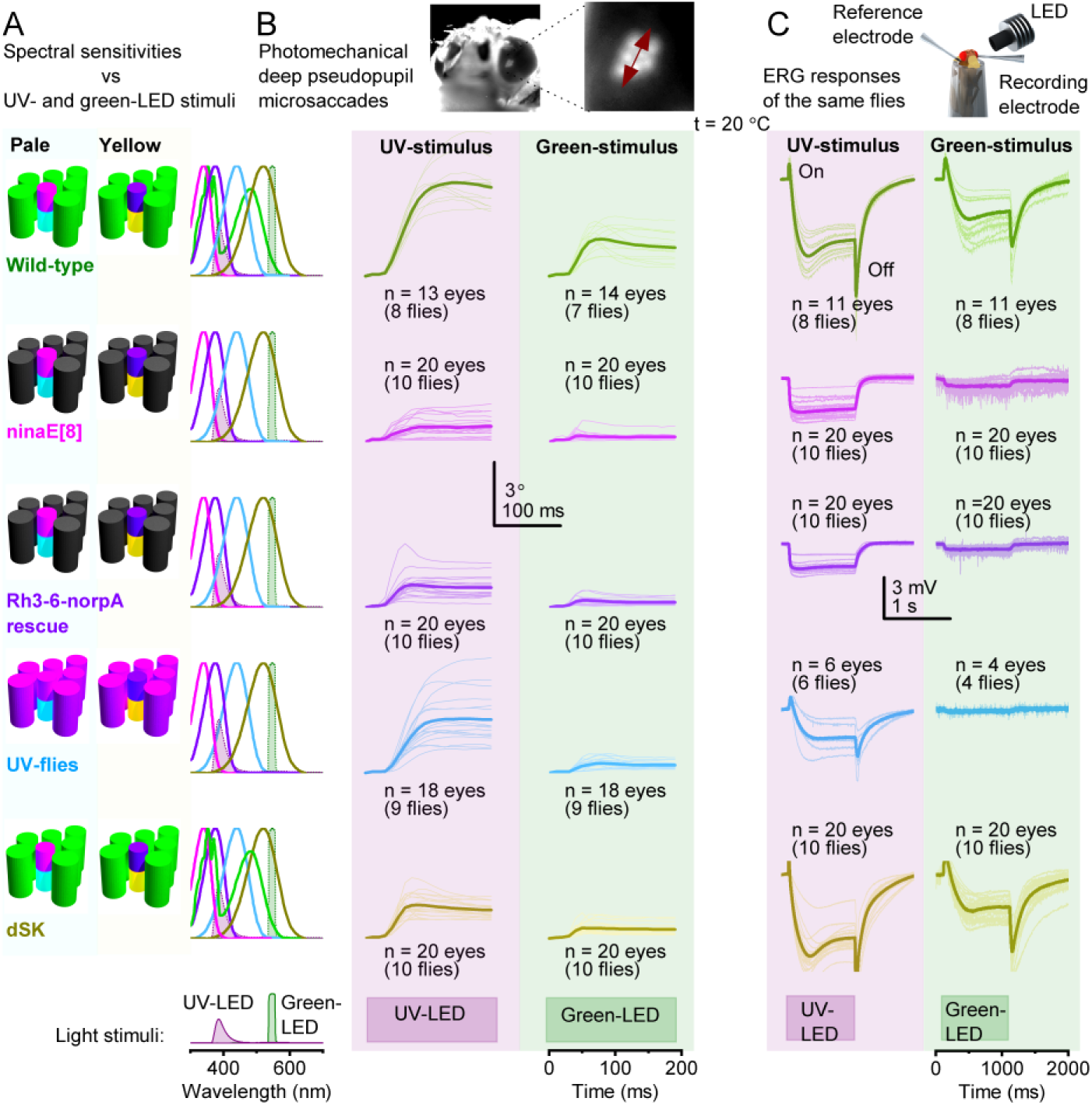
Deep pseudopupil (DPP) microsaccade dynamics combine the contributing photoreceptors’ spectral properties. (*A*) R1-R6 and R7/R8 photoreceptors in wild- type flies and their active and inactive spectral types in *ninaE^8^* mutants, Rh3-R6 *norpA* rescue flies, UV-flies, and *dSK* mutants, respectively. (*B* and *C*) Wild-type DPP microsaccades (middle columns) and ERG dynamics (right columns) integrate inputs from all functional photoreceptor R1-R6, R7_pale_, R7_yellow_, R8_pale,_ and R8_yellow_. *ninaE_8_* and Rh3-Rh6 rescued *norpA* dynamics integrate only R7/R8 inputs. UV-flies, in which R1-R6 photoreceptors express UV-sensitive Rh3 and have normal R7/R7 inputs, generate strong microsaccades to both UV and green stimuli but weak ERG to green light. *norpA* mutants lack both microsaccades and ERG responses. These flies’ respective rhabdomeres and nomograms (co-colored) are shown against the UV and green LEDs’ spectral emission (filled curves). All data are from pipette-tip-affixed flies.

We implemented these coupling dynamics in the detailed optical and biophysical modeling of how photoreceptors sample and integrate spatiotemporal information for dynamic super-resolution stereopsis (see Section V., below).

**Fig. S30.**
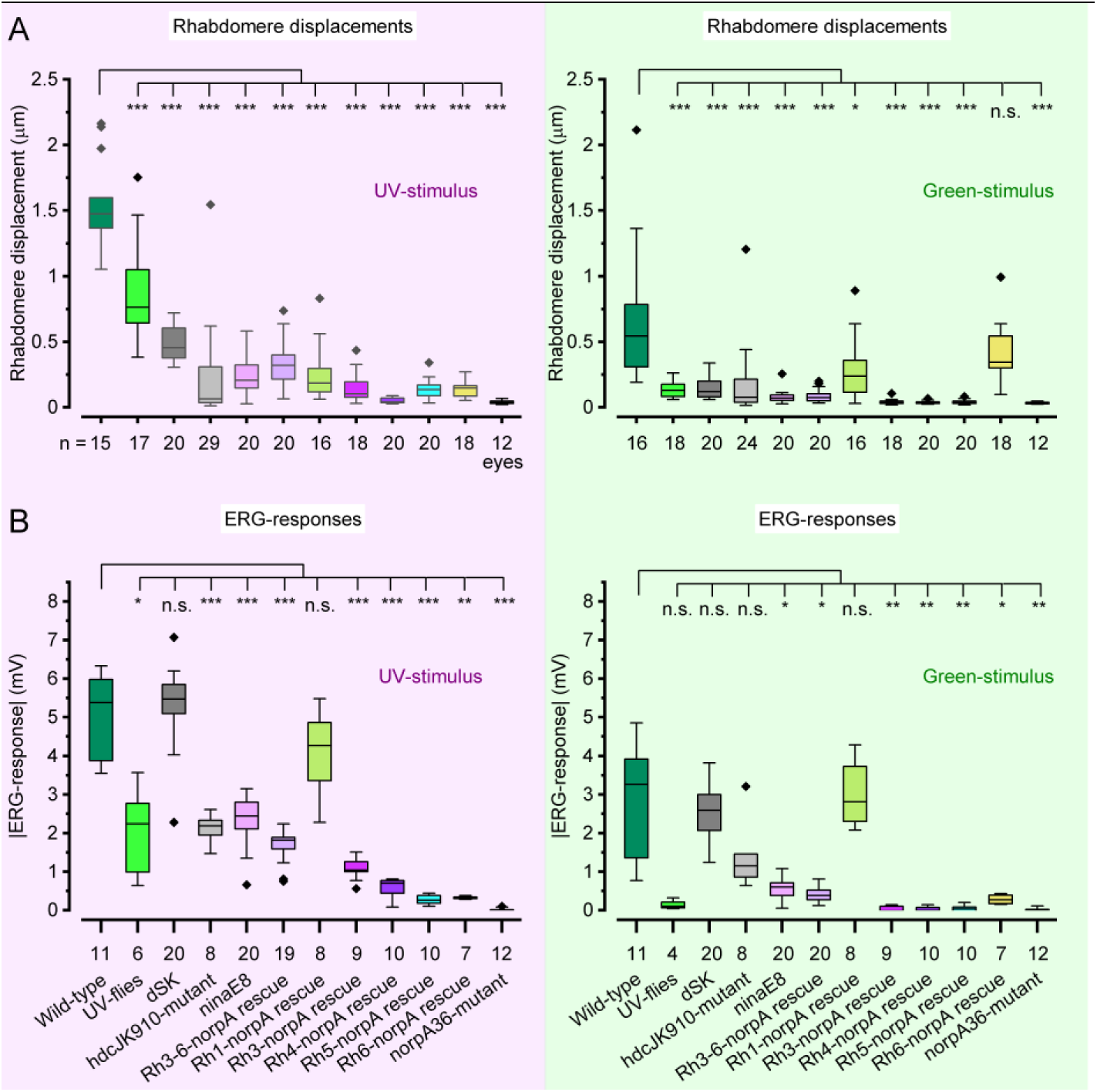
Comparing the Photoreceptor microsaccades (rhabdomere displacements) and ERG-responses between the different tested flies. (*A*) DPP photoreceptor microsaccades to 200 ms bright UV- and green- light pulses. (*B*) ERG-responses of the same flies. The black diamonds indicate each phenotype’s largest and smallest recorded responses. The largest responses were likely from the healthiest preparations; hence, not simply outliers.

We performed a suite of statistical tests to compare the observed DPP photoreceptor microsaccades and the ERG-responses between all tested fly groups. First, we used D’Agostino-Pearson’s normality test (86) to check if a group deviated from a Gaussian distribution with α = 0.05 significance level. If both groups were normally distributed, we used Welch’s adaptation of the two-sided t-test (86) to have higher reliability under unequal variances and sample sizes. If either group significantly deviated from a normal distribution, we used the Mann-Whitney U-test (86) instead. Finally, each statistics table was independently p-value adjusted using the Holm-Šidák step-down method (87) to control the family-wise error rate (Type 1 error) under multiple comparisons.

**Table S2.** DPP microsaccades to 200ms UV flash

| Group A | Group B | N_A | N_B | Mean difference A-B (μm) | Test | p-value (Holm-Sidak) |  |
| --- | --- | --- | --- | --- | --- | --- | --- |
| wild-type | UV-flies | 15 | 17 | $6.673 \times 10^{-1}$ | t-test | $3.584 \times 10^{-4}$ | *** |
| wild-type | *dSK* | 15 | 20 | 1.044 | t-test | $4.166 \times 10^{-8}$ | *** |
| wild-type | *hdc*<sup>JK910</sup> | 15 | 29 | 1.320 | Mann-Whitney | $8.980 \times 10^{-6}$ | *** |
| wild-type | *ninaE*<sup>8</sup> | 15 | 20 | 1.294 | t-test | $6.692 \times 10^{-10}$ | *** |
| wild-type | Rh3-6-*norpA* rescue | 15 | 20 | 1.210 | t-test | $1.661 \times 10^{-9}$ | *** |
| wild-type | Rh1-*norpA* rescue | 15 | 16 | 1.289 | Mann-Whitney | $5.096 \times 10^{-5}$ | *** |
| wild-type | Rh3-*norpA* rescue | 15 | 18 | 1.388 | t-test | $6.814 \times 10^{-10}$ | *** |
| wild-type | Rh4-*norpA* rescue | 15 | 20 | 1.487 | t-test | $2.414 \times 10^{-9}$ | *** |
| wild-type | Rh5-*norpA* rescue | 15 | 20 | 1.399 | Mann–Whitney | $1.531 \times 10^{-5}$ | *** |
| wild-type | Rh6-*norpA* rescue | 15 | 18 | 1.401 | t-test | $2.946 \times 10^{-9}$ | *** |
| wild-type | *norpA*<sup>36</sup>-mutant | 15 | 12 | 1.499 | t-test | $2.234 \times 10^{-9}$ | *** |
| UV-flies | *dSK* | 17 | 20 | $3.767 \times 10^{-1}$ | t-test | $3.015 \times 10^{-2}$ | * |
| UV-flies | *hdc*<sup>JK910</sup> | 17 | 29 | $6.524 \times 10^{-1}$ | Mann–Whitney | $1.852 \times 10^{-5}$ | *** |
| UV-flies | *ninaE*<sup>8</sup> | 17 | 20 | $6.270 \times 10^{-1}$ | t-test | $1.990 \times 10^{-4}$ | *** |
| UV-flies | Rh3-6-*norpA* rescue | 17 | 20 | $5.423 \times 10^{-1}$ | t-test | $9.350 \times 10^{-4}$ | *** |
| UV-flies | Rh1-*norpA* rescue | 17 | 16 | $6.213 \times 10^{-1}$ | Mann–Whitney | $2.350 \times 10^{-4}$ | *** |
| UV-flies | Rh3-*norpA* rescue | 17 | 18 | $7.203 \times 10^{-1}$ | t-test | $3.819 \times 10^{-5}$ | *** |
| UV-flies | Rh4-*norpA* rescue | 17 | 20 | $8.193 \times 10^{-1}$ | t-test | $1.134 \times 10^{-5}$ | *** |
| UV-flies | Rh5-*norpA* rescue | 17 | 20 | $7.319 \times 10^{-1}$ | Mann–Whitney | $6.468 \times 10^{-6}$ | *** |
| UV-flies | Rh6-*norpA* rescue | 17 | 18 | $7.334 \times 10^{-1}$ | t-test | $3.875 \times 10^{-5}$ | *** |
| UV-flies | *norpA*<sup>36</sup>-mutant | 17 | 12 | $8.314 \times 10^{-1}$ | t-test | $9.732 \times 10^{-6}$ | *** |
| *dSK* | *hdc*<sup>JK910</sup> | 20 | 29 | $2.757 \times 10^{-1}$ | Mann–Whitney | $3.106 \times 10^{-4}$ | *** |
| *dSK* | *ninaE*<sup>8</sup> | 20 | 20 | $2.503 \times 10^{-1}$ | t-test | $1.395 \times 10^{-4}$ | *** |
| *dSK* | Rh3-6-*norpA* rescue | 20 | 20 | $1.656 \times 10^{-1}$ | t-test | $2.820 \times 10^{-2}$ | * |
| *dSK* | Rh1-*norpA* rescue | 20 | 16 | $2.446 \times 10^{-1}$ | Mann–Whitney | $1.917 \times 10^{-3}$ | ** |
| *dSK* | Rh3-*norpA* rescue | 20 | 18 | $3.436 \times 10^{-1}$ | t-test | $2.037 \times 10^{-8}$ | *** |
| *dSK* | Rh4-*norpA* rescue | 20 | 20 | $4.426 \times 10^{-1}$ | t-test | $2.157 \times 10^{-10}$ | *** |
| *dSK* | Rh5-*norpA* rescue | 20 | 20 | $3.552 \times 10^{-1}$ | Mann–Whitney | $2.928 \times 10^{-6}$ | *** |
| *dSK* | Rh6-*norpA* rescue | 20 | 18 | $3.567 \times 10^{-1}$ | t-test | $1.366 \times 10^{-9}$ | *** |
| *dSK* | *norpA*<sup>36</sup>-mutant | 20 | 12 | $4.547 \times 10^{-1}$ | t-test | $1.490 \times 10^{-10}$ | *** |
| *hdc*<sup>JK910</sup> | *ninaE*<sup>8</sup> | 29 | 20 | $-2.539 \times 10^{-2}$ | Mann–Whitney | $4.148 \times 10^{-1}$ | ns |
| *hdc*<sup>JK910</sup> | Rh3-6-*norpA* rescue | 29 | 20 | $-1.101 \times 10^{-1}$ | Mann–Whitney | $3.792 \times 10^{-2}$ | * |
| *hdc*<sup>JK910</sup> | Rh1-*norpA* rescue | 29 | 16 | $-3.108 \times 10^{-2}$ | Mann–Whitney | $4.148 \times 10^{-1}$ | ns |
| *hdc*<sup>JK910</sup> | Rh3-*norpA* rescue | 29 | 18 | $6.785 \times 10^{-2}$ | Mann–Whitney | $7.898 \times 10^{-1}$ | ns |
| *hdc*<sup>JK910</sup> | Rh4-*norpA* rescue | 29 | 20 | $1.669 \times 10^{-1}$ | Mann–Whitney | $4.817 \times 10^{-1}$ | ns |
| *hdc*<sup>JK910</sup> | Rh5-*norpA* rescue | 29 | 20 | $7.948 \times 10^{-2}$ | Mann–Whitney | $7.898 \times 10^{-1}$ | ns |
| *hdc*<sup>JK910</sup> | Rh6-*norpA* rescue | 29 | 18 | $8.098 \times 10^{-2}$ | Mann–Whitney | $7.898 \times 10^{-1}$ | ns |
| *hdc*<sup>JK910</sup> | *norpA*<sup>36</sup>-mutant | 29 | 12 | $1.790 \times 10^{-1}$ | Mann–Whitney | $2.126 \times 10^{-1}$ | ns |
| *ninaE*<sup>8</sup> | Rh3-6-*norpA* rescue | 20 | 20 | $-8.473 \times 10^{-2}$ | t-test | $5.800 \times 10^{-1}$ | ns |
| *ninaE*<sup>8</sup> | Rh1-*norpA* rescue | 20 | 16 | $-5.689 \times 10^{-3}$ | Mann–Whitney | $8.295 \times 10^{-1}$ | ns |
| *ninaE*<sup>8</sup> | Rh3-*norpA* rescue | 20 | 18 | $9.324 \times 10^{-2}$ | t-test | $4.148 \times 10^{-1}$ | ns |
| *ninaE*<sup>8</sup> | Rh4-*norpA* rescue | 20 | 20 | $1.923 \times 10^{-1}$ | t-test | $7.883 \times 10^{-4}$ | *** |
| *ninaE*<sup>8</sup> | Rh5-*norpA* rescue | 20 | 20 | 1.049 x 10<sup>-1</sup> | Mann–Whitney | 1.251 x 10<sup>-1</sup> | ns |
| *ninaE*<sup>8</sup> | Rh6-*norpA* rescue | 20 | 18 | 1.064 x 10<sup>-1</sup> | t-test | 1.613 x 10<sup>-1</sup> | ns |
| *ninaE*<sup>8</sup> | *norpA*<sup>36</sup>-mutant | 20 | 12 | 2.044 x 10<sup>-1</sup> | t-test | 3.958 x 10<sup>-4</sup> | *** |
| Rh3-6-*norpA* rescue | Rh1-*norpA* rescue | 20 | 16 | 7.904 x 10<sup>-2</sup> | Mann–Whitney | 3.579 x 10<sup>-1</sup> | ns |
| Rh3-6-*norpA* rescue | Rh3-*norpA* rescue | 20 | 18 | 1.780 x 10<sup>-1</sup> | t-test | 1.128 x 10<sup>-2</sup> | * |
| Rh3-6-*norpA* rescue | Rh4-*norpA* rescue | 20 | 20 | 2.771 x 10<sup>-1</sup> | t-test | 1.701 x 10<sup>-5</sup> | *** |
| Rh3-6-*norpA* rescue | Rh5-*norpA* rescue | 20 | 20 | 1.896 x 10<sup>-1</sup> | Mann–Whitney | 3.163 x 10<sup>-4</sup> | *** |
| Rh3-6-*norpA* rescue | Rh6-*norpA* rescue | 20 | 18 | 1.911 x 10<sup>-1</sup> | t-test | 1.547 x 10<sup>-3</sup> | ** |
| Rh3-6-*norpA* rescue | *norpA*<sup>36</sup>-mutant | 20 | 12 | 2.892 x 10<sup>-1</sup> | t-test | 9.735 x 10<sup>-6</sup> | *** |
| Rh1-*norpA* rescue | Rh3-*norpA* rescue | 16 | 18 | 9.893 x 10<sup>-2</sup> | Mann–Whitney | 4.817 x 10<sup>-1</sup> | ns |
| Rh1-*norpA* rescue | Rh4-*norpA* rescue | 16 | 20 | 1.980 x 10<sup>-1</sup> | Mann–Whitney | 1.395 x 10<sup>-4</sup> | *** |
| Rh1-*norpA* rescue | Rh5-*norpA* rescue | 16 | 20 | 1.106 x 10<sup>-1</sup> | Mann–Whitney | 3.511 x 10<sup>-1</sup> | ns |
| Rh1-*norpA* rescue | Rh6-*norpA* rescue | 16 | 18 | 1.121 x 10<sup>-1</sup> | Mann–Whitney | 3.511 x 10<sup>-1</sup> | ns |
| Rh1-*norpA* rescue | *norpA*<sup>36</sup>-mutant | 16 | 12 | 2.101 x 10<sup>-1</sup> | Mann–Whitney | 2.699 x 10<sup>-4</sup> | *** |
| Rh3-*norpA* rescue | Rh4-*norpA* rescue | 18 | 20 | 9.909 x 10<sup>-2</sup> | t-test | 4.436 x 10<sup>-2</sup> | * |
| Rh3-*norpA* rescue | Rh5-*norpA* rescue | 18 | 20 | 1.162 x 10<sup>-2</sup> | Mann–Whitney | 8.295 x 10<sup>-1</sup> | ns |
| Rh3-*norpA* rescue | Rh6-*norpA* rescue | 18 | 18 | 1.313 x 10<sup>-2</sup> | t-test | 8.295 x 10<sup>-1</sup> | ns |
| Rh3-*norpA* rescue | *norpA*<sup>36</sup>-mutant | 18 | 12 | 1.112 x 10<sup>-1</sup> | t-test | 2.012 x 10<sup>-2</sup> | * |
| Rh4-*norpA* rescue | Rh5-*norpA* rescue | 20 | 20 | -8.746 x 10<sup>-2</sup> | Mann–Whitney | 3.469 x 10<sup>-4</sup> | *** |
| Rh4-*norpA* rescue | Rh6-*norpA* rescue | 20 | 18 | -8.596 x 10<sup>-2</sup> | t-test | 3.106 x 10<sup>-4</sup> | *** |
| Rh4-*norpA* rescue | *norpA*<sup>36</sup>-mutant | 20 | 12 | 1.210 x 10<sup>-2</sup> | t-test | 4.817 x 10<sup>-1</sup> | ns |
| Rh5-*norpA* rescue | Rh6-*norpA* rescue | 20 | 18 | 1.505 x 10<sup>-3</sup> | Mann–Whitney | 8.295 x 10<sup>-1</sup> | ns |
| Rh5-*norpA* rescue | *norpA*<sup>36</sup>-mutant | 20 | 12 | 9.956 x 10<sup>-2</sup> | Mann–Whitney | 6.222 x 10<sup>-4</sup> | *** |
| Rh6-*norpA* rescue | *norpA*<sup>36</sup>-mutant | 18 | 12 | 9.805 x 10<sup>-2</sup> | t-test | 6.262 x 10<sup>-5</sup> | *** |

**Table S3.** DPP microsaccades to 200ms Green flash

| Group A | Group B | N_A | N_B | Mean difference A-B (μm) | Test | p-value (Holm-Sidak) |  |
| --- | --- | --- | --- | --- | --- | --- | --- |
| wild-type | UV-flies | 16 | 18 | 5.415 x 10<sup>-1</sup> | Mann–Whitney | 3.122 x 10<sup>-5</sup> | *** |
| wild-type | *dSK* | 16 | 20 | 5.209 x 10<sup>-1</sup> | Mann–Whitney | 9.494 x 10<sup>-5</sup> | *** |
| wild-type | *hdc<sup>JK910</sup>* | 16 | 24 | $4.941 \times 10^{-1}$ | Mann–Whitney | $2.358 \times 10^{-4}$ | *** |
| wild-type | *ninaE<sup>8</sup>* | 16 | 20 | $5.928 \times 10^{-1}$ | Mann–Whitney | $1.286 \times 10^{-5}$ | *** |
| wild-type | Rh3-6-*norpA* rescue | 16 | 20 | $5.882 \times 10^{-1}$ | Mann–Whitney | $1.286 \times 10^{-5}$ | *** |
| wild-type | Rh1-*norpA* rescue | 16 | 16 | $3.935 \times 10^{-1}$ | Mann–Whitney | $3.543 \times 10^{-2}$ | * |
| wild-type | Rh3-*norpA* rescue | 16 | 18 | $6.311 \times 10^{-1}$ | Mann–Whitney | $1.917 \times 10^{-5}$ | *** |
| wild-type | Rh4-*norpA* rescue | 16 | 20 | $6.381 \times 10^{-1}$ | Mann–Whitney | $1.144 \times 10^{-5}$ | *** |
| wild-type | Rh5-*norpA* rescue | 16 | 20 | $6.325 \times 10^{-1}$ | Mann–Whitney | $1.144 \times 10^{-5}$ | *** |
| wild-type | Rh6-*norpA* rescue | 16 | 18 | $2.634 \times 10^{-1}$ | Mann–Whitney | $5.074 \times 10^{-1}$ | ns |
| wild-type | *norpA<sup>36</sup>*-mutant | 16 | 12 | $6.420 \times 10^{-1}$ | Mann–Whitney | $1.947 \times 10^{-4}$ | *** |
| UV-flies | *dSK* | 18 | 20 | $-2.065 \times 10^{-2}$ | t-test | $8.294 \times 10^{-1}$ | ns |
| UV-flies | *hdc<sup>JK910</sup>* | 18 | 24 | $-4.747 \times 10^{-2}$ | Mann–Whitney | $7.324 \times 10^{-1}$ | ns |
| UV-flies | *ninaE<sup>8</sup>* | 18 | 20 | $5.128 \times 10^{-2}$ | Mann–Whitney | $3.543 \times 10^{-2}$ | * |
| UV-flies | Rh3-6-*norpA* rescue | 18 | 20 | $4.669 \times 10^{-2}$ | t-test | $2.149 \times 10^{-1}$ | ns |
| UV-flies | Rh1-*norpA* rescue | 18 | 16 | $-1.481 \times 10^{-1}$ | Mann–Whitney | $2.518 \times 10^{-1}$ | ns |
| UV-flies | Rh3-*norpA* rescue | 18 | 18 | $8.956 \times 10^{-2}$ | Mann–Whitney | $3.046 \times 10^{-5}$ | *** |
| UV-flies | Rh4-*norpA* rescue | 18 | 20 | $9.655 \times 10^{-2}$ | Mann–Whitney | $7.654 \times 10^{-6}$ | *** |
| UV-flies | Rh5-*norpA* rescue | 18 | 20 | $9.093 \times 10^{-2}$ | Mann–Whitney | $1.286 \times 10^{-5}$ | *** |
| UV-flies | Rh6-*norpA* rescue | 18 | 18 | $-2.782 \times 10^{-1}$ | Mann–Whitney | $6.426 \times 10^{-5}$ | *** |
| UV-flies | *norpA<sup>36</sup>*-mutant | 18 | 12 | $1.004 \times 10^{-1}$ | t-test | $8.054 \times 10^{-5}$ | *** |
| *dSK* | *hdc<sup>JK910</sup>* | 20 | 24 | $-2.682 \times 10^{-2}$ | Mann–Whitney | $6.424 \times 10^{-1}$ | ns |
| *dSK* | *ninaE<sup>8</sup>* | 20 | 20 | $7.193 \times 10^{-2}$ | Mann–Whitney | $2.092 \times 10^{-2}$ | * |
| *dSK* | Rh3-6-*norpA* rescue | 20 | 20 | $6.734 \times 10^{-2}$ | t-test | $1.033 \times 10^{-1}$ | ns |
| *dSK* | Rh1-*norpA* rescue | 20 | 16 | $-1.274 \times 10^{-1}$ | Mann–Whitney | $3.071 \times 10^{-1}$ | ns |
| *dSK* | Rh3-*norpA* rescue | 20 | 18 | $1.102 \times 10^{-1}$ | Mann–Whitney | $1.645 \times 10^{-5}$ | *** |
| *dSK* | Rh4-*norpA* rescue | 20 | 20 | $1.172 \times 10^{-1}$ | Mann–Whitney | $2.606 \times 10^{-6}$ | *** |
| *dSK* | Rh5-*norpA* rescue | 20 | 20 | $1.116 \times 10^{-1}$ | Mann–Whitney | $6.041 \times 10^{-6}$ | *** |
| *dSK* | Rh6-*norpA* rescue | 20 | 18 | $-2.575 \times 10^{-1}$ | Mann–Whitney | $1.947 \times 10^{-4}$ | *** |
| *dSK* | *norpA<sup>36</sup>*-mutant | 20 | 12 | $1.211 \times 10^{-1}$ | t-test | $2.376 \times 10^{-4}$ | *** |
| *hdc<sup>JK910</sup>* | *ninaE<sup>8</sup>* | 24 | 20 | $9.875 \times 10^{-2}$ | Mann–Whitney | $8.294 \times 10^{-1}$ | ns |
| *hdc<sup>JK910</sup>* | Rh3-6-*norpA* rescue | 24 | 20 | $9.416 \times 10^{-2}$ | Mann–Whitney | $8.294 \times 10^{-1}$ | ns |
| *hdc<sup>JK910</sup>* | Rh1-*norpA* rescue | 24 | 16 | $-1.006 \times 10^{-1}$ | Mann–Whitney | $3.071 \times 10^{-1}$ | ns |
| *hdc<sup>JK910</sup>* | Rh3-*norpA* rescue | 24 | 18 | $1.370 \times 10^{-1}$ | Mann–Whitney | $1.264 \times 10^{-1}$ | ns |
| *hdc<sup>JK910</sup>* | Rh4-*norpA* rescue | 24 | 20 | $1.440 \times 10^{-1}$ | Mann–Whitney | $3.523 \times 10^{-2}$ | * |
| *hdc<sup>JK910</sup>* | Rh5-*norpA* rescue | 24 | 20 | $1.384 \times 10^{-1}$ | Mann–Whitney | $9.852 \times 10^{-2}$ | ns |
| *hdc<sup>JK910</sup>* | Rh6-*norpA* rescue | 24 | 18 | $-2.307 \times 10^{-1}$ | Mann–Whitney | $1.148 \times 10^{-3}$ | ** |
| *hdc<sup>JK910</sup>* | *norpA*<sup>36</sup>-mutant | 24 | 12 | $1.479 \times 10^{-1}$ | Mann–Whitney | $3.591 \times 10^{-2}$ | * |
| *ninaE<sup>8</sup>* | Rh3-6-*norpA* rescue | 20 | 20 | $-4.586 \times 10^{-3}$ | Mann–Whitney | $8.294 \times 10^{-1}$ | ns |
| *ninaE<sup>8</sup>* | Rh1-*norpA* rescue | 20 | 16 | $-1.994 \times 10^{-1}$ | Mann–Whitney | $1.349 \times 10^{-3}$ | ** |
| *ninaE<sup>8</sup>* | Rh3-*norpA* rescue | 20 | 18 | $3.828 \times 10^{-2}$ | Mann–Whitney | $2.640 \times 10^{-3}$ | ** |
| *ninaE<sup>8</sup>* | Rh4-*norpA* rescue | 20 | 20 | $4.527 \times 10^{-2}$ | Mann–Whitney | $8.054 \times 10^{-5}$ | *** |
| *ninaE<sup>8</sup>* | Rh5-*norpA* rescue | 20 | 20 | $3.965 \times 10^{-2}$ | Mann–Whitney | $1.002 \times 10^{-3}$ | ** |
| *ninaE<sup>8</sup>* | Rh6-*norpA* rescue | 20 | 18 | $-3.295 \times 10^{-1}$ | Mann–Whitney | $1.027 \times 10^{-5}$ | *** |
| *ninaE<sup>8</sup>* | *norpA*<sup>36</sup>-mutant | 20 | 12 | $4.913 \times 10^{-2}$ | Mann–Whitney | $4.332 \times 10^{-4}$ | *** |
| Rh3-6-*norpA* rescue | Rh1-*norpA* rescue | 20 | 16 | $-1.948 \times 10^{-1}$ | Mann–Whitney | $5.386 \times 10^{-3}$ | ** |
| Rh3-6-*norpA* rescue | Rh3-*norpA* rescue | 20 | 18 | $4.287 \times 10^{-2}$ | Mann–Whitney | $1.095 \times 10^{-2}$ | * |
| Rh3-6-*norpA* rescue | Rh4-*norpA* rescue | 20 | 20 | $4.986 \times 10^{-2}$ | Mann–Whitney | $2.376 \times 10^{-4}$ | *** |
| Rh3-6-*norpA* rescue | Rh5-*norpA* rescue | 20 | 20 | $4.424 \times 10^{-2}$ | Mann–Whitney | $3.973 \times 10^{-3}$ | ** |
| Rh3-6-*norpA* rescue | Rh6-*norpA* rescue | 20 | 18 | $-3.249 \times 10^{-1}$ | Mann–Whitney | $1.440 \times 10^{-5}$ | *** |
| Rh3-6-*norpA* rescue | *norpA*<sup>36</sup>-mutant | 20 | 12 | $5.371 \times 10^{-2}$ | t-test | $2.640 \times 10^{-3}$ | ** |
| Rh1-*norpA* rescue | Rh3-*norpA* rescue | 16 | 18 | $2.377 \times 10^{-1}$ | Mann–Whitney | $2.173 \times 10^{-4}$ | *** |
| Rh1-*norpA* rescue | Rh4-*norpA* rescue | 16 | 20 | $2.446 \times 10^{-1}$ | Mann–Whitney | $7.445 \times 10^{-5}$ | *** |
| Rh1-*norpA* rescue | Rh5-*norpA* rescue | 16 | 20 | $2.390 \times 10^{-1}$ | Mann–Whitney | $1.081 \times 10^{-4}$ | *** |
| Rh1-*norpA* rescue | Rh6-*norpA* rescue | 16 | 18 | $-1.301 \times 10^{-1}$ | Mann–Whitney | $3.071 \times 10^{-1}$ | ns |
| Rh1-*norpA* rescue | *norpA*<sup>36</sup>-mutant | 16 | 12 | $2.485 \times 10^{-1}$ | Mann–Whitney | $5.693 \times 10^{-4}$ | *** |
| Rh3-*norpA* rescue | Rh4-*norpA* rescue | 18 | 20 | $6.993 \times 10^{-3}$ | Mann–Whitney | $5.566 \times 10^{-1}$ | ns |
| Rh3-*norpA* rescue | Rh5-*norpA* rescue | 18 | 20 | $1.372 \times 10^{-3}$ | Mann–Whitney | $8.294 \times 10^{-1}$ | ns |
| Rh3-*norpA* rescue | Rh6-*norpA* rescue | 18 | 18 | -3.677 x 10<sup>-1</sup> | Mann–Whitney | 1.144 x 10<sup>-5</sup> | *** |
| Rh3-*norpA* rescue | *norpA*<sup>36</sup>-mutant | 18 | 12 | 1.085 x 10<sup>-2</sup> | Mann–Whitney | 3.071 x 10<sup>-1</sup> | ns |
| Rh4-*norpA* rescue | Rh5-*norpA* rescue | 20 | 20 | -5.622 x 10<sup>-3</sup> | Mann–Whitney | 6.424 x 10<sup>-1</sup> | ns |
| Rh4-*norpA* rescue | Rh6-*norpA* rescue | 20 | 18 | -3.747 x 10<sup>-1</sup> | Mann–Whitney | 5.004 x 10<sup>-6</sup> | *** |
| Rh4-*norpA* rescue | *norpA*<sup>36</sup>-mutant | 20 | 12 | 3.853 x 10<sup>-3</sup> | Mann–Whitney | 7.289 x 10<sup>-1</sup> | ns |
| Rh5-*norpA* rescue | Rh6-*norpA* rescue | 20 | 18 | -3.691 x 10<sup>-1</sup> | Mann–Whitney | 5.004 x 10<sup>-6</sup> | *** |
| Rh5-*norpA* rescue | *norpA*<sup>36</sup>-mutant | 20 | 12 | 9.475 x 10<sup>-3</sup> | Mann–Whitney | 3.071 x 10<sup>-1</sup> | ns |
| Rh6-*norpA* rescue | *norpA*<sup>36</sup>-mutant | 18 | 12 | 3.786 x 10<sup>-1</sup> | Mann–Whitney | 1.149 x 10<sup>-4</sup> | *** |

**Table S4.** DPP microsaccades to 200ms UV flash

| Group A | Group B | N_A | N_B | Mean difference A-B (mV) | Test | p-value (Holm-Sidak) |  |
| --- | --- | --- | --- | --- | --- | --- | --- |
| wild-type | UV-flies | 11 | 6 | -2.962 | Mann–Whitney | 1.450 x 10<sup>-2</sup> | * |
| wild-type | *dSK* | 11 | 20 | 3.121 x 10<sup>-1</sup> | Mann–Whitney | 5.316 x 10<sup>-1</sup> | ns |
| wild-type | *hdc*<sup>JK910</sup> | 11 | 8 | -2.909 | t-test | 4.876 x 10<sup>-5</sup> | *** |
| wild-type | *ninaE*<sup>8</sup> | 11 | 20 | -2.686 | Mann–Whitney | 1.601 x 10<sup>-4</sup> | *** |
| wild-type | Rh3-6-*norpA* rescue | 11 | 20 | -3.336 | Mann–Whitney | 1.601 x 10<sup>-4</sup> | *** |
| wild-type | Rh1-*norpA* rescue | 11 | 8 | -9.439 x 10<sup>-1</sup> | t-test | 4.097 x 10<sup>-1</sup> | ns |
| wild-type | Rh3-*norpA* rescue | 11 | 9 | -3.975 | t-test | 2.734 x 10<sup>-6</sup> | *** |
| wild-type | Rh4-*norpA* rescue | 11 | 10 | -4.460 | t-test | 1.077 x 10<sup>-6</sup> | *** |
| wild-type | Rh5-*norpA* rescue | 11 | 10 | -4.759 | t-test | 1.347 x 10<sup>-6</sup> | *** |
| wild-type | Rh6-*norpA* rescue | 11 | 7 | -4.711 | Mann–Whitney | 7.492 x 10<sup>-3</sup> | ** |
| wild-type | *norpA*<sup>36</sup>-mutant | 11 | 12 | -5.037 | t-test | 9.915 x 10<sup>-7</sup> | *** |
| UV-flies | *dSK* | 6 | 20 | 3.274 | Mann–Whitney | 6.316 x 10<sup>-3</sup> | ** |
| UV-flies | *hdc*<sup>JK910</sup> | 6 | 8 | 5.328 x 10<sup>-2</sup> | Mann–Whitney | 5.316 x 10<sup>-1</sup> | ns |
| UV-flies | *ninaE*<sup>8</sup> | 6 | 20 | 2.764 x 10<sup>-1</sup> | Mann–Whitney | 5.316 x 10<sup>-1</sup> | ns |
| UV-flies | Rh3-6-*norpA* rescue | 6 | 20 | -3.735 x 10<sup>-1</sup> | Mann–Whitney | 4.652 x 10<sup>-1</sup> | ns |
| UV-flies | Rh1-*norpA* rescue | 6 | 8 | 2.018 | Mann–Whitney | 6.217 x 10<sup>-2</sup> | ns |
| UV-flies | Rh3-*norpA* rescue | 6 | 9 | -1.012 | Mann–Whitney | 4.233 x 10<sup>-1</sup> | ns |
| UV-flies | Rh4-*norpA* rescue | 6 | 10 | -1.498 | Mann–Whitney | 6.217 x 10<sup>-2</sup> | ns |
| UV-flies | Rh5-*norpA* rescue | 6 | 10 | -1.797 | Mann–Whitney | 1.434 x 10<sup>-2</sup> | * |
| UV-flies | Rh6-*norpA* rescue | 6 | 7 | -1.749 | Mann–Whitney | 2.524 x 10<sup>-2</sup> | * |
| UV-flies | *norpA*<sup>36</sup>-mutant | 6 | 12 | -2.074 | Mann–Whitney | 1.056 x 10<sup>-2</sup> | * |
| *dSK* | *hdc*<sup>JK910</sup> | 20 | 8 | -3.221 | Mann–Whitney | 1.750 x 10<sup>-3</sup> | ** |
| *dSK* | *ninaE*<sup>8</sup> | 20 | 20 | -2.998 | Mann–Whitney | 1.362 x 10<sup>-5</sup> | *** |
| *dSK* | Rh3-6-*norpA* rescue | 20 | 20 | -3.648 | Mann–Whitney | 2.141 x 10<sup>-6</sup> | *** |
| *dSK* | Rh1-*norpA* rescue | 20 | 8 | -1.256 | Mann–Whitney | 4.152 x 10<sup>-2</sup> | * |
| *dSK* | Rh3-*norpA* rescue | 20 | 9 | -4.287 | Mann–Whitney | 5.395 x 10<sup>-4</sup> | *** |
| *dSK* | Rh4-*norpA* rescue | 20 | 10 | -4.772 | Mann–Whitney | 3.002 x 10<sup>-4</sup> | *** |
| *dSK* | Rh5-*norpA* rescue | 20 | 10 | -5.071 | Mann–Whitney | 3.002 x 10<sup>-4</sup> | *** |
| *dSK* | Rh6-*norpA* rescue | 20 | 7 | -5.023 | Mann–Whitney | 2.015 x 10<sup>-3</sup> | ** |
| *dSK* | *norpA*<sup>36</sup>-mutant | 20 | 12 | -5.349 | Mann–Whitney | 9.225 x 10<sup>-5</sup> | *** |
| *hdc*<sup>JK910</sup> | *ninaE*<sup>8</sup> | 8 | 20 | 2.231 x 10<sup>-1</sup> | Mann–Whitney | 3.927 x 10<sup>-1</sup> | ns |
| *hdc*<sup>JK910</sup> | Rh3-6-*norpA* rescue | 8 | 20 | -4.268 x 10<sup>-1</sup> | Mann–Whitney | 3.835 x 10<sup>-2</sup> | * |
| *hdc*<sup>JK910</sup> | Rh1-*norpA* rescue | 8 | 8 | 1.965 | t-test | 1.625 x 10<sup>-2</sup> | * |
| *hdc*<sup>JK910</sup> | Rh3-*norpA* rescue | 8 | 9 | -1.066 | t-test | 3.816 x 10<sup>-4</sup> | *** |
| *hdc*<sup>JK910</sup> | Rh4-*norpA* rescue | 8 | 10 | -1.551 | t-test | 7.458 x 10<sup>-6</sup> | *** |
| *hdc*<sup>JK910</sup> | Rh5-*norpA* rescue | 8 | 10 | -1.850 | t-test | 2.224 x 10<sup>-5</sup> | *** |
| *hdc*<sup>JK910</sup> | Rh6-*norpA* rescue | 8 | 7 | -1.802 | Mann–Whitney | 1.450 x 10<sup>-2</sup> | * |
| *hdc*<sup>JK910</sup> | *norpA*<sup>36</sup>-mutant | 8 | 12 | -2.128 | t-test | 2.364 x 10<sup>-5</sup> | *** |
| *ninaE*<sup>8</sup> | Rh3-6-*norpA* rescue | 20 | 20 | -6.499 x 10<sup>-1</sup> | Mann–Whitney | 2.332 x 10<sup>-3</sup> | ** |
| *ninaE*<sup>8</sup> | Rh1-*norpA* rescue | 20 | 8 | 1.742 | Mann–Whitney | 8.961 x 10<sup>-3</sup> | ** |
| *ninaE*<sup>8</sup> | Rh3-*norpA* rescue | 20 | 9 | -1.289 | Mann–Whitney | 2.214 x 10<sup>-3</sup> | ** |
| *ninaE*<sup>8</sup> | Rh4-*norpA* rescue | 20 | 10 | -1.775 | Mann–Whitney | 7.598 x 10<sup>-4</sup> | *** |
| *ninaE*<sup>8</sup> | Rh5-*norpA* rescue | 20 | 10 | -2.073 | Mann–Whitney | 3.002 x 10<sup>-4</sup> | *** |
| *ninaE*<sup>8</sup> | Rh6-*norpA* rescue | 20 | 7 | -2.026 | Mann–Whitney | 2.015 x 10<sup>-3</sup> | ** |
| *ninaE*<sup>8</sup> | *norpA*<sup>36</sup>-mutant | 20 | 12 | -2.351 | Mann–Whitney | 9.225 x 10<sup>-5</sup> | *** |
| Rh3-6-*norpA* rescue | Rh1-*norpA* rescue | 20 | 8 | 2.392 | Mann–Whitney | 9.760 x 10<sup>-4</sup> | *** |
| Rh3-6-*norpA* rescue | Rh3-*norpA* rescue | 20 | 9 | -6.389 x 10<sup>-1</sup> | Mann–Whitney | 1.056 x 10<sup>-2</sup> | * |
| Rh3-6-*norpA* rescue | Rh4-*norpA* rescue | 20 | 10 | -1.125 | Mann–Whitney | 6.595 x 10<sup>-4</sup> | *** |
| Rh3-6-*norpA* rescue | Rh5-*norpA* rescue | 20 | 10 | -1.423 | Mann–Whitney | 3.002 x 10<sup>-4</sup> | *** |
| Rh3-6-*norpA* rescue | Rh6-*norpA* rescue | 20 | 7 | -1.376 | Mann–Whitney | 2.015 x 10<sup>-3</sup> | ** |
| Rh3-6-*norpA* rescue | *norpA*<sup>36</sup>-mutant | 20 | 12 | -1.701 | Mann–Whitney | 9.225 x 10<sup>-5</sup> | *** |
| Rh1-*norpA* rescue | Rh3-*norpA* rescue | 8 | 9 | -3.031 | t-test | 2.015 x 10<sup>-3</sup> | ** |
| Rh1-*norpA* rescue | Rh4-*norpA* rescue | 8 | 10 | -3.516 | t-test | 8.761 x 10<sup>-4</sup> | *** |
| Rh1-*norpA* rescue | Rh5-*norpA* rescue | 8 | 10 | -3.815 | t-test | 7.377 x 10<sup>-4</sup> | *** |
| Rh1-*norpA* rescue | Rh6-*norpA* rescue | 8 | 7 | -3.767 | Mann–Whitney | 1.450 x 10<sup>-2</sup> | * |
| Rh1-*norpA* rescue | *norpA*<sup>36</sup>-mutant | 8 | 12 | -4.093 | t-test | 5.395 x 10<sup>-4</sup> | *** |
| Rh3-*norpA* rescue | Rh4-*norpA* rescue | 9 | 10 | -4.857 x 10<sup>-1</sup> | t-test | 1.768 x 10<sup>-2</sup> | * |
| Rh3-*norpA* rescue | Rh5-*norpA* rescue | 9 | 10 | -7.842 x 10<sup>-1</sup> | t-test | 4.833 x 10<sup>-4</sup> | *** |
| Rh3-*norpA* rescue | Rh6-*norpA* rescue | 9 | 7 | -7.367 x 10<sup>-1</sup> | Mann–Whitney | 1.130 x 10<sup>-2</sup> | * |
| Rh3-*norpA* rescue | *norpA*<sup>36</sup>-mutant | 9 | 12 | -1.062 | t-test | 1.250 x 10<sup>-4</sup> | *** |
| Rh4-*norpA* rescue | Rh5-*norpA* rescue | 10 | 10 | -2.985 x 10<sup>-1</sup> | t-test | 6.217 x 10<sup>-2</sup> | ns |
| Rh4-*norpA* rescue | Rh6-*norpA* rescue | 10 | 7 | -2.510 x 10<sup>-1</sup> | Mann–Whitney | 1.868 x 10<sup>-1</sup> | ns |
| Rh4-*norpA* rescue | *norpA*<sup>36</sup>-mutant | 10 | 12 | -5.763 x 10<sup>-1</sup> | t-test | 1.686 x 10<sup>-3</sup> | ** |
| Rh5-*norpA* rescue | Rh6-*norpA* rescue | 10 | 7 | 4.752 x 10<sup>-2</sup> | Mann–Whitney | 4.847 x 10<sup>-1</sup> | ns |
| Rh5-*norpA* rescue | *norpA*<sup>36</sup>-mutant | 10 | 12 | -2.778 x 10<sup>-1</sup> | t-test | 5.395 x 10<sup>-4</sup> | *** |
| Rh6-*norpA* rescue | *norpA*<sup>36</sup>-mutant | 7 | 12 | -3.253 x 10<sup>-1</sup> | Mann–Whitney | 6.316 x 10<sup>-3</sup> | ** |

**Table S5.**
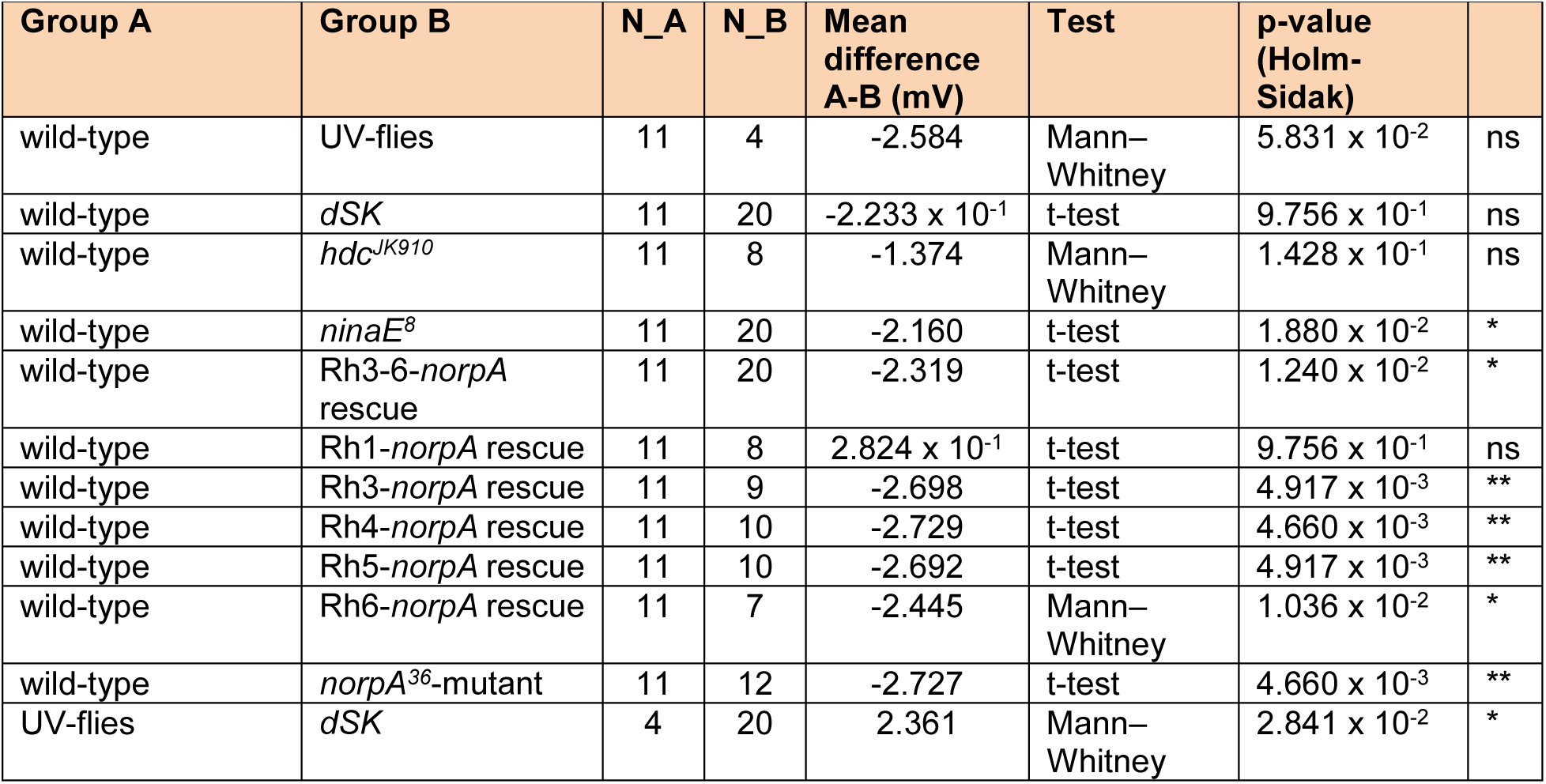
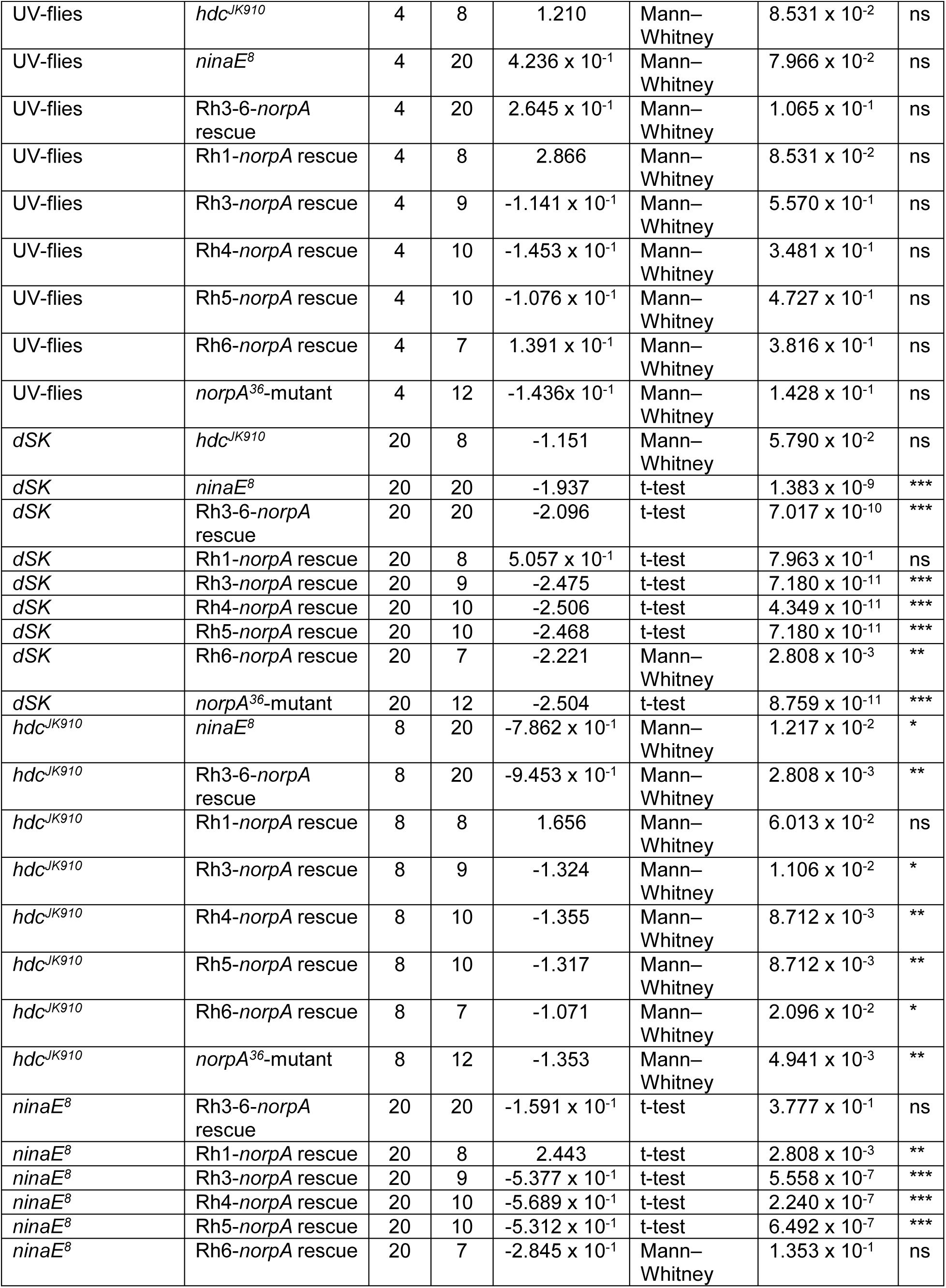
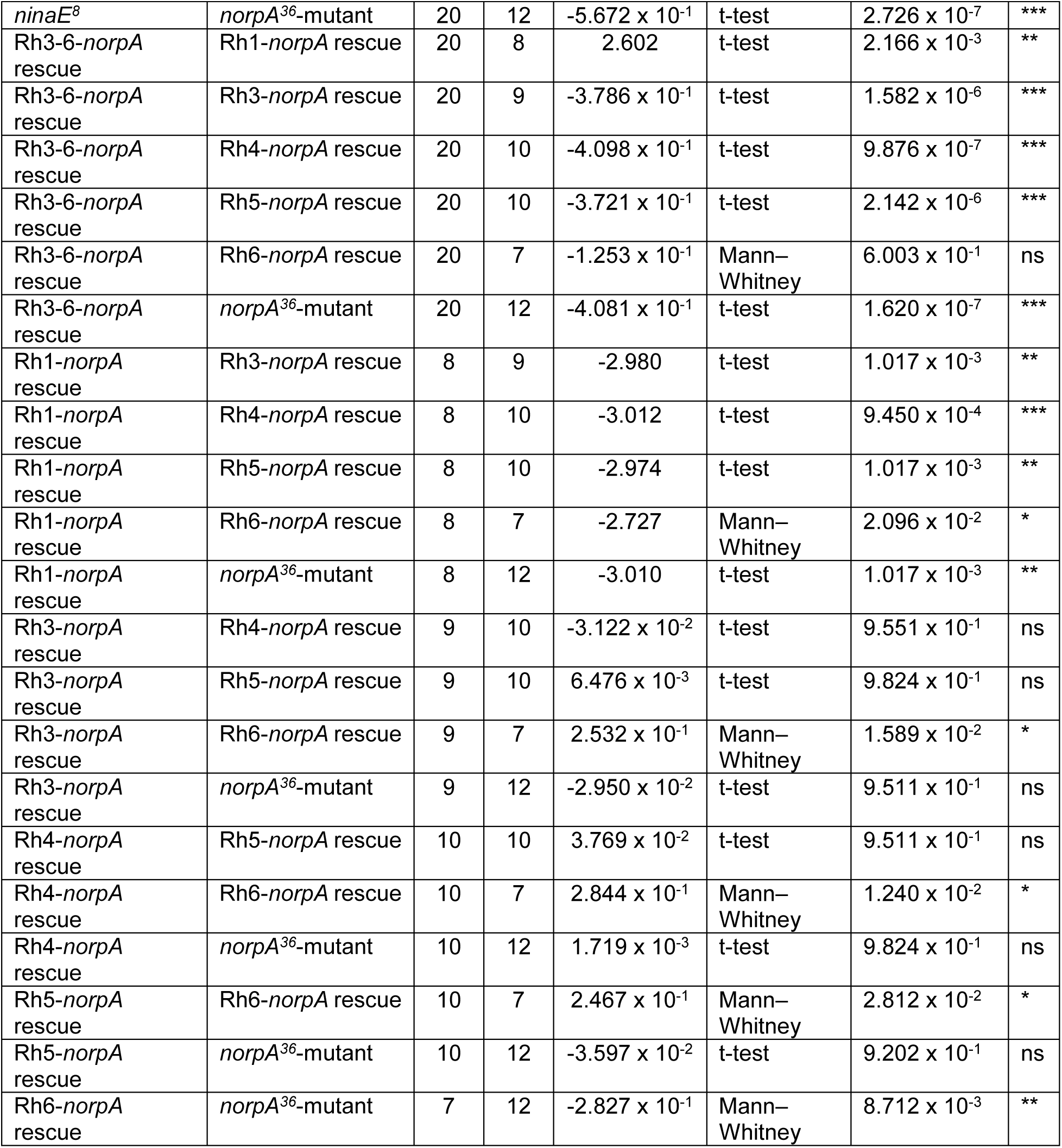
ERG-responses to 200ms Green flash

###### II.8.ii Cornea-neutralization imaging R1-R7/8 photomechanics inside individual ommatidia

To complement DPP imaging, which merges optically superpositioned individual rhabdomeres of different ommatidia into a single virtual image (21, 24), we further used the cornea-neutralization method (24) to examine light-induced Drosophila R1-R7/8 rhabdomere movements inside individual ommatidia directly (Fig. S31). The purpose of these experiments was to test how two different spatially-restricted light patterns (field or spot) activate photoreceptor microsaccades on the eye surface locally. The experiments were done with a separate bespoke imaging system built around an upright microscope (Olympus BX51), secured to an x,y-stage on an anti-vibration table (MellesGriot, UK). The system was light-shielded inside a black Faraday cage with black lightproof curtains covering its frontal opening, and the experiments were done in a dark room to minimize light pollution. Rhabdomeres were viewed with a 40x water immersion objective (Zeiss C Achroplan NIR 40x/0.8 w, ∞/0.17, Germany) and recorded with a high-speed camera (Andor Zyla, UK) at 100 frames/s.

A *Drosophila* was gently fastened to an enlarged fine-end of a 1 ml pipette tip, as explained in Section I.1. The fly was then positioned with a remote-controlled x,y,z-fine resolution micromanipulator (Sensapex, Finland) underneath the water immersion objective (Fig. S31 *A* and *B*), using a live video stream on a computer monitor.

***Rhabdomere imaging*** Antidromic illumination from a high-power IR light source (740 nm LED with 720 nm high-pass edge-filter, driven by Cairn OptoLED, UK) was delivered transcuticularly through the fly head, revealing R1-R7/8 rhabdomeres inside local ommatidia (Fig. S31*C*). R1-R7/8 are effectively insensitive to >720 nm red light (11).

**Fig. S31.**
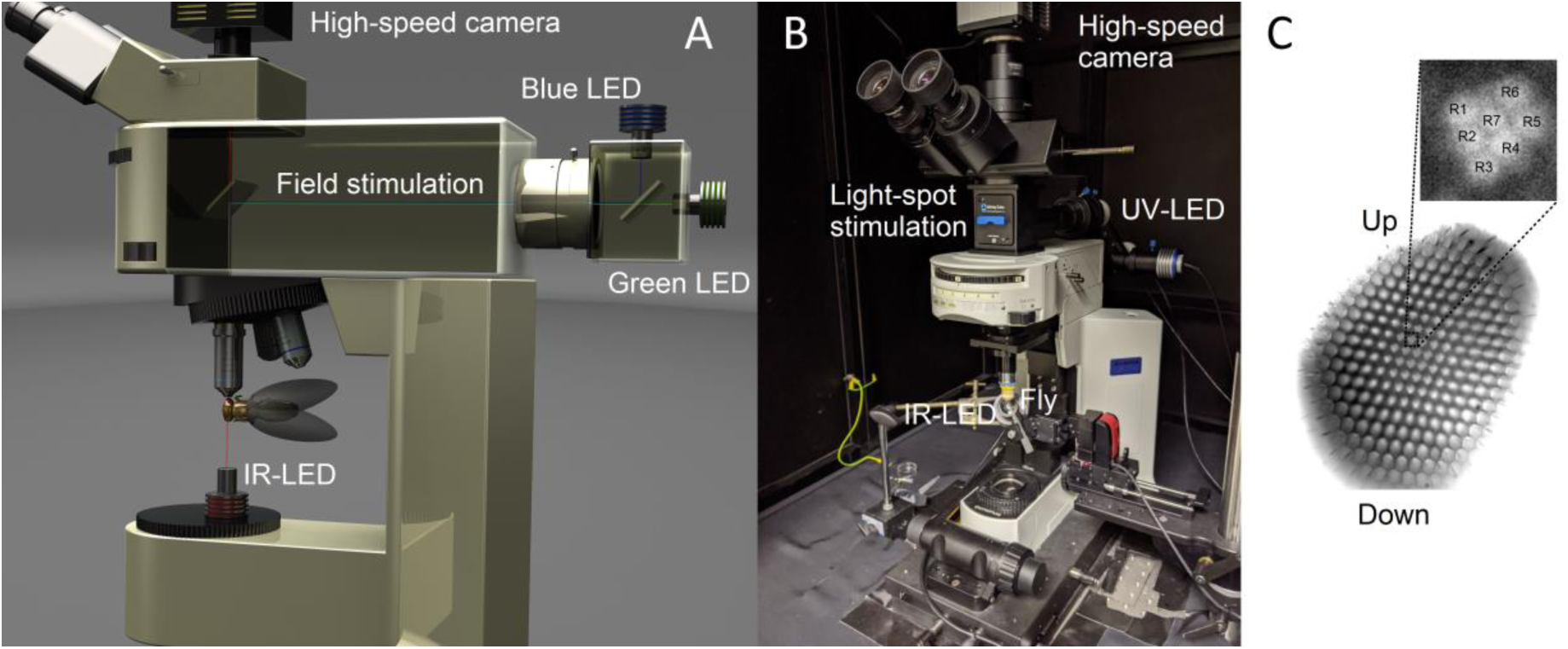
*In vivo* orthodromic light stimulation of local ommatidial rhabdomeres in cornea-neutralized *Drosophila* eyes. (*A*) *Light-field stimulation* was delivered by blue and green LEDs, mounted in the microscope’s optical back-port, and controlled by two LED drivers. Blue and green stimuli were first fused (by a beam-splitter) and then directed (via another beam-splitter) to the 40x-objective’s center and focused on the rhabdomere tips. (*B*) *Light-spot stimulation* was delivered by a Blue (470 nm) or UV (365 nm) LED through an optical pinhole contraption (Infinity-Cube, Cairn Research, UK) mounted between the microscope’s turret and ocular pieces. This stimulation mode enabled highly localized flash-activation of only a few photoreceptors at a time. (*C*) A typical high-speed camera’s field of view of local neighboring ommatidial rhabdomeres, recorded under continuous IR-LED illumination, which does not activate photoreceptors (11). IR-imaging allowed us to capture unhindered local microsaccades (rhabdomere movements) to both the blue/green-field and blue-spot stimulation with minimal recording artifacts. Insert: the photoreceptors, which participated in the resulting microsaccades, could be identified across the ommatidia at the resolution of single rhabdomeres at each time-point (video frame), and their local dynamics revealed by cross-correlation analyses. *C* modified and adapted from (11).

***Microsaccade activation*** We flashed two different orthodromic stimuli: (**i**) *light-field* and (**ii**) *light-spot* through the 40x-objective onto the left *Drosophila* eye to evoke local photoreceptor microsaccades.

i. Two high-power LEDs delivered the field stimulation: 470 nm (blue) and 545 nm (green), each separately controlled by its own driver (Cairn OptoLED, UK) (Fig. S31*A*). These peak wavelengths were selected to activate R1-R6s’ rhodopsin (Rh1) and its meta-form near maximally. Thereby, through their joint stimulation, we minimized desensitization by prolonged depolarizing after-potentials (PDA) (88). Light from the two LEDs was merged into one focused beam by a 495 nm dichroic mirror and low-pass-filtered at 590 nm. The ommatidial rhabdomere images were split spectrally by another dichroic mirror (600 nm). As a result, effectively, only red image intensity information (≥600 nm) was sampled by the high-speed camera.
ii. x,y-position adjustable pinhole/beam-splitter optics (Cairn Infinity-Cube, UK) produced a ∼5 µm light-spot on rhabdomeres inside single ommatidia (Fig. S31*B*). This contraption was placed in the light path between the high-speed camera and the objective. It shaped and split the light from a high-power blue (470 nm) or UV (365 nm) LED (controlled by a Cairn OptoLED driver) to rhabdomeres while letting IR images be sampled before, during, and after their microsaccade activation.

***Microsaccades to light-field stimulation.*** We delivered 10 ms blue/green field stimulus flashes, separated with ∼3-minute dark periods, on a dark-adapted fly eye’s local surface area. The field stimulus covered the 40x-objective’s field of view, which was simultaneously imaged under continuous IR-light, exposing R1-R7/8 rhabdomere tips inside about 90-150 ommatidia. The exact configuration varied from one fly preparation to another, as limited by the fly mounting, pipette positioning angles, and the local eye curvature at the different imaged eye locations.

Individual rhabdomere movements inside single ommatidia were analyzed from the high-speed light-field video recordings offline. We hand-marked 90-150 individual ommatidia (Fig. S31*C*) in these videos, and the cross-correlation analysis was performed separately for each ommatidium’s rhabdomeres (see Supplement II.2.iii. for further details). Characteristically, the intra-ommatidial rhabdomere contractions (photoreceptor microsaccades) to light-field stimulation peaked within 80-140 ms after the 10 ms light flash. The rhabdomere movement noise, as analyzed 40-160 ms before the flash, was subtracted from the maximum photoreceptor microsaccade values. The ommatidia that showed smaller-than-noise motion were considered to be still (not photomechanically contracting), with their rhabdomeres not being light-activated.

The field-stimulus flashes (Fig. S32, *A* to *C*) evoked the strongest photoreceptor microsaccades in the ommatidia at the stimulus/image center, pointing directly towards the orthodromic light-field stimulator and thus experiencing direct incident light (Fig. S32, *D* to *F*). Further away from the stimulus center the intraommatidial rhabdomeres resided, the smaller (Fig. S32*E*) and slower (Fig. S32*F*; *cf.* the lognormal fits to WT fly #1 microsaccades) their microsaccades were. Thus, the rhabdomeres in the ommatidia, which were about 100-µm-distance from the stimulus center, remained practically still, producing no noticeable photomechanical movements (Fig. S32, *D* to *F*). Nonetheless, owing to the imaging system’s extreme sensitivity, some preparations/configurations inadvertently generated minute (10-70 nm) mechanical jitter. This slight extrinsic resonance superimposed the same temporal (synchronized) noise pattern on all the simultaneously recorded intraommatidial photoreceptor microsaccades across the eye (Fig. S32; *cf.* WT flies #2-3). However, such sporadic recording noise did not bias the general results of the local spatially-constrained microsaccade activation dynamics, which were repeatedly observed in different fly preparations.

The *Drosophila* compound eyes’ two well-known architectural factors (12) best explain the observed spatiotemporal microsaccade-waning over the stimulated/imaged area:

i. because the ommatidial tiling follows each compound eye’s small radius of curvature (Fig. S32*C*), their photoreceptors’ RFs increasingly direct away from the brightest (incident) light
ii. the ommatidial screening pigments in the ommatidial walls block non-incidental light scatter from being absorbed by the rhabdomeres

**Fig. S32.**
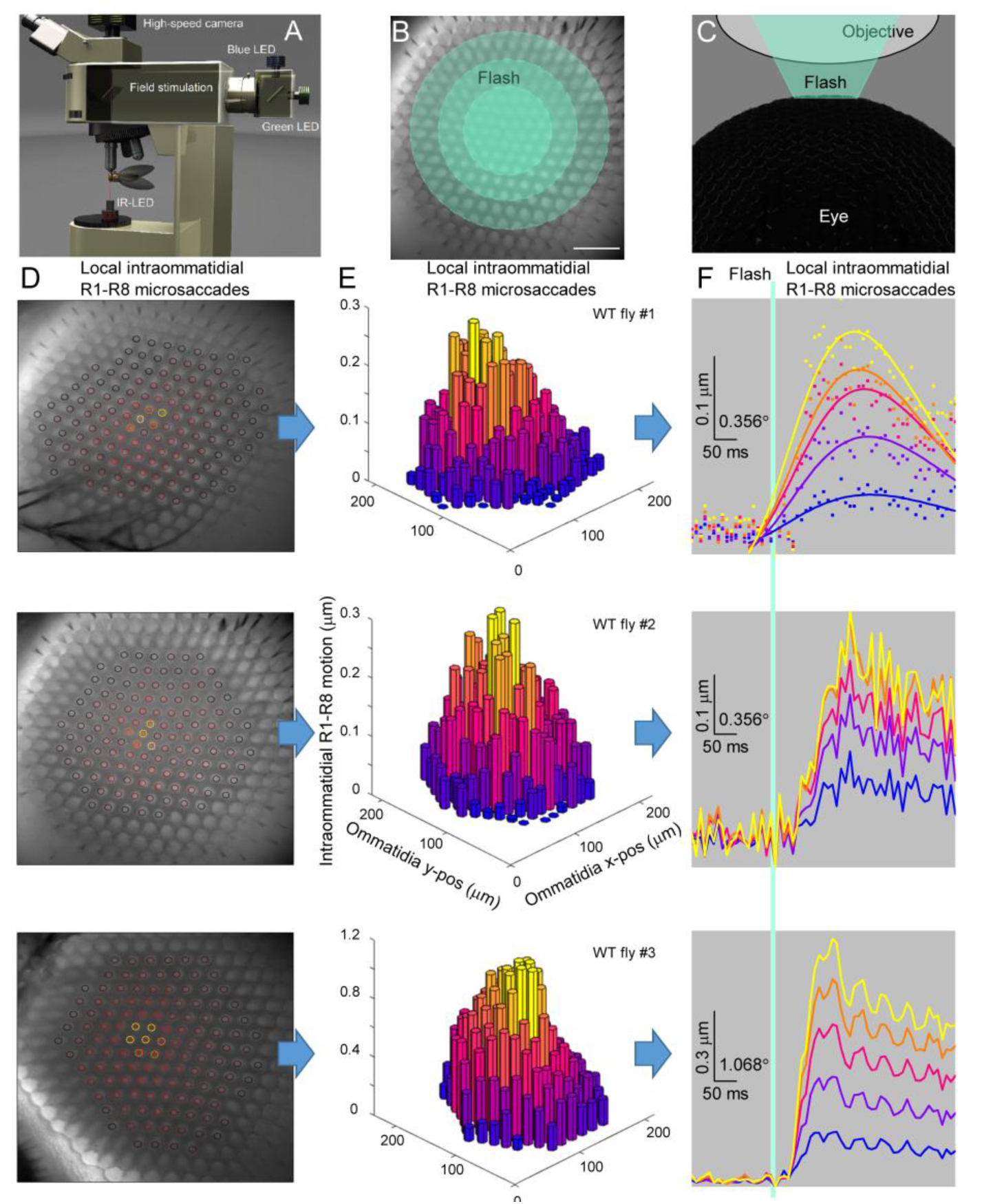
Light-field evoked the strongest photoreceptor microsaccades in the ommatidia directly facing it. (*A*) High-speed cornea-neutralized *Drosophila* eye imaging. Orthodromic blue/green field-flash was used to activate the local rhabdomeres within the microscope’s field of view. (*B*) Ommatidial rhabdomere tips at the image center viewed by the high-speed camera. The concentric green disks represent the decreasing field intensity over the eye surface with the brightest light at the center. Scale bar: 50 µm. (*C*) Because the *Drosophila* eye is spherical, the further away its ommatidia are from the image center, the more their rhabdomeres’ RFs point away from the stimulus. Thus, fewer become light-activated. (*D*) Light-field stimulus evoked the most apparent photomechanical R1-R7/8 rhabdomere movements inside the central ommatidia that directly faced the incident light. These photomechanical hotspots are highlighted by bright yellow and orange circles at the eye image examples (WT flies #1-3), which show three different eye locations/imaging positions. (*E*) Maximal microsaccades inside the ommatidia petered out as a function of distance from the image center, as analyzed from *D*. This positional microsaccade decay resulted from the eye curvature gradually shifting the photoreceptors’ RFs away from the brightest (incident) light. At the same time, the ommatidial screening pigments reduced light scatter. (*F*) Local eye-position-dependent microsaccades followed the characteristic light-intensity-dependent dynamics, decreasing amplitude away from the image center. Intraommatidial microsaccades were sorted by their amplitude into five groups (as in *E*). The traces show the group’s average response waveforms and their lognormal fits for fly#1. Mechanical jitter in the fly preparation/imaging system caused the synchronized (oscillating) 10-70 nm noise patterns on fly#2’s and fly#3’s microsaccadic responses. The green bar indicates a 10 ms field-stimulus flash.

The observed local microsaccades’ movement directions matched those mapped by DPP imaging (see Section II.1., above). Such spatiotemporal local and global eye-map correspondences connote that intrinsic eye muscle activity, which would have moved all retinal rhabdomeres together, had little influence in these and the DPP recordings (Fig. S16). Therefore, the observed field-stimulus-induced microsaccades were photomechanical, with the photoreceptors’ light absorption probability regulating their strength and velocity (Fig. S32 *D* and *E*). However, the maximum microsaccade amplitudes varied trial-to-trial and between individual flies (Fig. S32 *D* and *E*; *cf.* WT flies #1-3), sometimes considerably. Therefore, it is not inconceivable that *Drosophila*’s intrinsic/diurnal activity state, via feedback synapses from the higher brain centers (30, 32), could co-regulate R1-R7/8-microsaccade gain, similar to what has already been shown for R1-R6 photoreceptor voltage responses (20, 33, 35, 36, 78, 79).

These observations and results, confirmed by imaging many *Drosophila* eyes (n = 15 flies; both the left and right eyes) at different corneal locations (Fig. S32), concur with the results from the X-ray and the DPP imaging experiments (see Section I and Sections II.1-7, above), respectively. They are also consistent with our earlier published data (11).

***Microsaccades to light-spot stimulation*** We managed to light-activate the photoreceptor rhabdomeres in single ommatidia with the light-spot stimulation, generating 0.1-0.15 µm microsaccades (Fig. S33), while the intraommatidial rhabdomeres across the rest of the eye remained practically still. These exceedingly local microsaccades reached their peak amplitudes ∼40-80 ms after the flash onset (Fig. S33*E*). They showed somewhat faster dynamics than the microsaccades to the field stimulus flashes (Fig. S32*E*), which peaked 80-140 ms after the flash and had longer decay times (> 100 ms).

Single photoreceptor light-activation caused small-amplitude microsaccades where all R1-R7/8s moved collectively inside one ommatidium. These technically challenging results are consistent with the *norpA*- rhodopsin-rescue results (see Fig. S28 in Section II.8.i, above). However, we only obtained dominant “single-ommatidium” microsaccades in 3 out of 15 tested wild-type flies (Fig. S33 *D* and *E*), as in the other 12 preparations, photoreceptor microsaccades were either also seen in the near-neighboring ommatidia (n = 2) or could not be accurately resolved (n = 10). Notably, all the 15 flies - including the shown examples (WT flies #1-3) - showed consistent photoreceptor microsaccades to light-field stimulation within a broader ommatidium population (*cf*. Fig. S32*E*). We found two primary reasons for the light-spot stimulation experiment’s low success:

i. In some fly preparations, the photoreceptor microsaccades to the light-field stimulation were already relatively small (≤0.3 µm). Consequently, the much dimmer light-spot stimulation failed to evoke reliable/measurable responses (*i.e.,* microsaccades larger than the recording noise).
ii. When the maximum microsaccade amplitudes to spot-stimulus were ≥0.15 µm, the rhabdomeres in the adjacent ommatidia also contracted photomechanically, although the light-spot was smaller than the ommatidium, as seen at the microscope’s focal plane. However, the conical light beam (from the microscope objective) penetrated 30-40 µm into the eye (Fig. S33*C*). Therefore, unavoidably, some scattered light reached the near-neighboring ommatidia, making their photoreceptors generate photomechanical microsaccades together with the photoreceptors in the directly stimulated center ommatidium.

Here, the largest single-ommatidium-activated microsaccades were evoked by a spot-stimulus, which was focused right at its next-door ommatidium (Fig. S33). These findings further indicate that a light-spot at the stimulated sub-ommatidial area could cross over its microscope-observed focal-plane boundaries (due to scattering) in the used experimental configuration. It then light-activated photoreceptors also in its nearest ommatidial neighbor.

**Fig. S33.**
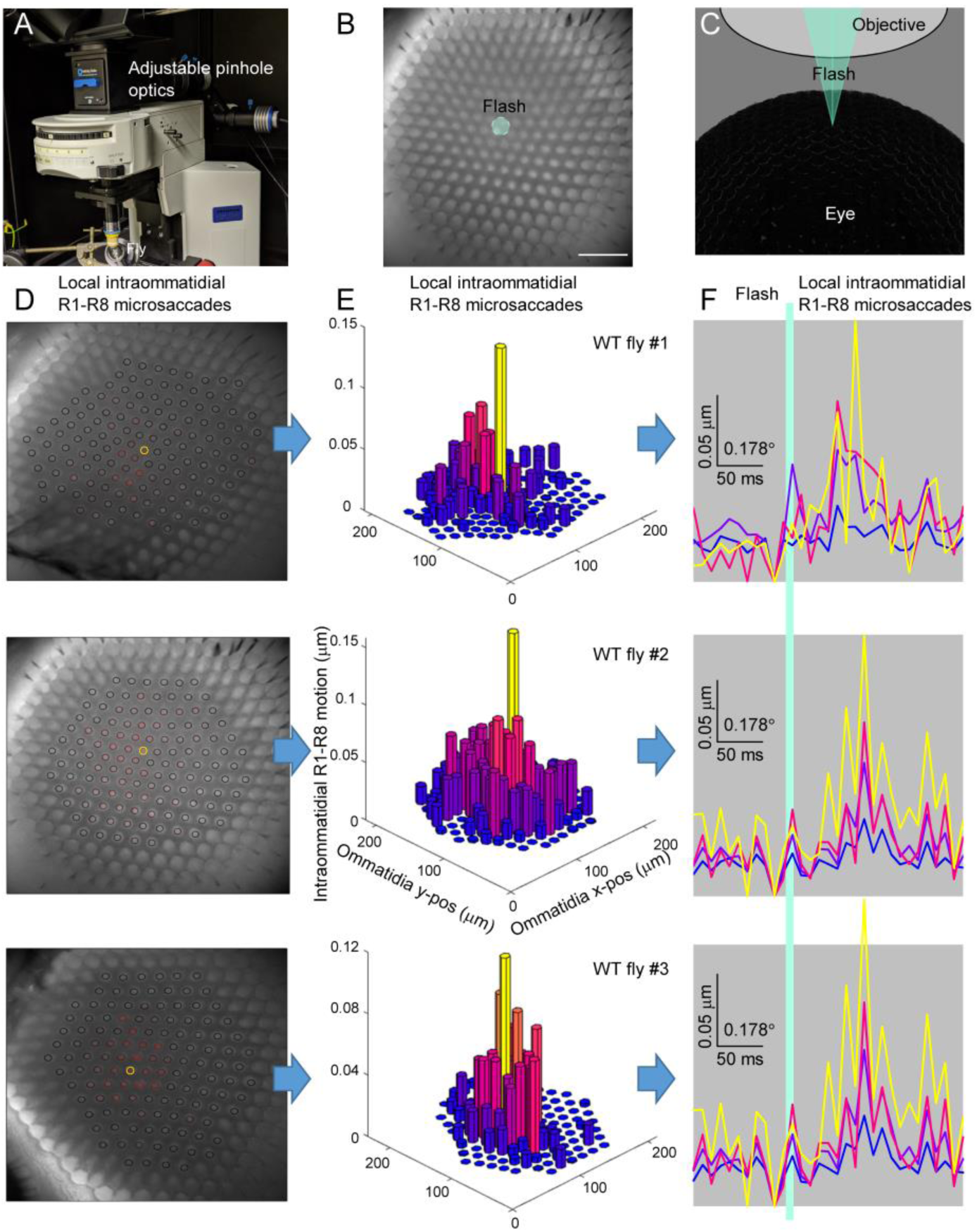
Light-spot evoked photoreceptor microsaccades only in the ommatidia experiencing incident light. (*A*) Blue LED light (470 nm) was passed through a small iris (pinhole) to generate a small spot, which was directed to the ommatidia by a dichroic mirror. IR light was passed through the fly head to the camera via the microscope objective. (*B*) Transparent green disk indicates a typical local eye surface area, containing only a few ommatidia, inside which the spot stimulation was tested. The actual stimulus spot (diameter: ∼5 µm) showed only faintly in the IR image (now hidden inside the green disk). Scale bar: 50 µm. (*C*) Because of the narrow spot-stimulus beam and the eye’s curvature, only the rhabdomeres in the very central ommatidia are light-activated. (*D*) Spot stimulus evoked the largest photoreceptor microsaccades in a single ommatidium (shown in yellow), while photoreceptors in most neighboring ommatidia remained still. (*E*) The maximum photoreceptor microsaccades inside a single ommatidium (shown in yellow) were significantly higher than in the rest of the ommatidia. (*F*) Single ommatidial photoreceptor microsaccades to 10 ms light-spot flash (green bar) lasted only about ∼50 ms. The microsaccades were sorted into five groups by their amplitudes (as in *D*). The microsaccades at the center of each imaged eye (yellow traces) were significantly larger than in the neighboring ommatidia (red to blue) further away.

#### III High-speed optical imaging of eye-muscle-induced whole retina movements and antennae castings

**Overview**

This section describes the deep pseudopupil (DPP)-based approaches to measure whole retinal movements in living head-immobilized *Drosophila*. It gives central background information and additional supporting evidence for the results presented in the main paper, including:

- In our recording configurations, the whole-retina movements occur rarely. And when they do, they show considerably slower dynamics than the photomechanical photoreceptor microsaccades. Therefore, eye-muscle-induced whole-retinal movements could only have negligible effects on the microsaccadic sampling dynamics presented in this paper.
- Light-stimulus can trigger antennal “casting movements” in some flies. But these movements only happen after the photoreceptor microsaccades, at the earliest about 40-50 ms later.
- Puffing air to the antennae invariably triggers their casting movements. But neither air puffs nor antennae casting evokes photoreceptor microsaccades or whole retina movements.

It is not known if a fly can - by will - induce DPP movements. Photoreceptor microsaccades (via synaptic feedbacks), eye-muscle-induced whole retina movements (via attentive top-down regulation), or both, might also be intrinsically elicited by other sensory inputs such as airflow and olfaction. For example, a fly might use retinal movements and other directional sensing (such as antennal casting) to get a better idea of the object it is just encountering. Theoretically, neural control of such time-locked information sampling could be voluntary or involuntary and vary with a fly’s attentive state. After all, while integrating multisensory information reduces uncertainty, increasing fitness, its execution costs extra energy. Therefore, this optimization is likely complex, non-generic, and suboptimal for both information and energy, leading to different adaptive behaviors in different conditions.

To test these general concepts, we performed high-speed imaging of DPP and antennae movements using pipette-tip-held *Drosophila* in the goniometric imaging system (see Fig. S10) in different experimental configurations.

##### III.1 Recording eye-muscle induced DPP movements in darkness and under steady illumination

Long-term (10 min long) high-speed DPP recordings (Fig. S34*A*) in darkness (Fig. S34B, blue traces, darker = left eye, lighter = right eye) and ambient light (Fig. S34*C*, green) revealed sporadic eye-muscle-induced whole retina movements in which dynamics varied from fly to fly. These movements included slow drifts (fly 1), synchronized binocular vergence motion (fly 2), unsynchronized binocular vergence motion (fly 3), saccadic jumps (fly 3 and 5), and any combinations of these (flies 4 to 6). Generally, the dynamics in the dark- and light-adapted eyes were slow, occurring mainly in the seconds-to-minutes time scale. Consequently, their power spectra (Fig. S34*D*), having 0.1 Hz median frequencies (Fig. S34*E*), are about 10-times more low-passed (slower) than the corresponding photoreceptor microsaccade metrics (*cf*. Fig. S24). Thus, when inspected in a shorter time scale, it becomes evident how much slower and smaller the spontaneous eye-muscle-induced slow retina drifts (Fig. S34F) and saccadic movements (Fig. S34*G*) are than the light-flash triggered photomechanical photoreceptor microsaccades (Fig. S34*H*).

**Fig. S34.**
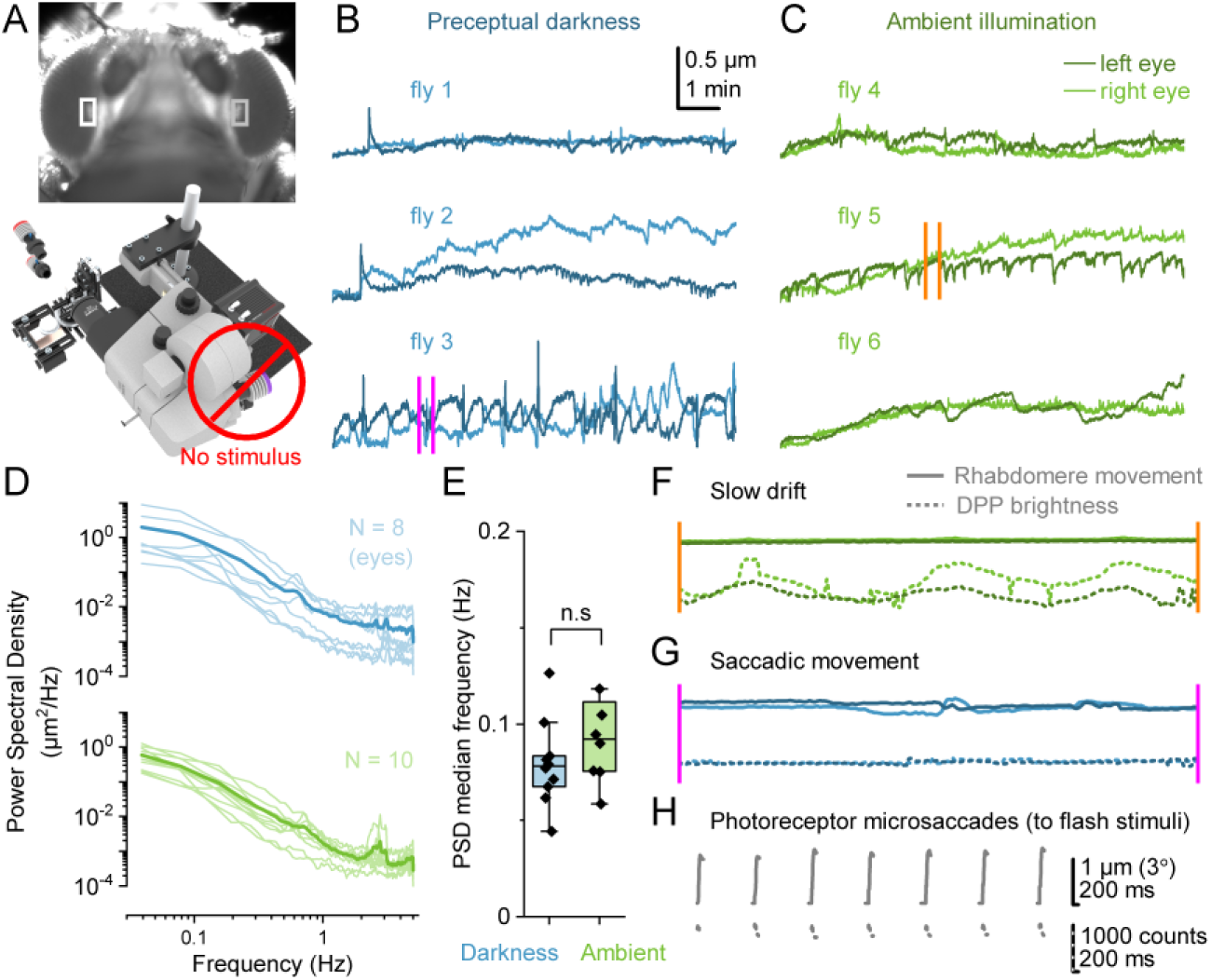
Head-immobilized flies show slow spontaneous eye-muscle-induced whole retina movements. (*A*) Using the goniometric high-speed imaging system, we recorded deep pseudopupil (DPP) movements induced by spontaneous eye-muscle activity (*i.e*., no light-flash stimulation) in darkness (blue traces) and ambient light (green). Each recording lasted 10 min. (*B*) Three examples of eye-muscle induced whole retina movements in relative darkness, showing slow drifts, binocular and monocular vergence motion, and saccadic events. (*C*) Three similar examples of eye-muscle induced whole retina movements in ambient light. (*D*) The power spectra of these whole retina movements in darkness and ambient light are strongly low-passed, indicating the dominant presence of slow (low-frequency) events. (*E*) The corresponding median frequencies are around 0,1 Hz with no significant difference between darkness and the ambient light background. (*F* and *H*) In a 20-s-long time window, the whole retina drifts (*F*) and saccadic movements (*G*) appear strikingly slower and typically smaller than the fast flash-triggered photoreceptor microsaccades (*H*). Examples are taken from fly 3 and 5 recordings, marked by orange and pink lines in *B* and *C*. Continuous and dotted lines show the corresponding lateral and axial DPP movement dynamics. Notice the eye-muscle-induced retina movements are dominated by very slow axial DPP darkening/lightening drifts.

##### III.2 Simultaneous recording of DPP microsaccades and light-triggered antennal casting

Flies can cast their antennae following light stimulation. Recording DPP microsaccades and any co-occurring antennae movements, we found the light-triggered antennae castings not a reflex but a fly-dependent (intrinsic) phenomenon (Fig. S35*A*). Some flies showed antennae castings to UV light flashes; others did not (i, right). Moreover, even in a single fly, these dynamics were highly variable (ii, left). However, crucially, the light-triggered antennal castings happened only after the DPP microsaccades (Fig. S35*B*), showing an absolute 40-50 ms neural delay. The DPP microsaccade (blue) and antennae casting (yellow) statistics quantified this delay in their correspondingly shifted latencies (Fig. S35*C*) and half-rise times (Fig. S35*D*).

**Fig. S35.**
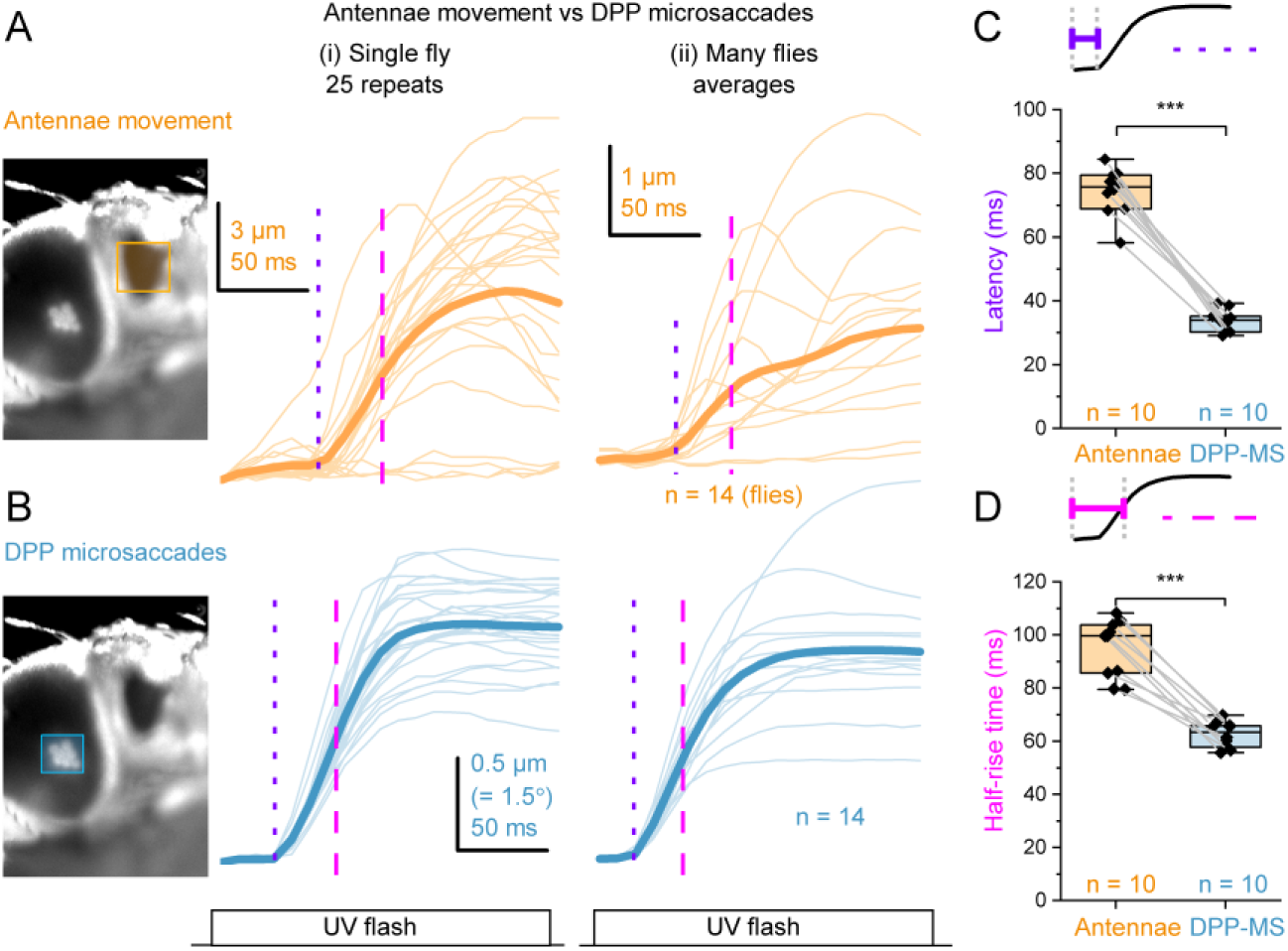
Some *Drosophila* generate antennae casting 40-50 ms after photoreceptor microsaccades. We simultaneously high-speed video recorded (*A*) antennae movements and (*B*) deep pseudopupil (DPP) microsaccades to a bright repeated 190 ms UV flash. (*A*) A light flash can trigger antennae casting, but these movements vary significantly in their size and time course. The thick yellow lines show the means for a single fly (left) and the tested population (right). Sometimes a flash triggers antennae casting, other times not. Equally, some flies show them, others do not. There are also sporadic (rare) “anticipatory” castings happening before the light flash. The given µm-scale relates to the recorded image pixels. (*B*) A UV flash always (invariably) evokes a DPP microsaccade that happens about 40-50 ms before the antennae casting (A). The vertical dotted and dashed lines indicate the corresponding latencies and high-rise times, respectively. However, the absolute DPP microsaccade size varies between flash repetitions and the tested flies. The thick blue lines show the means. Note, the given µm-scale relates the physical rhabdomere size, (*C*) The latency from the UV stimulus onset to the apparent start of the DPP microsaccade (blue) is always about 40-50 ms shorter than the start of the antennal casting (yellow). (*D*) The DPP microsaccade high-rise time (blue) is always about 40-50 ms shorter than that of the antennal casting (yellow).

##### III.3 Simultaneous recording of DPP and air puff triggered antennal casting in darkness

We next asked the opposite question - whether other sensory stimuli causing antennae casting could also induce DPP movements. In these 40-s-long experiments, performed in darkness (Fig. S36*A*), we repeatedly puffed air to the fly head while simultaneously video recording both the resulting antennae casting (Fig. S36B, left) and any DPP movements (right) that may follow this stimulation.

We found that air puffs invariably caused antennae casting, but neither the puffs nor the antennae movements triggered apparent DPP movements (Fig. S36*C*). When inspected in a briefer time-scale (Fig. 36*D*), the antennae casting revealed variable dynamics, occurring both during (time-synchronized) and between the air puff pulses. Nevertheless, the same recordings showed no apparent DPP microsaccades. Finally, the corresponding pairwise comparisons quantified the statistical significance of these findings (Fig. 36*E*).

**Fig. S36.**
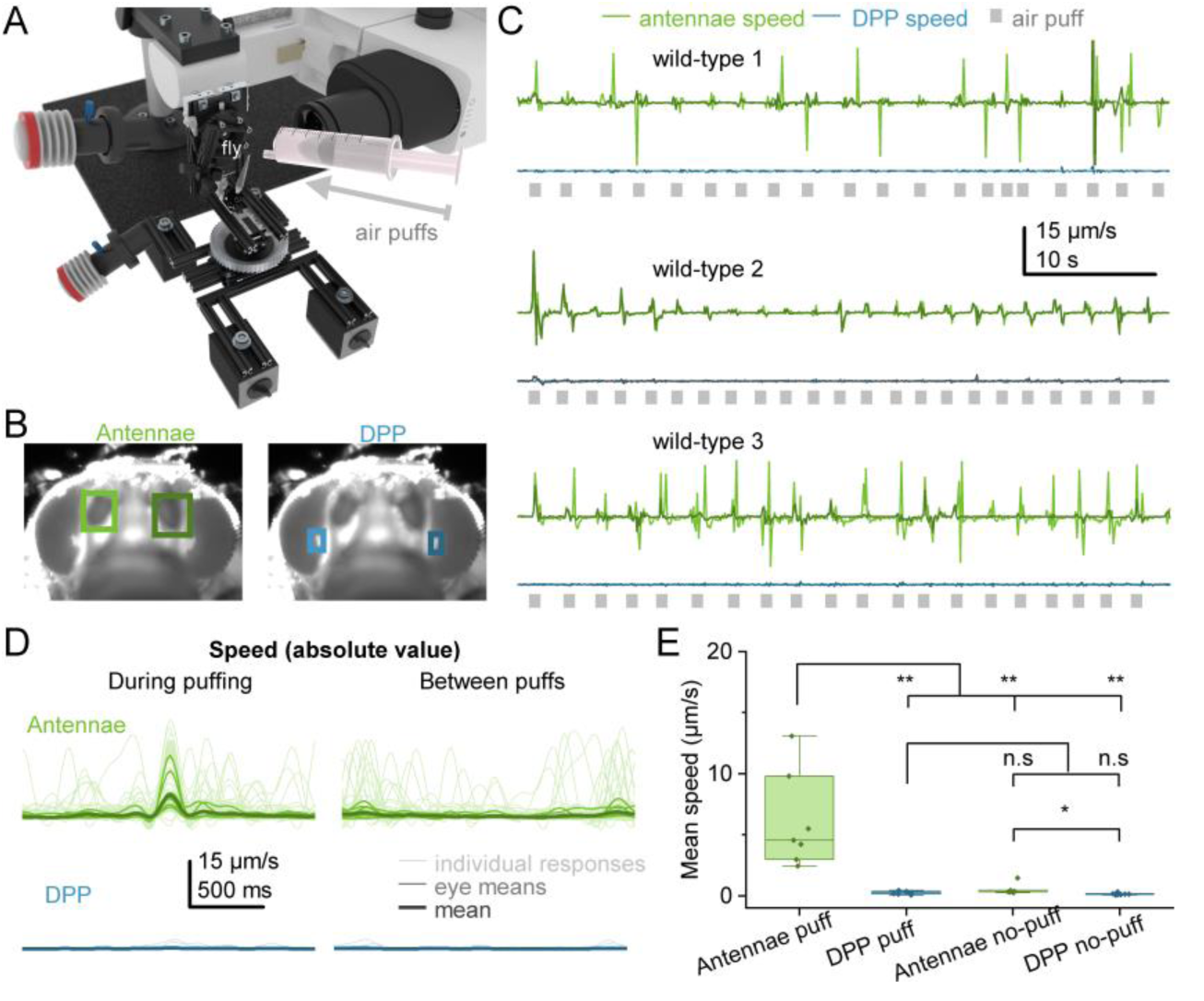
Air-puff-triggered antennae casting does not evoke photoreceptor microsaccades in head-immobilized *Drosophila*. (*A*) Using pipette-tip-held *Drosophila* in the goniometric *in vivo* imaging system (see Fig. S10), we puffed air from a syringe to their antennae. (*B*) We high-speed video recorded the resulting antennae movements (left) and any deep pseudopupil (DPP, right) dynamics that may correlate with them. These movement dynamics were then cross-correlated into time-series recordings (*C*). (*C*) Characteristic long-lasting recordings from three individual flies show ∼1-s-long air puffs (gray squares) reliably triggering antennae casting (green). But neither the air puffing nor the antennae movements resulted in fast DPP microsaccades or slower whole retina movements (blue). Note, the given µm-scale relates to the image pixel-scale (not to the physical rhabdomere size, which is about 10x-times smaller). (*D*) When viewed in a briefer time-scale, antennae movements reveal highly variable dynamics, happening both during (time-synchronized) and between the air puff pulses. However, the DPP recordings showed no apparent fast microsaccades time-locking to air puffing or antennae movements between the puffs. (*E*) Statistical analyses confirmed no link between the air-puff-triggered or the spontaneous antennae movements and the fast DPP microsaccades.

#### IV *In vivo* 2-photon Ca^2+^ imaging L2-neuron responses to hyperacute stimuli

**Overview**

This section describes the experimental and theoretical approaches to measure visual acuity and direction sensitivity of *Drosophila* L2 large monopolar cell terminals in the left and right *medulla*-neuropil using *in vivo* high-speed 2-photon Ca^2+^-imaging. It gives central background information and additional supporting evidence for the results presented in the main paper, including:

- The L2-terminals transmit hyperacute visual information over a broad range of velocities.
- The L2-terminals’ motion-direction sensitivity is broadly co-linear with the microsaccade direction of the photoreceptors transmitting visual information to them.
- Therefore, L2 neurons participate both in encoding and processing hyperacute stereoscopic and optic flow information and channeling these signals to downstream neurons.

We performed 2-photon Ca^2+^-imaging from L2 monopolar cells in UV-flies^13^ or transgenic flies, with natural WT R1-R7/8 photoreceptor visual pigments (Fig. 4). These flies show normal photomechanical photoreceptor microsaccade dynamics (Fig. S37). GCaMP6f was expressed selectively in L2s, and activity changes (fluorescence signals) to visual motion stimuli were imaged at L2 medulla terminals using a laser resonance-scanning microscope (TrimScope, La Vision Biotech, Germany) with 1 NA 40XW objective. The 2-photon excitation source was a mode-locked Ti:Sapphire Mai Tai SP Laser tuned to 920 nm. Fluorescence was collected by a photomultiplier (Hamamatsu H7422-40-LV, Japan) after bandpass filtering by a 525/50 nm emission filter. Images (approximately 150 x 1024 pixels) were acquired with ImSpectorPro software (La Vision Biotech, Germany), typically 20-25 frames/s. Besides, when imaging smaller areas (*e.g.,* 32 x 512 pixels), the used sampling rates were considerably higher (∼50-200 frames/s). The laser intensity was kept below 240 mW (measured at the back aperture) to avoid heat-induced artifacts.

**Fig. S37.**
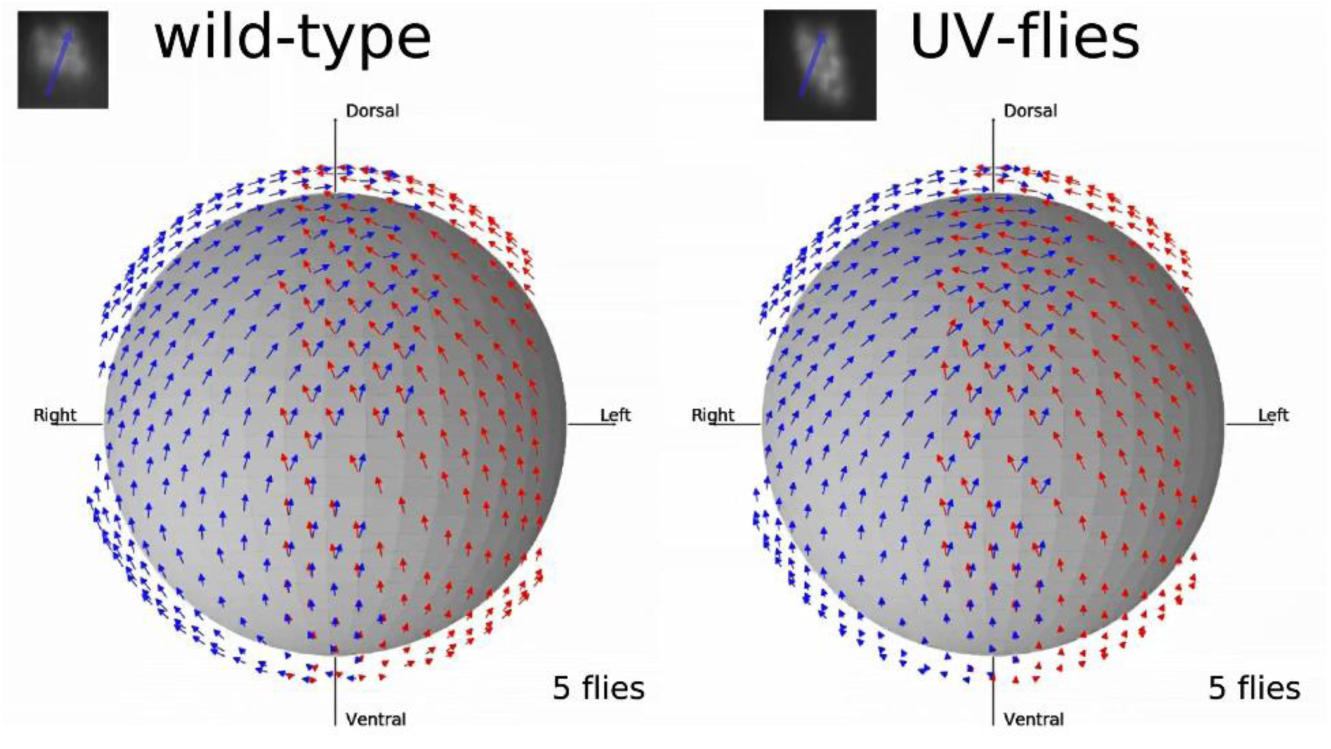
UV-flies show standard photoreceptor microsaccade directions across their eyes. The left (red arrows) and right eye (blue) photoreceptor microsaccade movement trajectories of wild-type (left) and UV-flies (right) flies match. The trajectories were calculated through image cross-correlation from the light-triggered high-speed microsaccade video recordings (see Section II.1). Average directions are shown; data recorded from five tethered flies.

##### IV.1 *In vivo Drosophila* preparation

2-to-4-day-old cold-anesthetized flies (usually males) were prepared for the experiments much as described before (22, 89, 90). A fly was waxed to a 0.001-inch-thick folded stainless steel shim holder, which allowed access to the back of the head through a 0.8 mm opening (Fig. S38*A*). The head was tilted forward approximately 60°, exposing its back at the opening, and left the retina below the shim (Fig. S38*B*). We cut a small hole at the back of the head cuticle with a fine tungsten needle and removed connective tissue, including the trachea, to obtain optical access to the left and/or right medulla L2 axon terminals (Fig. S38*A*). The fly was positioned over an air-suspended 6.13 mm Æ polypropylene ball within the 2-photon imaging system, facing panoramic visual stimulation screens to enable motor activity recording (Fig. S38*C*). Closed-loop temperature-controlled (25 °C) oxygenated fly ringer solution (containing in mM: 120 NaCl, 5 KCl, 10 TES, 1.5 CaCl_2_, 4 MgCl_2_, and 30 sucrose) was perfused over the back of the head, keeping the preparation alive/healthy for hours-long experiments

**Fig. S38.**
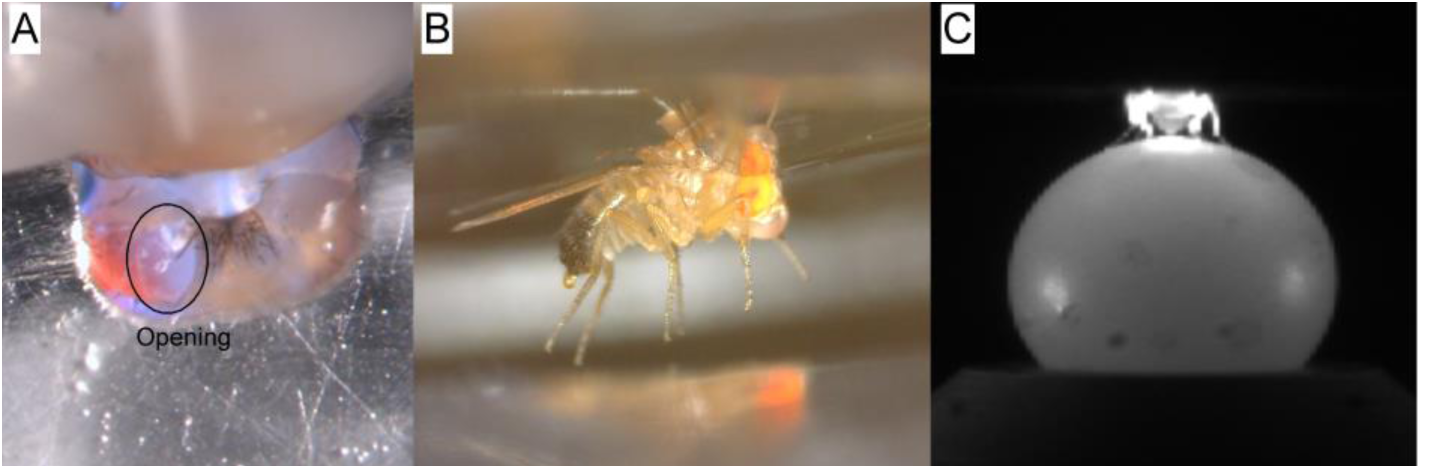
*in vivo Drosophila* preparation for 2-photon Ca^2+^- imaging. (*A*) Using a bespoke preparation micromanipulation system, we fixed a fly in the predetermined position and orientation to a 0.001-inch-thick folded stainless steel shim (of a disk-shaped fly-holder) to access the back of the head through a 0.8 mm gap, comparable to (89). A small opening was cut at the head’s back cuticle through an oxygenated fly ringer bath that covered the back of the head only, giving a visual view of the left or right medulla L2-terminals. (*B*) The fly’s positioning inside the portable disk-shaped fly-holder. The fly-holder was transported to the 2-photon imaging system, where it was rotationally adjusted by hand to center the fly facing the panoramic visual stimulation screen (Fig. S39, below). Notice the semi-transparent beeswax droplet underneath the fly’s eyes, immobilizing its proboscis. (*C*) A tested fly could walk on a track-ball during the 2-photon imaging of its L2-terminals’ neural responses (Ca^2+^ fluorescence signals). In the experiments, the fly faced the visual stimulation screen inside a black-fabric chamber, which blocked outside light leakage and minimized scatter and internal reflections.

##### IV.2 Visual stimulation

Fig. S39 and Movie S8 show how *fast ultrafine* video stimulation was presented to *Drosophila* during the 2- photon Ca^2+^-imaging experiments. The effective stimulus resolution viewed by *Drosophila* was 7.5-11.25- times finer than its eyes’ average (∼4.5°) interommatidial angle (12), which was long thought as the visual resolution limit. We used a digital light projector (EKB DLP® LightCrafter™ Fiber-E4500MKII™ development module, EKB Technologies, Israel), equipped with a powerful 385 nm UV-LED, to provide 360 Hz UV-video stimulation with native 912 x 1140 pixel resolution (Fig. S39*A*). The UV-video images were projected on a back-projection (diffuser) screen. The whole system was inside a black, fluffy-fabric enclosed cage (Fig. S39*B*) to block outside light and minimize internal reflections and scatter. Three short focal length achromatic doublet camera-lenses (MVL6WA, Thorlabs, USA) were then used to focus the projected images onto one end of three 7 x 7 mm coherent bundles of optical fibers (IB ASSY QA x 24”, Schott, USA), with ∼108 x 108 pixels (as the counted average) projecting onto each bundle (Fig. S39*C*). These images were transmitted and magnified by three optical tapers (Schott, USA). The tapers formed three Parafilm-capped panoramic fiber-optic screens (virtual reality stimulation screens), surrounding a tested fly frontally (Fig. S39*D*). Parafilm diffused light and damped reflections related to the numerical aperture of the taper/bundle fibers. The three fiber-optic screens accurately reproduced the video images into three angled vertical sections, positioned 38 mm from the fly eyes, filling large central parts of their left and right visual fields (total area: 135° x 45°). Therefore, with 108 x 324 pixels spread across the three screens, the angular resolution was ∼0.6° at the point closest to the eyes and ∼0.4° near the corners.

**Fig. S39.**
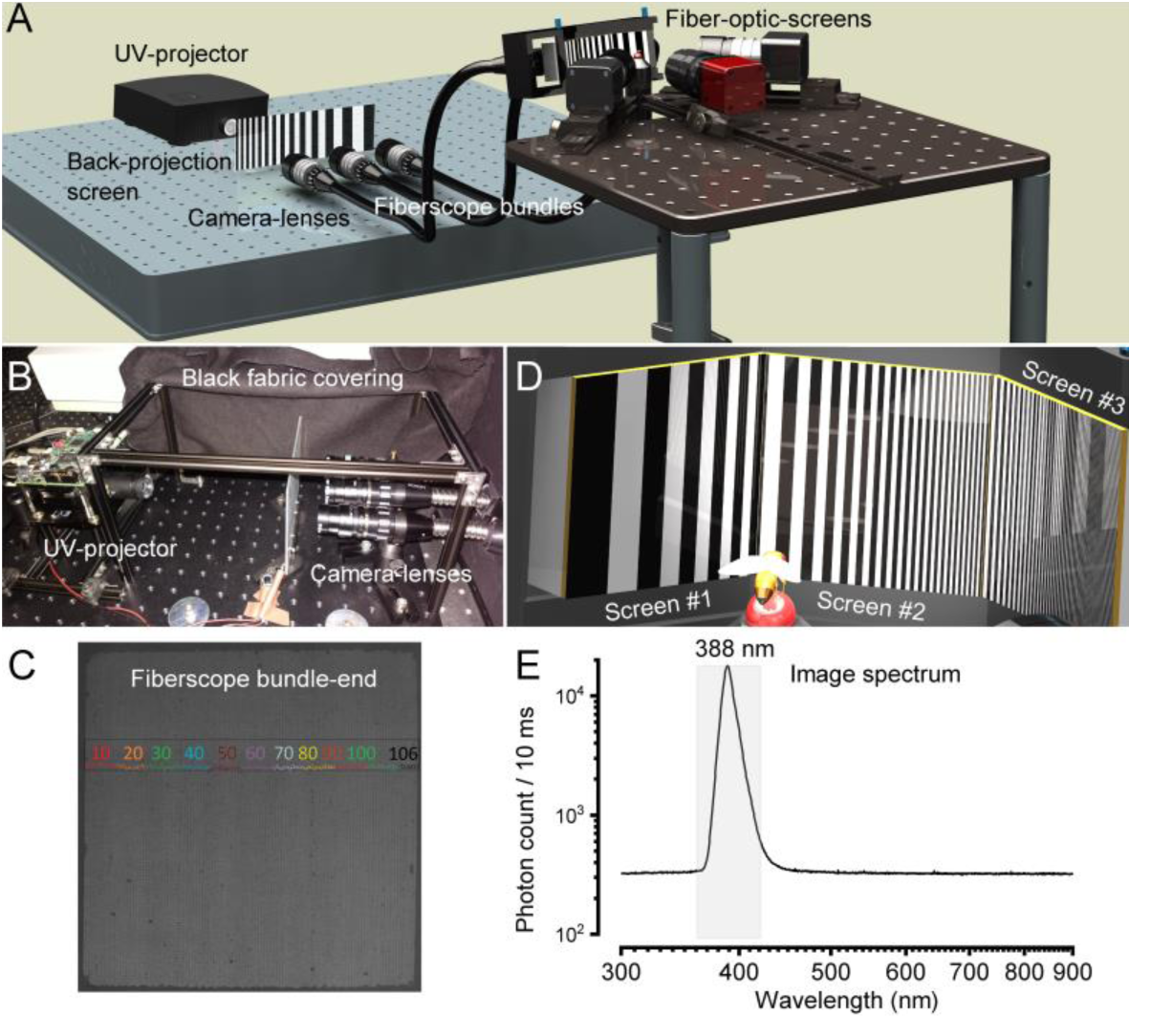
The bespoke high-resolution UV-video-display system (attached to the 2-photon imaging system) used for stimulating L2 neurons visually. (*A*) The optical path, from the high-speed UV-projector to three high-resolution fiber-optic-screens (taper ends), for presenting *Drosophila* with UV-video stimuli. (*B*) UV-stimuli were projected on a UV- preserving back-projection screen. Three camera lenses sampled the focused back-projected video images on three high-resolution ordered fiberscope bundles. This optical path was kept inside a light-proof cage (covered by a thick, fluffy black fabric) to minimize light scatter and internal reflections. (*C*) One fiberscope bundle end, with the highlighted fiber count for one of its rows. (*D*) The panoramic visual stimulation screen assembly was made out of three high-resolution optical tapers (fiber-optic-screens), in which angles and position could be precisely and freely adjusted and fixed around the tested fly (by the instrument design). (*E*) The video-display system’s spectral output, as directly measured at the visual stimulation screen facing the tested *Drosophila*. The visual stimulation was dominated by UV-light, peaking at 388 nm.

Visual stimuli were created using custom-written Matlab code, partly using the Psychophysics toolbox, in which the renderer updated images at 360 Hz, with a nominal 8-bits of DLP intensity at each pixel, and accurately projected them onto the three taper-screens. Additional UV-band-pass filters (Edmund Optics, UK; 377 nm, bandwidth 50 nm, OD 6) and adjustable apertures, interposed between the back-projection screen and the bundles, allowed us to cut off long (non-UV) tail wavelengths of the images and adjust their overall intensities. The spectrum used in experiments is shown in Fig. S39*E*. We estimate that R1-R6 photoreceptors that faced the optic taper screens were presented with 10^5^-10^6^ UV-photons/s, causing moderate to high light adaptation. Notice that because of the refractory photon sampling and intracellular pupil, which cause a dramatic drop in quantum efficiency (11, 40, 59), most photons are lost during light adaptation. Consequently, an R1-R6 photoreceptor’s effective photon absorption rate is actively maintained at ∼1.5-8.0 x 10^5^ to maximize its information transfer rate for high-contrast stimuli (11).

##### IV.3 Measuring L2-terminal sensitivity to stimulus velocity and orientation

The images about medulla L2-terminal fluorescence responses were analyzed by custom-written Python scripts (K. Razban Haghighi). The fluorescence intensity variations were quantified after background subtraction. Ca^2+^-signal variations were obtained by subtracting the basal fluorescence, *F*_0_, calculated as the mean intensity before the visual stimulation, from the observed intensity, *F*, (Δ*F* = *F* − *F*_0_) and giving this difference as the relative fluorescence change (Δ*F*/*F*_0_).

***The use of “UV-flies” minimizes antidromic sampling artifacts*** Because the basement membrane between the lamina and retina lacks screening pigments, photoreceptors can be stimulated antidromically by shining light through the fly brain (91). Equally, during Ca^2+^-imaging, fluorescence signals from the brain circuits propagate towards the photoreceptors. Therefore, in *Drosophila* with wild-type spectral sensitivities, the green-light-activated R1-R6s and R8_yellow_ photoreceptors inadvertently multiplex light stimuli from the world with the L2 green-fluorescence signals from the lamina, potentially obfuscating downstream visual processing (as recorded by two-photon imaging). We used “UV-flies” (22) to overcome this problem.

**Fig. S40.**
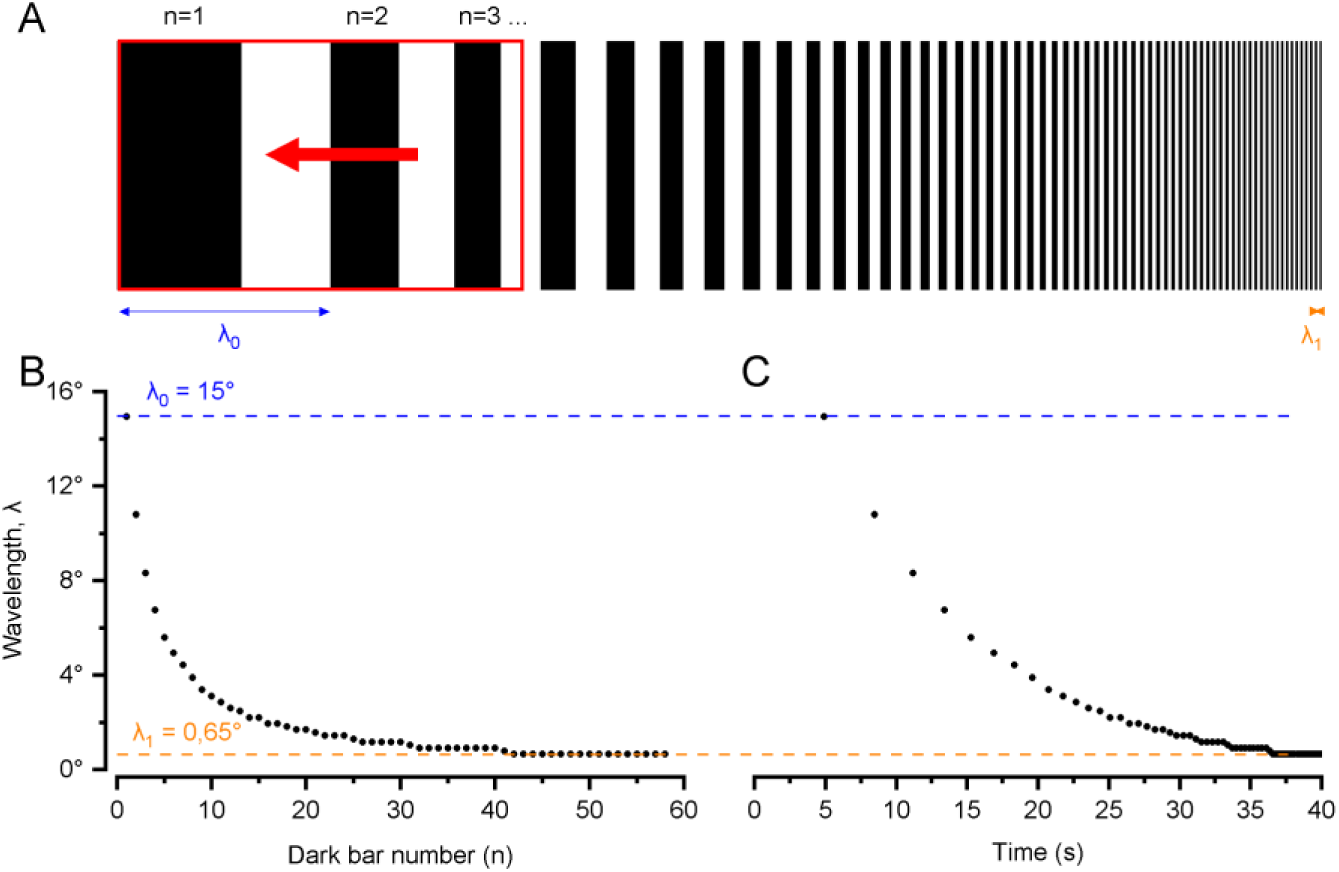
Graphical description of the four parameters used for dynamically narrowing the black-and-white bar grating stimulation in time. (*A*) Grating stimulus design. Wavelength narrows in time from *λ*_0_to *λ*_1_. Red rectangle: the stimulus screen as seen by the fly. Red arrow: Grating motion direction, running through the screen at a constant speed. (*B*) The wavelength at each black bar. (*C*) Wavelength over time. The orange dashed line indicates the smallest tested wavelength, 0.65° (*B* and *C*).

***Testing individual L2-terminals’ speed and orientation sensitivity to moving stripes and bars*** L2 neurons’ medulla terminals respond strongly to light-OFF stimuli (28, 29, 34). Therefore, a bright moving bar crossing an L2 neuron’s receptive field (RF) evokes a transient response. Here, we used two types of moving stimuli to measure L2 speed and orientation sensitivity.

One stimulus type was made of two parallel bars crossing an L2 neuron’s RF. These bars induced a two-peaked change in the observed L2-terminal calcium fluorescence as a response. We can measure how well this intraneural calcium response resolved the two moving stimuli using the Rayleigh criterion:

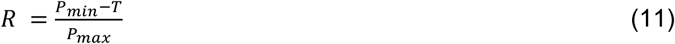

, where *T*, *P*_*min*_and *P*_*max*_are the trough, the smallest peak, and the highest peak, respectively.

We further measured single L2 neurons’ resolvability to dynamically narrowing bar gratings (of continuously decreasing wavelength; Fig. S40 *A* and *B*) using a novel four-parameter bar-grating stimulus (as constructed in Matlab). The stimulus parameters were the speed, motion direction, initial wavelength, and final wavelength (*s*, *θ*, *λ*_0_ and *λ*_1_, respectively). The inter-bar wavelength, which entered the tested *Drosophila*’s field of view, followed the geometric sequence update:

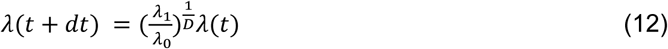

, where *D* was the duration of the stimulus (Fig. S40*C*). This way, the wavelength was divided by a constant factor, frame after frame, enabling an accurate estimate of the wavelength/time point when the L2 neuron could no longer resolve the adjacent moving bars. A more intuitive formula representing the wavelength over time is the following:

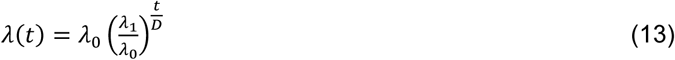

Importantly, this spatiotemporal stimulation enabled us to simultaneously monitor how the neighboring L2- terminals, in which RFs were covered by the same visual display (see above), encoded the same directional motion stimulation in different angular resolutions.

Similar to the moving two-bar stimulation (above), the dynamically narrowing bar grating stimulation induced a Ca^2+^-fluorescence signal, showing a succession of peaks. To each pair of peaks, we can attribute resolvability. Since this stimulus induces a response with a dynamic baseline, we applied the Rayleigh criterion on the relative peak heights:

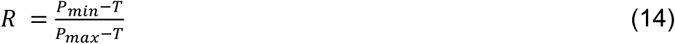

To make resolvability estimation consistent and free of human observer bias, we built a six hyper-parameter algorithm in Python that takes the Ca^2+^-fluorescence signal as input and returns the smallest resolvable angle (SRA). Two of the parameters enable accurate peak detection, considering the noise in the data. One parameter is the noise-threshold: *R* = 0, if *P*_*min*_− *T* is smaller than the threshold. The other parameter is the inter-peak noise threshold: *R* = 0, if the inter-peak noise is higher than the threshold. Two separate parameters were used to detect false negatives. The last pair of peaks where *R* ≠ 0 is taken as the SRA. In separate tests, the algorithm generated highly similar resolvability estimates to those provided by trained experimentalists.

**Fig. S41.**
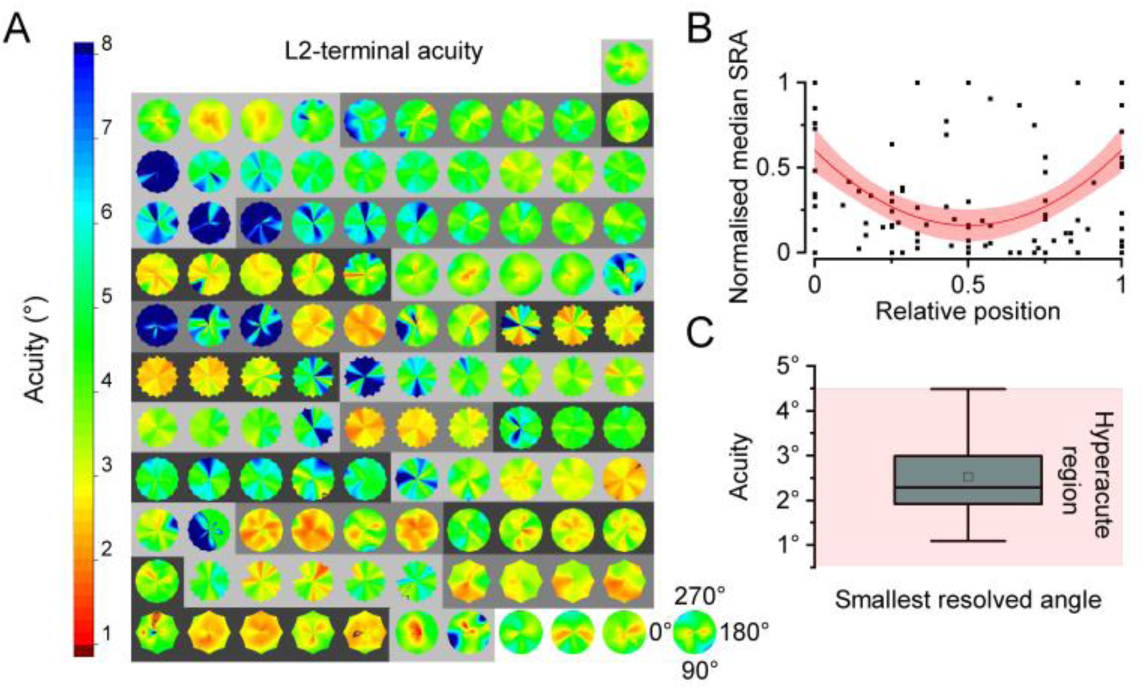
L2-terminal Ca^2+^-fluorescence responses show hyperacute speed and motion-direction sensitivity to moving bar-grating stimulation. (*A*) The figure presents the collective motion direction sensitivity maps (*i.e.,* acuity maps) of 20 flies, tested with the four-parameter bar-grating stimulation protocol. Each fly displayed at least one and at most twelve actively responding neighboring L2 medulla terminals. Each fly’s maps are shown in their physical order, following their terminals’ medulla positions, plotted on the same gray background. The L2-terminals closest to the edge of the imaging window, bordering the dissected tissue area, typically showed the least sensitive responses (from cyan to blue). Whilst the L2-terminals in the center showed the highest sensitivity and hyperacute stimulus resolvability (from light-green to dark-red), with many neurons encoding less than 1.5° apart bars moving along a specific direction(s). (*B*) Each point represents the median value of a single L2-terminal’s *smallest resolved angle* (SRA) map vs. its relative position in the recording window. The values are normalized within each fly. The red line shows a quadratic fitting of the points, with 95% confidence and prediction intervals (red and pink areas, respectively). Two possible reasons may account for a better acuity in the center than on the edges: (i) the peripheral terminals might be closer to the dissected tissue, implying health issues; (ii) the recording plane and the neural plane are not parallel, and local fluorescence signal-to-noise ratio (SNR) along the axon is not constant. These reasons suggest that only the central axons and terminal can be sectioned at the highest SNR region. (*C*) Distribution of all the L2-terminal’ minimum SRA (highest acuity). Mean (dot), median (line) are shown. Box range: 25-75%. The pink background indicates the hyperacute stimulus motion resolvability range (<4.5°).

For each recording, we could monitor several (between 1 and 12) L2-terminal responses simultaneously (Fig. S41 and S42). The stimuli were presented multiple times to the fly by varying the speed (usually *s* = 20, 30, 60°/s) and the motion direction (usually every 15° or 30°, covering 360°). Hence, this gave us an SRA polar heat map (acuity map) for each recorded neuron in the fly preparation (Fig. S41*A* and S42*A*). These SRA polar heat maps almost always suggested the best-resolved direction (the direction of highest acuity; or the stimulus direction for which SRA is smallest). To calculate it accurately and quantify the accuracy, we fitted the SRA (modulo 180°) using a 180° fixed-wavelength sine-function with Levenberg-Marquardt iteration algorithm (Fig. S43 and S44). The reason for this choice is that we expect periodic SRA values with minima at an angle *α* and *α* + 180°, and a maximum at *α* + 90° and *α* − 90°. The phase (subtracted by 45°) of the fitting gives us the “preferred” highest-acuity direction. We used the Levenberg-Marquardt error values as error margins (Fig. S43*B*). We also evaluated these fits with the R^2^ value. Given that Gaussian noise sinusoidal fitting has an R^2^ distribution with mean = 5.8% and rarely reaches 15%, we considered that a clear preferred direction for L2 SRA fitting was when *R* > 25% (∼*Err* < 12°) (Fig. S42, *B* to *D* and S44).

We calculated each recorded L2 neuron’s receptive field (RF) location using two stimuli: a single light bar moving back and forth horizontally and another vertically. We considered each terminal’s peak responses induced by the bar leaving its receptive field (characteristic of an OFF response). This correspondence enabled us to reconstruct a good approximation of the RF boundaries.

Therefore, for each tested fly, we attained a map of its L2-terminals’ highest-acuity directions positioned at the corresponding receptive field locations (Fig. S42*C* and S44).

We used data from the best-dissected (or healthiest) fly preparations in the main results (Fig. 4), which displayed at least eight consecutive neurons with consistent activity (Fig. S42). We found that:

- The L2-terminals’ most preferred motion directions (i.e., the orientation axes of their motion-direction sensitivity) are collinear to the connected photoreceptors’ microsaccadic motion directions (Fig. S42*C*). This assessment excluded the most peripherally recorded terminals. These outliers typically showed inconsistent responses. Such inconsistency could be caused by compromised health at the dissected tissue boundary (Fig. S41*B* and S42, *B* to *D*). Or, it could reflect variable SNR along the axons, where the highest SNR cannot be recorded on every axon because the recorded section plane, and the actual neural plane, were not parallel.
- The preferred motion directions shifted systematically about 5° from neighbor to neighbor (Fig. S42, *B* to *D*), similar to the gradual shifting of the photoreceptor motion directions (Fig. S37).

**Fig. S42.**
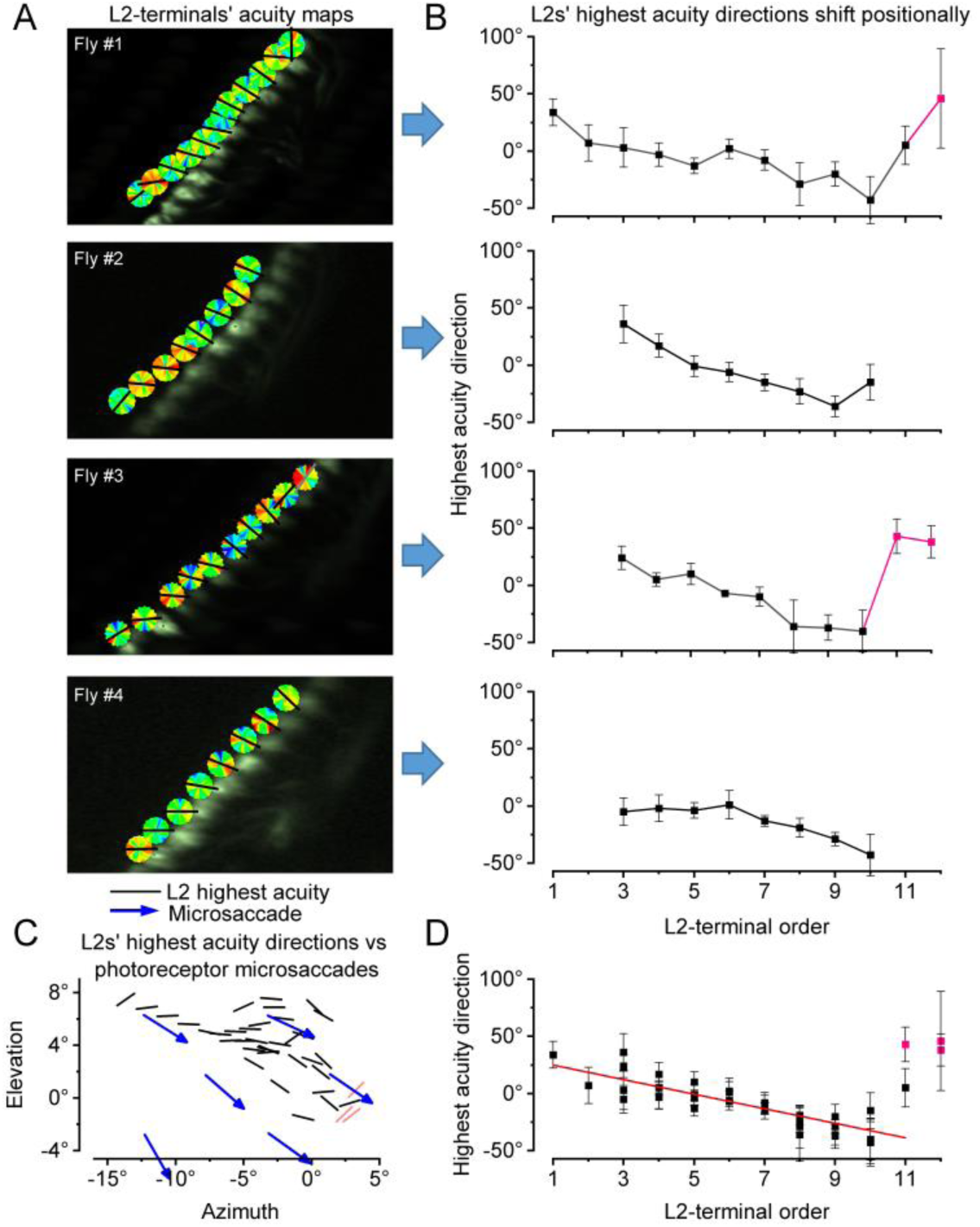
L2-terminals’ motion-direction sensitivity axis, showing the highest acuity, aligns with their photoreceptors’ microsaccade directions in healthy fly preparations. (*A*) Two-photon GCaMP6f- fluorescence images of L2 axon terminals from four *Drosophila* (#1-4) that provided long-lasting stable recording conditions. Next to each terminal is its corresponding acuity map, with a black line indicating its “preferred” highest-acuity motion direction (fitted orientation axis). (*B*) L2-terminals’ preferred motion directions shift gradually and systematically across their retinotopically organized medulla layer. Only a few peripheral terminals (red), closest to the surgically prepared recording window’s edge, showed inconsistent, possibly dissection-affected responses. The error bars give the Levenberg-Marquardt error range for each fitted highest-acuity direction; see Fig. S43 and S44, below). (*C*) The locations of the L2 receptive fields (RF; shown for the fly’s right eye) with their respective highest-acuity directions (black lines shows their fitted orientation axis) aligned broadly with the corresponding photoreceptor microsaccades’ biphasic motion directions (blue arrows; *cf.* Fig. S37.). (*D*) L2-terminals’ highest-acuity motion directions aligned regarding their RF locations.

**Fig. S43.**
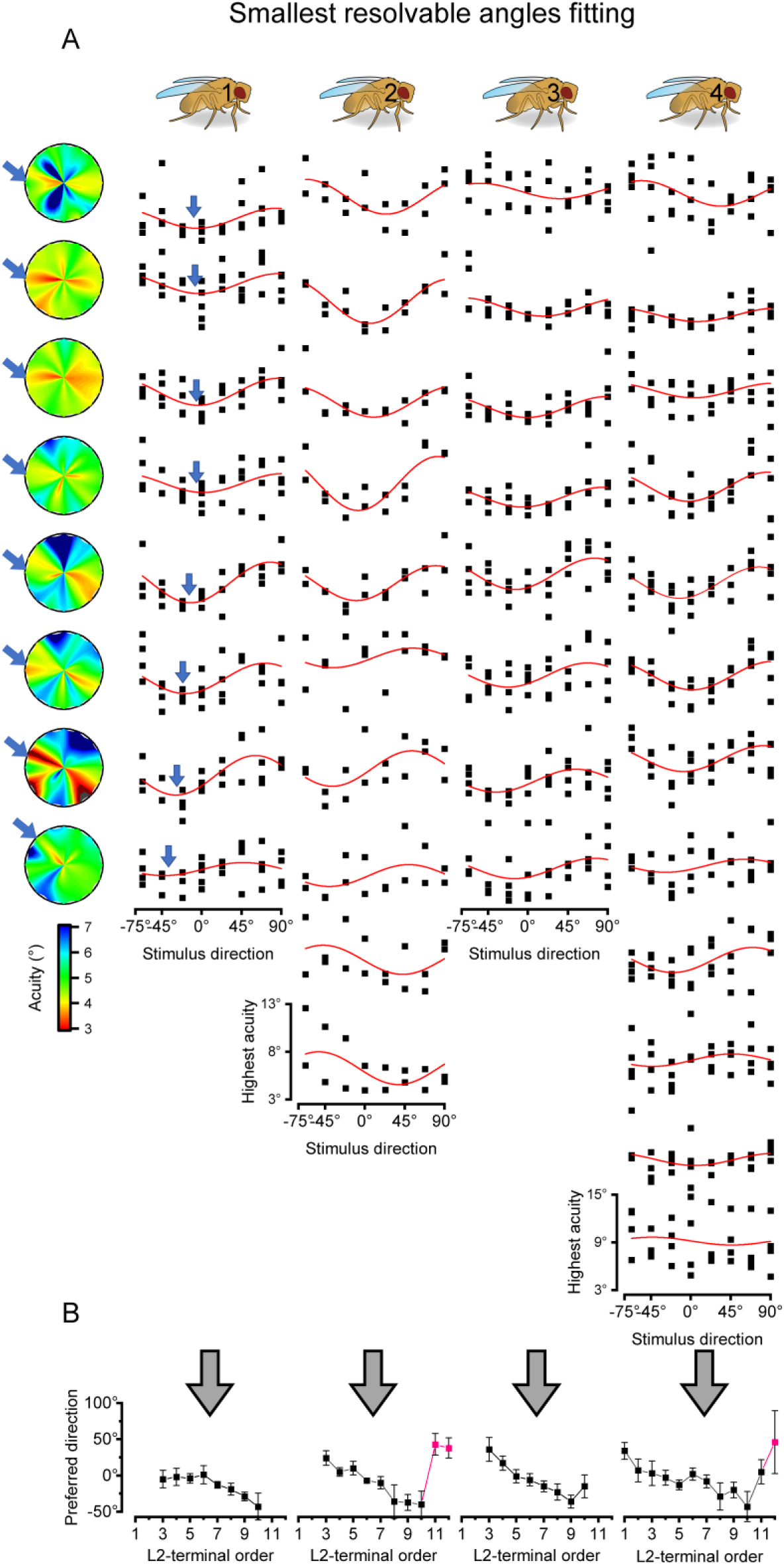
Determining L2- terminal’s highest-acuity direction by sinusoidal fitting (examples from four healthy flies). (*A*) The obtained SRA plotted against the stimulus direction for each recorded L2-terminal. Red curves indicate 180°- wavelength sinusoidal fitting, shown for each L2-terminal (rows) of the four flies (columns), one column per fly. Blue arrows are the “preferred” highest-acuity directions; chosen as the sinusoidal fits’ minima. Heat maps (SRA acuity maps) are shown for the #1 fly’s every L2-terminal. (*B*) The minima of each fit for the consecutive (neighboring) L2- terminal, plotted per fly. Error bars give the Levenberg-Marquardt error margin for each fit.

**Fig. S44.**
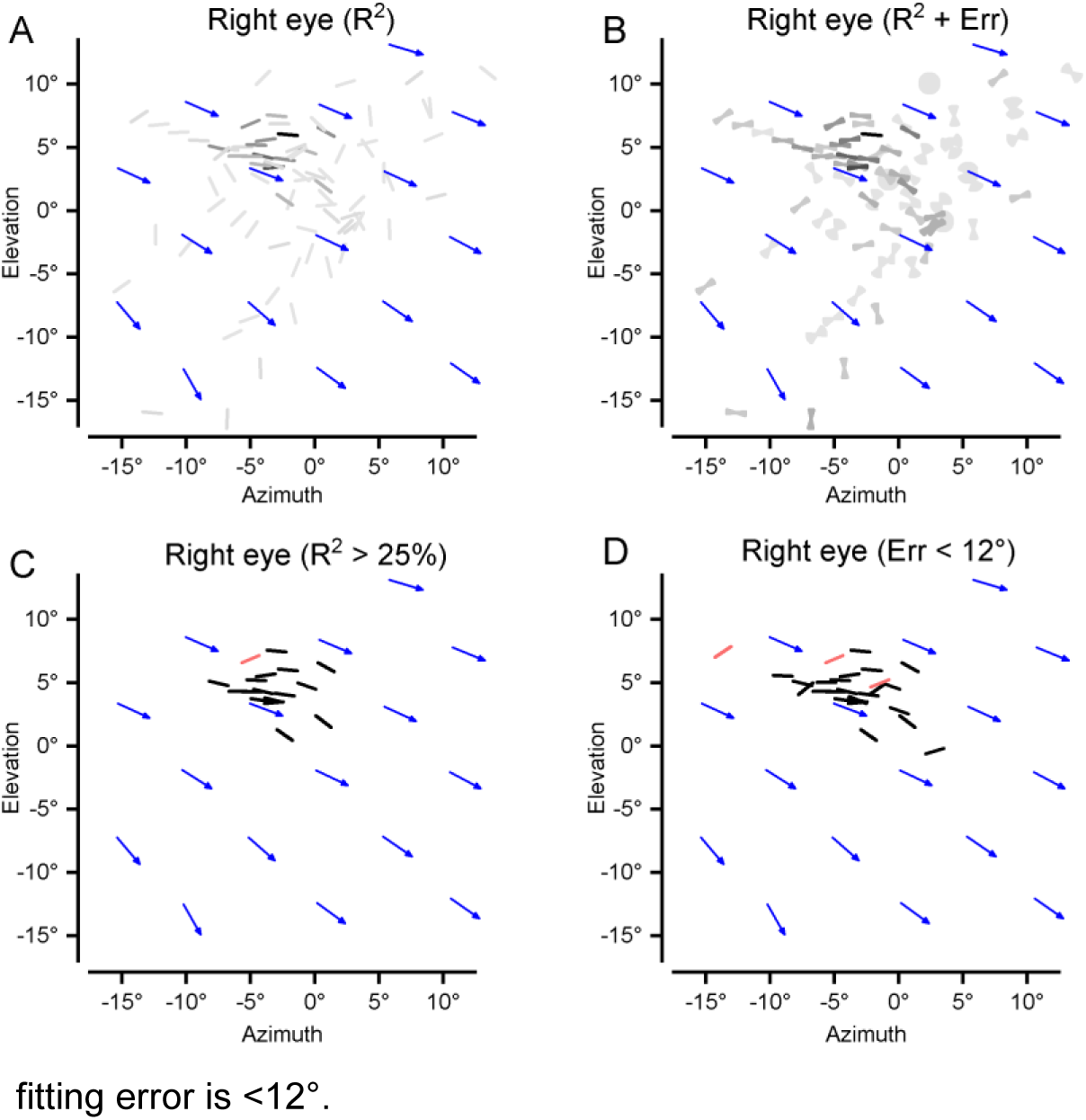
All the measured right eye L2-terminals’ highest acuity directions, plotted regarding their receptive field (RF) locations. (*A*) R^2^ of the sinusoidal fits are shown in a linear grayscale: the lightest 0%; the darkest 100%. The darker the line, the better the reliability and predictive value of the estimated highest-acuity direction. Red: peripherally recorded L2- terminals (possible outliers). Blue arrows: the photoreceptor micro-saccade directions at their corresponding RF locations. (*B*) R^2^ of the sinusoidal fits shown with the same reliability-dependent coloring as in *A*. Error of the sinusoidal fits shown with their direction margins. (*C*) L2-terminals’ highest acuity directions when the sinusoidal fits’ R^2^ is >25%. (*D*) L2-terminals’ highest acuity directions when the sinusoidal fitting error is <12°.

***Sampling aliasing prevention*** To concurrently image many L2-terminals with a high signal-to-noise ratio, we used relatively low frame-rates of 20-25 fps (*i.e.,* each complete image frame was sampled at ∼20 Hz). Such rates could be prone to sampling aliasing; if the actual light-stimulus-induced fluorescence changes happened faster than the sampling. However, several factors ensured that aliasing effects on the data were minimal:

- Each image frame is not an instant snapshot but built up by scanning its pixels line-by-line at ultra-high speed (each pixel in ∼50 ns). Thus, both the used resonant scanner’s line-scan rate and the recorded local Ca^2+^-signals’ (pixel-wise) spatiotemporal correlations are much faster than the full image frame rate and the underlying Ca^2+^-fluorescence dynamics.
- The Shannon-Nyquist sampling theorem states that no information is lost if the sampling rate is higher than twice the signal’s maximum frequency. Hence the minimum consistent value for SRA (smallest resolvable angle) follows the rule:

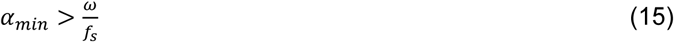

where *α*_*min*_, ω and *f*_s_ are the minimum inter-bar distance used for the SRA, stimulus motion speed, and sampling rate. Those minimum values for the SRA were rarely reached, so the risk of aliasing was minimal.
- The sampling rate was never kept constant in the recordings, thus minimizing any systematic aliasing effects. Theoretically, aliasing causes central symmetrically spreading patterns in the recorded images, such as fake rigs or harmonic ringing (11), which never occurred in the SRA maps.
- Control experiments with much higher frame rates (85-145 fps) generated even higher L2-terminal acuity maps than those with 20 fps sampling, but with similar directional selectivity trends, showing clear hyperacuity and specific highest acuity motion directions. The acuity map trends for the 20 fps and >85 fps sampling started to differ only at the highest tested velocity stimuli (60°/s). One acuity map for 85 fps sampling was included in Fig. S41. Overall, we found a suggestively higher L2-terminal hyperacuity for the higher sampling rate data (Fig. S45):

○ *High fps*: 2.20° ± 0.25° (mean ± SD); SRA = 1.93°, Median = 2.17°, Max = 2.5° (n = 6 L2-terminals)
○ *Low fps*: 2.53° ± 0.82° (mean ± SD); SRA = 1.09°, Median = 2.31°, Max=6° (n = 117 L2-terminals)

Therefore, in light of all this evidence, together with *Drosophila*’s striking hyperacute visual behaviors in a flight simulator system (11, 78) (Section V, below) and faster intracellular voltage responses (22, 33–36, 77–79), we are confident that we present reliable and conservative estimates (lower bounds) of the L2- terminals’ motion direction-sensitive hyperacuity (for the given experimental conditions, instrumental noise, and sampling limitations). A freely flying *Drosophila*’s visual acuity can only be better in natural environments and could even be significantly higher.

**Fig. S45.**
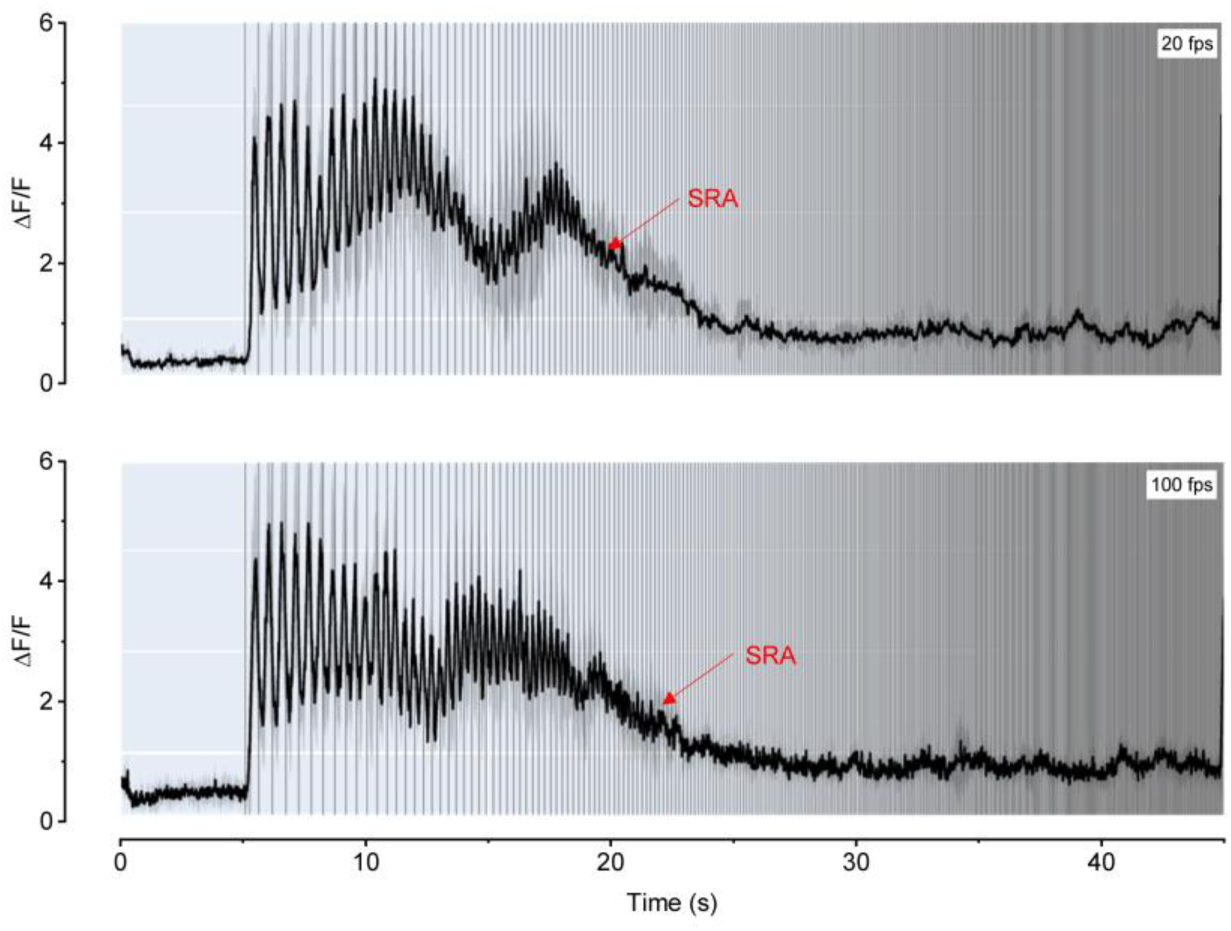
Using a higher imaging frame rate (*i.e.,* Ca^2+^- signal sampling rate) increases the recorded L2-terminals’ resolvability for fast-moving stimuli. (*A*) Average of 4 repeats of the same L2-terminal (ROI) responses at 20 fps to a 60°/s narrowing (13° to 0.65°) grating stimulus moving upwards following 5 s of a gray frame (as in Fig. S40) (*B*) Average of 4 repeats of the same L2-terminal (ROI) responses at 100 fps to the same stimulus as in *A*. Gray margins: ± SD. Red arrows: the first pair of peaks with null resolvability, edging the smallest resolved angle (SRA). Gray vertical bars: times for which a dark bar of the stimulus crosses the L2’s receptive field.

***Removing motion artifacts in L2-terminals’ Ca^2+^-signals*** Both photomechanical photoreceptor microsaccades and spontaneous intraocular muscle contractions can move the fly brain during 2-photon imaging. We used a computer vision and machine learning library (open-cv) to write a stabilization algorithm in Python. Two main functions were needed: one (*goodFeaturesToTrack*) finds the most prominent corners in the image or the specified image region, as described in a proposed algorithm that uses Newton-Raphson style search methods (92). The other (*calcOpticalFlowPyrLK*) calculates an optical flow for a sparse feature set using the iterative Lucas-Kanade method with pyramids.

We used this technique on recordings where the motion artifacts moved the L2-terminal away from the region of interest (ROI) window (typically ∼2µm). This technique enabled the ROI fluorescence average to be coherently correlated with the neural activity and not affected by physical displacement. Fig. S46 shows the resulting displacements for some cases. Interestingly, the displacements were sometimes stimulus-locked: Fig. S46*B* shows slower displacement at the beginning but faster around the end of the stimulus. A high sampling rate (∼85 Hz) shows a robust synchronization between the displacement and the stimulus (Fig. S46*A*). Two phenomena could explain this:

- The stimulus-induced fluorescence variations themselves may fool the stabilization algorithm by faking a motion. However, this phenomenon is unlikely because applying the stabilization algorithm on the stabilized video only resulted in small and noisy motion residuals.
- The fast stimulus-locked L2-terminal displacements are likely induced by the photoreceptor microsaccades as these movements are analogous to the photomechanical tissue displacement recorded during the X-ray imaging experiments. Indeed, as seen in Fig. S37, photoreceptors move photomechanically back-and-forth along the main axis each time a bar crosses their receptive fields, and such motion could similarly drive L2-terminal displacement in Fig. S46*A*. The collective evidence from separate experiments using different assays is already compelling. But for conclusive proof, an additional displacement analysis on activity-independent fluorescence (such as Tomato dye) can be done in the future. Note that the small L2-terminal displacements, such as the one seen in Fig. S46*A*, had no real effect on the recorded fluorescence signals, so subtracting them made no difference in the analyses.

**Fig. S46.**
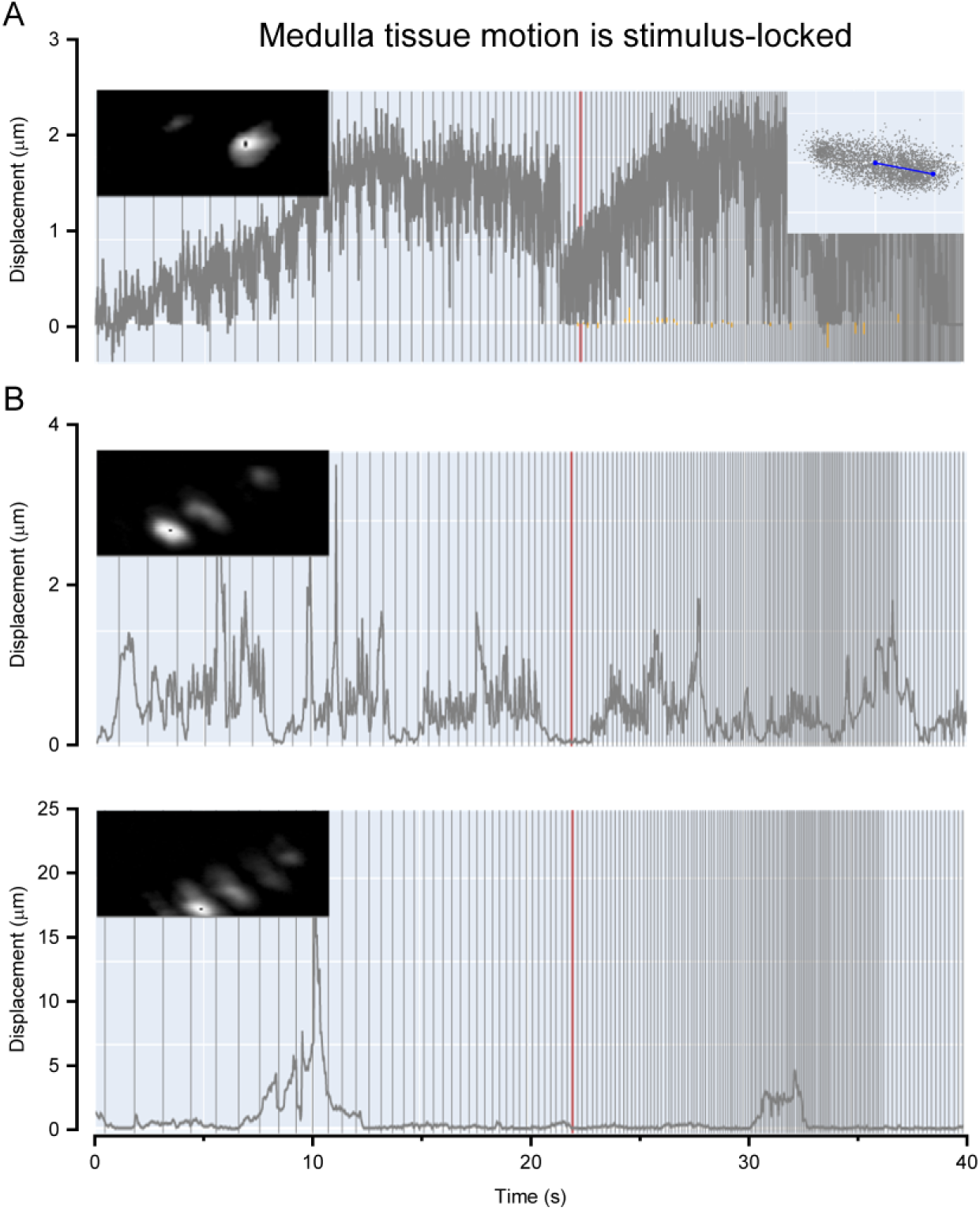
During 2-photon imaging, L2-terminals can show mechanical stimulus-synchronized jitter. We used a stabilization algorithm to subtract this jitter from the fluorescence video recordings if it was deemed too large. (*A*) A L2-terminal (ROI; region of interest) displacement during 85 fps imaging. In the inset, the L2- terminal’s position is projected in the principle direction (blue line). Given the regularity and size of these small movements (<<1 µm), they likely resulted from the photoreceptor microsaccades bouncing the optic lobes in a stimulus-synchronized manner. Similar optic-lobe-displacement dynamics were seen during the X-ray imaging (see, *e.g.,* Fig. S3) (*B*) Two examples of larger mechanical displacements of the medulla L2-neuron terminals, obtained with low (∼20 Hz) sampling rates (20 fps). The larger movements (>1 μm) are likely caused by intraocular muscle activity (13) that can move the retina in slow bursts. The smaller movements (<<1 μm), superimposed on the bursts, are likely caused by the stimulus-synchronized photoreceptor microsaccades moving the retinal tissue. The three images depict the studied ROI pixels’ standard deviation; *i.e.,* showing how the L2-terminal physically moved during the dynamically narrowing bar grating stimulation (Fig. S40). The red vertical lines in *A* and *B* indicates GCamp6f resolvability limit, as obtained from separate flash-stimulation tests.

Furthermore, the larger (>1 µm) and more sporadic L2-terminal movements in the medulla, as seen in the analyses (Fig. S46*B*), likely reflected intrinsic intra-ocular muscle activity (13).

The scripts to process and analyze the 2-photon images are downloadable from the repository: https://github.com/JuusolaLab/Hyperacute_Stereopsis_paper/tree/main/AnalyzeL2Data

#### V Multiscale modeling the adaptive optics and photoreceptor signaling

**Overview**

This section describes the theoretical multiscale approaches to simulate the *Drosophila* ommatidium/compound eye optics and biophysically model how its R1-R7/8 photoreceptor cells sample spatiotemporal light information morphodynamically. It deals with three general cases:

- *Point-source light stimulation simulations*. We calculated the light power a *Drosophila* photoreceptor absorbs from the stimuli using the following two-step optical calculations. The first step consists of applying ray tracing to propagate the incoming light through the lens, followed by applying the Fourier transform beam propagation method (FTBPM, (37)) for propagation through the crystal cone and the rhabdomere. In contrast to the earlier ommatidium wave-optical modeling (75, 93, 94), this approach gives more flexibility to analyze the optical structures’ individual contributions and combined effect on morphodynamic light information sampling when R1-R7/8 photomechanics (10, 11) shift the rhabdomeres axially and sideways (11). Moreover, because each ommatidial R1-R7/8 rhabdomere has its unique size (11, 12) (Fig. S47), the optical simulations were tailored to produce the specific dynamically absorbed light power inputs of their transductions. In the subsequent (biophysically tractable) four-parameter photon sampling model simulations (11, 39, 40), the light inputs were converted to refractory quantum bumps (QBs), which integrated each R1-R7/8’s light-induced current (LIC). The photoreceptor voltage output simulations were then converted from their LICs (11, 39, 40) by using the Hodgkin-Huxley-type photoreceptor membrane model (78, 95) (the HH-model module (11, 39, 40)).
- *Complex stimulus pattern simulations*. R1-R7/8 responses to moving objects were simulated within their receptive fields (RF), estimated from point source simulations with corresponding rhabdomere size and axial/lateral positions. Similar membrane voltage calculations were then conducted, as above, with the full photoreceptor model.
- *Stereo vision.* We simulated frontal stereo-information sampling using stereoscopic photoreceptor arrays in both the left and right eyes and their measured morphodynamic microsaccade dynamics (see Section II.6. and Fig. S23, above). We propose a new theory/method based on neurophysiologically feasible cross-correlation computations to estimate object depth by the subsequent neural networks.

**Fig. S47.**
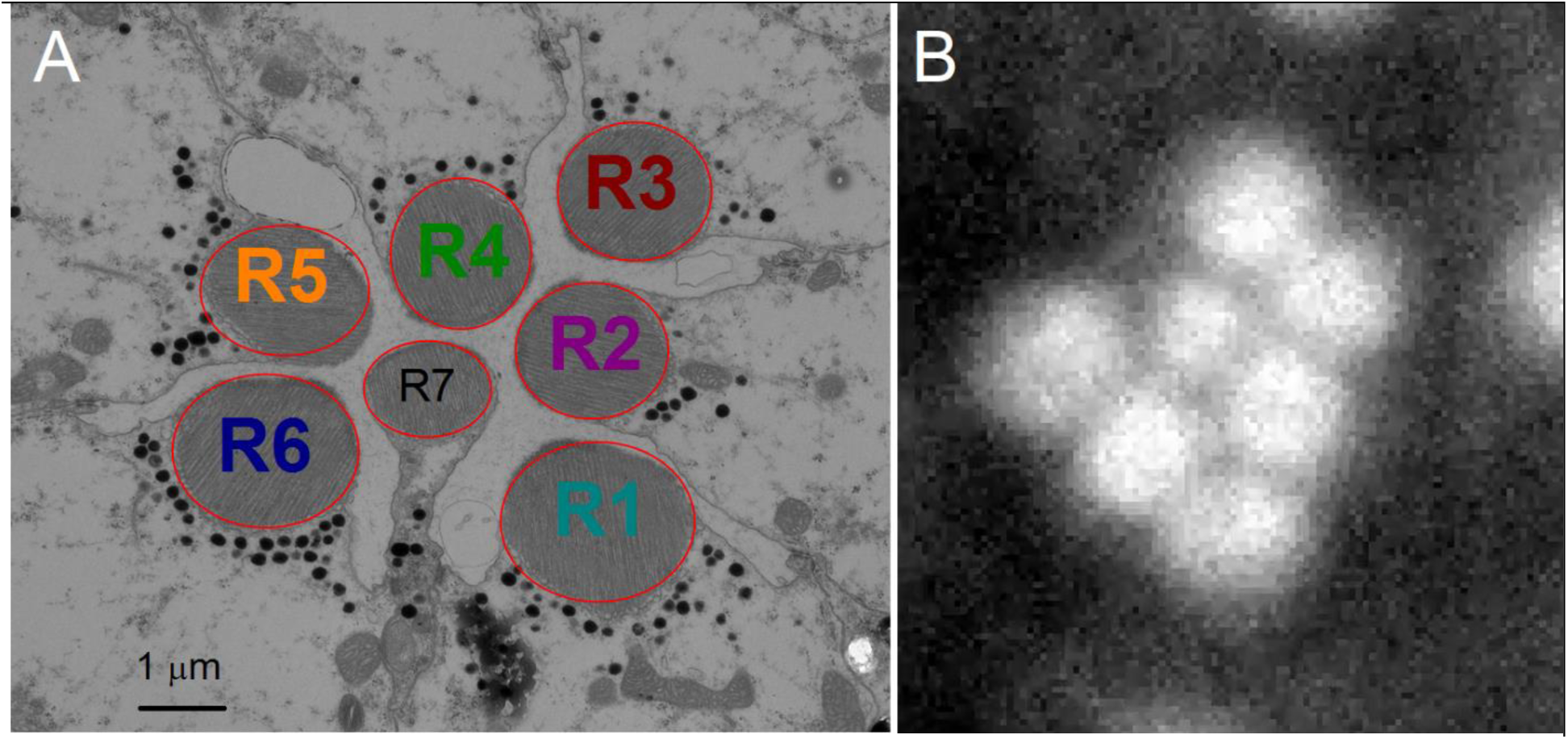
R1-R7/8 rhabdomere shapes vary from oblong to round and have different sizes. (*A*) Typical ommatidial R1-R7/8 rhabdomere pattern as seen in transverse section EM cut. (*B*) Antidromically IR-illuminated ommatidial R1-R7/8 rhabdomeres as recorded during high-speed *in vivo* imaging through cornea neutralized eyes. Images modified and adapted from (11).

##### V.1 Ray tracing through the *Drosophila* ommatidium lens

In this and the following two Sections, we define the *Drosophila* ommatidium optical structures and how they are parameterized for realistic photomechanical R1-R7/8 photoreceptor light sampling simulations, starting with the ommatidium lens.

To analyze the *Drosophila* optics for a light point source stimulus, we used a ray-tracing method (37) to simulate the average 16 μm diameter ommatidium lens (Fig. S48). Rays were cast to a regular square grid (31 x 31 rays) at the thick convex lens’ front (outer) surface plain (16 x 16 μm) from a distant point source, 1 m away. The rays were then traced to the lens’s back (inner) surface by calculating their intersection points with the outer and inner lens surfaces. Only rays hitting the front lens surface were considered. Finally, the intersection points with the outer plain were calculated. The results of the above served as an input to the FTBPM, discussed below.

The main lens parameters were obtained from the previous optical study (75): thickness, 8 μm; outer and inner surface curvatures, 11 μm and -11 μm, respectively; refractive index, 1.45; and the underlying crystal cone refractive index, 1.34.

**Fig. S48.**
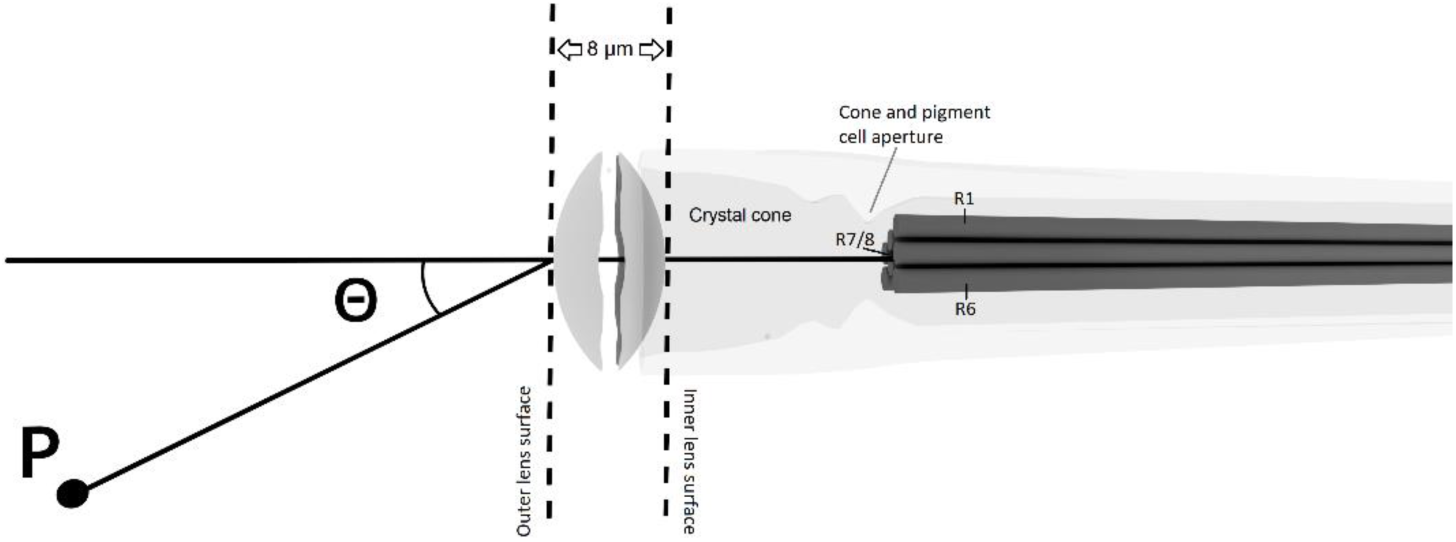
The ommatidium lens system. Ommatidium lens (left) and R1-R7/8 rhabdomeres (dark-gray rods, right). The optical *z*-axis goes through the lens center and R7/8 rhabdomeres. R1-R6 rhabdomeres do not lay at this axis but about 2 µm off-center. The crystal cone (gray) and the cone and pigment cell aperture, which narrows the light pathway (light-gray), are at the front of the rhabdomeres (11). Rays are cast from a point, *P*, with an incident light angle, θ, to the lens’s optical center axis. Rays hit a regular grid at the outer lens surface (the left dashed line). Each ray is then traced to the inner lens plain (the right dashed line).

##### V.2 Beam propagation through the *Drosophila* crystal cone and rhabdomere

Owing to a rhabdomere’s complex lightwave properties, we used FT BPM (37) to simulate the field propagation through the crystal cone and the rhabdomere. The FT BPM is easily applicable and does not need analytical solutions to simulate the behavior of light in a rhabdomere’s complex optical structure. The method is quite suitable to deal with paraxial propagation in structures with low index contrasts.

For monochromatic light, the 3D scalar wave-equation, with an assumed time dependency *e*^*iωt*^, is:

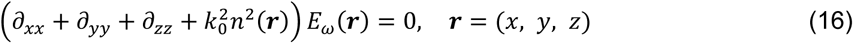

with *ω* the angular frequency, *k*_0_(= *ω*/*c* = 2*π*/*λ*) the vacuum wavenumber, *λ*(= 450nm) the wavelength and *E*_*ω*_(***r***) is the complex electrical field. The true electrical field is:

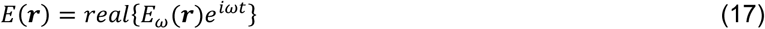

When considering light, which mostly propagates at small angles with, say, the positive *z*-axis (Fig. S49*A*), we can use the slowly varying envelope (SVE) approximation (SVEA), with SVE Ψ:

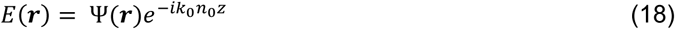

as explained next. The *n*_0_ is a constant defined as 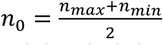. For the rhabdomere, *n_max_*, its refractive index is 1.363, while *n*_*min*_, the refractive index around the rhabdomere is 1.34. Crystal cone was estimated to be homogeneous material (75) with index *n*_0_ = 1.34, which was used in corresponding region. For the SVEA to accurate, the difference between *n*_*max*_and *n*_*min*_needs to be small. As this does not hold for the lens region, the ray-tracing method was used instead of the FT BMP.

**Fig. S49.**
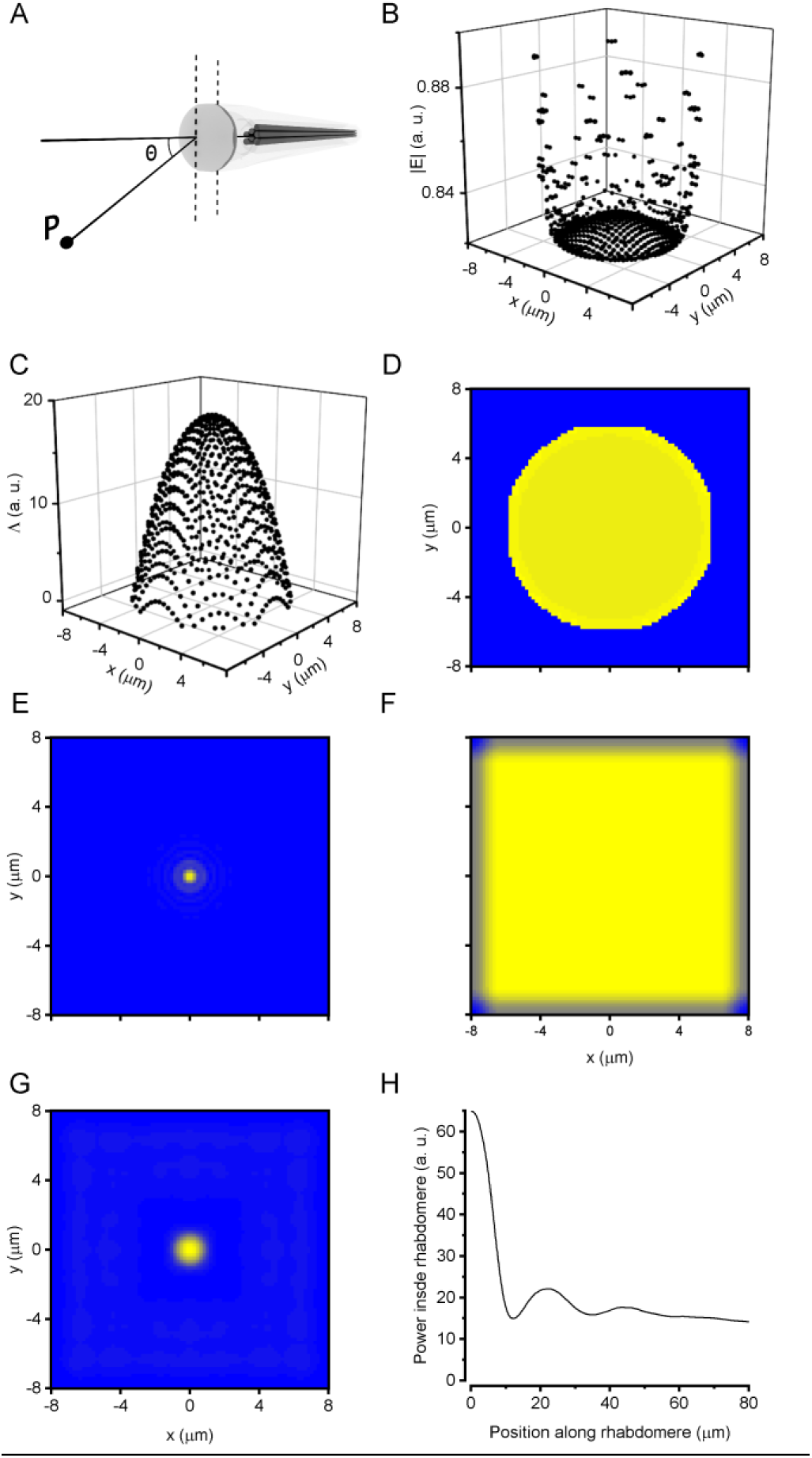
Optical simulation *Drosophila* lens and rhabdomere. (*A*) Rays were cast from a distant point to the ommatidium lens with incident angle 0°. (*B* and *C*) We calculated the rays’ electrical field strength using (Eq. 27) and the rays’ optical distance (Eq. 25) at the inner lens surface. (*D*) These ray tracing results were converted (Eq. 28) and interpolated to the beam propagation electrical field. (*E*) The electrical field size decreased drastically with the beam propagating 17 µm from the inner lens surface to the center (R7) rhabdomere tip. (*F*) The rhabdomere transmittance part (Eq.20) at Δz = 125 nm. The side absorbing boundaries prevent the re-inflow light from leaving the window because of FFT cyclicity. The center rhabdomere has minimal absorption (barely visible here) because of its small 1.0 μm diameter. (*G*) The electrical field strength at the proximal rhabdomere end after traveling 80 μm towards the eye center. (*H*) Absorption light power (inner summation of Eq. 23) along rhabdomere length. The position is 0 μm at the rhabdomere tip and 80 μm at the rhabdomere’s proximal end.

Substituting Eq.16 to Eq.14 leads to an expression:

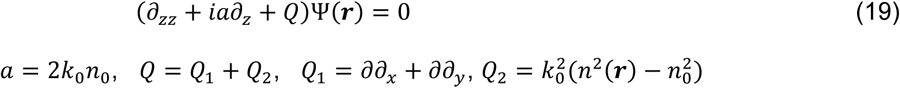

It can be shown that for paraxial propagation in low contrast structures, one may neglect the operator *∂*_*zz*_in the above (37) leading, assuming sufficiently small step sizes *Δz*(> 0), to the following solution to Eq. 19:

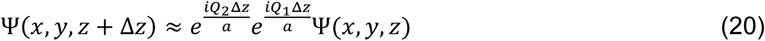

The requirement of small *Δz* values stem from the fact that the exponential operators in Eq. 20, with non-commuting operators *Q*_1_ and *Q*_2_ are applied in succession.

For a practical implementation of Eq. 20, a discretization of the SVE Ψ is required, for which we introduce the matrix ***M***(*z*), containing the field values on a regular grid in the *x-y* plane. The first step - the application of the operator *exp*( *iQ*_1_ *Δz*/*a*) - can now be written as

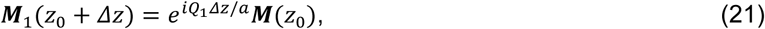

with ***M***_1_ an intermediate result. It can be performed most efficiently in Fourier space, owing to the presence of second-order differential operators, as follows:

1. 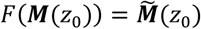, Fourier transform, the elements of 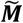 correspond to certain values for the wave vector along *x* and *y*, denoted by *k*_*x*_and *k*_*y*_.
2. Multiply each of the elements of 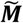 with the appropriate phase factor, *e*^*i*(*kz*−*k*0*n*0)*Δz*^, with 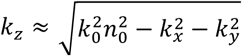; the latter follows from simple manipulations using *k* ≈ *k n*, owing to the paraxial approximation. We note that higher *k*_*x*_and *k*_*y*_values may correspond to imaginary values for *k*_*z*_. To prevent unphysical field blow-up, one should always choose *lm*( *k*_*z*_) > 0 to attain damping (of the high spatial frequency components).
3. Back transform to the desired intermediate result: 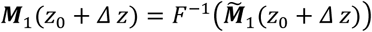

A consequence of the above procedure is that light running out of the computational window is re-entering at the other side, owing to the Fourier window periodicity. To that end, small absorbing layers were applied, corresponding to a small imaginary part of the refractive index, in stripes at the boundary. Its magnitude was slowly increasing from zero to some suitable value at the boundary to prevent back-reflection. The latter absorption was made effective via the second operator in Eq. 20, as explained next.

The second step of the FTBPM can be written as

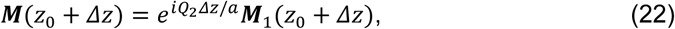

with a multiplication of all field components in real space with a corresponding factor, depending on *x* and *y*, *exp*( *Q*_2_*Δz*/*a*), which can be applied straightforwardly. It is noted that absorption is introduced via an imaginary part of the index, say, *n*(*x*, *y*) = *n*^′^(*x*, *y*) + *in*^″^(*x*, *y*), leading to *lm*(*Q*_2_*Δz*/*a*) ≈ *k*_0_*n*^″^*Δz*, with *n*^″^ > 0 corresponding to absorption and an absorption coefficient given by *κ* = 2*k*_0_*n*^″^. The factor of 2 is because *κ* refers to power decay. It is further noted that step indices, as at the boundaries of rhabdomeres, are smoothed in FTBPM to prevent unphysical scattering at the transitions, which may occur in particular if the structure is varying along *z*. A smoothing term 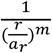 was applied to both exponent functions in Eq. 22; where *r* is the distance from the rhabdomere center, *a*_*r*_is the rhabdomere radius, and *m* is a constant defining the spatial width of the smoothing (Fig. S49 *F* and *G*).

In the case of the crystal cone, its constitutive material was considered homogenous (*Q*_2_ = 0 and *κ* = 0). Thus, Eq. 22 could be skipped, and Δ*z* in Eq. 21 set equal to the propagation length through the crystal cone. In the crystal cone (96) simulations, Δ*z* = 17 μm (Fig. S49*E*), if not specified otherwise, and its refractive index was 1.34. Owing to the cyclical nature of FFT with step regarding *Q*_1_, we added an absorption layer around the x- and y-simulation boundaries, preventing the electrical field from traveling over them.

Light propagation in the ∼80-μm-long R1-R6 (and R7+R8) rhabdomeres was simulated with Eq. 22, using 125 nm steps, which was a sufficiently small value (results remained virtually the same on lowering this value). The rhabdomere cross-section is a roundish disk, having 0.005/μm absorbance (97, 98) and 1.34 refractive index around it (75). Importantly, each R1-R7/8 has its specific rhabdomere diameter (11), with R1’s and R6’s being 1.8 μm; R2-R5s’ 1.6 μm; and R7/R8’s 1 μm. From the rhabdomere simulations, total absorbed power was calculated by integrating power *P*(*r*) = |Ψ(*r*)|^2^ over the whole rhabdomere (Fig. S49*H*):

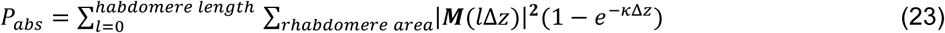

Absorbed photon flux, which is possible to measure electrophysiologically from photoreceptors using bump calibration, is related to absorbed power:

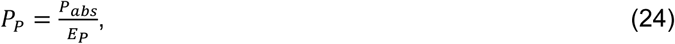

where 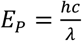 is the single-photon energy at 450 nm.

From the ray-tracing results above, we calculated FT BPM simulation electrical field at the lens inner surface. The optical distances (Fig. S49*B*) and field strengths (Fig. S49*C*) were calculated from the ray-tracing simulations. The ray optical distance was calculated for the electrical field phase:

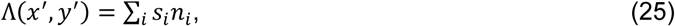

where each distance the rays traveled *s*_*i*_was multiplied by the material’s refractive index. *x*^′^ and *y*^′^ are the ray x- and y-positions, respectively, at the lens inner plain (Fig. S49*B*).

The relative power represented by a certain ray (being a ray resulting from the ray-tracing calculations) is (approximately) inversely proportional to the area it represents in the plane perpendicular to that ray, which 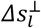 denotes, with

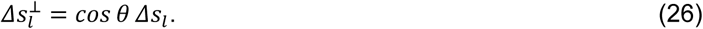

In the above, *l* is a label for the rays, *θ* is the angle between the ray and the *z*-axis and *Δs*_*l*_is 1/4^th^ of the area enclosed by the 4 nearest rays at the inner plain (near the lens).

So, the considered ray’s absolute value of the resulting relative field strength is given by

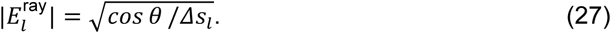

By evaluating the above and the corresponding phase, see Eq. 25, we know the field distribution as a result of the ray propagation. These serve as an input to calculate the input field for the FTBPM, introduced above

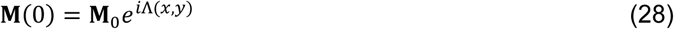

The field values, being the entries of ***M*** , have been interpolated from 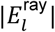 and the corresponding phase Λ(*x*, *y*) from the corresponding ray phases (Eq. 25), using Matlab procedure ‘*scatteredInterpolant*’ (Mathworks, USA) in which 512 x 512 points covered 16 x 16 μm lens area (Fig. S49, *B* to *D*).

##### V.3 Simulating R1-R7/8 photoreceptors’ optical spatial properties (*static cases*)

R1-R7/8 photoreceptor rhabdomeres’ spatial light-collecting properties were calculated by optical simulations. We varied the incident light angle between the point source and the lens optical axis, spanning ± 20.4° with 1.7° resolution to be comparable to previous intracellular recordings (11). From the simulations with varying point source angles, a rhabdomere’s total absorbed power, *P*_*abs*_(Eq.21), was calculated in each simulation point. Then the incident light angles’ total absorption curve was fitted with a Gaussian function to determine the tested rhabdomere’s optical receptive field (RF) shape, its center, width at half-maximum (*static* half-width or acceptance angle, Δ*ρ*^*s*^) and amplitude. Specifically, we examined two *static* scenarios of how a fixed rhabdomere position affects its optical RF shape; *i.e.,* the distribution of light rays it collects from the lens:

1. We analyzed a suite of RF simulations, where R1-R7/8 rhabdomeres were fixed at *different axial positions* away from the lens (Fig. S50). The axial distance between the lens and the rhabdomere tip was increased by varying the crystal cone thickness (the distance between the lens’s inner surface and the outer rhabdomere tip).
2. We analyzed a suite of RF simulations where R1-R7/8 rhabdomeres were fixed at *different lateral positions* by increasing the radial distance between the lens center axis and the rhabdomere tip position (Fig. S51).

**Fig. S50.**
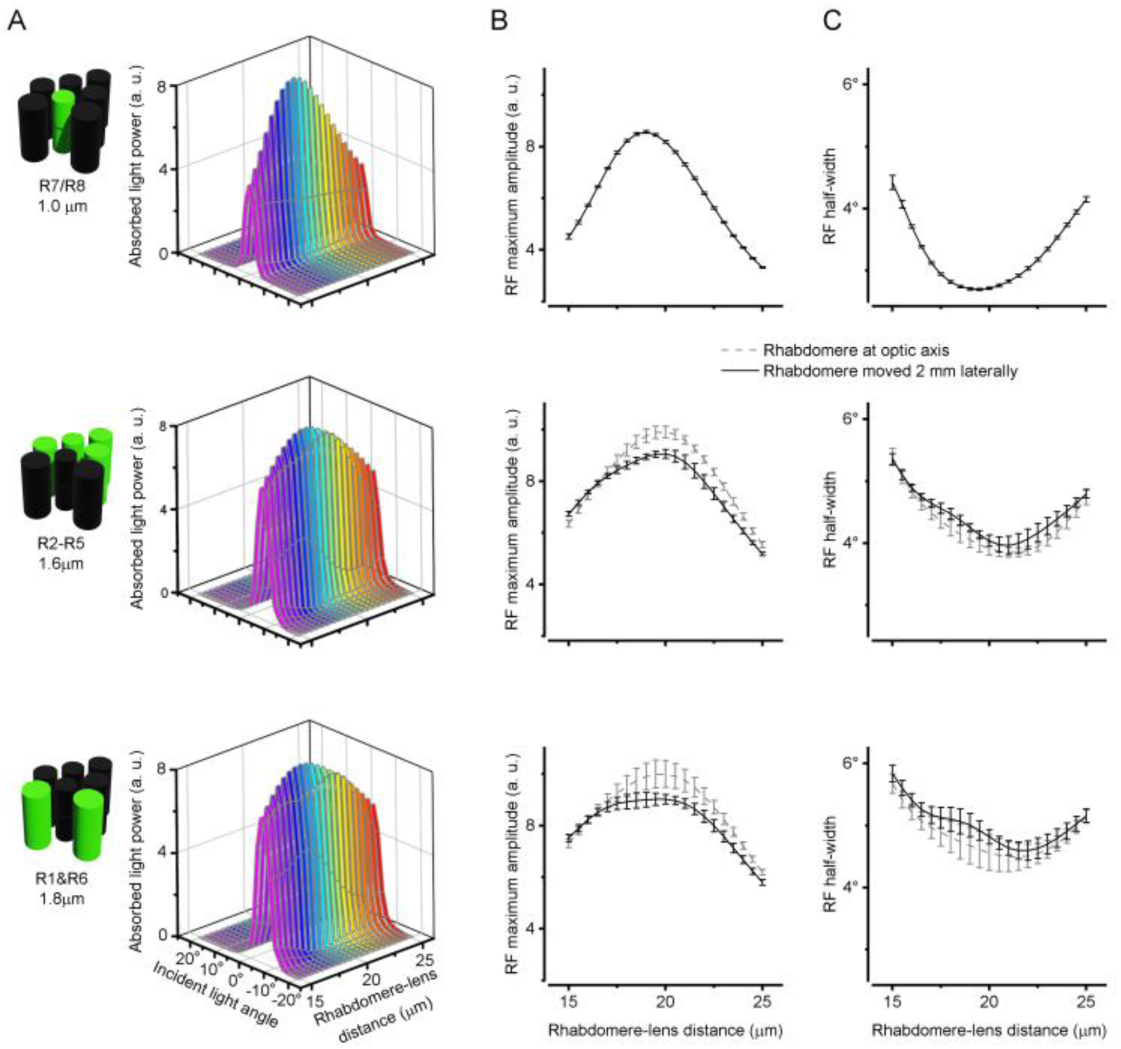
Changing a rhabdomere’s distance to the ommatidial lens changes its optical receptive field (RF) dimensions. (*A*) RF shape changes as a rhabdomere moves inward (away from the lens). Top: R7/R8 rhabdomere ∅ = 1.0 μm. Note, cone/pigment-cell aperture does not affect the lens center where R7/R8 rhabdomeres reside. Middle: R2-R5 rhabdomere ∅ = 1.8 μm; aperture ∅ = 5 μm. Bottom: R1 and R6 rhabdomere ∅ = 1.6 μm. aperture ∅ = 5 μm. (*B*) Total absorbed maximum power varies with rhabdomere-to-lens distance. Top: R7/R8 rhabdomere Middle: R2-R5 rhabdomere; Bottom: R1 and R6 rhabdomere. ((*C*) RF half-width (Δ*ρ*_*ls*_) varies with rhabdomere-to-lens distance. Top: R7/R8 rhabdomere. Middle: R2-R5 rhabdomere. Bottom: R1 and R6 rhabdomere. (*B* and *C*) Middle and Bottom: R1-R6s’ RFs simulated either the corresponding rhabdomeres at the lens center axis (gray) or their normal off-axis positions (black).

For both of these scenarios (Fig. S50 and Fig. S51), we tested three specific rhabdomere diameters (11): 1μm R7/R8 (Top rows); 1.6 μm R2-R5 (Middle); and 1.8 μm R1 and R6 (Bottom); see also Table S6. Moreover, in the RF simulations, we considered the (static) *aperture effect of cone and pigment cells* (Fig. S52) on the R1-R6’s optical input (black traces, with the aperture; gray, without). These densely pigmented cells border the crystal cone opening just above the rhabdomeres, forming an aperture (11). The outer edges of R1-R6 rhabdomere tips either touch or are just outside this aperture.

**Fig. S51.**
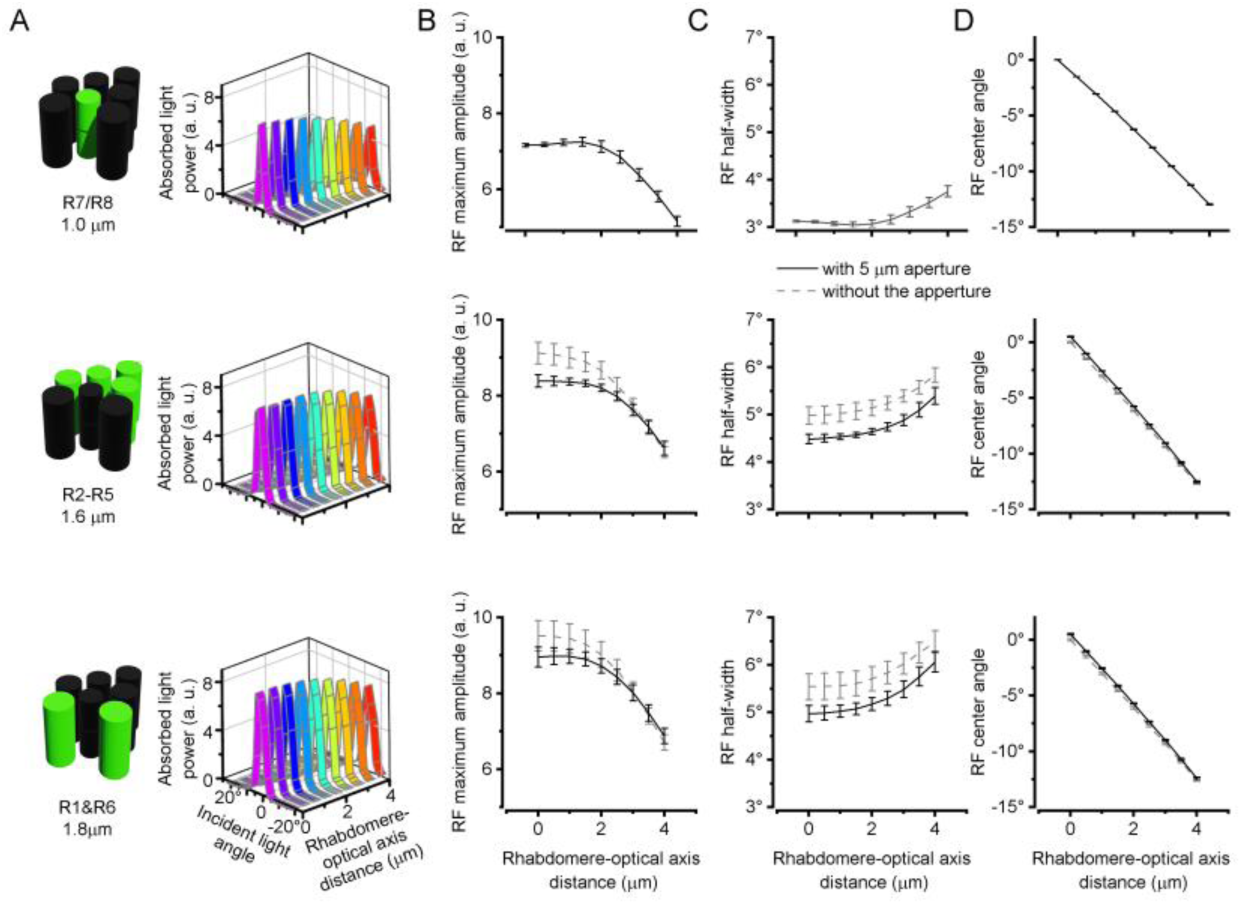
Changing a rhabdomere’s lateral (off-center axis) position changes its optical receptive field (RF) dimensions. (*A*) RF shape varies with a rhabdomere’s sideways positioning. Top: R7/R8 rhabdomere ∅ = 1.0 μm. Middle: R2-R5 rhabdomere ∅ = 1.6 μm. Bottom: R1 and R6 rhabdomere ∅ = 1.8 μm. (*B*) RF half-width (Δ*ρ*_*l*_^*s*^) varies with a rhabdomere’s sideways positioning. Top: R7/R8 rhabdomere. Middle: R2-R5 rhabdomere. Bottom: R1 and R6 rhabdomere. (*C*) Total absorbed power max amplitude varies with a rhabdomere’s sideways positioning . Top: R7/R8 rhabdomere. Middle: R2-R5 rhabdomere. Bottom: R1 and R6 rhabdomere. (*D*) Total absorbed power’s center position varies with a rhabdomere’s sideways positioning. Top: R7/R8 rhabdomere. Middle: R2-R5 rhabdomere. Bottom: R1 and R6 rhabdomere. (*B* to *D*), Middle and Bottom rows: with Æ 5 μm cone/pigment-cell aperture (black traces) and without it (gray).

**Fig. S52.**
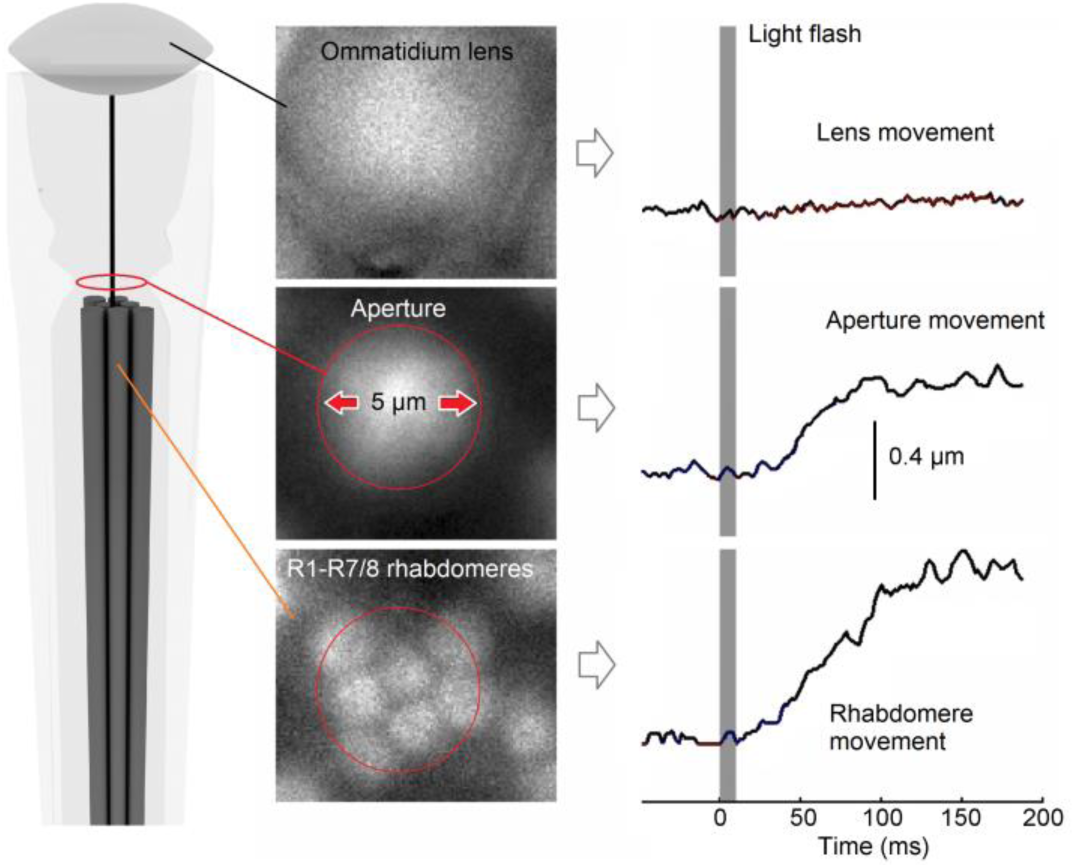
Ommatidial cone and pigment cell aperture – located between the crystal cone and the rhabdomere tips - shapes the light input to R1-R6 rhabdomeres. The R1-R6 rhabdomere adherens-junctions connect to the cone cells (99). Therefore, during a photoreceptor microsaccade, with rhabdomeres contracting, the aperture drags behind, moving about half as much sideways as the rhabdomeres. We call this delayed aperture movement the “swing effect” (11). These local structural photomechanical movements to green flashes were measured using the IR- cornea-neutralization method (see Section II.8.ii, above) while raising and lowering a fly underneath the microscope objective with piezo steps. Thereby, we could change the focus from the lens surface (above) to the cone and pigment cell aperture (middle) to the rhabdomeres (below) while light-activating the photoreceptors. Notice that the ommatidium lens remains stationary throughout the experiment, similar to X-ray imaging (Fig. S3E). Images modified and adapted from (11).

The aperture was simulated as a 5 μm diameter round opening, estimated from the light microscopy images (Fig. S52) (11). Its thickness (12) was 2 μm with 2.8% total transmittance. In the previous wave-optical modeling studies (75, 93, 94), a different type of aperture, which tightly surrounds the rhabdomere with the same diameter, inadvertently arises from the mode simulation equations. But to our knowledge, the real cone and pigment cell aperture effect on a rhabdomere’s optical receptive field shape had not been considered before. 0.4 µm

Fig. S50 shows how changing the rhabdomere-to-lens distance (Fig. S50*A*) changes the optical RF shape (Fig. S50 *B* and *C*). The simulations indicated that Δ*ρ*_*l*_^*s*^, a rhabdomere’s optical acceptance angle (RF half-width; Fig. S50*B*) is at its narrowest at ∼21 μm from the lens inner surface. At this point, the rhabdomere’s light absorption power reaches its maximum (Fig. S50*C*) for all the three simulated rhabdomere diameters. Note that as the rhabdomeres contract during *in vivo* light stimulation, their axial component moves their tips ∼2 μm away from the lens (11), which is just a fraction of the total range (10 μm) simulated here.

Seven rhabdomere tips (with R7/R8 counted as one) make the characteristic lopsided pattern behind the ommatidium lens. Naturally, with R1-R6 photoreceptors positioned off-center, the lens center optical axis never passes through their rhabdomeres. Fig. S51 shows how changes in a rhabdomere’s lateral position, away from the lens optical center axis, change its optical RF shape. This offset causes the optical RF centers to tilt 3°/μm (Fig. S51*D*) in all the simulated rhabdomeres (of different diameters). The RF tilts to the opposite way of the offset direction because the ommatidium lens inverts the rays. As the offset becomes larger, the optical RF acceptance angle (Δ*ρ*_*l*_) broadens, and the maximal absorbed power reduces. Such physics happens because the lens obscures some fraction of the light that enters with large angles (Fig. S51 *B* and *C*). To establish how the cone-pigment cell aperture shapes R1-R6 rhabdomeres’ RFs, we simulated their light input with and without the 5 μm cone-pigment cell aperture in front of them. In all tested conditions, the aperture narrowed R1-R6 rhabdomeres’ RFs in respect to the corresponding simulations without it.

##### V.4 Generating light current

Light-induced current (LIC) responses were simulated from the absorbed photon flux, *P*_*P*_(Eq. 24), using a four-parameter stochastic photon sampling model (Fig. S53), a mathematical representation of phototransduction in microvilli (40). It closely reproduces the real *in vivo* sampling/integration dynamics, generating realistic simulations (40). For a given light stimulus, the model converts the successfully absorbed photons to quantum bumps (QBs) and integrates them to a LIC, as set by (i) the number of a *Drosophila* photoreceptor’s photon sampling units (Fig. S53*B*, 30,000 microvilli in an R1-R6), (ii) the microvilli refractoriness distribution (Fig. S53*H*), (iii) the QB latency distribution (Fig. S53*F*) and (iv) the adapting QB waveform (Fig. S53*E*). Because *P*_*P*_was not directly related to the point source power, we chose the maximal absorbed photon flux, based on intracellular recordings (11), and *P*_*P*_was scaled to this. We used the established QB latency (Fig. S53*F*) and refractory Gamma distributions (Fig. S53*H*) (40) at 20 °C. Gamma distributions contain *n* and *τ* parameters:

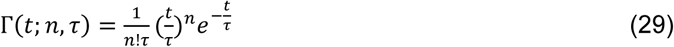

Temperature affects (82) much the latency (Q_10_ = 3.4) and refractory distribution half-widths, and thus the refractory distribution *τ* parameter in the simulations. To compare the simulations to the typical recordings (11) at 25 °C, which is *Drosophila*’s preferred temperature (11), the simulations were Q_10_-scaled (11, 39, 40) when needed.

**Fig. S53.**
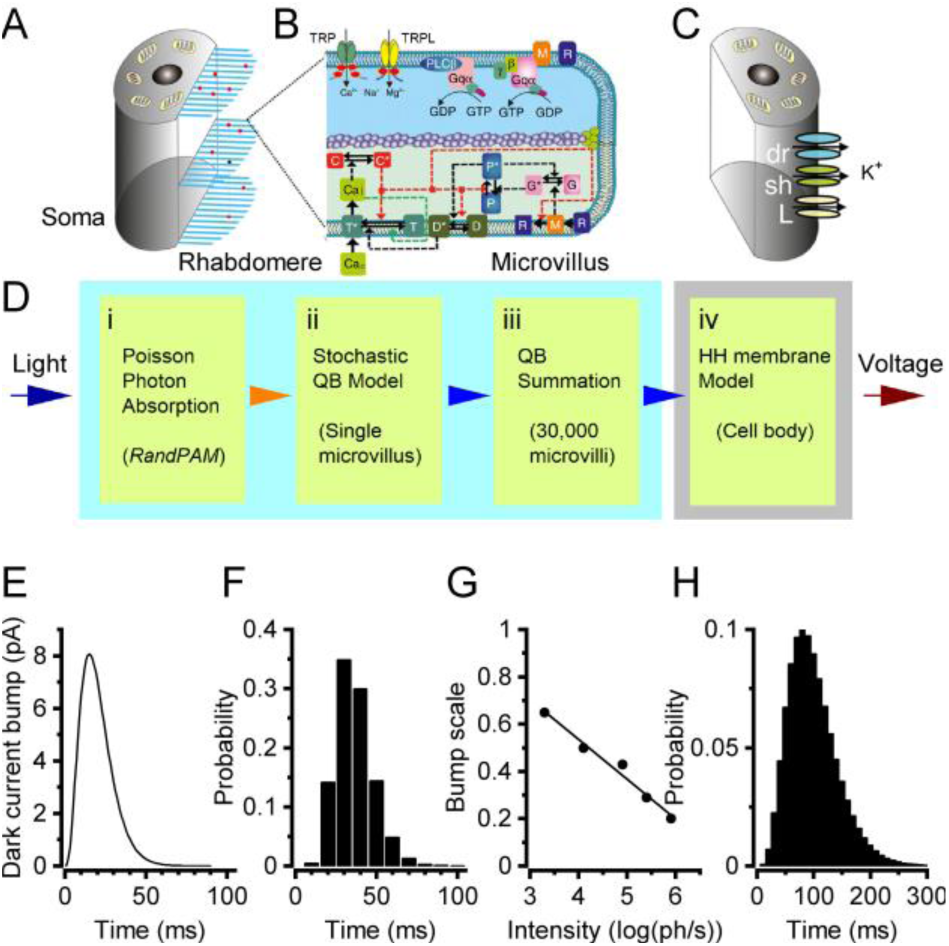
Schematic structure of a *Drosophila* photoreceptor sampling model. (*A*) A fly photoreceptor is functionally divided into the photo-sensitive membrane (rhabdomere) and the photo-insensitive part (soma). (*B*) The critical phototransduction cascade molecules inside a single microvillus (above) and their differential equation directions (arrows, below) (40). (*C*) Voltage-gated K^+^ conductances on the soma (78, 100). (*D*) The model contains four modules (i-iv) (40). (*E*) Dark current quantum bump (QB). (*F*) QB latency distribution at 20 °C. (*G*) QB feedback adaptation. Linear fit for Log-linear relationship between experiments (101) and simulations. (*H*) QB refractory period distribution.

An average dark-adapted LIC QB waveform (Fig. S53*E*) was modeled as a Gamma function (40, 60, 82, 102). Light-adaptation (negative feedback) was modeled by reducing the QB’s amplitude and shortening its duration (60, 82). The QB duration is controlled by the Gamma-function’s *n*-variable, which fastens the QB’s rising phase, similar to light-adaptation (60, 82, 101). These QB adaptations were controlled by one parameter used as a multiplier to the QB amplitude and *n*. For short stimuli, we fitted the published LIC responses to 5 ms flash experiments (101) at 20 °C. We selected the multiplier by hand so that the macroscopic current was the same in the simulations as in the actual recordings. After fitting the individual data points, we could perform a linear fit in a log-linear scale between the light intensity and the multiplier (Fig. S53*G*). For dynamic simulations, the total photon flux for calculating the QB multiplier from the fit was obtained as the sum of absorbed photons between the start of simulations and the QB generation time point, using a 5 ms time-bin to match the measured QB latency data (101). The QBs were appropriately light-adapted by a controlled pre-simulation photon exposure. This procedure ensured that the QB light-adaptation dynamics and range followed physiologically accurately the simulations’ light intensity modulation (photon flux changes). The maximal photon flux was set to 11.1 x 10^6^ph/s.

##### V.5 Light current to voltage response conversion

The macroscopic LIC response was converted to a voltage response using the *Drosophila* photoreceptor HH-membrane model (Fig. S53*C*) (78, 95, 100). The model consists of several ion channels: the two LIC- channels (*trp* and *trpl* – here, combined), in which conductance (101) was calculated by dividing the light current with the -80 mV driving voltage (in the voltage-clamp-configuration), three K^+^-channels and two passive leak channels, which approximate the mean synaptic feedback effect (36, 78). The voltage responses were simulated with an improved HH-model, which could now directly compute light-induced conductances instead of using a global voltage-feedback as was done before (11, 39, 40). This modification simplified and expedited the simulations while producing similar results as the old model. We obtained the RF’s peak voltage response from the voltage simulations while fitting the total RF shape with a Gaussian function. This procedure further gave us the RF half-width and peak position. The used parameter values were the same as in the early *Drosophila* membrane HH-models (100), except that the total input-resistance was set to ∼200 MΩ, which better approximates the more commonly recorded values (11, 78).

##### V.6 Estimating R1-R7/8 voltage response RF half-width for *static* rhabdomere positions

An R1-R7/8 photoreceptor’s *voltage output* receptive field (RF) half-widths (acceptance angles, Δ*ρ_v_*^*s*^) were estimated similar to their optical *light input* RF half-widths (Δ*ρ_l_*^*s*^), but now using the corresponding voltage simulations. The stochastic four-parameter model generated realistic LIC responses to light flashes of 22,000 photons/s (maximum intensity at the RF peak (11)). We modeled R7/R8 rhabdomeres at the on-center (the ommatidium lens’s optical axis) and R1-R6 rhabdomeres 2.0 μm off-center, with their outer edges touching the cone-pigment cell aperture. We used 10 ms flashes, in which intensity was set by the corresponding relative intensity (based on optical simulations with varying incident light angles).

At the R1-R7/8 rhabdomeres’ measured dark-resting positions, with respect to the ommatidium lens, their *static* voltage response RF half-widths (Δ*ρ*^*s*^_*v*_) were: 4.6° ± 0.2 for 1.0 µm diameter R7/R8 rhabdomere, 6.4° ± 0.4 for 1.6 µm diameter R2-R5 rhabdomeres and 7.1° ± 0.4 for 1.8 µm diameter R1 and R6 rhabdomeres, respectively (Fig. S54). These values were predictably larger than their corresponding optical absorption power RF half-widths (Δ*ρ*^*s*^_*l*_): 3.12° ± 0.02, 4.5° ± 0.1 and 5.0° ± 0.1 (Table S6), respectively. These differences result from the compressive nonlinearities in the transformation from photon absorptions to voltage responses, such as slow QB-waveform dynamics and membrane conductances (40, 60, 100), which grant higher gain to weaker light changes, fattening the RFs’ midriff and tails.

In the actual electrophysiological voltage response recordings (during relative dark adaption) (11), the estimated average *dynamic* wild-type R1-R6 photoreceptors’ RF half-width (Δ*ρ_v_*^*d*^) was even wider: 9.65° ± 1.06, being about twice the simulated *static* optical RF half-widths (Δ*ρ_l_*^*s*^). The differences between the simulated and recorded voltage response RFs must arise from the *dynamic* processes that the *static* simulations lack. For example, the voltage RF simulations lacked the rhabdomere movements and slow QB adaptations, which do not fully recover (11) during the short experimental (500-1,000 ms) stimulus intervals. We know from the goniometric *in vivo* rhabdomere imaging (Fig. 3 in the main paper, and Section II.6, above) that during *in vivo* electrophysiological recordings with repeated light-flash stimulation (11), the rhabdomeres and their RF centers must continuously shift in different positions in respect to the start state, widening the RF estimates. Moreover, during the experiments between the flash stimuli, there can be additional rhabdomere movements caused by spontaneous intraocular muscle activity (11, 13).

Interestingly, in the dark-adapted intracellular electrophysiological recordings, with the R1-R6 rhabdomeres positioned about 2 µm laterally off the ommatidium lens optical center axis (Fig. S47A), the voltage RFs often showed skewness/asymmetry (11). This phenomenon is readily reproduced in the RF simulations when R1-R6 rhabdomere is positioned 2 µm off the ommatidium lens’ optical center axis (Fig. S54 *B* and *C*).

**Fig. S54.**
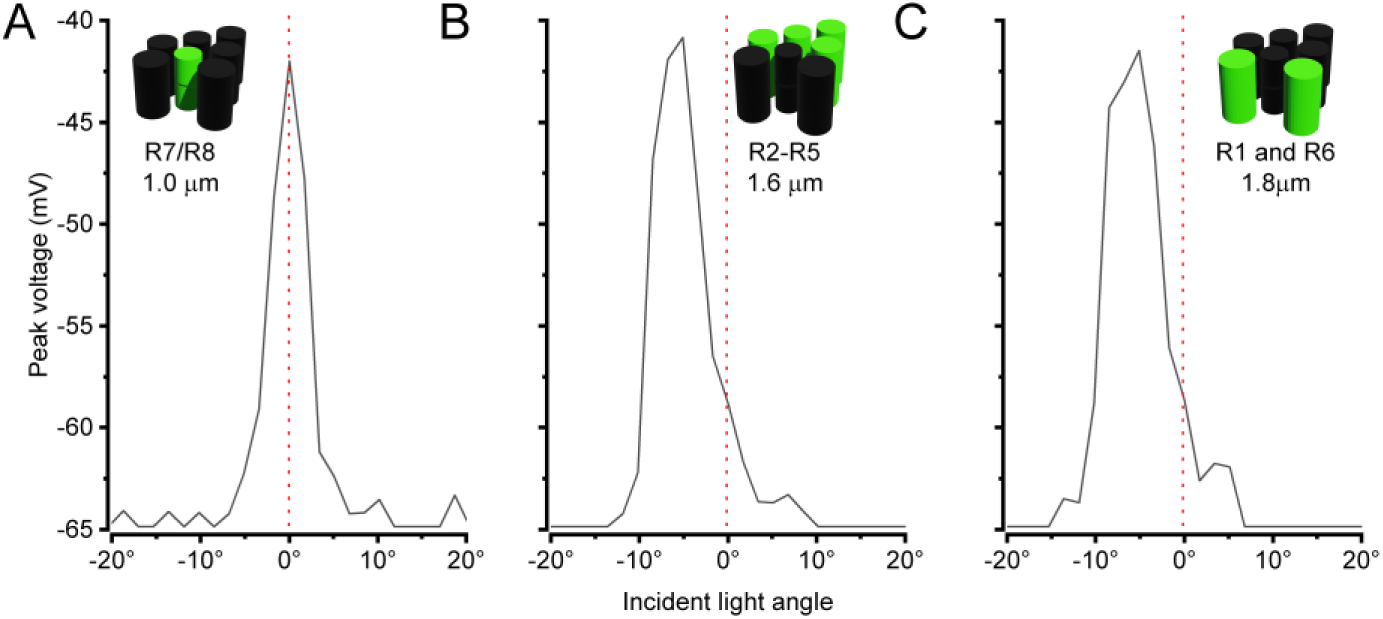
R1-R7/8 photoreceptor receptive fields (RFs) based on simulated voltage responses. (*A*) R7/8 Δ*ρ*_*v*_^*s*^= 4.6 ± 0.2° with 1.0 μm rhabdomere diameter. R7/8s’ RFs are Gaussian. (*B*) R2-R5 Δ*ρ*_*v*_^*s*^= 6.4 ± 0.4° with 1.6 μm rhabdomere diameter. rhabdomere diameter. Because R2-R5 rhabdomeres are off-center in respect to the ommatidium lens, their RFs are skewed. *(C)* R1 and R6 Δ*ρ*_*v*_^*s*^= 7.1 ± 0.4° with 1.8 μm rhabdomere diameter. Because R1 and R6 rhabdomeres are off-center in respect to the ommatidium lens, their RFs are skewed.

##### V.7 Photomechanical rhabdomere movements

Our previous studies (10, 11) revealed the biphasic R1-R7/8 photoreceptor microsaccade photomechanics to a flash stimulus consisting of a fast contraction phase (rise) followed by a slower relaxation phase (decay). Here, we further measured the microsaccades’ frequency response function (see Section II.6., above). Moreover, we showed in Section II.8. (above) that in each ommatidium, R1-R7/8 rhabdomeres were structurally coupled - possibly by cross-connecting R1-R7/8 tip-links. Therefore, even a single photoreceptor light-activation made all its ommatidial sister photoreceptors contract/move in unison. Based on these results, the microsaccadic rhabdomere motion (*x*_*d*_) was modelled as a spring-dampener system, which closely reproduced the measured dynamics (Fig. S55):

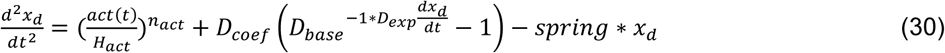

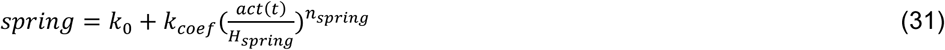

The microsaccadic movement system consisted of three forces: (i) the mechanical activation force, connected to the photoreceptors’ photon absorptions; (ii) the dampener force, resisting the change in the resulting photomechanical movement; and (iii) the spring force, returning the rhabdomeres to their original positions. The equations lacked the mass term as the other terms accounted for this. The dynamic simulations used the Euler method to solve, numerically, the differential equation with 1 ms step. The various model parameters were fitted to match the recordings (Fig. S55):

- Light activation *act*(*t*) connected the four-parameter-photoreceptor-model (see Section V.4, above) to the activation force. The activation *act*(*t*) was the absorbed photons leading to PIP_2_ cleavage (10), following the four-parameter model’s latency distribution dynamics. In each ommatidium, the light input was the sum of its seven (R7 and R8 fused) rhabdomeres’ total absorption. *H*_*act*_(= 9,000 ph/µm^1/2^) controlled the rhabdomere movement amplitude when maximal photon flux was 900 ph/ms. The activation co-operation parameter, *n_act_*, was set to 2, which reproduced the rhabdomeres’ photomechanical creep-up and creep-down behaviors seen in Fig. S55*A*.
- The dampener resisted the change in the rhabdomere speed (*dx*_*d*_/*dt*). We set the dampener to have a maximal force with the positive movement speeds, *D_coef_* =0.0001 μm/ms^2^. The dampener base (*D_base_* = 2) and the dampener exponent (*D_exp_* = 3,900 μm/ms) defined the dampener’s rectifier shape (its fast rise and slow decay). The dampener made the movement model unstable with brighter than 900 ph/ms light stimuli. Consistently, at such high light intensity levels, the photoreceptors’ intracellular pupil mechanism and the ommatidial screening pigments actively filter off any brighter photon flux to maintain appropriate QB production rates, enabling maximum information flow while preventing saturation (11, 39).
- The spring constant, *spring*, depended on activation *act*(*t*), increasing with light input. We set the spring constant without activation (k_0_=0.0001/ms^2^) so that its decay was slow and the impulse response peaked in a reasonable time. The average microsaccade dynamics of characteristic recordings (measured from five wild-type photoreceptors) to brief positive and negative contrast changes (Fig. S55 *B* and *C*) were used to adjust the activation-dependent spring constant: *H_coef_* =0.00115 1/ms^2^, *H*_*spring*_= 200 ph/ms and *n_act_* =1.3.

**Fig. S55.**
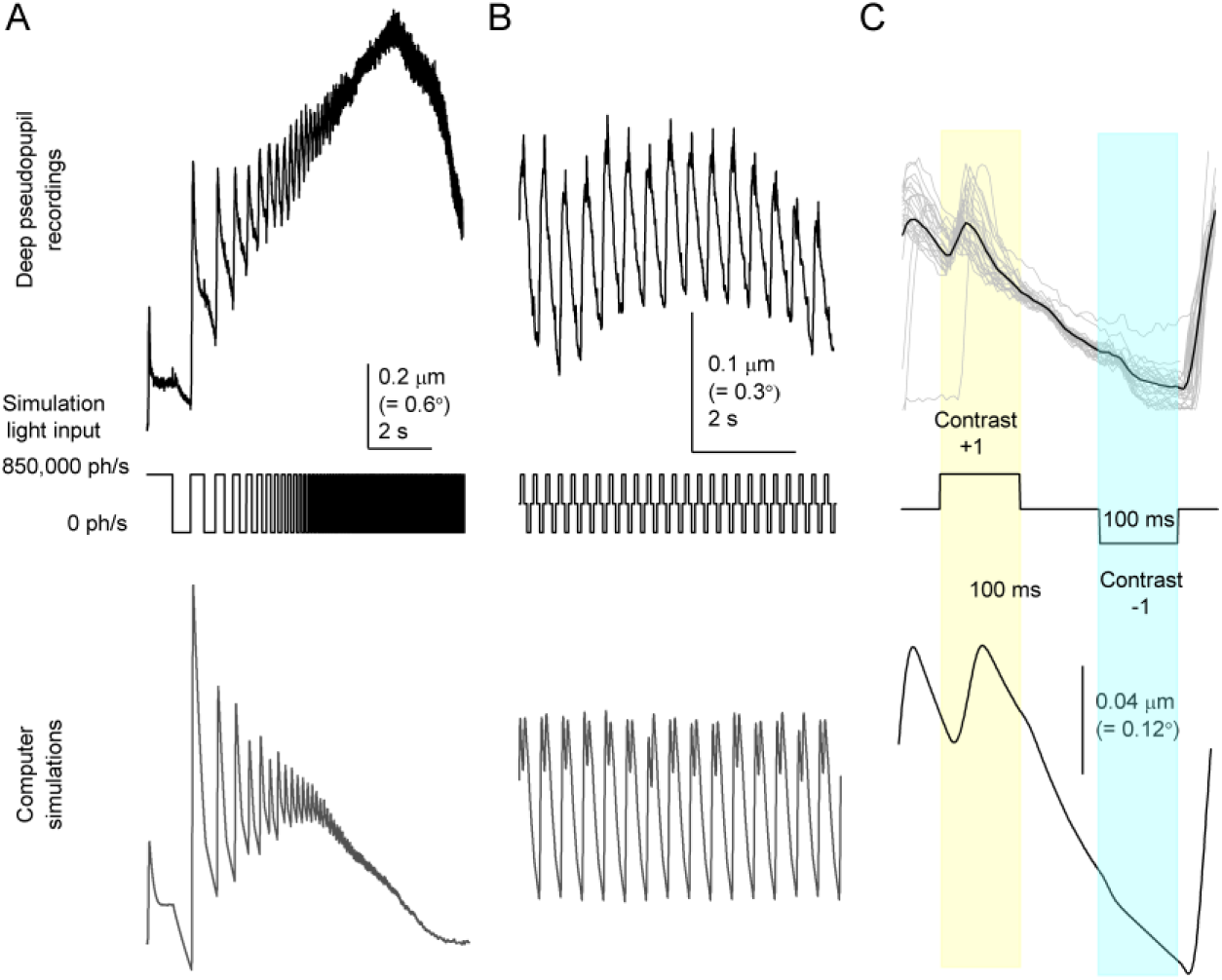
Modeling ommatidial rhabdomere photomechanics as a spring-dampener system. (*A*) Above: a typical recording of wild-type photoreceptor microsaccades to frequency accelerated pulsed +/-1 contrast stimulus. Below: a corresponding microsaccade simulation. In the simulation, *n_act_*-parameter was adjusted to mimic the rhabdomeres’ photomechanical creep-up and creep-down. (*B*) Above: a typical photoreceptor microsaccade recording to repeated 100-ms- long +/-1 contrast pulses. Below: Simulation to the same stimulation. (*C*) Above: recorded microsaccade waveforms to repeated 100-ms-long +/-1 contrast pulses show characteristic variability (gray traces); black trace gives their mean. Below: the corresponding mean simulated microsaccade waveform is very similar, showing fast activation-phase (up-surge) and slower recovery-phase (down-surge) dynamics superimposed on a longer adapting trend (downwards slope).

Only those rhabdomeres, which saw (*i.e.,* their RFs directly experienced) light changes, generated an intra-ommatidial R1-R7/8 movement locally. Whereas in those ommatidia, which did not experience (see) light changes, the rhabdomeres were still (*cf*. Fig.1F-I in the main paper; Fig. S31 to S33 in Section II.8.ii, above; Movie S10).

##### V.8 Photoreceptor responses to spatiotemporal stimuli

To study how *Drosophila* photoreceptors respond to moving stimuli, we simulated R1-R7/8 voltage responses to spatiotemporal visual objects crossing their receptive fields. For generating light inputs to the models, we ray-traced R1-R7/8 rhabdomere RFs - using their measured intraommatidial positions (Fig. S47*A*) - onto a virtual surface; with the rays being cast from the center of the ommatidium lens (its outer face). The resulting RFs at the virtual surface (Fig. S56*A*) were interpolated from the RF rays, divided by their surface areas. For generating the rhabdomeres’ light inputs, we convolved their RFs fields with the stimulus image/video at the virtual surface, assuming that the screen is Lambertian (*i.e.,* with every angle having an equal light power output). The resulting light series was normalized by the maximum absorbed photon flux. This outcome was then fed, as the input, to the combined four-parameter/HH-model to generate the simulated voltage response to the given stimulation (Fig. S56 *C* and *D*).

Fig. S56 *E* and *F* show examples of two dots moving across two similar R5 photoreceptors’ RFs (one located in the right eye and the other in the left eye), in which movement directions were along (in the same way; in the right eye) or against (in the opposite way; in the left eye) the given dot-movement direction. Thus, effectively, these two cases also simulate the corresponding R5 rhabdomere movements in the binocular left and right eye ommatidia; sampling light from the same small frontal area at the distance, where their RF fields overlap (near) perfectly.

**Fig. S56.**
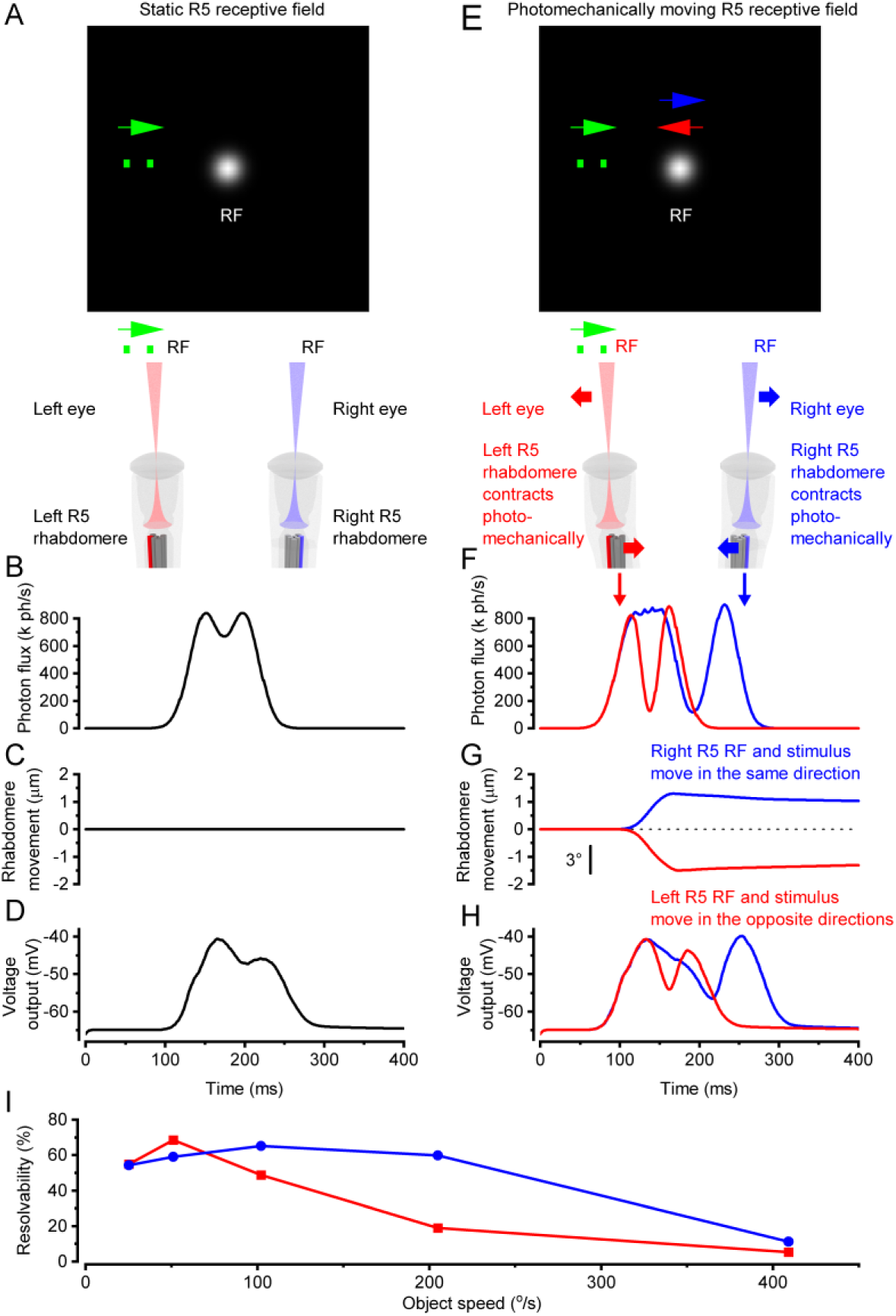
Photoreceptor photomechanics dynamically shift and narrow their receptive fields (RF) and voltage responses. (*A*) Rays were cast from characteristic ommatidial optics to a virtual screen to simulate an R5 photoreceptors’ RF. In *static simulations*, the RF remained immobile. The 5 x 5 cm screen was 5 cm away. Resolution: 0.1°/px The R5 rhabdomere was 17 μm from the lens with 2 µm off the center-axis, having the cone/pigment-cell aperture in front of it. Stimulus: two dots (green) moving at 102°/s and crossing the R5 RF. Dot size is 1.7 x 1.7° and inter-dot-distance 6.8°. (*B*) Optical rhabdomere input for two dots crossing a *static* (immobile) R5 RF (Δ*ρ*_*l*_^*s*^= 4.46°), generating a two-peaked light input when maximal. (*C*) In the naïve *static* case, the R5 rhabdomere was considered immobile. (*D*) Voltage response to the two dots crossing a *static* R5 RF; the resultant light input is used in (B). The maximal photon flux ∼8 x 10^5^ absorbed photons/s at 25 °C. The QB was pre-light adapted by 45,000 photons (see section V.5) to generate ∼20 mV responses. Thus, *if the fly compound eye optics were static* (8, 9), R5s in the left or right eye would generate identical voltage responses irrespective of the stimulus movement direction. (*E*) In *dynamic simulations*, the moving light stimulus enters the RFs and thus begins to excite an R5 photoreceptor. Consequently, its RF narrows and moves - with the photoreceptor contracting both axially and sideways. Notably, light evokes mirror-symmetric (opposing) photomechanical left and right eye photoreceptor microsaccades (as quantified experimentally *in vivo* in Sections I. and II., above). (*F*) Optical rhabdomere input for two dots crossing a *dynamic* R5 RF (Δ*ρ*_*l*_^*d*^= 4.05°). The right eye R5 RF moves in the same (blue) direction, and the left eye R5 RF in the opposite (red) direction as the dots. Notice how axially contracting and sideways moving rhabdomere improves optical resolution (the dip between the peaks) compared to the stationary rhabdomere in *B*. (*G*) Photomechanical R5 rhabdomeres’ sideways movement, 𝜈. The right RF is moving in the same (blue) and the left RF in the opposite (red) direction, in relation to the dot movement. (*H*) Voltage response to two dots crossing *dynamic* R5 RFs; the resultant light inputs in (F). The maximal photon flux: ∼8 x 10^5^ photons/s; the temperature: 25 °C. The bump size was adapted by 45,000 photons (see Section V.5) to generate ∼20 mV responses, matching the real intracellular recordings (11). The right R5 photoreceptor resolved the dots better (as quantified by the larger dip between the peaks; Rayleigh criterion (11)) because these moved in the same direction as its RF (blue), giving its phototransduction more time to separate them. Notice how the responses of the moving rhabdomeres resolved the moving dots better than the stationary rhabdomere in *D*. (*I*) Photoreceptor voltage response resolvability improves when the RF and stimulus (dots) move in the same direction.

The moving R1-R7/8 rhabdomeres’ Gaussian RFs were controlled by their intra-ommatidial photomechanical movements in the virtual screen simulations. How a rhabdomere’s intra-ommatidial light capture and the subsequent microsaccade (of axial and lateral movements) affected and moved its RF (at the virtual screen with ommatidial lens inverting the directions) was estimated from the optical light-point-source simulation results (Section II.6.). In Fig. S51 (above), we showed that the receptive field center moved 3°/μm. A rhabdomere moved simultaneously inwards and sideways (11), with its distal tip’s starting position being 17 μm from the inner ommatidium lens surface (see also Fig. S15, above). Table S6 catalogs how the RFs of different sized rhabdomeres behaved, giving their dynamic light input acceptance angle (Δ*ρ*_*l*_^*d*^) estimates, when the rhabdomere-to-lens-distance increased from 17 to 19 µm (Fig. S50).

**Table S6.**

| Rhabdomere diameter | R1-R7/8 rhabdomeres' *optical* light input RF half-widths (acceptance angles) |  | R1-R7/8 rhabdomeres' *optical* light input RF maximum amplitude |  |
| --- | --- | --- | --- | --- |
| | **Hypothetical static case ($\Delta\rho_l^s$):**<br>no R1-R7/8 photomechanics.<br>Fixed 17 $\mu\text{m}$ rhabdomere-to-lens distance | **Realistic dynamic case ($\Delta\rho_l^d$):**<br>Rhabdomere-to-lens distance increases from 17 to 19 $\mu\text{m}$ | **Hypothetical static case:** no R1-R7/8 photomechanics.<br>Fixed 17 $\mu\text{m}$ rhabdomere-to-lens distance | **Realistic dynamic case:**<br>Rhabdomere-to-lens distance increases from 17 to 19 $\mu\text{m}$ |
| **R7/8: 1 $\mu\text{m}$** | **3.12°** | **2.7°**; RF half-width reduces by -0.42° | **7.16 (a.u.)** | **7.60**; collects more photons by +1.4 |
| **R2-R5: 1.6 $\mu\text{m}^*$** | **4.48°** | **4.05°**; RF half-width reduces by -0.43° | **8.39 (a.u.)** | **9.75**; collects more photons by +1.36 |
| **R1 and R6: 1.8 $\mu\text{m}^*$** | **4.99°** | **4.67°**; RF half-width reduces by -0.32° | **8.96 (a.u.)** | **9.90**; Collects more photons by +0.94 |
| **\*5 $\mu\text{m}$ aperture touching the rhabdomere tip's outside edge** |  |  |  |  |

Notably, these are realistic but conservative *mean estimates for dark-adapted R1-R7/8 photoreceptors with round-tip cylindrical rhabdomeres*. Our previous study (11) compared the R1-R6 photoreceptors’ electrophysiologically measured angular sensitivity functions to their two-dot separation responses. These were measured immediately, one after the other from the same cells (11). We deduced that the highest acuity photoreceptor’s acceptance angle would need to narrow down to ≤3.7° dynamically to achieve its two-dot response resolution. Whereas for the most R1-R6s, their acceptance angles would need to contract to ∼4-4.5°. We attributed these Δ*ρ*_*l*_^*d*^-differences to the natural variations in the individual R1-R6s rhabdomere diameters and their eye-location-dependent orientation in respect to the given stimuli – *i.e.,* whether the two dots crossed their *oblong rhabdomere tips* (Fig. S47) along the long (ð larger acceptance angle) or short diameter (ð smaller acceptance angle). Moreover, Table S6 simulations do not include the RF narrowing by the intracellular pupil mechanism during light adaptation (94, 103, 104). Thus, the corresponding light-adapted acceptance angles should be smaller yet.

##### V.9 Neural superposition

For neural superposition (Fig. S57*A*), we simulated neighboring ommatidia on a virtual screen. A single ommatidium’s RF pattern is shown in Fig. S57*B*, with parameters taken from the optical simulations (Fig. S49 to S51). Lens positions and the standard hexagonal lens patterns were calculated based on the known parameters (75, 93): 16 µm distance between neighboring lenses (12), 5° angle between the lens centers (results from the eyes’ hexagonal ommatidia tiling with 4.5° interommatidial angle). The R1-R7/8 rhabdomere pattern in the ommatidia was taken from high-resolution EM and live microscopy images (11). The rhabdomere center positions were measured, with R7/8 rhabdomeres expected to be on the lens optical center axis during dark-adaptation. For the best overlap in the neural superposition pattern, the distances from R7 were multiplied by 0.85 (Fig. S57 *C* and *D*; Table S1) (R1-R6 are 0.2 μm closer to the center at the rhabdomere distal tip (12)). Because R1, R2, R3, R4, R5, and R6 rhabdomeres have different diameters (11) and are different distances away from the ommatidium lens center, the neural superposition pattern cannot align perfectly, as shown in Fig. S57 *C* and *D*. These results directly equate to Pick’s (23) findings, which showed that photoreceptor optical angles vary between ommatidia, leading to imperfect neural superposition tiling. The slight discrepancies with photoreceptors’ positions and photomechanical movements caused voltage responses in superpositional photoreceptors (Fig. S57*E*) to be slightly misaligned and effectively increase the over-completeness of the photoreceptor matrix (11, 23).

**Fig. S57.**
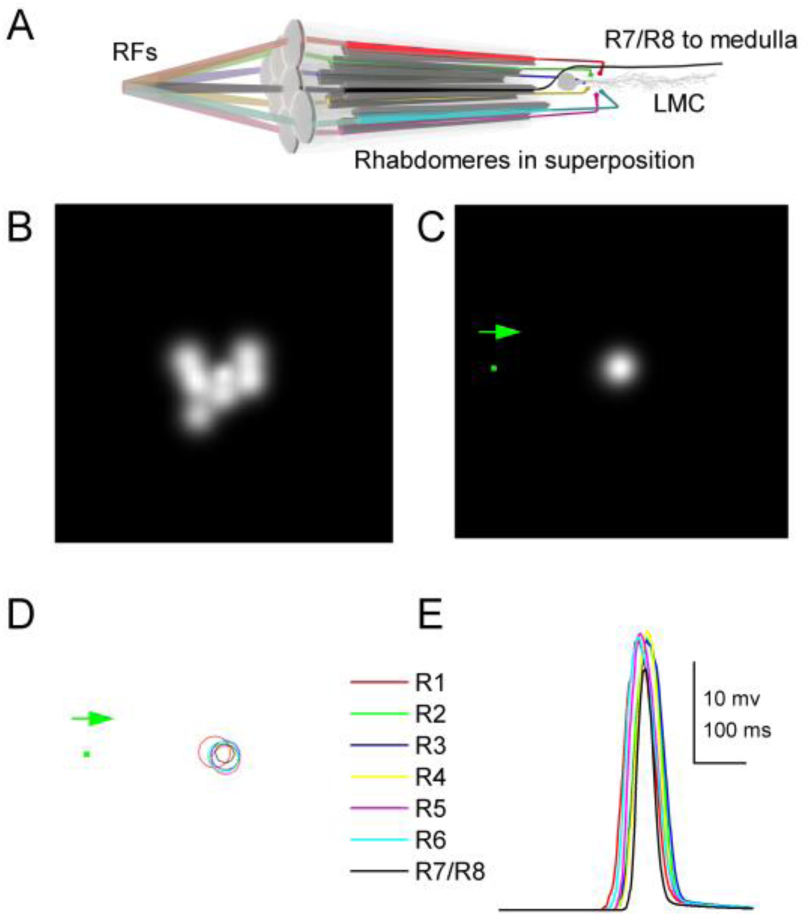
R1-R7/8 receptive fields (RFs) as seen at a 5 x 5 cm virtual screen, 5 cm away. (*A*) In neural superposition wiring, R1-R6 photoreceptors from six neighboring ommatidia optically collect light information from overlapping receptive fields (RFs) and transmit it to large monopolar cells (LMC, L1-L3) and an amacrine cell (30, 32) in the lamina. R7/R8 photoreceptors share some information with R1 and R6 through gap junctions (22, 105) but form their synapses in the medulla. (*B*) Single ommatidium R1-R7/8 rhabdomeres’ optical RFs. (*C*) Optically superpositioned R1-R7/8 RFs of the neighboring ommatidia and a point object (green dot) traveling towards them. (*D*) R1-R7/8 rhabdomere RF half-widths (circles) of the neighboring ommatidia in optical superposition with the same approaching dot (green). (*E*) Imperfectly aligned superpositional R1-R7/8 voltage responses for 1.7° x 1.7° dot crossing them with 100 °/s (the green dot in *C* and *D*). The simulations of the superpositional R1-R7/8 photoreceptors ommatidia included their photoreceptors’ photomechanical microsaccadic movements. The photoreceptors’ voltage responses were adjusted to have maximal light input of 350,000 photons/s and pre-adaptation of 35,000 photons in 25°C.

##### V.10 New theory for mirror-symmetric microsaccadic sampling of dynamic stereo-information

We now extend the theoretical framework from simulating a single ommatidium’s spatiotemporal sampling dynamics to simulating an ommatidia group’s sampling dynamics within the binocular (stereo) eye regions. *Drosophila*’s microsaccadic sampling of stereo-information was simulated using two 4 x 5 ommatidia grids, representing its two eyes’ frontal sampling matrixes at the fly head’s central (antenna) level. Each ommatidium’s seven rhabdomeres’ RFs (R7/R8 fused) were simulated on a virtual screen. The distance between the nearest left and right eye ommatidia is 440 μm (inter-eye distance), and their lenses diverge 2°, as determined from the X-ray images (Fig. S2B). Furthermore, their rhabdomeres moved out- and downward at a 45° angle, as determined by the goniometric measurements (Fig. S25; Movie S4).

**Fig. S58.**
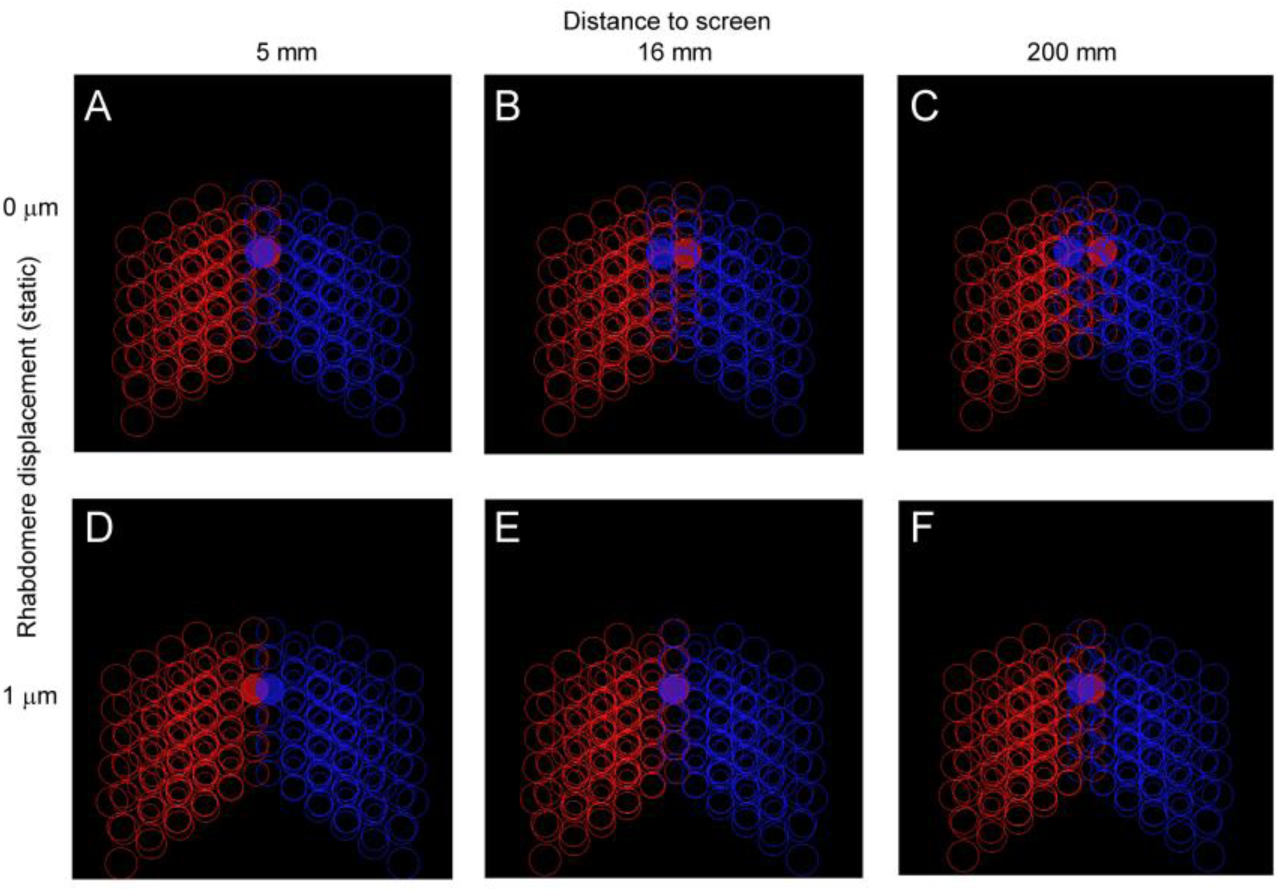
Forward-facing left and right eye rhabdomeres’ optical receptive field (RF) half-widths at different distances. (*A*) RF half-widths (disks) of the corresponding left and right eye R6 photoreceptors, as projected 5 mm away from the eyes with no added rhabdomere displacement. Red circles are the RFs of the neighboring photoreceptors in the left visual field, blue circles in the right visual field. (*B*) RF half-widths 16 mm away with no rhabdomere displacement. (*C*) RF half-widths 200 mm away with no rhabdomere displacement. (*D*) RF half-widths 5 mm away with an added 1 μm static rhabdomere displacement. (*E*) RF half-widths 16 mm away with an added 1 μm rhabdomere displacement. (*F*) RF half-widths 200 mm away with an added 1 μm static rhabdomere displacement.

Fig. S58 shows the rhabdomeres RFs (half-widths) and their photomechanically induced shifts, as projected at different virtual screen depths. Maximally 2.5 photoreceptor rows (∼11.5° binocular RF half-width overlap) are overlapping in stereo vision (Fig. S58*C*). The overlap was over-complete, as multiple RFs tiled up around the same position. With 1 μm movement, the overlap decreased to 1.5 rows (Fig. S58*F*). As the virtual screen was brought closer, the overlap became smaller. The crossing point of rows changed from roughly 5 mm away at rest (Fig. S58*A*) to 16 mm (Fig. S58*E*), when all rhabdomeres had moved 1 μm in 45° away direction. With dynamic stimulus (Movie S9), the degree of overlap changed over time, increasing the visual fields’ over-complete tiling.

##### V.11 Estimating distance using both eyes

Based upon its eyes’ *static* (immobile) anatomical dimensions, *Drosophila*’s estimated horizontal stereo vision field is small, containing maximally photoreceptor 2.5 rows. A similar constraint also arises in other compound eyes with a small overlapping field of view, such as bees (106). The small stereo vision field will make the conventional *static* stereo parallax - the left and right eye image disparity - distance estimator have a low depth resolution and a short depth range.

**Fig. S59.**
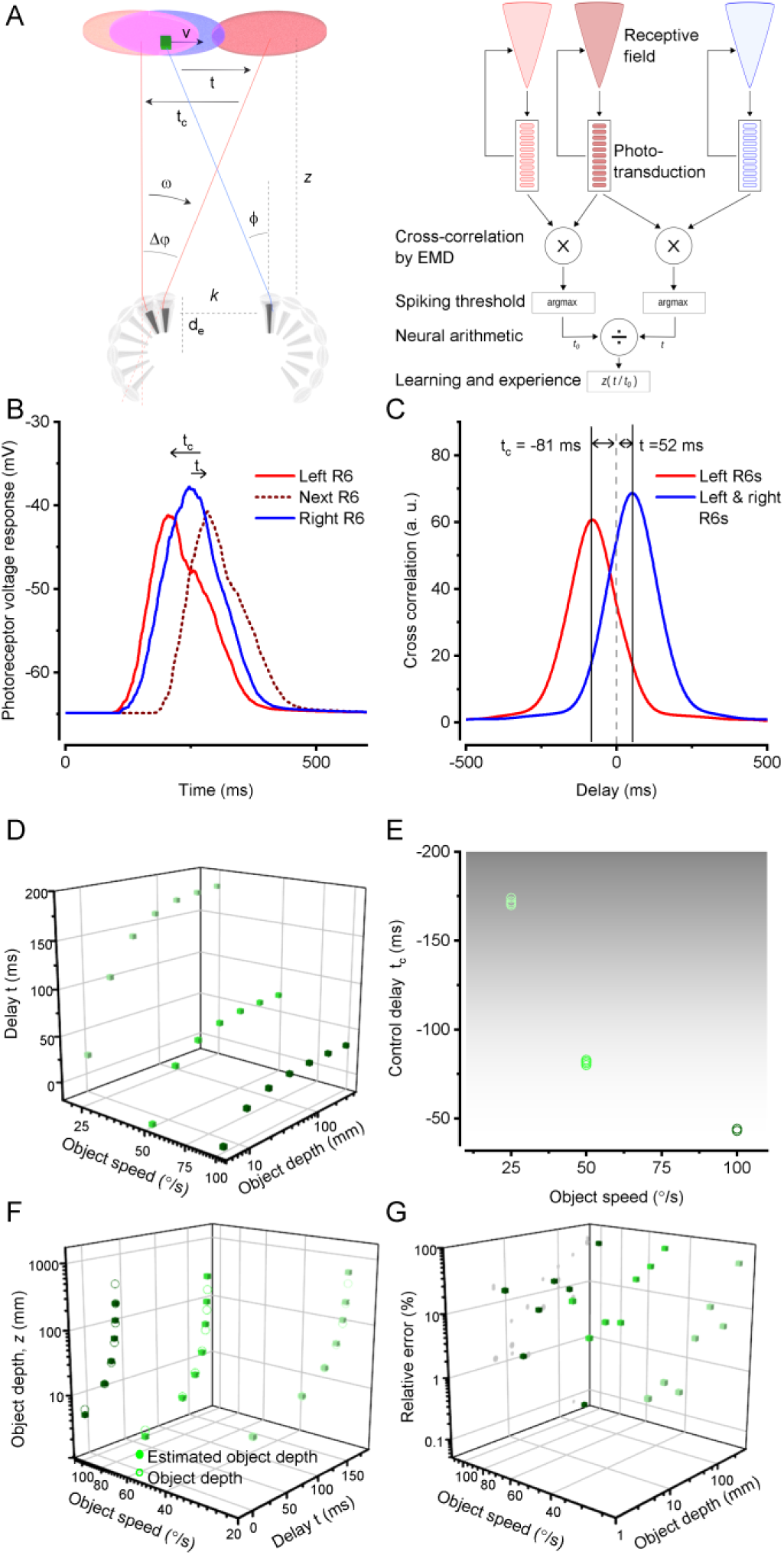
Object depth estimation from dynamic binocular R6-photoreceptor outputs. (*A*) Schematic of structural depth perception constraints in *Drosophila* compound eyes and the parameters and neural computations needed for calculating the object depth (*z*) in (*dynamic*) stereovision. Red indicates the left-eye and blue the right-eye receptive fields (RFs) and sampling. (*B*) Simulated voltage responses of three R6- photoreceptors at 25°C when a 1.7° x 1.7° object (a light-point) passes their overlapping RFs 50°/s, 25 mm away. These photoreceptors experienced maximal light input of 350,000 photons/s and were pre-adapted by 35,000 photons. (*C*) Cross-correlations calculated from the same responses in *B*. The red correlation is between the two left-eye R6-photoreceptor responses in the neighboring ommatidia. This pixel-wise correlation withstands the transmission in the optically/neurally superpositioned adjacent cartridges, from the photoreceptors to the lobula plate H1-neurons (107)). The blue correlation is calculated over the binocularly-shared RFs (overlapping pixels); between the corresponding right- and left-eye R6-photoreceptor responses. Such binocular correlations likely happen in the retinotopically organized neural cartridges of the lobula optic lobe, where the location-specific ipsi- and contralateral photoreceptor information is pooled (see Section V, below). The time delays occur between the maximum correlations (vertical lines) and the object crossing the left R6-photoreceptor’s RF center (vertical dashed line). See Movie S9 and S10. (*D*) Simulated delays, *t*, between the corresponding left and right-eye R6-photoreceptors (with overlapping RFs) when varying the object (1.7° x 1.7° light point) distance and speed. The screen resolution was 0.1°/pixel in all simulations. (*E*) Corresponding changes in the control delay, *t_c_*, when varying the object depth (7 different object depths taken from D) for the three different tested object speeds (25, 50, and 100°/s). The control delay is not dependent on the object depth, as all simulations with the same speed show little variance. (*F*) Comparison between the real object depth (open circles) and the corresponding model estimated object depth (disks); calculated from the estimated delays using Eq. 37. (*G*) The relative error in the model estimated object depth with respect to the real object depth. The error was calculated between (*D*) and (*F*).

Here, we suggest a new *dynamic* depth estimation method (Fig. S59) based on an object moving in the stereo field. An animal perceives motion when an object moves in its visual field and/or when itself or its eyes move (self-motion). For diurnal insects, praying mantis has been shown to estimate the distance to a moving object (45, 108). The distance between the left and right eye causes a depth (*z*) dependent delay between the corresponding left and right eye photoreceptor responses when their RFs collect light information from the same small visual area in space (Fig. S59*A*). The time difference (*t*), when an object moves with speed (*v*) over a distance (*s*) (*s* = *vt*), can be estimated from the delay in the peak cross-correlation between the photoreceptor responses. From the geometry between the corresponding left and right eye R6 photoreceptors (Fig. S59*A*), we have the following relationship:

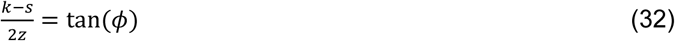

, where *k* = 440 µm is the distance between the eyes and *ϕ* is the photoreceptor convergence angle. With *s* = *vt* substitution, we obtain the object depth as:

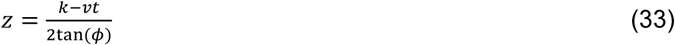

For determining the object speed, we used two neighboring photoreceptors (which also correspond to neurally superimposed neighboring LMC pixels in the lamina sampling matrix (107)) in the left eye as inputs to a simplified elementary motion detection circuit. In this scheme, we presume that the inputs from the corresponding binocular photoreceptor RFs (of the ipsi- and contralateral eyes) are brought together and compared in the lobula, in which connectivity indicates such circuits (see Section V, below). We calculated the delay *t*_*c*_between these photoreceptors using cross-correlation. Then the following equation is true:

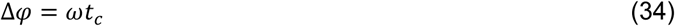

, where Δ𝜑 is the interommatidial angle (4.5°), and *ω* is the object’s angular speed:

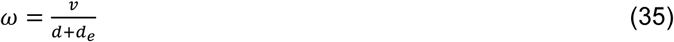

, where *d*_*e*_is the eye radius. Thus, the object speed is

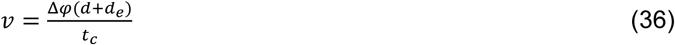

By substituting the speed in Eq. 33, the object depth is

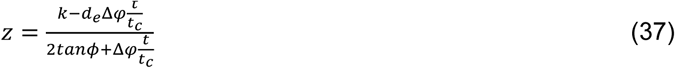

The convergence angle, *ϕ* , is dependent on rhabdomere movement (Fig. S58). The movement amplitude is dependent on object speed (Δ𝜑/*t*_*m*_). When an object moves through the field, the photoreceptor convergence-angle gets smaller (Fig. S58). The exponential function with negative exponent was found as the best fit for approaching the dependency:

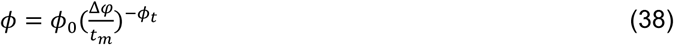

, where *ϕ*_0_ is the starting photoreceptor converge angle (5.8°) and the speed-dependent exponent *ϕ*_*t*_is 0.26565. As the object moved faster through the receptive field, the rhabdomere had greater movement amplitude (given the same stimulus light strength). Thus, the convergence angle was smaller.

We simulated three ommatidia (*e.g.,* two in the left eye and one in the right eye) in the stereo vision field. We calculated the delays: *t*_*c*_(or control delay) between two neighboring R6-photoreceptors in the left eye and *t* between the corresponding (and mirror-symmetrically aligned) R6-photoreceptors in the left and right eye (Fig. S59*A*). Fig. S59*B* shows an example of such R6 voltage responses for the three ommatidia, and Fig. S58C shows the cross-correlation curves based on Fig. S59*B* data. The delays *t*_*c*_and *t* are the delays with the maximal cross-correlations in respect to the left R6-photoreceptor’s RF-center (zero time-point). Fig. S59*D* shows how the delay *t* increases exponentially as a function of the object depth (the distance from the eyes). The *t*_*c*_-delay, which mimics that seen in the classic elementary motion detectors (107, 109, 110), shortens as a function of the object speed (Fig. S59*E*) with its slight variations coming from the noise (stochastic variations) generated by the QB summation in the four-parameter model. Fig. S59*F* shows the estimated depth by Eq. 37, and Fig. S59*G* shows the corresponding error.

Recordings from real neurons suggest that the depth estimation requires a change in the object’s visual distance, as shown for the praying mantis (45). If the object distance did not change, the suggested depth neurons would operate like many neurons along the motion detection pathway, responding most strongly to some preferred motion-direction yet showing less clear speed- or intensity-dependency.

##### V.12 Estimating *Drosophila*’s dynamic stereo vision range

Given that the *Drosophila* left and right eyes are ∼440 µm apart, the corresponding binocular photoreceptor pairs’ receptive fields (RFs) converge, move mirror-symmetrically and cross a certain distance in the front of the eyes, the accuracy of the dynamic stereoscopic depth estimation is limited. The absolute and relative depth error (Fig. S60 *A* and *B*) increased with the object distance because the angular differences become negligibly small far away. The relative error was, in general, >10% when the distances were >10 cm. The depth error can be explained by the hyperbolic shape (Fig. S59*D*, Eq. 37) of delay (*t*) combined with the phototransduction model’s noise (the four-parameter model’s stochastic variations in QB integration).

In Fig. S60 *C* and *D*, we tested a case where one eye’s rhabdomeres were stationary (immobile). The monocular photomechanical movements led to a significant depth overestimation because the delay (*t*) increased in these conditions. The object speed estimate became miss-calculated when one eye’s photomechanical movements were stopped.

**Fig. S60.**
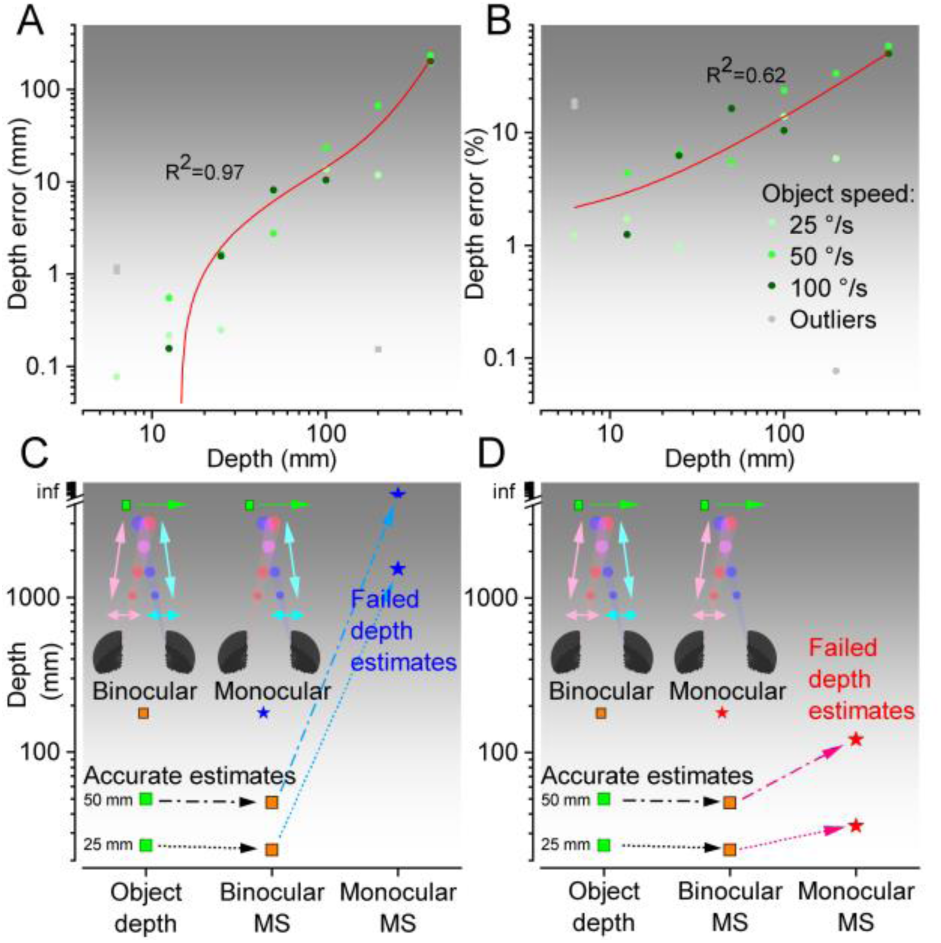
High-accuracy moving object depth estimation requires binocular mirror-symmetric microsaccades and decreases gradually with the increasing object depth. (*A*) The depth estimates’ absolute error and (*B*) relative error increase with the object (1.7° x 1.7° light-point) distance from the fly eyes; shown for three different object speeds. (*A* and *B*) The depth estimates were taken from Fig.54. Cubic and linear fits (red lines) show the estimates’ accuracy decreasing with the increasing object distances. Outliers (not used in the fits) are marked in light gray. (*C* and *D*) The real object depth and the model estimated object depths for the binocular mirror-symmetric and the monocular unidirectional microsaccadic sampling (while the other eye’s photoreceptors are motionless). The binocular sampling captured the object depth accurately. In striking contrast, the object depth estimation (Eq.35) failed drastically for the monocular unidirectional microsaccadic sampling. The model’s depth estimation error was more extensive (*C*; blue stars) when the object moved against the monocular microsaccadic RF direction (with the other eye’s photoreceptor RFs being stationary) than when moving with the RF motion (*D*; red stars). Corresponding behavioral 3D-object learning tests, based on these theoretical predictions, are shown in Section VI.6., Fig. S74.

##### V.14 Stimulus size and movement direction differentially shape R1-R7/8 outputs

We simulated how a collective photomechanical R1-R7/8 microsaccade in a single ommatidium affects each contributing photoreceptor’s power to resolve moving object details (Fig. S61). Because each R1-R6 rhabdomere has (i) different size and (ii-iii) lays a specific lateral distance off the ommatidium lens center-axis and the cone/pigment cell aperture’s outer rim, every R1-R6 samples light input during the microsaccade differently. Whilst, correspondingly, the stacked R7/R8 rhabdomeres move away from the lens center axis but not far enough for their responses to be shaped by the aperture’s light clipping.

**Fig. S61.**
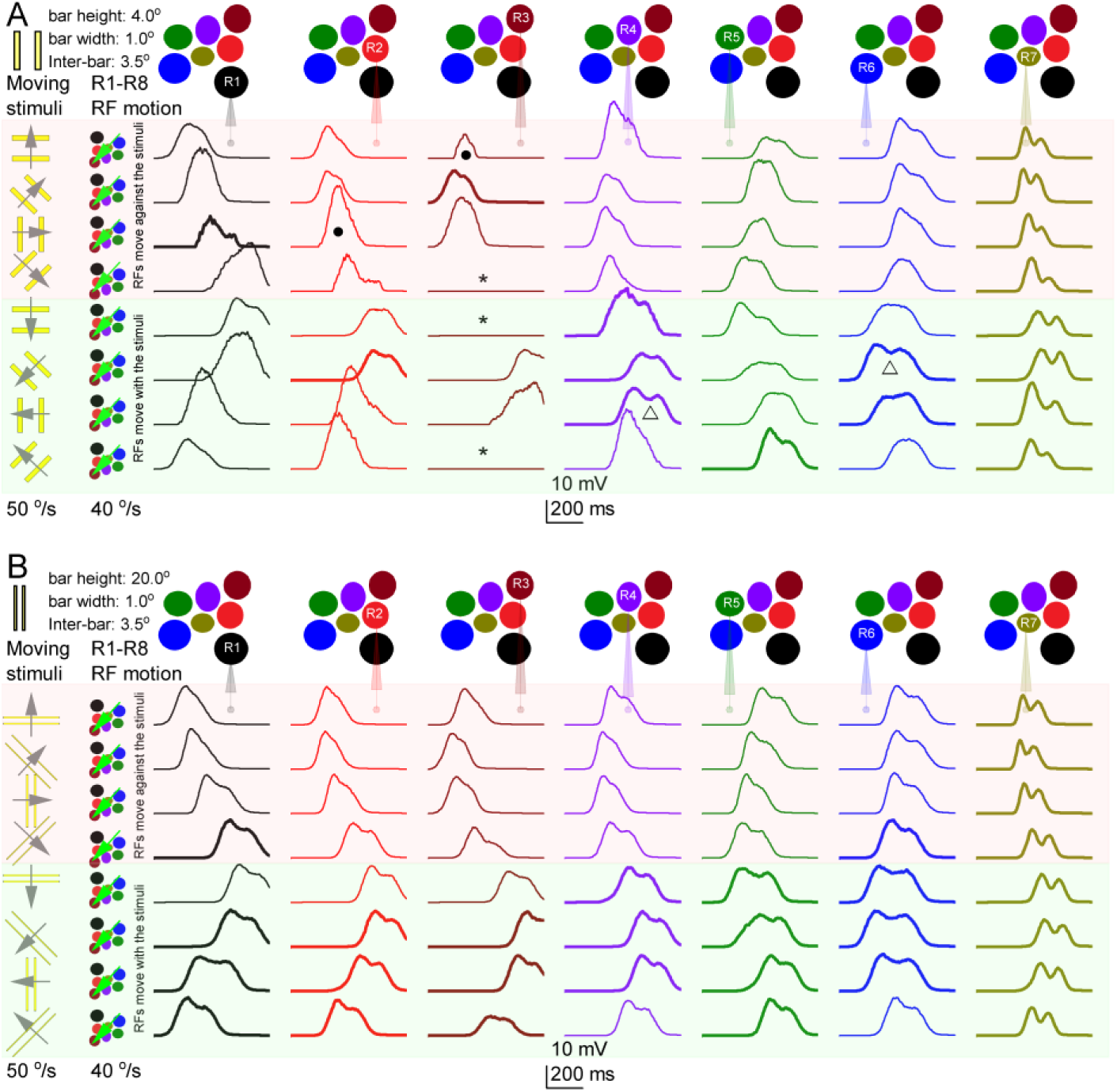
Stimulus size and direction. (*A*) R1-R7/8 photoreceptors best resolve the stimuli that move in the same direction as their receptive field (RF). Two short hyperacute bars (1.7° x 1.7°, 3.3° apart with screen resolution: 0.22°/px) cross R1-R7/R8s’ RFs at eight different directions, covering 360°. Light-red background highlights the R1-R7/R8 voltage responses to the stimuli that move against the microsaccadic RS motion. Light-green background groups the photoreceptor responses to the stimuli that move along the microsaccadic RS motion. The thick lines indicate the responses, in which the two bars caused two peaks, and the thin lines those with less clear or no peak separation. Above: colored disks indicate the R1-R7 rhabdomeres in a single ommatidium. Below: their voltage responses. Left: the ommatidium lens x/y-flipped R1-R7s’ RFs (colored disks) and their microsaccadic fast-phase movement direction regarding the given stimuli. *, · and Δ indicate the interesting cases where the combined microsaccadic movement (here, as initiated either by R1 or R5 photomechanics) could pull the R3, R2, R4, and R6 RFs either entirely or partially away from the stimulus movement path (explanations in the main text). The photoreceptors’ voltage responses were adjusted to have maximal light input 350k ph/s and pre-adaption of 35k ph 25°C. (*B*) Two long hyperacute bars (20° x 1.7°, 3.3° apart with screen resolution 0.22°/px) cross R1-R7s’ RFs at eight different directions, covering 360°. Note, these photoreceptor acuity simulations are deliberately conservative. We used dark-adapted acceptance angles (Table S6) without implementing the intracellular pupil mechanism, which would further improve photoreceptor resolvability in diurnal conditions. Nevertheless, we still obtained clear two-peaked responses to the moving hyperacute two bars.

To explore the consequences of these structural and positional dynamics in R1-R7/8 signaling, we tested how well each photoreceptor’s voltage response separates in time two short (Fig. S61*A*) or two long hyperacute (Fig. S61*B*) bars. These crossed the photoreceptors’ receptive fields (RFs) in a different direction at 50°/s. The simulations revealed that:

- R7/R8, with the narrowest rhabdomeres, resolve hyperacute moving stimuli better than R1-R6, which have wider rhabdomeres.
- Irrespective of the ommatidial photoreceptor position, the combined R1-R7/8 microsaccade enhances the resolution of objects that move broadly in the same direction (Fig. S61. light green background) as the R1-R7/8 RFs; in contrast to moving in the opposite direction (light red background). But even when opposite, vision is still hyperacute. Thus, the simulations predict (or, at least, are consistent with) the observed L2-terminal responses’ hyperacute orientation axes (*cf.*, Fig. 4*F* and Section IV.3., Fig. S41 *A* and *C*).
- For small hyperacute two bar stimuli, in which dimensions are less than the 4.5° interommatidial angle (Δφ) and are moving in a specific direction relatively slowly, the photomechanical activation of a single R1-R6 (as the stimulus first enters its RF) alone can cause a microsaccade that drags:

○ some of its neighbors’ RFs out of the stimulus light path (Fig. S61*A*, *), causing a null-signal (no light-induced depolarization).
○ some of its neighbors’ RFs only partially out of the stimulus light path (Fig. S61*A*, Δ), so that both bars cross their rhabdomere tips fractionally, causing a transient slit-effect. As if a slit appeared on the top of a rhabdomere to narrow its angular sensitivity, improving the resulting response’s two-bar resolution (superfine-signals).
○ some of its neighbors’ RFs temporarily out of the stimulus light path (Fig. S61*A*, l), so that the first bar is seen but the second one not.

These concurrent null-, single-peak- and superfine-signals may enhance visual objects’ spatiotemporal contrasts (dynamic edge-enhancement) at the lamina (the next optic neuropil), as the optically superimposed R1-R6 voltage signals from the seven neighboring ommatidia are pooled in synaptic transmission.

##### V.15 Theoretical predictions

Our new theory and its simulations - about the corresponding left and right eye photoreceptor arrays sampling depth-information in time - suggest that such *dynamic* sampling of image disparities gives three critical benefits for *Drosophila* vision in respect to using *static* (non-moving) photoreceptor arrays:

- *It enlarges the stereoscopic field of view*. In the static case, only 2.5 ommatidial rows of frontal (the left and right eyes’) photoreceptors could sample a tiny slice (∼11.5°) of the world horizontally in stereo. With mirror-symmetric photoreceptor microsaccades sweeping their receptive fields (RFs) side-to-side, this binocular slice (the stereoscopic horizontal field of view) expands at least to >30° (Fig. S14; for the experimental test and conformation, see Section II.1.ii, above.
- *It improves the retinal image resolution.* In the binocular region, one-half of the photoreceptors (say, ipsilateral) sample information while moving along with the object, and the other-half (contralateral photoreceptors) sample while moving against this motion. With microsaccades moving and narrowing the photoreceptors’ RFs, their responses encode much finer (hyperacute) object details (<1° (11); Fig. 4 and Fig. 6; Fig. S41) than what static photoreceptors ever could (∼4.5°, limited by the ommatidial spacing). However, crucially, during the dynamic sampling, these photoreceptor response time-differences also simultaneously carry the object depth information to the fly brain.
- *It improves visual image reliability and combats aliasing*. Because the ommatidial photoreceptor rhabdomeres are of different sizes and different distances from the center-axis (Fig. S47) and mechanically interconnected (Fig. S32 and Fig. S33, possibly by tip-links), their RFs tile the eyes’ binocular field over-completely (Fig. S58) and their voltage responses to moving visual objects vary (Fig. S61). This organization means that when pooling the photoreceptor responses in neural superposition, each LMC receives 6 (R1-R6) + 2 (R7/R8 – through gap-junction before the synapse (22, 105)) slightly differing samples of the same local visual object/event (Fig. S57). As we have shown before for the stochastic QB integration (40), such variability in spatiotemporal sampling improves the accuracy/reliability of the transmitted neural messages (*cf.* wisdom of the crowds (111, 112)) and combats aliasing (11, 59). See Fig. S67 and Section VII.3, below, for the behavioral test and confirmation.

In most seeing animals, because the photoreceptor sampling matrix and the underlying visual circuitry maps the world retinotopically, the spatial information of the neighboring visual points is already genetically encoded in the eye/brain network structure. Therefore, dynamic changes and correlative linking of the objects and their movements in the visual world can be efficiently replayed/represented as temporal differences in the networks’ phasic neural responses.

Interestingly, our theory further predicts that *Drosophila* would have “short-sighted” stereo vision, seeing close-by objects in higher resolution than those further away from them (Fig. S59 and Fig. S60). In Section VII., below, we test and verify this prediction.

##### V.16 Estimating responses to hyperacuity stimuli with classic stationary eye models

We estimated how well a hypothetical *Drosophila*, having *static eye structures* with *sampling limited by interommatidial angles* (as is the dominant/classic view in the literature), could differentiate hyperacute contrast differences between two neighboring photoreceptors’ receptive fields (single non-overlapping “pixels,” with 5.4° half-width) (Fig. 6G). Both test images contained 1° black-and-white stripes, but one also had a single 0.98° black dot in the center. The eye’s distance to the screen was the same as in the flight simulator experiments: 25 mm (Fig. 6), and the screen resolution was 0.01°/pixel. The black intensity was half of the white with maximal photon flux: 500,000 photons/s at 25° C. The resulting intensity difference (transient contrast change) between the two images is ∼1.6%. We simulated photoreceptor responses to a 100 ms negative light pulse, comparable to the image intensity of the black dot in the background and the stripes images alone, using the four-parameter photoreceptor model to generate the light current. From the corresponding light currents, we simulated the voltage responses. These simulations made it clear that it would be practically impossible for a *static* pixelated *Drosophila* eye to neurally differentiate the 0.98° black dot response from the black-and-white stripe background, which was smaller than the simulation noise.

The scripts to simulate and analyze *Drosophila* ommatidial optics are downloadable from: https://github.com/JuusolaLab/Hyperacute_Stereopsis_paper/tree/main/OpticalSimulations

#### VI Anatomical Rationale

In the insect brain, the lobula complex neuropile pools visual information from ipsi- and contralateral eyes (43–45). In the praying mantis, the *coCOM* neuron has been shown to carry information relevant to stereopsis bilaterally (45). The LC14 neurons are thought to be homologous to the mantis *coCOM* neurons (6, 45). In *Drosophila*, they project from one lobula (and the medulla for LC14b neurons (113, 114) to the lobula on the contralateral side. As such, they represent one possible class of neurons that integrates visual information across hemispheres and allows stereopsis to occur.

To assess the pattern of projection (i.e., do the neurons project from one area of the lobula to the same area on the other side), we selected MCFO images from the flylight database that were identified in the NeuronBridge (115) tool as expressing in at least one LC14 neuron. After manual quality control to look for datasets containing low misexpression, we collated the data (Fig. S62).

The LC14 neurons appear to project from one area of the lobula to an approximately similar area on the contralateral side, although this cannot be ascertained to a fine degree. Since the lobula is organized retinotopically (e.g. (116)), this suggests that the neurons are integrating information from roughly the same regions of visual space in each eye. Hence, there is a plausible anatomical reason to think that visual information can travel across the *Drosophila* brain, although we cannot at this stage conclude that these specific neurons carry out this role.

**Fig. S62.**
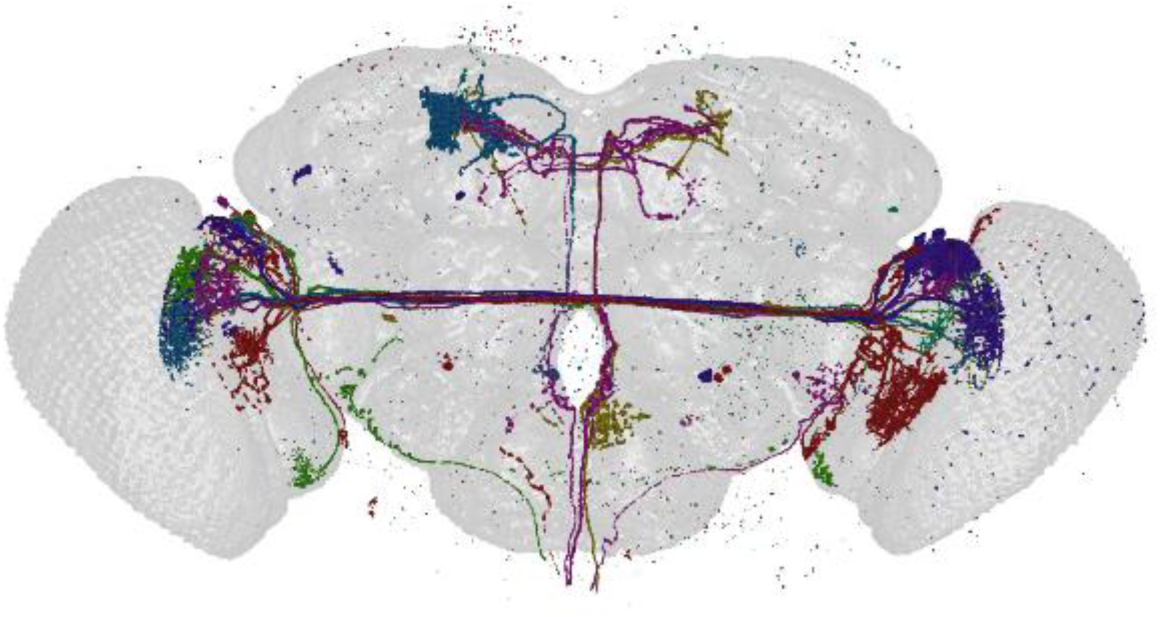
LC14 neurons and other similar lobula neurons may participate in processing stereoscopic visual information. LC14 neurons were identified in Neuprint and cross-linked with gal4 expression in NeuronBridge. Matching lines were then taken from the flylight Generation 1 MCFO Collection, separated by a channel, thresholded, and collated. Lines used: R12F03, VT047848, VT062633. Each color represents a different confocal stack. The neurons are labelled sparsely; however, in some instances, multiple LC14 neurons are labelled in each stack. The image resolution is limited by the resolution of the original confocal stacks they were taken from. Neurons appear to project from roughly the same area of the lobula on each side.

#### VII Flight simulator experiments

**Overview**

This section describes flight simulator experiments to measure (i) *Drosophila* optomotor behavior from hyperacute to coarse 2D stimuli, (ii) visual salience to hyperacute 2D and 3D stimuli, and (iii) associative avoidance learning of these stimuli. It gives central background information and additional supporting evidence for the results presented in the main paper, including:

- ptomotor responses are stronger to the closer hyperacute rotating scenes of the same angular resolution (2.5 vs. 5.0 cm away from the fly eyes), indicating short-sighted *Drosophila* vision/stereopsis (*i.e.,* the flies are seeing nearby objects in higher resolution). These results are consistent with the theoretical predictions; see Sections V.12. and V.15. above (Fig. 5 and Fig. S60).
- well-known optomotor response reversal to a rotating ∼7° stripe pattern originates from the mirror-symmetric left and right eye photoreceptor microsaccades, whereby one eye’s microsaccades move with, and the other eye’s against the screen rotation, causing a neural imbalance in the optic flow perception.
- optomotor response reversal is velocity-dependent - occurs when the field rotation speed approaches the eyes’ microsaccade speed (∼40-50°/s) - and can be stopped by painting one eye black, eliminating the eyes’ optic flow imbalance driving the behavior. Thus, the optomotor response reversal does not result from spatial sampling aliasing (the eyes’ ommatidial photoreceptor spacing) but perceptual aliasing. These results pair with the theoretical predictions; see Section V.15. above.
- *Drosophila* has super-resolution stereoscopic vision:

○ It finds hyperacute 3D objects more salient than the same area/contrast 2D objects.
○ It needs two eyes to see hyperacute 3D objects.
○ It needs binocular mirror-symmetric microsaccades to see 3D objects.
○ It uses both R1-R6 and R7/8 photoreceptors cells for stereopsis.

##### VII.1 *In vivo Drosophila* preparation

*Drosophila* were raised on molasses-based food at 25°C on a 12-h light/12-h dark cycle. 2- to 9-day-old female flies (vast majority 4-day-old flies) were briefly cold-anesthetized (on a bespoke Peltier cooling/preparation-making stage) for fixing a small copper-wire hook (0.06 mm Æ) with UV-light-curable glue (Loctite) between the head and thorax (11).

##### VII.2 *Drosophila* flight simulator system

A tethered flying fly was connected to the torque-meter by a small clamp holding the copper-wire hook, which fixed its head in a rigid position and orientation while transducing the fly’s yaw torque (left and right rotation attempts) into a voltage signal (Fig. S63 *A* and *B*). The fly was positioned in the center of a hollow plastic transparent cylinder (cup - its flight arena). This cup displayed high-resolution visual stimuli: black laser-printed patterns (Sharp MX-5141 printer; 1,200 × 1,200 dpi resolution) and/or small 3D objects attached on white paper, surrounding the fly’s long axis. We either used a small cup (inner Ø 50mm; Fig. S63*C*) or a large cup (Ø 100 mm; Fig. S63*D*), which kept the stimuli at 25 or 50 mm from the fly eye, respectively. In either case, the cups were rotated around the vertical axis by a stepping-motor, moving the stimuli free of flashing or aliasing. Outside, the cups faced a layer of surrounding diffusers. Behind them was a ring-shaped flicker-free light tube (special full-band: 350-900 nm), which uniformly illuminated the stimuli with no visible or only negligible shadows. Although perceptually bright, this background intensity was, nevertheless, 0.5-1.5 log units less than the maximum used in the L2-neuron Ca^2+^-imaging recordings (Fig. 4) and previous intracellular recordings (11); measured by Hamamatsu Mini C10082CAH spectrometer (Japan). Notably, here, our “drum stimuli” *were not testing visual parameter changes affecting the fly vision during translation*, such as angular size, spatial wavelength composition, or distance-dependent velocity.

**Fig. S63.**
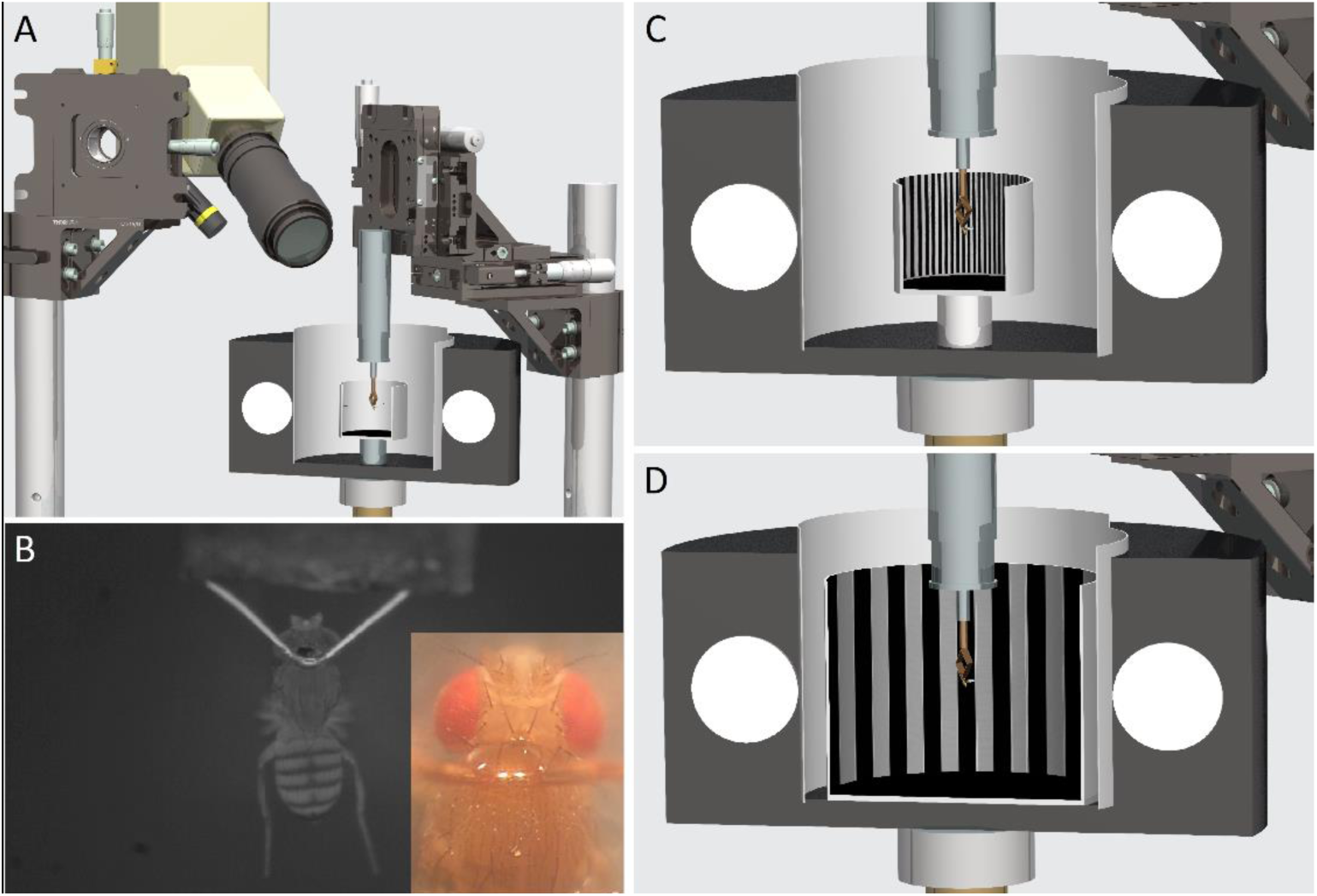
Schematic cross-sections of the *Drosophila* flight simulator system. (*A*) A *Drosophila* is tethered from a torque-meter, flying in the center of a panoramic arena (a cup), which is back-illuminated through layers of diffusers by a ring-shaped high-intensity lamp (white discs). The fly’s yaw-torque signal controls in a closed-loop the panoramic scene it faces in the learning experiments. When the fly viewed the test stimulus, the infra-red laser (yellow stripe) was activated automatically to condition the test stimulus with heat punishment (unconditioned stimulus, US) to the fly head. During the experiment, the fly’s behavior could be further recorded with a macro lens video camera (*B*). (*B*) A small holder was used to clamp the tethered flying fly from a copper-wire hook (glued between the head and thorax), connecting it to the torque meter. (*C* and *D*) To test how a fly’s visual perception depends on its distance to the stimulus, we used both a small cup (*C*) and a large cup (*D*) visual arenas. The visual patterns/objects in the small cup were 25 mm from the fly eyes; they were 50 mm from the fly eyes in the big cup.

The flight simulator system was mounted on a vibration isolation table inside a black-painted and light-proofed steel-walled Faraday cage, with a black roller curtain at the front to block any outside light (potential visual cues) affecting the experiments.

##### VII.3 Optomotor behavior (open loop)

A fly saw a continuous panoramic black-and-white stripe-scene of a specific angular resolution on the given cup’s inner wall. After 1 s of viewing the still scene, the scene was spun to the right (clockwise) by a stepping motor for 2 s, stopped for 2 s, before rotating to the left (counterclockwise) for 2 s, and stopped again for 1

This 8 s stimulus was repeated 10-25 times, and each trial, together with the fly’s yaw torque responses, was sampled at 1 kHz and stored in a hard drive for later analysis. Typically, a tethered flying fly attempts to follow the moving panorama, generating optomotor responses (yaw rotation signals), the strength of which is thought to reflect the strength of its motion perception in respect to the used stimulus parameters.

If a fly stopped flying during trials, it would be encouraged to start flying again immediately with puffs of air or provided with a paper ball soaked with 30% sucrose solution. Tests were stopped if flies stopped flying >5 times during a 2 m period.

***Testing hyperacute vision distance range***

Owing to the left and right eyes’ mirror-symmetric photomechanical photoreceptor microsaccades (Figs 1-3; see Sections II. and IV., above) and the resulting phase differences in the binocular receptive field dynamics (Figs 4-5; see Section V., above), our theory predicts that a fly should see the nearby world in hyperacute 3D. But it should see the more distant world in blurry 2D.

- *Notably, such dynamics would offer a Drosophila a way to sense object size.* For instance, a small nearby object, another *Drosophila* - seen frontally by the left and right eye, would generate a stereoscopic pair of separate images (with phasic time differences in their neural representations), signaling no danger. But a distant object of the same angular size would have little or no such stereo-neural cues. Therefore, it could be perceived as further away, signaling that this object is bigger and potentially dangerous.

**Fig. S64.**
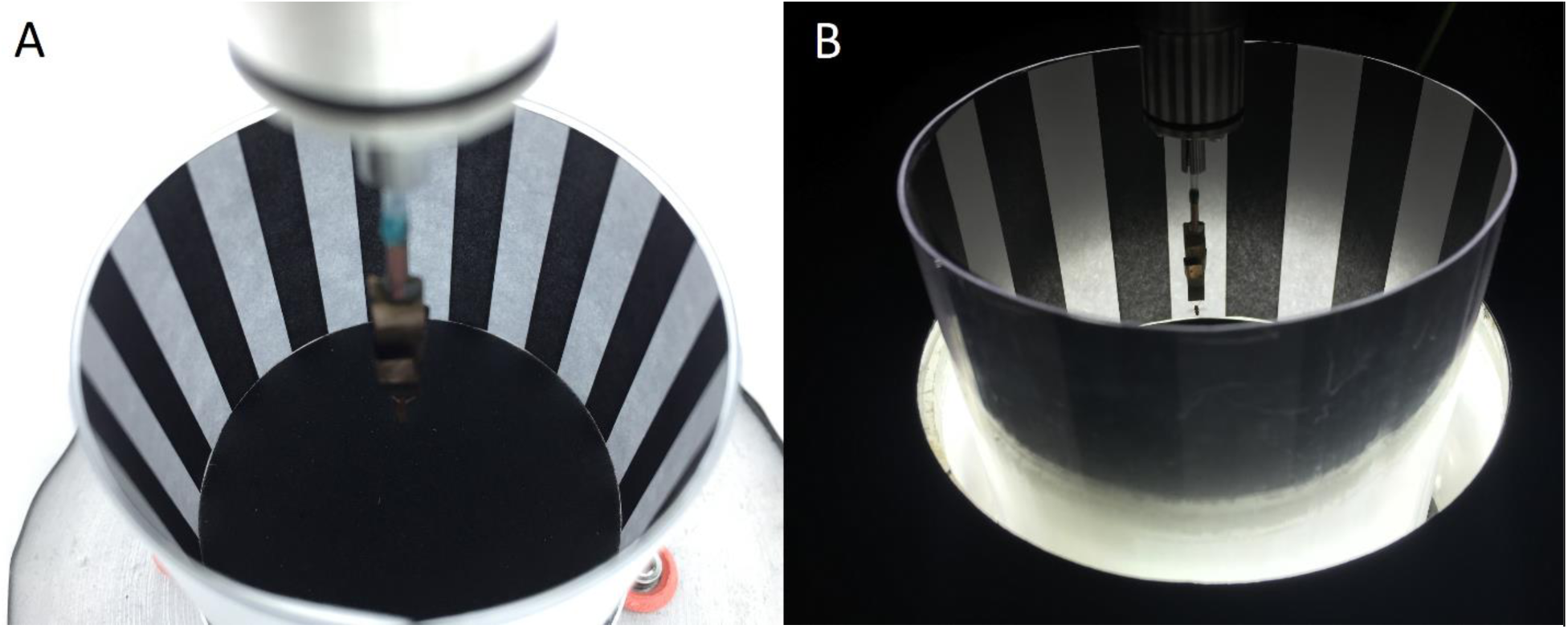
Testing optomotor behavior with two different size arenas: the small and the big cup. Their black stripe patterns were printed on white paper so that the resulting angular stripe widths were similar, as seen by the tested flies. (*A*) A tethered *Drosophila* viewing the stripe patterns in the small cup. (*B*) A tethered *Drosophila* viewing the stripe patterns in the large cup.

To test whether the flies’ visual acuity, as defined by their optomotor response strength, depended on how far the presented stimulus was (*i.e.,* the distance from the fly eyes to the stripe scene), we used both the small and large cup (Fig. S64). For the small cup, its stripe patterns (25 mm from the eyes; Ø = 50 mm) were within *Drosophila*’s estimated stereo vision range (0-30 mm), whereas for the large cup, its patterns (50 mm from the eyes; Ø = 100 mm) should lie closer to the outer edge of this range. The fixed stimulus parameters for moving stripe scenes, as shown in the figures, were: azimuth ±360°; elevation ±45° (small cylinder) or ±40° (large cylinder); contrast, 1.0, as seen by the fly. The large cylinder’s top was less illuminated because it extended further away from the surrounding ring-light (Fig. S64*B*). However, as we kept each fly at the same vertical position regarding the ring-light, they experienced similar light intensity changes with both the cups.

Notice that because the big and small cup’s angular speed and spatial wavelength were made as close as possible identical, the temporal frequency (i.e., the ratio between angular velocity and spatial wavelength of the pattern) was constant for each paired experiment.

***Optomotor tests with the small cup.*** Black-and-white stripe-scenes (spectral full-width: 380-900 nm) of five different spatial resolutions (wavelength [bar-to-bar-distance]: 2.34° [1.17°], 4.68° [2.34°], 6.43° [3.21°], 12.86° [6.43°] and 25.71° [12.35°]) were rotated at 45 and 300 °/s (Fig. S65, *A* to *D*). As the *light* control, to examine whether airflow or some hidden features in the stimulus panorama affected optomotor responses, we used either white paper or a separate white diffuser cup of the same size or both, rotated at the same two speeds. As the *dark* control, the same flies’ optomotor responses were recorded to the scene rotations in complete darkness. The *light* and *dark* controls evoked either no or only minimal torque responses.

*Optomotor tests with the large cup*. We tested the flies’ torque responses to 2.43° [1.215°], 4.86° [2.43°], 6.92° [3.46°], 13.84° [6.92°] and 27.69° [13.845°] wavelength [bar-to-bar-distance] black-and-white stripe-scenes, rotated at 45 and 300°/s (Fig. S65, *E* to *H*). Thus, to a tethered fly, the black-and-white bars in the corresponding large and small cup stripe-scenes had broadly similar angular widths, but these images were now twice as far from its eyes. *Light* and *dark* controls were adapted for the large cup, as explained above. Again, these control stimuli evoked either no or only minimal torque responses.

**Fig. S65.**
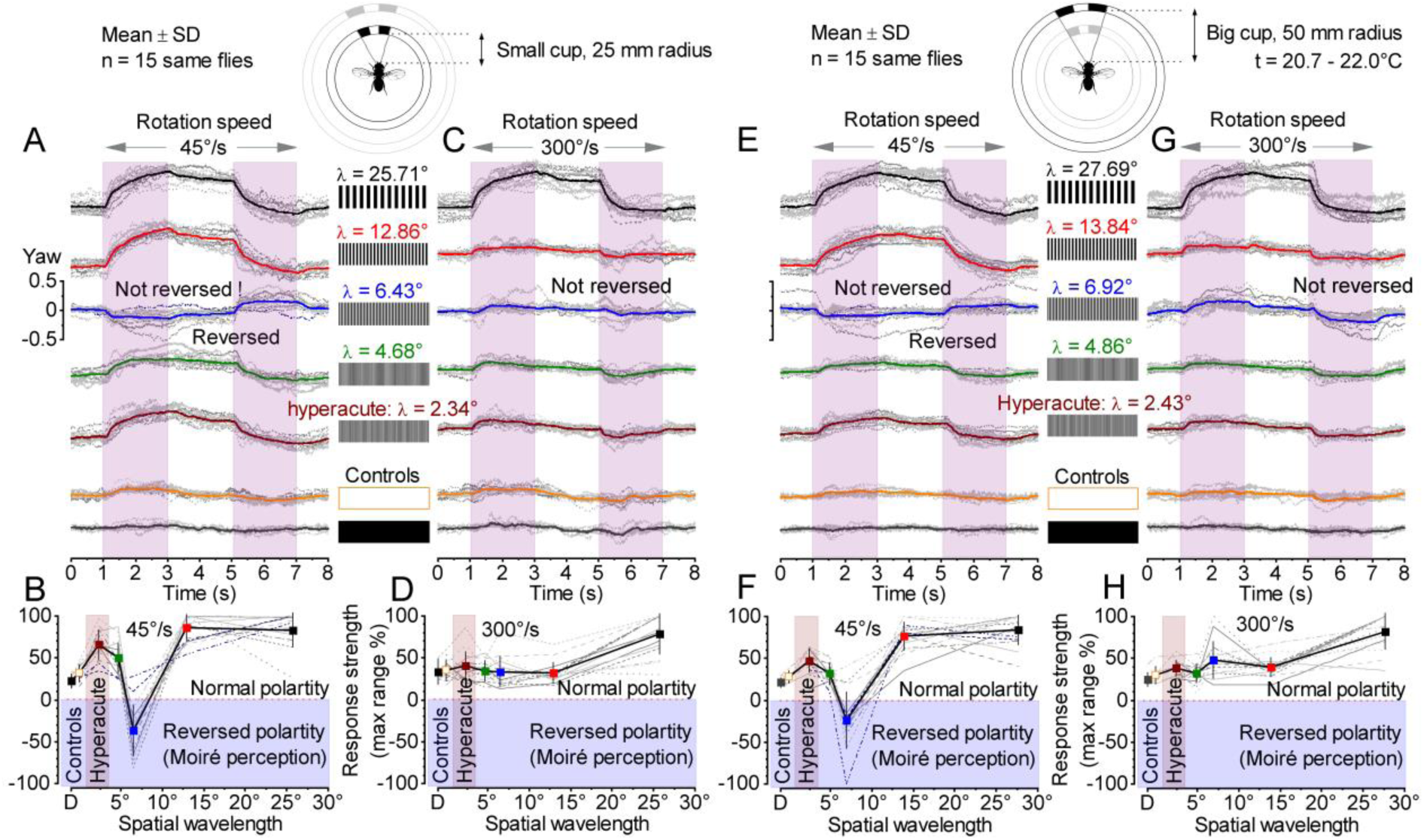
Optomotor responses to clock-wise and counter-clockwise rotated visual stimuli of different wavelengths and velocities; presented either 25 mm (small cup: *A* to *D*) or 50 mm (large cup: *E* to *H*) from the *Drosophila* eyes. (*A* and *C*) Small cup: Each fly was tested with five black-and-white stripe patterns of different wavelengths and two control stimuli (white-paper and dark), rotated at two different velocities (*A*: 45°/s and *C*: 300°/s). In each case, the specific stimulus was presented 10-times. For each fly, the resulting optomotor responses (yaw-torque) were first averaged and then scaled by normalizing them with their largest average response for the most sensitive stimulus. The thin traces show these stimulus strength-normalized averages for each fly and the thick traces their corresponding population means (n = 15 flies). (*B* and *D*) Small cup: Optomotor response strength depends on the stimulus wavelength and velocity (*B*: 45°/s and *D*: 300°/s rotations). Overall, *Drosophila* tracked most vigorously both coarse (black, 25.71°) and hyperacute (wine, 2.34°) stimulus rotations. 13 out of 15 *Drosophila*’s optomotor responses reversed during 6.43° (*B*, blue) black-and-white stripe rotations at 45°/s. But this reversal never occurred during 300°/s (*D*, blue) rotations. (*E* and *G*) Big cup: Each fly was tested with five black-and-white stripe patterns of different wavelengths and two control stimuli (white-paper and dark), rotated at two different velocities (*E*: 45°/s and *G*: 300°/s), with each of these stimuli presented 10-times. The resulting optomotor responses were averaged and normalized for every fly, as in *A* and *C*. The thin traces show these averages, and the thick traces the population means (n = 15 flies). (*F* and *H*) Big cup: Optomotor response strength depends on the stimulus wavelength and velocity (*F*: 45°/s and *H*: 300°/s rotations). Overall, *Drosophila* tracked most vigorously coarse (black, 27.0°) and hyperacute (wine, 2.43°) stimulus rotations. 12 out of 15 *Drosophila*’s optomotor responses reversed during 6.92° (*F*, blue) black-and-white stripe rotations at 45°/s, but this reversal never occurred during 300°/s rotations (*H*), with some flies’ responses being unexpectedly strong (blue, 6.92°)

The optomotor responses of individual flies to repeated field rotations vary in strength and repeatability (Fig. S65, thin traces), but their visual performance to different spatial resolution stripe scenes is different. These differences can be quantified by measuring the mean torque response of a single fly to stimulus repetitions and by averaging the mean responses of the many flies of the same stripe scene resolution (thick traces). This procedure reduces noise and non-systematic (arbitrary) trends of single experiments, revealing the underlying response strength and optomotor behavior characteristics. Characteristically, a fly’s torque response returns gradually to baseline after the optomotor stimulus stops, but this can take seconds, varying with individual flies (11, 22). Accordingly, in our experiments, which comprise only brief 2-s-long inter-stimulus-intervals, the torque responses typically recovered only fractionally (10-70%) during these still periods toward the baseline. Therefore, for comparing the optomotor behavior at different stripe scene resolutions, we used the maximum range (or peak-to-peak) of the torque response evoked by the combined leftward and rightward field rotation stimulus.

Consistent with our previous results (11), the optomotor responses to the small cup’s hyperacute (wavelength: 2.34°), fine (4.68°) or coarse (12.86° and 25.71°) stripe-scenes (at 25 mm from the eyes), irrespective of the tested rotation speeds, showed no aliasing, which otherwise would have been perceived as slowed down image rotation, eventually reversing to the opposite direction (the reverse rotation effect). However, in clear contrast, we found that ∼80-87% of the flies showed response reversing (51) to a 6.43° stripe-scene when rotated at 45 °/s (Fig. S65 *C* and *G*, thin traces), indicating that with these stimulus settings, the flies likely *perceive* Moiré-like visual effects. Yet notably, with high rotation speeds, such as 300 °/s, the optomotor responses to the same 6.43° stripe-scene did not reverse but normally followed the rotations (Fig. S65 *D* and *H*, blue squares). Moreover, ∼13-20% of the flies never reversed their optomotor responses to any test stimuli. Such a fly- and velocity-dependent selective motion perception reversal (for a narrow stimulus wavelength range only) suggests that this behavior unlikely resulted from eye size differences - the average inter-ommatidial angle would be the same for small or large compound eyes - or spatial sampling-aliasing attributable to 3.5-4.5° interommatidial angles.

Crucially, the flies generated stronger optomotor responses to the hyperacute stripe patterns of similar angular widths when closer to their eyes (*cf.* Fig. S65 *C* and *G*, wine squares; Fig. S66). This finding is consistent with our theory about how the mirror-symmetric left and right eye microsaccades sample 3D- information (see Sections V.13. and V.15., above), predicting that *Drosophila* has short-sighted (stereo) vision.

**Fig. S66.**
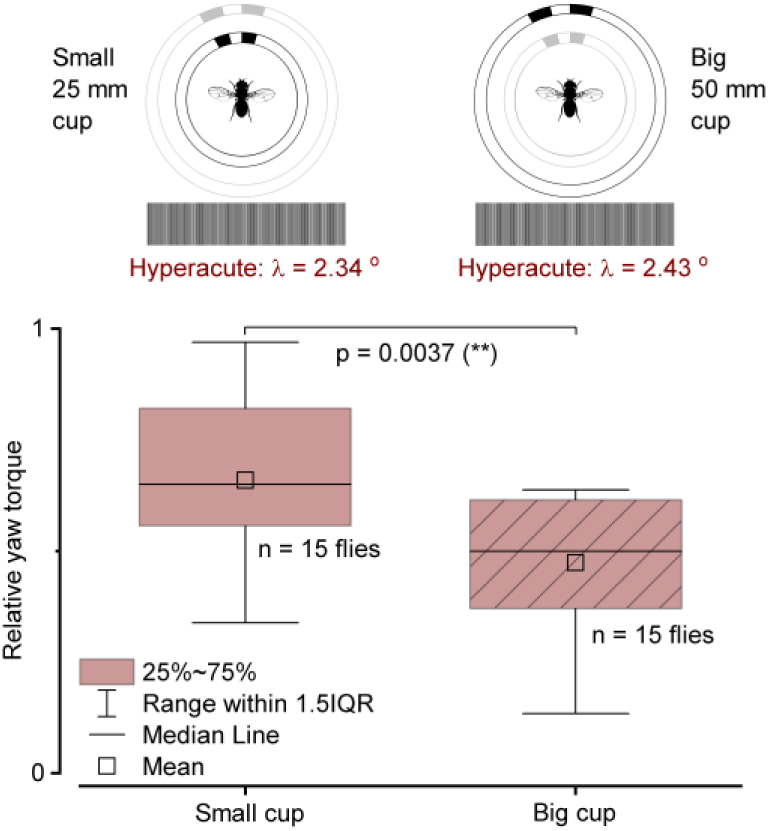
Optomotor responses to hyperacute black- and-white stripe pattern rotation, with similar wavelengths and speeds, are stronger when the stimuli are closer to the fly eyes.

***Mirror-symmetric photomechanical photoreceptor contractions reverse optomotor perception to slowly rotating (45* °*/s) ∼6.5-7.5*° *vertical stripe pattern scenes.*** As we and others have shown earlier, *Drosophila* eyes’ sampling matrixes are not fully orderly. R1-R7/8 rhabdomere sizes and positional off-sets differ (11), their optically superimposed microsaccades track local light intensity changes (see Section II, above), R7/8 pigmentation is stochastically distributed over the majority of the eye surface (41), and the photoreceptor’s connectivity matrix is asymmetric (11, 30). Therefore, we can be confident that selective pressures have tailored the eyes’ neural images at the level of photoreceptors and first interneurons to be *free of sampling aliasing* (11, 117–119). Nevertheless, in certain unusual stimulus conditions, which the flies would not normally encounter in the natural environment, mirror-symmetric left and right eye photoreceptor microsaccades can lead to imbalanced image cross-correlation later at the motion detection computations, causing *perceptual aliasing* (120).

Theoretically, the dominant contributing factor for the observed *perceptual aliasing* to the 45°/s rotating ∼7° stripe cup should come from the left and right eye’s mirror-symmetric photoreceptor microsaccades, which themselves travel 40-50°/s. For one eye’s photoreceptors, their microsaccadic speed and direction would broadly match the stimulus rotation, causing their RFs to rapidly lock to the moving stripes. Thus in the retinal mosaic, those neurally superimposed near-neighbor LMC pixels paired 6-8° apart for retinotopic depth/motion detection (see Section V.12., above)(107) would point to similar stripe patterns, seeing little stimulus change; signaling little or “no-movement.” At the same moment, the other eye’s photoreceptor microsaccades would make their RFs travel against the rotation, seeing “double-fast” moving stripes flashing by. For the fly brain, this *perceptual* “dynamic imbalance” between the left and right eye inputs may appear as if the stimulus rotated in the opposite direction, triggering an optomotor response against the actual stimulus rotation. Alternatively, the fly may perceive the stimulus approaching one side and thus turn away from it to re-center itself and balance the optic flow (121). Of course, the eyes’ input imbalance would reverse during their photoreceptor microsaccades’ slow-phase, which moves the RFs in the opposite directions (Fig. 3F; see also Section II.6., above). But as the refractory recovery slow-phase motion is weaker than the transient fast-phase, it may impact the fly perception less.

**Fig. S67.**
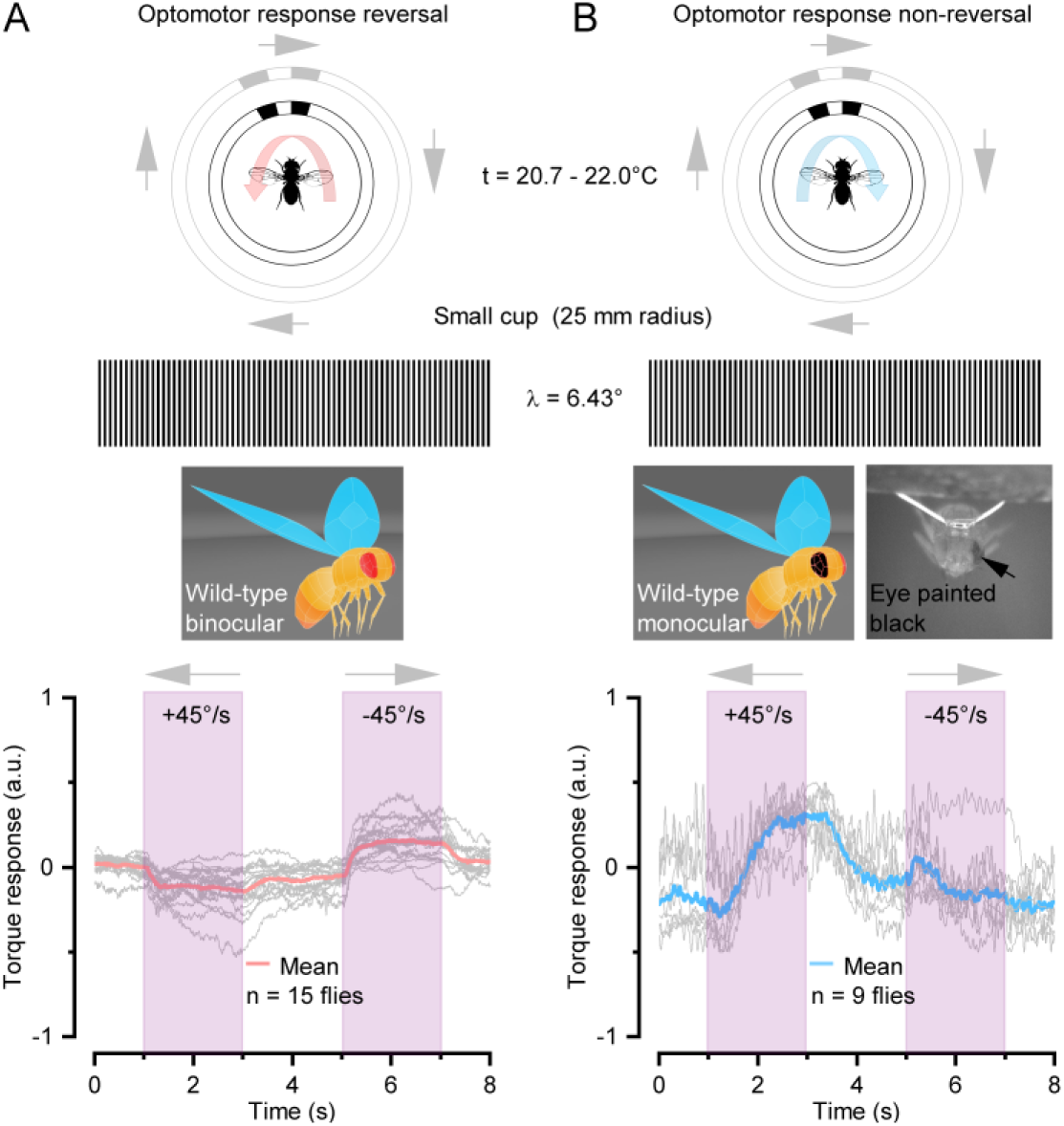
Monocular flies - having one eye painted black - do not reverse their optomotor responses during 6.43°-wavelength black-and-white stripe scene rotation. (*A*) Most binocular wild-type flies show an optomotor response reversal (thick red trace) to 45°/s rotating 6.43°-stripe stimulus. (*B*) In clear contrast, monocular wild-type flies follow 45°/s rotating 6.43°- stripe stimulus without reversing their optomotor response direction (thick blue trace). Together these (*A*-*B*) results demonstrate that optomotor response reversal results from the dynamic sampling imbalance between the left and right eyes’ optic flow inputs - as predicted by our theory; and not from arbitrary static spatial aliasing in the photoreceptor matrix. Each thin gray trace is a mean of 15 to 22 trials.

To test this concept directly, we painted *Drosophila*’s one eye black (left or right; see Section VII.6., below, for paint details), eliminating its counter-rotating microsaccadic RF movements affecting the optomotor behavior, and repeated the experiments (Fig. S67). As predicted, we now found that the monocular black-eye-flies turned along with the 45°/s rotating 6.43° stripe stimulus, in contrast to the normal two-eyed flies, which in most cases turned against it. In total numbers, 7/9 black-eye-flies consistently followed the rotating stimulus direction (100%, in every trial), whereas 2/9 of them followed the stimulus ∼90% of the trials (turning against the rotation only ∼10% of the time). Such slight hesitancy (or variation) might have resulted from these two flies’ painted-eyes perhaps being less-perfectly light-proof. Overall, this experiment demonstrated that the *reverse optomotor turns, as tested in a conventional flight simulator system, result from perceptual aliasing; and not from sampling aliasing* in the photoreceptor matrix (51). Nevertheless, the full neural mechanism and dynamics behind such *perceptual aliasing* are likely to be more complicated and may involve other factors and even other senses.

##### VII.4 Studying stereopsis using the *Drosophila* flight simulator system

In this study, we used real object depth rather than prisms or colored filters (as in the praying mantis work (45, 108)) or mirrors or goggles (as in the mammal/bird work (5, 7, 122)) to test the visual stereopsis behavior. Could a *Drosophila* use monocular or other cues in the flight simulator experiments, such as motion parallax or air currents, to distinguish the hyperacute 3D objects, accounting for the visual salience (Fig. 6A-J) and learning results (Fig. 6K-P) shown in the main paper?

**Fig. S68.**
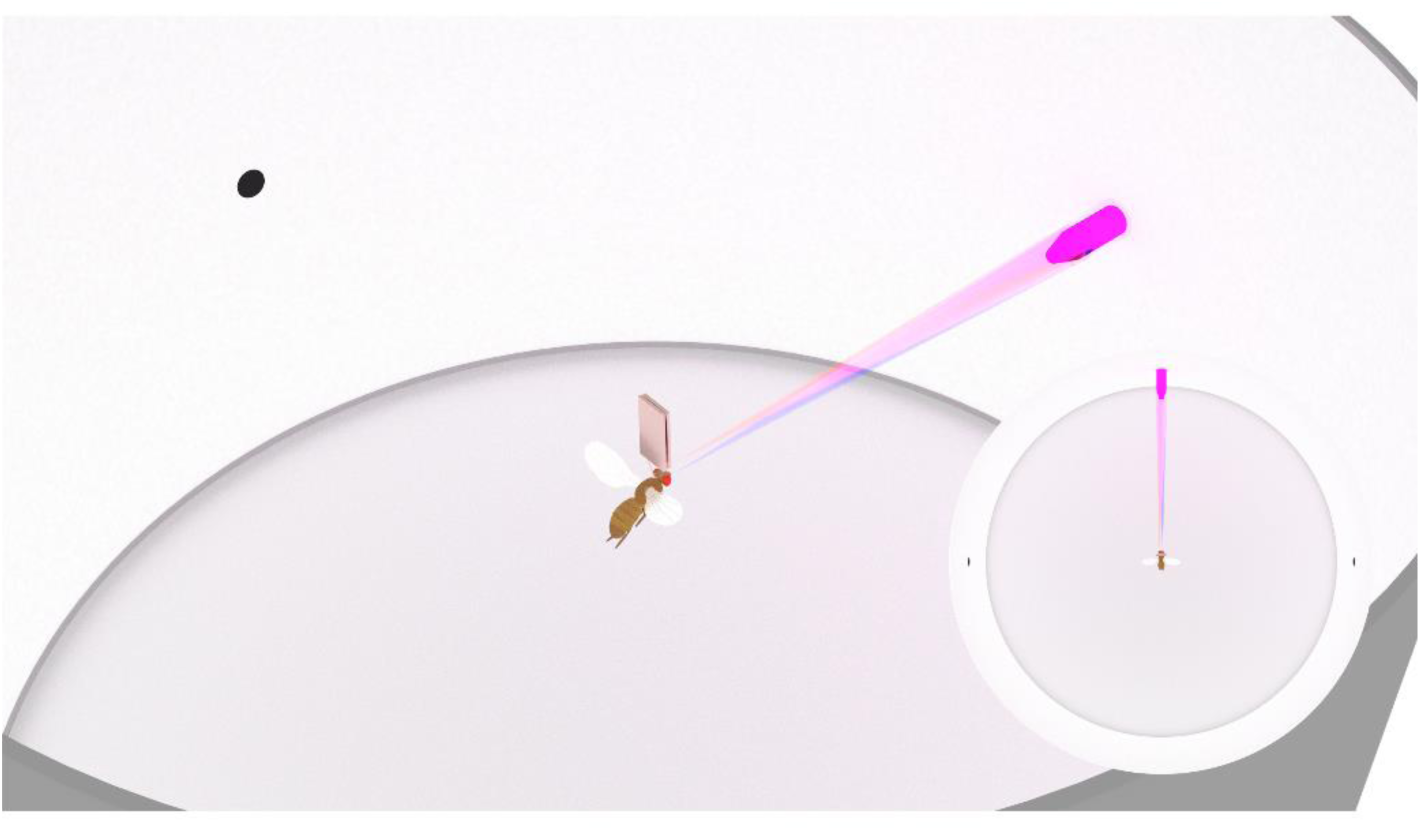
A tethered *Drosophila* must use both eyes to resolve and recognize the hyperacute 3D-pin in our flight simulator system. (*A*) In the closed-loop configuration, a fly fixates on the 3D-pin from a fixed distance (2.5 cm), making the pin fall within single corresponding left (red beam) and right (blue) eye photoreceptor receptive fields (RF, or pixels). The fly’s fixation behavior, as measured by the torque-meter, drives minute left-to-right-to-left pin movements (left-to-right-to-left cup rotations), evoking mirror-symmetric photomechanical microsaccades in the left and right photoreceptor. Because the pin moves with one RF and against the other, its movement causes phasic differences in the photoreceptor voltage responses (see Section V.9., above). The fly brain can use this dynamic neural image disparity to work out the pin size and distance from the fly eyes. Importantly, in this experimental design, as the fly is clamped to the torque-meter with its head immobilized, it cannot generate monocular cues, such as motion parallax, by approaching the pin or moving around it. Therefore, the fly needs two eyes to visually differentiate the 3D-pin-attractor from the 2D-dot-distractors of the same contrast and 2D-size. The RF sizes and angles are the same as extrapolated in Section V.9. above. See also Fig. S69 and Fig. S70, below.

The control measures in our experimental design eliminated these concerns. In our flight simulator system, a tethered fly saw the tested objects from a fixed distance (2.5 cm) and could not move its head to generate translational motion parallax (Fig. S68). Therefore, as the fly could not approach the object frontally by orienting towards it, there were no monocular cues it could use to construct a 3D representation of the object neurally. Moreover, if the fly-eye optics presented the world spatially with 4.5° pixelation (interommatidial angle) and the tested dots/pins were <3°, monocularly, each tested object would fall within a single pixel. Therefore, to have seen such a small 3D object, the fly must have used both its left and right eyes. This theoretical axiom was experimentally demonstrated in Fig. 6 *K* and *L*, while the non-visual cues, including the air current, were eliminated using the blind controls (Fig. 6*M*). Crucially, these results, together with those from further binocular (Fig. 6 *N* and *P*) and monocular (Fig. 6*O*) microsaccade controls, confirmed that *Drosophila* left and right eye photoreceptors must generate mirror-symmetric synchronous microsaccades to see small 3D objects; making the compound eyes stereopsis dynamic and phasic along with the core theory of this paper.

##### VII.5 Salience experiments (closed-loop)

*Drosophila* yaw torque responses were used to control the cup rotation, enabling the fly to choose what visual features/patterns in the panoramic scene it wanted to see. When a fly sees something interesting that it intends to inspect more closely, it characteristically brings that object in the frontal (stereo) view, “fixating to it” with small left and right rotations that keep the object simultaneously visible to both its left and right eye. In contrast to what has been shown for LED-arena type of stimulation (123), the flies find small dots attractive (and not aversive) in our flight simulator system, which uses printed visual objects and is free of LED pulse-width intensity-modulation that might scare *Drosophila*. Another key difference is the small dot sizes. We used 1° dots, which are a lot smaller than the “small” square objects (30°) used in the previous study (123).

**Fig. S69.**
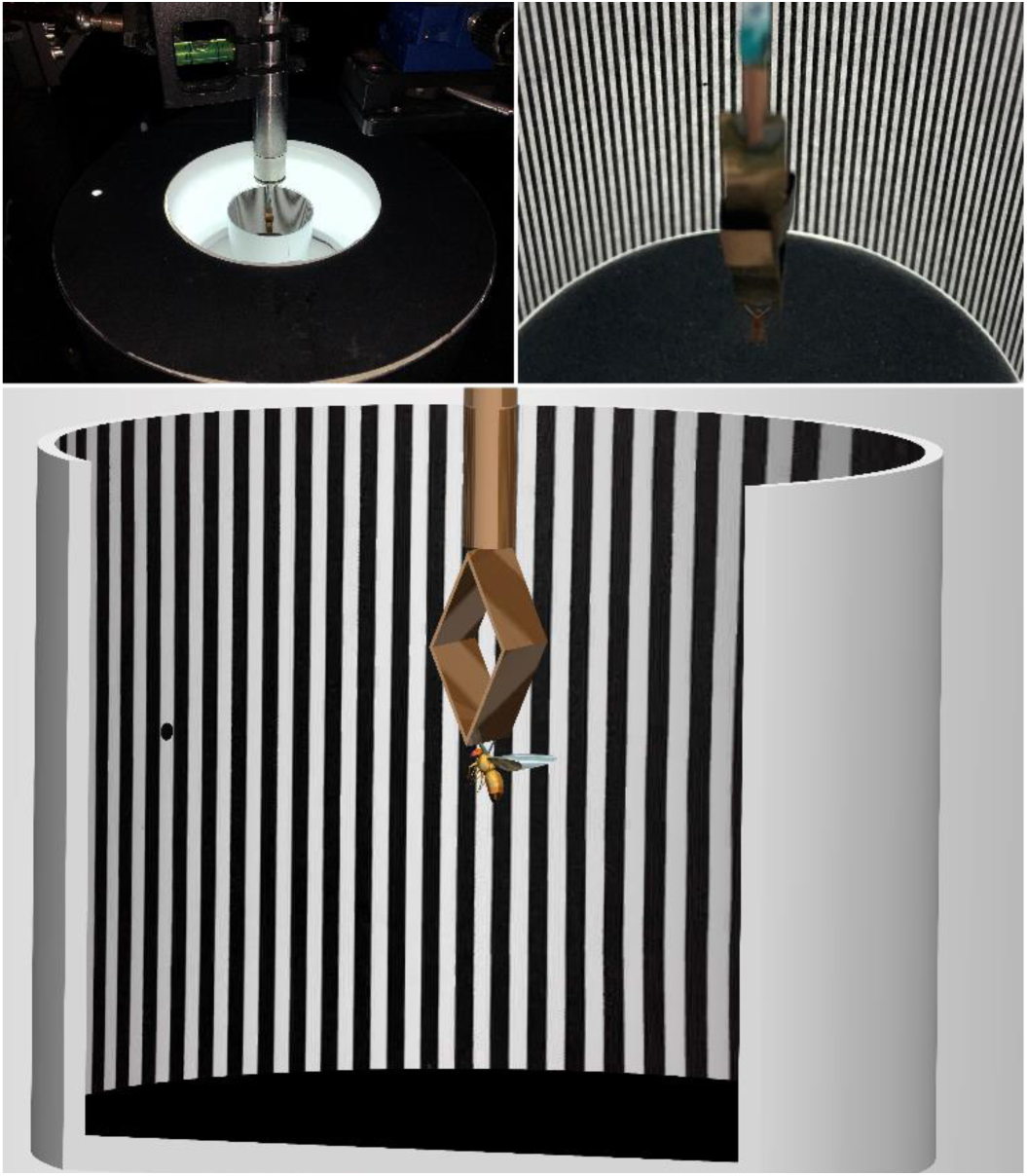
Hyperacute dot salience test in a flight simulator system, running in the closed-loop where the flying tethered *Drosophila* controls the panoramic scene position. A *Drosophila* fixates on a hyperacute dot - hidden amongst hyperacute stripes - by keeping the dot within its frontal view. The distance from the fly eyes to the panoramic screen is 25 mm (small cup).

***Testing hyperacute vision by salient 2D- and 3D-objects***

We presented different combinations of hyperacute objects at three different positions to test whether *Drosophila* saw hyperacute (<4.5° inter-ommatidial angle) stimuli at 25 mm from the eyes (the small cup).

- First, we tested visual behavior to a small black 2D-dot (0.98°) hid within a hyperacute panoramic stripe scene (with 1.17° inter-black-bar-distances) (Fig. S69). The dot was either at the scene center (0°), left (-90°), or right (90°) relative to the paper seam. The control stripe scene lacked the dot. We recorded 8 minutes of tethered flight for each case, measuring at each ms the panoramic position the fly was facing (or fixating). Each fly’s orientation behavior (relative fixation) over the panoramic scene was then given as probability.
- Second, we tested visual behavior to three black dots (3.9° Ø) on a white 360° background; The dots were at the center (0°), left (-90°), and right (90°). One of them had a small black 3D-pin (4 mm long) center (2.7° Ø) (Fig. S70). Even for a single human eye, all the dots looked the same (no clear contrast difference; Fig. S70 *B* and *C*). Thus, to see the 3D-pin dot, a fly must have stereo vision. For each fly (n = 20), we tested all three pin-positions and a blank-control white scene separately, one after another. In each of these four experiments, conducted in a random order, we recorded 8 minutes of tethered flight, continuously measuring the fly’s fixation positions. Fixation over the 360° scene was then given as probability. Fig. S71 shows an example of how five single flies performed in these separate experiments.

**Fig. S70.**
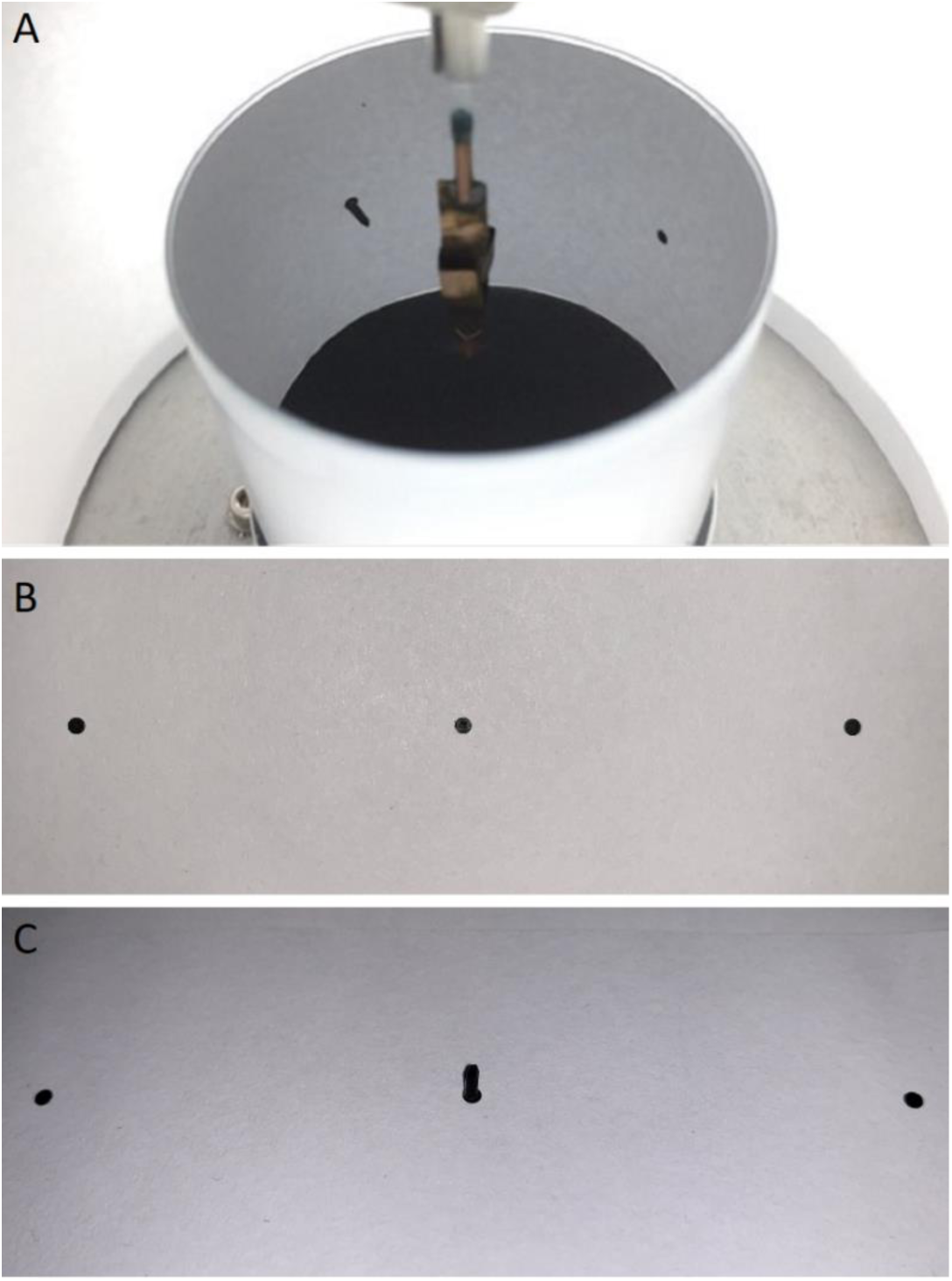
Testing visual salience of hyperacute 3D-pin vs. 2D-dots in closed-loop settings. (*A*) The stimulus configuration in the visual salience paradigm. In the experiments, a fly saw a hyperacute black pin and two hyperacute back dots 90° apart, and we measured its fixation probability of the whole 360° visual scene. (*B*) Two 3° Æ black dots (on the side) and a central black pin on the white paper background as used in the visual salience experiments. When viewed monocularly at the center of the image – parallel to the pin’s long axis – the center bin is very difficult to resolve, even for the human eye. (*C*) The black pin is visible binocularly and becomes apparent monocularly if the viewer moves sideways, as this camera image shows. However, because the fly head is immobile, clamped to the torque-meter at the center of the panorama, and cannot approach the pin or move sideways to generate motion parallax, it can only see the pin through dynamic stereopsis, sampled by mirror-symmetric photomechanical photoreceptor microsaccades.

**Fig. S71.**
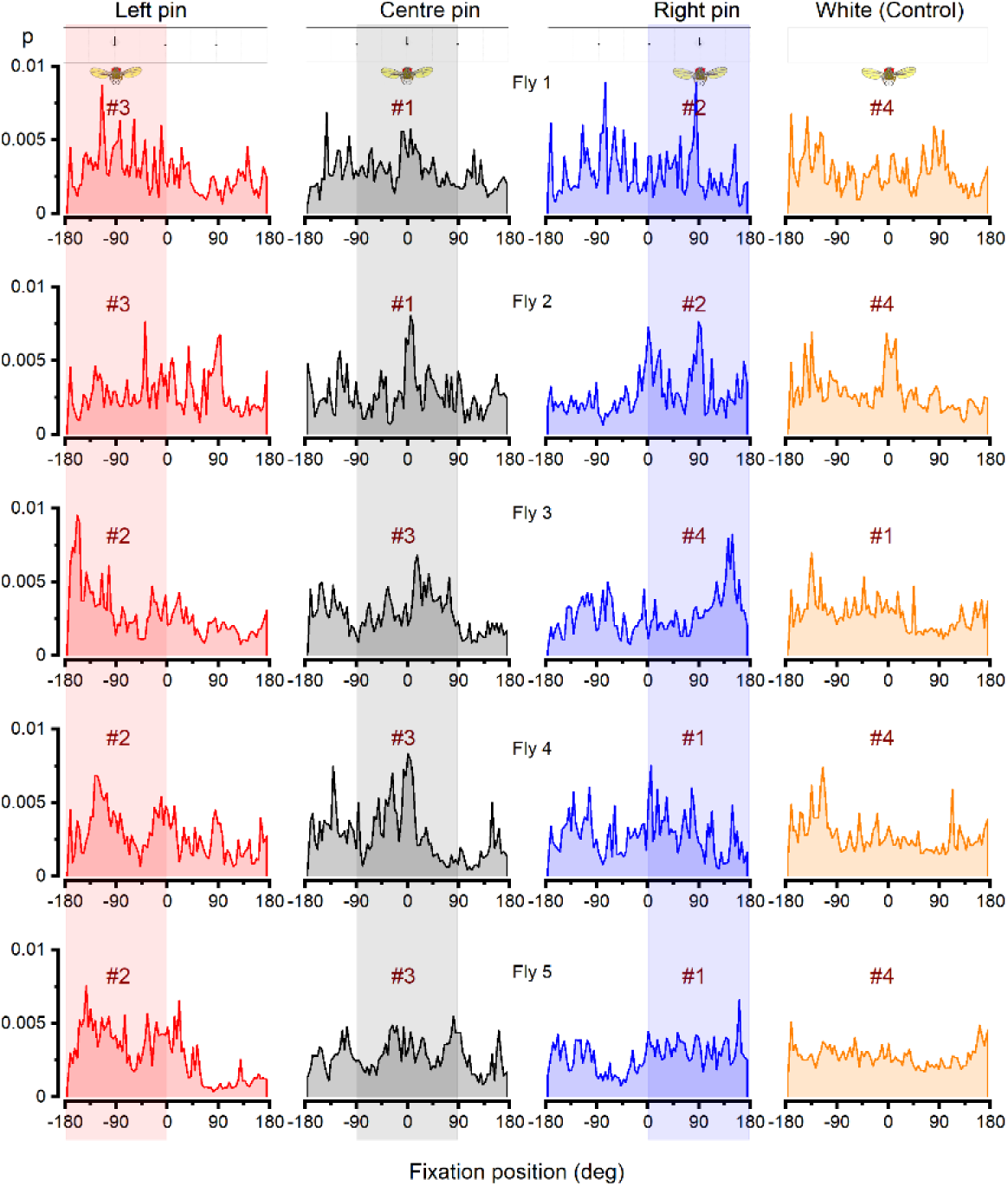
Fixation probabilities of five tethered wild-type *Drosophila* for four separate choice behavior experiments in a flight simulator system. (*A*) In three stimulus-choice tests (3 left columns), each fly saw a pin and two dots (of equal contrast and area) at three different positions (-90°, 0°, 90°). Each of these tests and the white control experiment (blank-scene; right column) lasted 8 minutes of closed-loop flying, during which the fly controlled the scene position it was facing (fixating at) by its yaw-torque responses. The fly’s fixation positions over each 360° scene were then given as probability. The test and the control scenes were presented in a random order (#1, #2, #3, and #4). The transparent red, gray, and blue bars indicate the three stimulus areas in which fixation probabilities were compared.

***Saliency analysis***. We hypothesized that:

**H1.** A fly finds a hyperacute 3D-pin (attractor) more salient than two competing hyperacute 2D-dots (distractors).
**H2.** A fly finds and fixates (is attracted) to a hyperacute 2D-dot hidden amongst hyperacute stripes.

For testing either of these hypotheses, each fly performed four consecutive experiments in random order (H1: Fig. 6C and H2: Fig. 6*H*). Three of the experiments quantified a fly’s probability density function for viewing (fixating on) the main attractor (H1: black pin; H2: black dot) when it was placed in the left (-90°), middle (0°), or right (90°), while the black dots (competing distractors) occupied the other two positions (H1) or the whole scene contained hyperacute stripes (H2) (Fig. S71, H1: left position, red; middle, gray; right, blue). The fourth (control) experiment (Fig. S71, H1: orange) quantified each fly’s *intrinsic-fixation probability density function* in exploring the homogeneous 360° background; either a white (H1) or stripe scene (H2). The *intrinsic fixation probability density function* can reveal additional visual or sensory cues in the flight simulator system that could systematically bias the fly behavior during the saliency tests. For an unbiased flight simulator system, this function should be flat over the 360° scene, as calculated using the whole tested fly population.

For comparing the flies’ fixation probabilities of the three (left, middle, and right) attractor positions, we:

- Calculated each fly’s unbiased fixation probability density function for each tested attractor position. These functions were obtained by subtracting the fly population’s mean *intrinsic-fixation probability density function* (n = 20 flies) from each fly’s fixation probability density function for each attractor experiment.
- Calculated the fly populations’ mean fixation probability density function (n = 20 flies) for the left, middle and right attractors (H1: Fig. 6*D* and H2: Fig. 6*I*).
- Calculated each flies’ fixation probability for the three attractor positions, using 180°-scene-sections with 90° section overlaps. This procedure gave each fly three mean fixation probabilities for each attractor experiment: one for the attractor (H1: pin; H2: dot with stripes) position and two for the competing distractors (H1: dot; H2: stripes alone) positions. Thus together, each fly’s three attractor experiments (left, middle, and right) gave us nine mean fixation probabilities.
- Pooled all tested flies’ (n = 20) mean fixation probabilities for the left, middle and right attractor positions into the corresponding nine groups and performed their statistical mean comparisons (H1: Fig. 6*E*; H2: Fig. 6*J*).

With each group not being tested against itself, we obtained 24 relevant mean probability comparisons (Table S7-S10) for testing statistically two questions related to H1 and H2, using one-way ANOVA:

**Q1.** Is a fly’s fixation probability at any one of the three attractor positions (say, the left position) higher when the attractor is there (the pin is at left) in comparison when it is in one of the other positions (the pin is at right or middle) (H1: Fig. 6*E*, the row above)?
**Q2.** In each experiment, is a fly’s fixation probability for the attractor (say, the left-pin) higher than its probability to fixate at the distractors (middle- and right-dots) (H1: Fig. 6*E*, the row below)?

**Table S7.** **For each test position, is the flies’ fixation probability higher when occupied by a pin-attractor?** (one-way ANOVA statistics)

| **Table S7. For each test position, is the flies' fixation probability higher when occupied by a pin-attractor?** |  |  |  |
| --- | --- | --- | --- |
| (one-way ANOVA statistics) |  |  |  |
| **Testing Q1 (pin vs dot in the same position)** | Left-pin (attractor)<br>vs | Middle-pin (attractor)<br>vs | Right-pin (attractor)<br>vs |
| Left-dot (distractor)<br>(when Middle-pin) | P = 6.590 x 10<sup>-3</sup> (**) |  |  |
| Left-dot (distractor)<br>(when Right-pin) | P = 5.623 x 10<sup>-8</sup> (***) |  |  |
| Middle-dot (distractor)<br>(when Left-pin) |  | P = 7.422 x 10<sup>-5</sup> (***) |  |
| Middle-dot (distractor)<br>(when Right-pin) |  | P = 1.593 x 10<sup>-2</sup> (*) |  |
| Right-dot<br>(when Left-pin) |  |  | P = 5.055 x 10<sup>-8</sup> (***) |
| Right-dot<br>(when Middle-pin) |  |  | P = 1.402 x 10<sup>-2</sup> (*) |

**Table S8.**
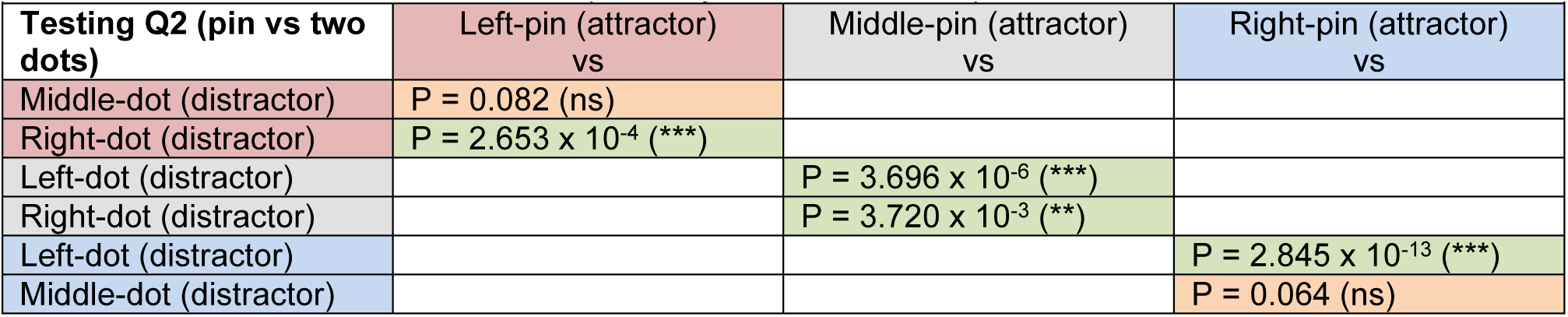
**Do the flies fixate more at a pin-attractor than the competing dot-distractors?**

**Table S9.**
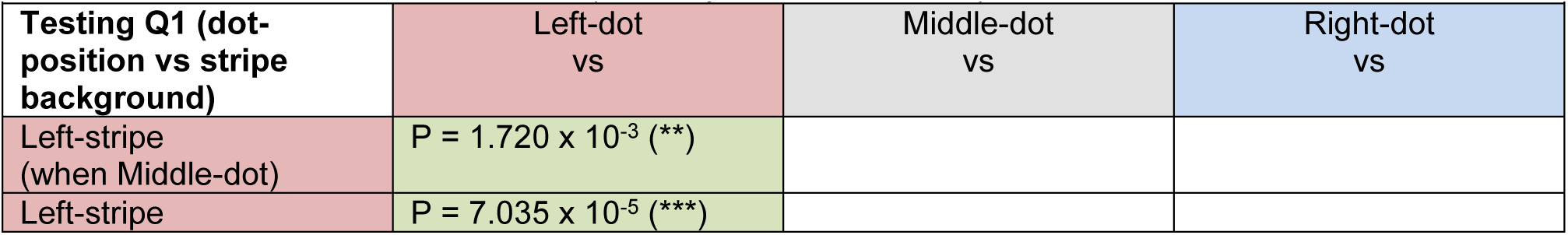
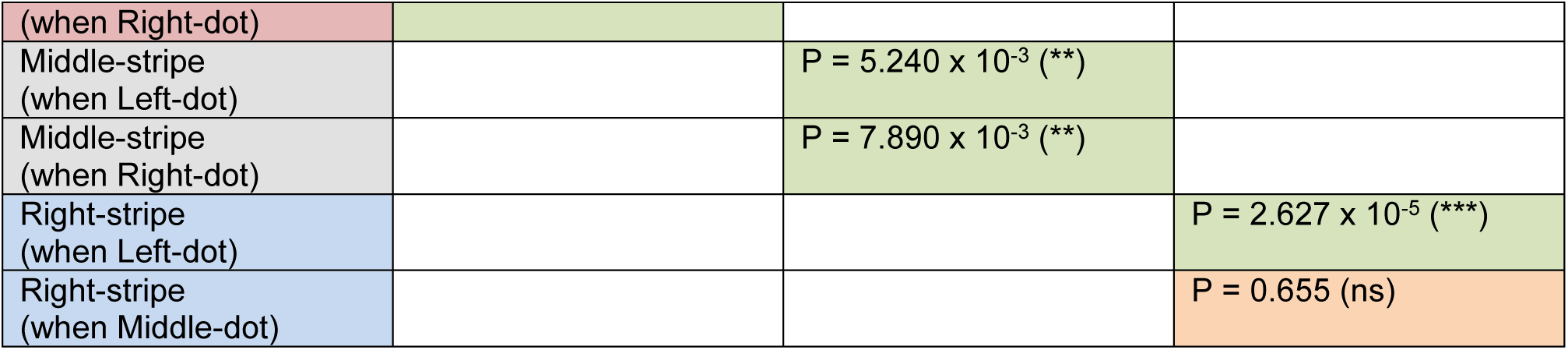
**For each test position, is the flies’ fixation probability higher when occupied by a dot-attractor?** (one-way ANOVA statistics)

**Table. S10.** **Do the flies fixate more at a dot-attractor than the competing background?**

| **Table. S10. Do the flies fixate more at a dot-attractor than the competing background?**<br>(one-way ANOVA statistics) |  |  |  |
| --- | --- | --- | --- |
| **Testing Q2 (dot-attraction vs stripe background)** | Left-position (Left-dot)<br>vs | Middle-position (Middle-dot)<br>vs | Right-position (Right-dot)<br>vs |
| Middle-position (Left-dot) | $P = 0.351 (ns)$ | | |
| Right-position (Left-dot) | $P = 5.952 \times 10^{-6} (***)$ | | |
| Left-position (Middle-dot) | | $P = 1.09728 \times 10^{-5} (***)$ | |
| Right-position (Middle-dot) | | $P = 5.090 \times 10^{-3} (**)$ | |
| Left-position (Right-dot) | | | $P = 3.803 \times 10^{-4} (***)$ |
| Middle-position (Right-dot) | | | $P = 0.920 (ns)$ |

##### VII.6 Learning experiments (closed-loop)

The avoidance associative learning experiment was automatized and recorded in 1 ms time resolution in the PC’s hard drive. The experiment consisted of a sequence of 9 blocks of 2-min duration each. During the first two blocks, the fly adapted to the flight simulator conditions without heat punishment. During training (light gray blocks in Fig. 6 *K* to *P* and Fig. S72), infrared laser light (heat) to the fly head was turned on (or off) by the computer, depending on the fly’s flight direction choice for the visual patterns at the arena wall. Under software control, the panorama was sectioned into four 90° quadrants, each having its pattern (either test or control) in its center. Identical patterns were placed in opposite quadrants. Whenever the fly’s longitudinal body axis crossed one of the panorama’s invisible quadrant-boundaries, heat (unconditioned stimulus, US) was turned either on or off. An infrared laser delivered the heat-punishment (825 nm, 150 mW), directed (using a piezo 3-axis micromanipulator; Sensapex, Finland) from the front and above onto the fly’s head and thorax. This heat-punishment (unconditioned stimulus, US) led to significant avoidance learning of the visual patterns.

Between every 2 min block, the panorama was span both clockwise and counterclockwise with a random duration that lasted for 5 s. This maneuver randomized the starting scene position for each block in respect to the fly head.

We tested both binocular (normal eyes) and monocular (either the left or right frontal eye section painted with non-toxic black acrylic paint: Winsor & Newton, Winton Oil Colour, Ivory Black – 1414331) avoidance learning. The eye was painted immediately before tethering (to the flight simulator from the copper-wire hook between the head and thorax), followed by instantly testing the fly. This procedure reduced the fly disrupting the paint coverage over their eye by attempting to rub the paint with their legs. However, many flies were able and willing to fly immediately after tethering, with minimum observable discomfort attributable to the paint. Therefore, only flies that did not repeatedly attempt to remove the paint from their eye were included in the dataset. In these experiments, we measured the *Drosophila* learning performance index (PI) for the following patterns (3D hyperacute object pairs of equal gamma-corrected contrast and size):

- A black 3D-pin at a black dot center vs. a black 2D-dot
- A black 3D-pin at a black vertical 2D-stripe center vs. a black vertical 2D-stripe (3.9° width)

As a control experiment, we measured both binocular and monocular learning performance indexes for the classic large 2D T vs. Ʇ objects (symbols), with each being 40° (height) × 40° (width) with 10° bar width. This base-metric was then compared to the corresponding hyperacute 3D learning performance indexes.

**Measuring associative learning of hyperacute 3D-objects**

A *Drosophila* controlled the panorama, which showed two opposing test objects (*e.g.,* black dots with a black center-pin, called 3D-dots) and orthogonally to them two control objects (*e.g.,* black dots, called 2D-dots). For each 18-min-long experiment, we calculated PI for each of its 2-min-blocks: as the time (in seconds) the fly selected to face CS+ (the heat-punishment associated object; the conditioned stimulus) minus the time the fly selected to face CS- (the neutral object; the non-conditioned stimulus) divided by the total time.

Because the flies learned to avoid either one of the tested objects (during the last two blocks: short-term learning: PI >0.2), as quantified after two bouts of training (*i.e.,* teaching) with heat-punishment (high avoidance, performance index, PI >0.8), they must have seen the small 3D differences between the objects, as required for hyperacute stereopsis. Importantly, *Drosophila* learned similarly well (blue: PI >0.2) in the classic T vs. Ʇ paradigm.

For each genotype, we tested the flies’ learning performance for all the predetermined test objects. *E.g.,* in one-half of the 3D-pin vs. 2D-dot experiments, the heat-punishment was associated with the 3D-pin (10/20 wild-type flies) and the other half with the 2D-dot (10/20 flies). Predictably, as learning required distinguishing (seeing) the two patterns as different, the flies learned to avoid 3D-pin and 2D-dot equally well, with similar PIs - and the data were pooled.

The two rims, joining the paper strip’s short ends, caused a faint narrow seam (∼0.1°) in the white background panorama. However, this seam did not affect *Drosophila* visual object learning; *i.e.,* the flies did not use it as a positional learning cue. Furthermore, we kept the same paper strips in both binocular and monocular (one eye painted black) experiments as the tested hyperacute 2D and 3D objects’ backgrounds. Therefore, if the flies used the seam as a visual learning cue, both binocular and monocular flies would have shown a positive learning performance index. However, because only the binocular flies learned to avoid the hyperacute 2D and 3D objects (Fig. S72*A*, the row above *binocular* vs. the row below *monocular*), the seam had no role in the measured learning performances, and the flies used stereo vision to differentiate and memorize the tested objects.

Interestingly, after the first object training (after the 3^th^-4^th^ block heat-avoidance training spout), many flies (Fig. S72) showed small but insignificant PI, indicating that the 1^st^ teaching spout caused only a transient change in their behavioral choices. This finding is consistent with the theory of dynamic learning. To improve survival, animals would need to continually question the learned information as the world is not static but changes continuously. In other words, it would be beneficial to check whether a recently seen predator was still there rather than believe that nothing had changed, and if the predator had moved (was no longer there), then change the behavior. Similarly, our data suggest that after the 1^st^ teaching spout, the flies soon changed their behavior (in respect to their avoidance PI during training), as if to check whether they would still be heat-punished when looking at the object. And since the punishment no longer occurred, they could actively forget the learned association between the tested object and the heat punishment. However, in clear contrast, the 2^nd^ heat-punishment teaching spout caused a highly significant and longer-lasting object avoidance in the flies’ behavioral choice (Fig. S72 and Table S11-S14).

**Fig. S72.**
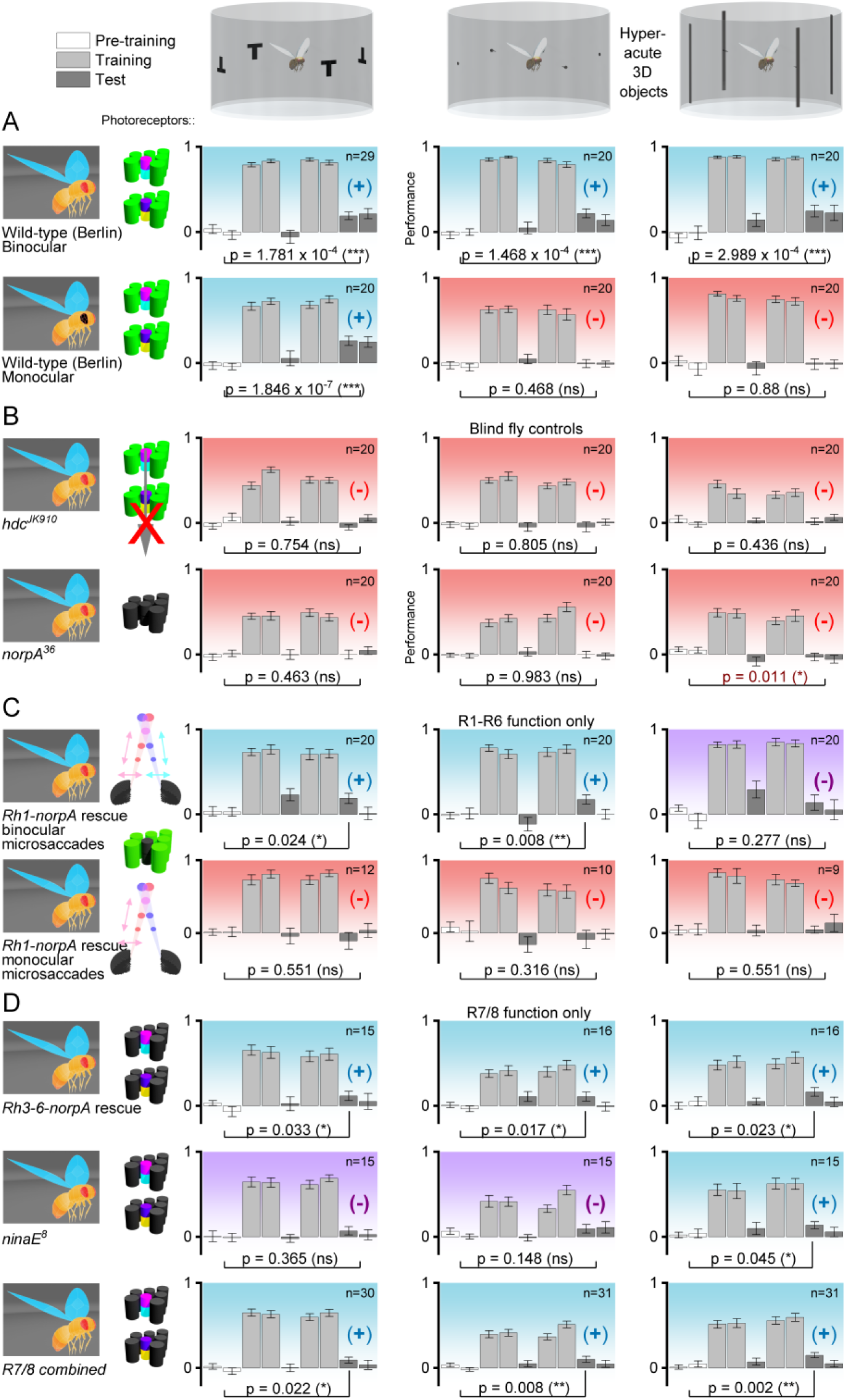
*Drosophila* with binocular synchronous mirror-symmetric microsaccades learn to avoid hyperacute 3D visual stimuli associated with heat punishment. (*A*) In a flight simulator system, wild-type flies’ learning performance for hyperacute 3D objects (dot vs. pin-dot; stripe vs. pin-stripe) is similarly positive to large 2D objects (T vs. Ʇ) indicating that the flies must see the tested nearby objects in super-resolution stereo. *Drosophila* could not learn 3D hyperacute objects monocularly (with one eye painted black) but learned large 2D objects, meaning that two eyes are needed for super-resolution stereopsis. (*B*) Blind control mutants: *hdc^JK910^* (lacks photoreceptor neurotransmitter, histamine) and *norpA^36^*(faulty phototransduction), in which other senses ought to function normally, did not learn to avoid the tested visual objects, meaning that the wild-type learning was predominantly visual; *i.e.,* not based upon auditory, tactile or olfactory cues. (*C*) *Rh1*-*norpA* rescue flies (only R1-R6 functioning), which had normal ERGs in both eyes but showed monocular microsaccades (lateral photoreceptor microsaccades only in one eye; left or right), could not learn the tested visual objects. These results demonstrate that synchronous mirror-symmetric binocular photoreceptor microsaccades are necessary for super-resolution stereo vision. (*D*) Both Rh3-6-*norpA* rescue flies (only R7/R8 functioning) and *ninaE^8^* mutants (only R7/R8 functioning) learned the 3D hyperacute but less well than the wild-type flies, meaning that both R1-R6 (in **C**) and R7/R8 contribute to high-resolution stereo vision. Blue panels and (+) indicate significant visual learning; red panels and (-) indicate no learning; purple panels and (-) (three cases) indicate positive learning performance indexes, which were not significant. For each fly group, the significance of learning was calculated between the pre-training and test responses. In the R1-R7/8 photoreceptor insets (left), the bright colors indicate the functioning photoreceptors with their normal photopigments; dark gray indicates the blind photoreceptors.

**Table S11.**
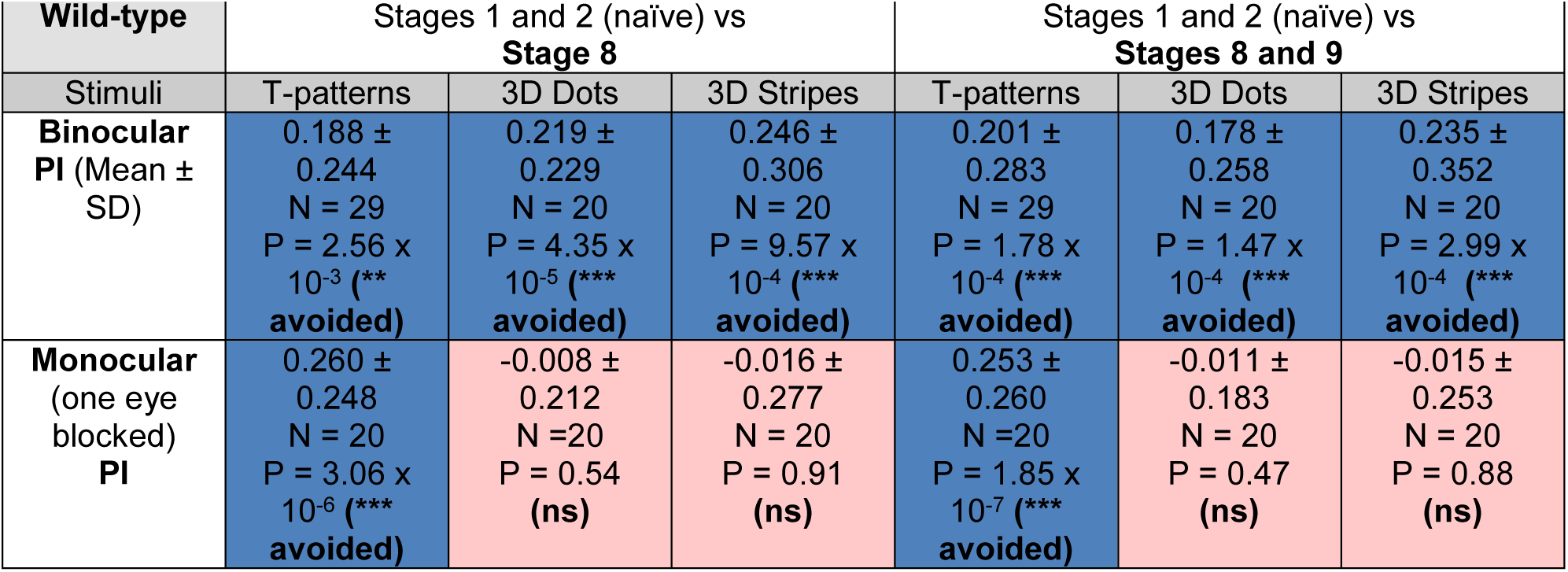

| Wild-type | Stages 1 and 2 (naïve) vs<br>Stage 8 |  |  | Stages 1 and 2 (naïve) vs<br>Stages 8 and 9 |  |  |
| --- | --- | --- | --- | --- | --- | --- |
| Stimuli | T-patterns | 3D Dots | 3D Stripes | T-patterns | 3D Dots | 3D Stripes |
| **Binocular**<br>**PI** (Mean ±<br>SD) | 0.188 ±<br>0.244<br>N = 29<br>P = 2.56 x<br>10<sup>-3</sup> (**<br>avoided) | 0.219 ±<br>0.229<br>N = 20<br>P = 4.35 x<br>10<sup>-5</sup> (***)<br>avoided) | 0.246 ±<br>0.306<br>N = 20<br>P = 9.57 x<br>10<sup>-4</sup> (***)<br>avoided) | 0.201 ±<br>0.283<br>N = 29<br>P = 1.78 x<br>10<sup>-4</sup> (***)<br>avoided) | 0.178 ±<br>0.258<br>N = 20<br>P = 1.47 x<br>10<sup>-4</sup> (***)<br>avoided) | 0.235 ±<br>0.352<br>N = 20<br>P = 2.99 x<br>10<sup>-4</sup> (***)<br>avoided) |
| **Monocular**<br>(one eye<br>blocked)<br>**PI** | 0.260 ±<br>0.248<br>N = 20<br>P = 3.06 x<br>10<sup>-6</sup> (***)<br>avoided) | -0.008 ±<br>0.212<br>N = 20<br>P = 0.54<br>(ns) | -0.016 ±<br>0.277<br>N = 20<br>P = 0.91<br>(ns) | 0.253 ±<br>0.260<br>N = 20<br>P = 1.85 x<br>10<sup>-7</sup> (***)<br>avoided) | -0.011 ±<br>0.183<br>N = 20<br>P = 0.47<br>(ns) | -0.015 ±<br>0.253<br>N = 20<br>P = 0.88<br>(ns) |

**Table S12.**

| R1-R6<br>function | Stages 1 and 2 (naïve) vs<br>Stage 8 |  |  | Stages 1 and 2 (naïve) vs<br>Stages 8 and 9 |  |  |
| --- | --- | --- | --- | --- | --- | --- |
| Stimuli | T-patterns | 3D Dots | 3D Stripes | T-patterns | 3D Dots | 3D Stripes |
| *norpA* Rh1<br>rescue<br>Binocular<br>Saccades<br>PI (Mean ±<br>SD) | 0.187 ±<br>0.268<br>N = 20<br>P = 2.42 x<br>10<sup>-2</sup><br>(* avoided) | 0.172 ±<br>0.240<br>N = 20<br>P = 7.75 x<br>10<sup>-3</sup><br>(** avoided) | 0.138 ±<br>0.407<br>N = 20<br>P = 0.141<br>(ns) | 0.098 ±<br>0.311<br>N = 20<br>P = 0.286<br>(ns) | 0.086 ±<br>0.263<br>N = 20<br>P = 0.105<br>(ns) | 0.094 ±<br>0.466<br>N = 20<br>P = 0.277<br>(ns) |
| Monocular<br>Saccades<br>PI | -0.104 ±<br>0.389<br>N = 12<br>P = 0.214<br>(ns) | -0.084 ±<br>0.394<br>N = 10<br>P = 0.315<br>(ns) | 0.05 ± 0.148<br>N = 9<br>P = 0.968<br>(ns) | -0.032 ±<br>0.360<br>N = 12<br>P = 0.551<br>(ns) | -0.047 ±<br>0.311<br>N = 10<br>P = 0.316<br>(ns) | 0.095 ±<br>0.268<br>N = 9<br>P = 0.551<br>(ns) |

**Table S13.**
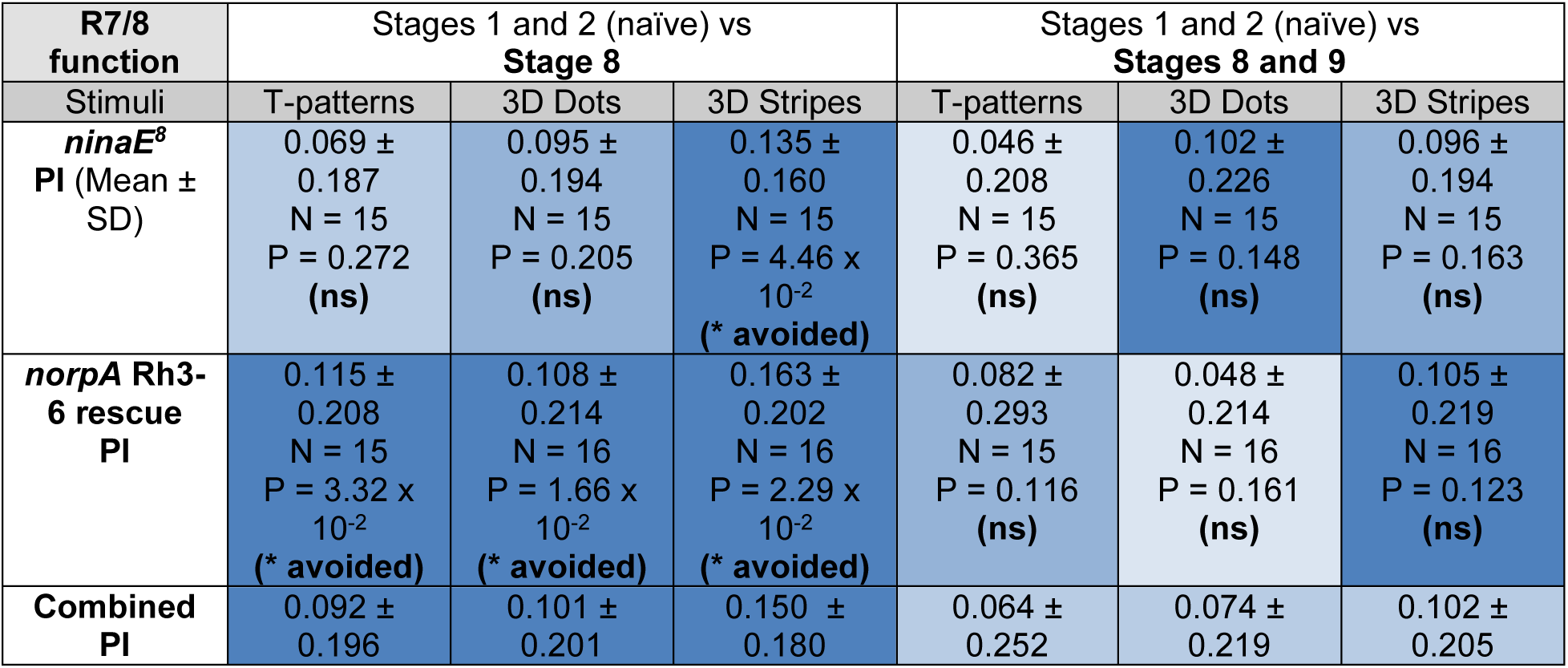
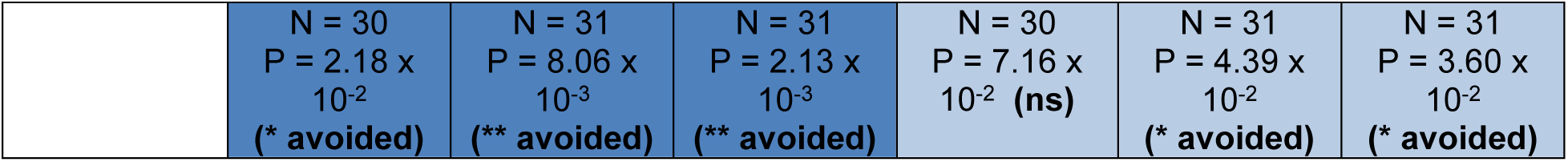

**Table S14.**

| Blind mutants | Stages 1 and 2 (naïve) vs Stage 8 |  |  | Stages 1 and 2 (naïve) vs Stages 8 and 9 |  |  |
| --- | --- | --- | --- | --- | --- | --- |
|  | T-patterns | 3D Dots | 3D Stripes | T-patterns | 3D Dots | 3D Stripes |
| *hdc<sup>JK910</sup>*<br>PI (Mean ± SD) | -0.053 ± 0.142<br>N = 20<br>P = 0.151 (ns) | -0.048 ± 0.259<br>N = 20<br>P = 0.841 (ns) | 0.019 ± 0.172<br>N = 20<br>P = 0.903 (ns) | 0.002 ± 0.164<br>N = 20<br>P = 0.754 (ns) | -0.019 ± 0.214<br>N = 20<br>P = 0.618 (ns) | 0.042 ± 0.168<br>N = 20<br>P = 0.436 (ns) |
| *norpA*<br>PI | -0.002 ± 0.240<br>N = 20<br>P = 0.874 (ns) | -0.002 ± 0.174<br>N = 20<br>P = 0.790 (ns) | -0.032 ± 0.206<br>N = 20<br>P = 0.051 (~* attracted) | 0.022 ± 0.221<br>N = 20<br>P = 0.463 (ns) | -0.012 ± 0.168<br>N = 20<br>P = 0.983 (ns) | -0.045 ± 0.204<br>N = 20<br>P = 0.01083 (* attracted) |

| Heat punishment avoidance learning performance index (PI) scale |  |  |  |  |  |  |
| --- | --- | --- | --- | --- | --- | --- |
| PI > 0.1 | 0.1 > PI > 0.08 | 0.075 > PI > 0.05 | 0.05 > PI > 0.025 | 0.025 > PI > 0 | 0 > PI > -0.025 | -0.025 > PI |

Clear avoidance
Slight avoidance
Random
Slight attraction

***Choosing a heat-punishment direction.*** We further found that the heat-punishment direction and the fly’s body location receiving it - here, directed from the up-front to its head; see above - contributed to the flies’ PI. In control experiments, in which the heat-punishment was directed from behind and above onto the fly’s head and thorax, the learning performance indexes were somewhat higher, closely matching with the previous results (49, 124) using a similar delivery (Fig. S73). Thus, suggestively, for the flies to form aversive object associations, targeting the heat-punishment to the head’s back provides a more potent unconditioned stimulus than heat-punishment to the head’s front. Nevertheless, in this study, we settled to the above-described tethering and frontal heat-punishment direction because it minimized the possibility of the flies seeing additional visual cues, such as the opening and closing of light-proof curtains and any experimenter activity during the experiments. Thus, this arrangement ensured that the obtained statistical differences between the different test and control experiments became undisputable within the given experimental settings and their limitations.

**Fig. S73.**
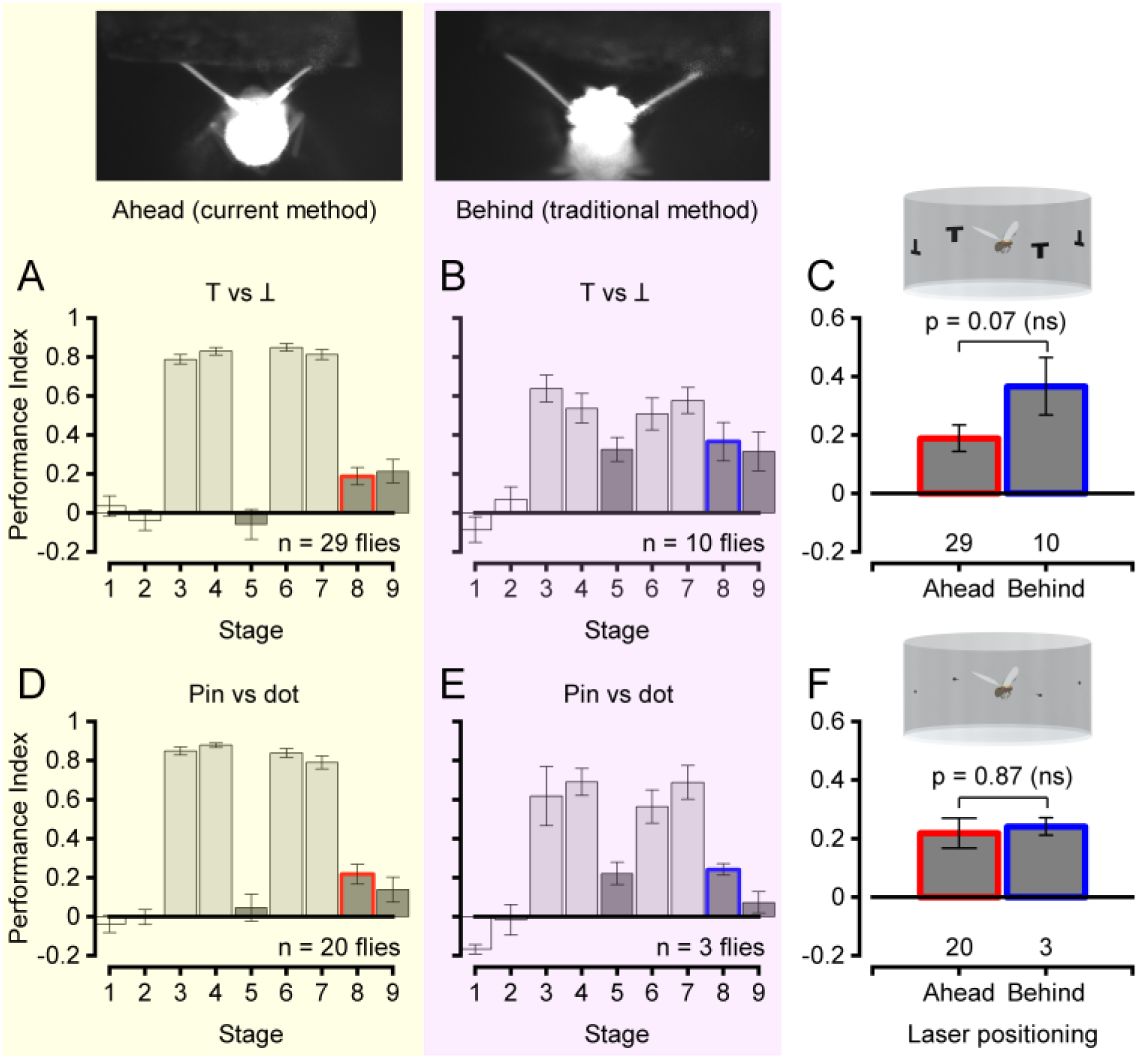
The heat-punishment direction onto the fly head affects the visual object avoidance learning. (*A*) Wild-type flies’ object avoidance training (light gray bars) and learning (dark gray) scores for the classic T vs. Ʇ paradigm when IR-heat-punishment was directed onto the fly head from the up-front direction. (*B*) Wild-type flies’ object avoidance training and learning scores for the same T vs. Ʇ patterns, but now IR- laser heat punishment was directed onto the fly head and thorax from behind. The flies’ heat-punishment avoidance is more robust for the front- direction (*A*), yet their object learning performance index is higher for the back-direction (*B*). The back-direction heat-punished flies’ performance index values for the T vs. Ʇ paradigm match those of the earlier studies (49, 124). (*C*) The learning performance indexes for stage 8, immediately after the last two spouts of training, appeared somewhat higher for the traditional heat-punishment back-direction. (*D*) Object avoidance training (light gray bars) and learning (dark gray) scores for the hyperacute pin vs. dots paradigm when IR-heat-punishment was directed onto the fly head from the up-front direction. (*E*) Object avoidance training (light gray bars) and learning (dark gray) scores for the hyperacute pin vs. dots paradigm when IR-heat-punishment was directed onto the fly head from behind. (*F*) The learning performance indexes for stage 8 (*D* and *E*) seemed overall similar for the front head and back heat-punishment.

**Measuring associative learning of flies showing monocular photoreceptor microsaccades**

Serendipitously, while collecting data for the different genotypes’ photoreceptor microsaccade and electroretinogram (ERG) statistics (see Section II.8., above), we found some Rh1-*norpA* rescue flies lacking the sideways-moving microsaccades in one of their eyes. And, intriguingly, since both of their eyes showed normal ERG responses, presumably, something must have gone wrong during the development of the mechanical linkages guiding the rhabdomeres lateral microsaccade movements in one of their eyes (see Section II.4., above). We realized the importance of these flies, as they enabled us to test the role of mirror-symmetric microsaccades in stereopsis directly. To do this systematically, we established a 3-pronged experimental protocol (*multi-method paradigm*) for testing every *Rh1-norpA rescue fly*. The protocol included separate DPP and ERG recordings of the flies’ left and right eyes and flight-simulator learning experiments, all performed on the same day within about 2 hours. These combined experiments enabled us to identify:

i. Flies with normal R1-R6 phototransduction and binocular mirror-symmetric lateral photoreceptor microsaccades.
ii. Flies with normal phototransduction but monocular asymmetric lateral photoreceptor microsaccades.
iii. Blind flies without photoreceptor microsaccades and flat ERGs.

Therefore, we could reliably link the **i**- and **ii**-grouped flies’ hyperacute 3D object learning performance to their normal or faulty photoreceptor microsaccade function.

***Multi-method paradigm.*** First, the flies were tethered and tested for associative avoidance learning with one of the three hyperacute 2D or 3D patterns in the flight simulator. Then, we generally unhooked the flies and fixated them on a pipette tip for faster and less error-prone handling, although few flies were tested tethered. In the deep pseudopupil setup, photoreceptor microsaccades were recorded to 200 ms green- or UV-flashes repeated 25 times every 2 s for additional statistics. These recordings were performed from two fixed locations on the ventral left and right eyes: +28° and -28° horizontal rotations from the midline with constant -37° vertical rotation from the antennae. Finally, we stimulated and recorded the ERG-responses approximately from the same locations where the microsaccades were imaged, although only the right eye was used for a minority of the flies. We measured the ERGs last to avoid any Ringer solution spillage on the fly-eye or minor damage from the eye-touching electrodes, both of which could have influenced the learning and the microsaccades. Further details of the ERG and deep pseudopupil recording methods are presented in Section II.3. and Section II.1, respectively. The details of the avoidance learning testing can be found in Section VII.6.

***Binocular microsaccades.*** Initially, we assumed that the Rh1-*norpA* rescue would generate a homogenous group of flies with similar eyesight, but based on the photoreceptor microsaccades and the ERG-responses, these flies clustered into three groups with very distinctive visual capabilities (Fig. S74). Most flies (∼80%) showed binocular microsaccades (Fig. S74*B*, green) and regular ERG-responses of approximately 3 mV with transient On- and Off-responses (Fig. S74*C*, green). These flies could learn to avoid both the 2D (T vs. Ʇ) and the 3D (dot vs. pin-dot and stripe vs. pin-stripe) hyperacute testing patterns (Fig. S74*A*, green). However, compared to the wild-type, the binocular Rh1-*norpA* flies’ 3D avoidance learning performances seemed somewhat weaker. Nevertheless, the difference was not statistically significant (Table S15-S20, below) for any of the un-pooled or pooled patterns, demonstrating that normal binocular microsaccades are sufficient for hyperacute 2D/3D avoidance learning even without the functioning R7/8s.

**Fig. S74.**
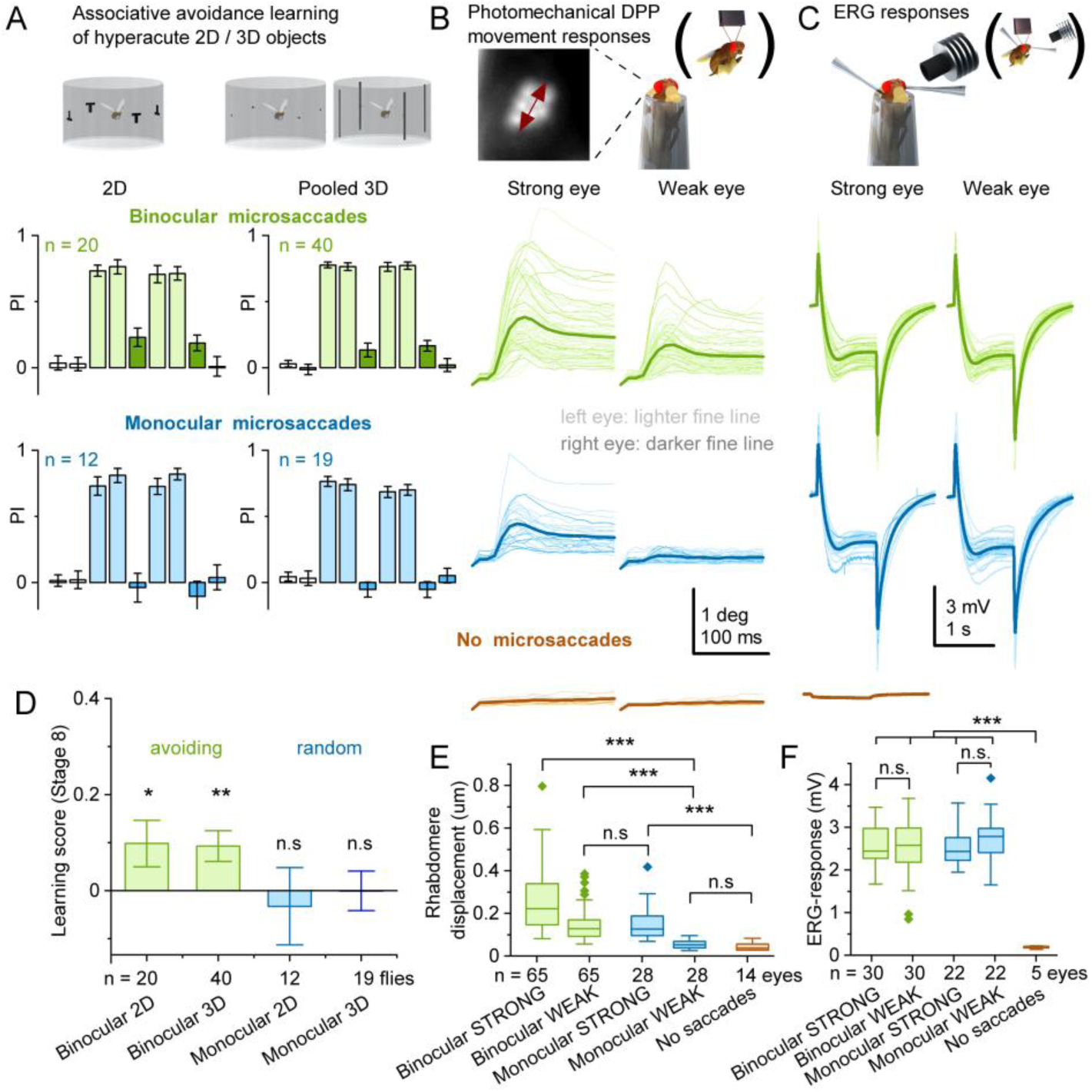
Rh1-*norpA* rescue flies with monocular microsaccades fail to learn hyperacute 2D and 3D patterns. (*A*) 2D and pooled 3D learning scores of *Rh1-norpA* rescue flies with binocular (green) or monocular lateral microsaccades (blue) or no microsaccades (brown). The binocular microsaccade flies learned both the 2D and 3D patterns, whereas the monocular ones failed to learn these. (*B*) Deep pseudopupil microsaccades (DPP-MS), recorded to 200 ms bright UV light pulses, divided into strongly and weakly responding eyes. (*C*) ERG-recordings of the same flies, divided into strongly and weakly responding eyes by the DPP-MS responses. The ERGs indicate that the no-microsaccade flies are blind. (*D*) Quantified learning scores for the data in (A) show that normal, binocular microsaccades enable successful learning, whereas the flies with monocular microsaccades failed to learn. (*E*) The summarized microsaccade data from (B) shows that the weak monocular microsaccade eyes do not statistically differ from the no-saccade flies, but the weakly binocular microsaccade eyes do, supporting the presented grouping to binocular, monocular, and no-saccade flies. (*F*) ERG-responses of binocular and monocular flies’ weak eyes are significantly larger than the ERGs in no saccade flies, indicating that the monocular microsaccades are not a result of insufficient Rh1 expression.

***Monocular microsaccades.*** Besides the binocular microsaccade flies, about 10% of the randomly selected Rh1-*norpA* rescue flies showed monocular microsaccades (Fig. S74*B*, blue) and normal ERGs (Fig. S74*C*, blue). Interestingly, these flies could neither learn the 2D testing pattern nor the 3D patterns (Fig. S74*A*, blue).

To acquire a sufficient number of these flies, we ran a preselection program where hundreds of Rh1-*norpA* rescue flies were first checked in the deep pseudopupil (DPP) setup for their microsaccades, discarding the flies with binocular and no microsaccades while proceeding on with the monocular microsaccade flies. We classified the flies with one eye microsaccade movement smaller than one camera pixel as monocular because movements of this size or larger can be confidently distinguished from the no-movement. To maximize the preselection throughput, we used the pipette-tip fixation method over the more time-consuming tethering. However, because the found monocular microsaccade flies were soon to be tethered for the flight simulator experiments, we only applied a small blob of barely-melting beeswax on the fly thorax pipette tip interface, leaving the head free to move during the preselection. This single blob of wax was easily removed using tweezers if the fly turned out to have monocular microsaccades. After the flight simulator experiments, the DPP and ERG recordings were performed as described earlier (see *Multi-method paradigm*, above). Unexpectedly, a total of 4 flies changed from showing monocular to binocular microsaccades between the preselection and the final pseudopupil recordings, potentially reflecting additional neural activity modulation from the fly brain (see microsaccade variability in Section II.8.ii., above). Considering that these experiments were immensely onerous and that these four flies showed similar “no-learning” scores to the monocular flies, we decided to include them in the monocular group’s learning data.

***No microsaccades.*** Besides the binocular and monocular microsaccades, we observed <10% of Rh1- *norpA* rescue flies with the total absence of microsaccades (Fig. S74*B*, brown). Crucially, these flies were also unresponsive to both green- and UV-flashes in the ERG recordings (Fig. S74*C*, brown), indicating that they were, indeed, blind. Because their blindness - but not the lack of microsaccades - would explain any discrepancies in the visual avoidance learning observed between the binocular and monocular microsaccade flies, we did not investigate these flies further, and their learning was not tested systematically. In this small minority of the Rh1-*norpA* rescue flies, the Rh1 expression presumably failed during the development.

Overall, our multi-method paradigm with Rh1-*norpA* rescue flies demonstrated that normal binocular photoreceptor microsaccades are necessary for hyperacute 2D/3D avoidance learning (Fig. 6N-P and Fig. S72*C*). The monocular microsaccades almost certainly broke the spatiotemporal correlations between left and right eyes’ neural images, making visual learning difficult. Because the no-saccade-flies were blind, we could not examine if the total microsaccade absence affected the learning, but perhaps this can be probed in the future by genetic or pharmacological interventions. It appears, however, that the absolute photoreceptor microsaccade size predicts the flies’ learning on the population level (Fig. S75), although other factors and differences between the groups are likely playing a role as well. Video-file showing examples of monocular microsaccades can be downloaded from: https://github.com/JuusolaLab/Hyperacute_Stereopsis_paper/tree/master/MonocularMS

**Fig. S75.**
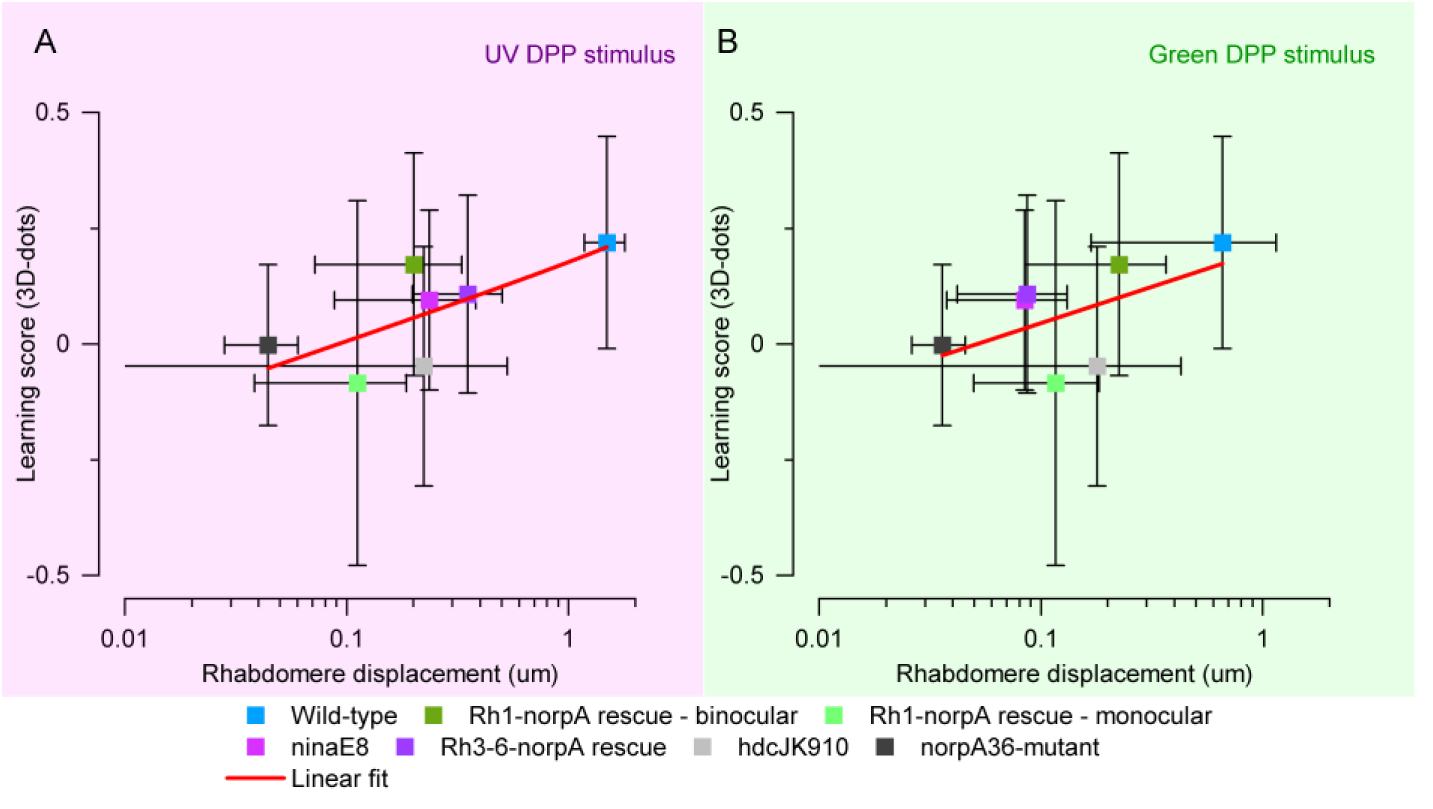
Population- level regression implies a correlation between microsaccade amplitude and visual learning. (*A* and *B*) Population-level 2D-dots-vs-3D-pins learning scores as functions of the microsaccade amplitude - deep pseudopupil (DDP) rhabdomere displacement magnitude - to a UV- and green-flash, respectively. The red lines indicate linear regressions with Pearson’s r = 0.704 for the UV- (*A*) and r = 0.550 for the green-light DPP stimulation (*B*).

##### VII.7 Comparable learning experiment statistics

The statistical (one-way ANOVA) comparisons between the different *Drosophila* geno- and phenotypes’ learning performance indexes at stage-8 for hyperacute 3D- and large 2D-objects are shown in Fig. S76 (group-wise) and Fig. S77 (pooled) and listed in Table S15-S20.

**Fig. S76.**
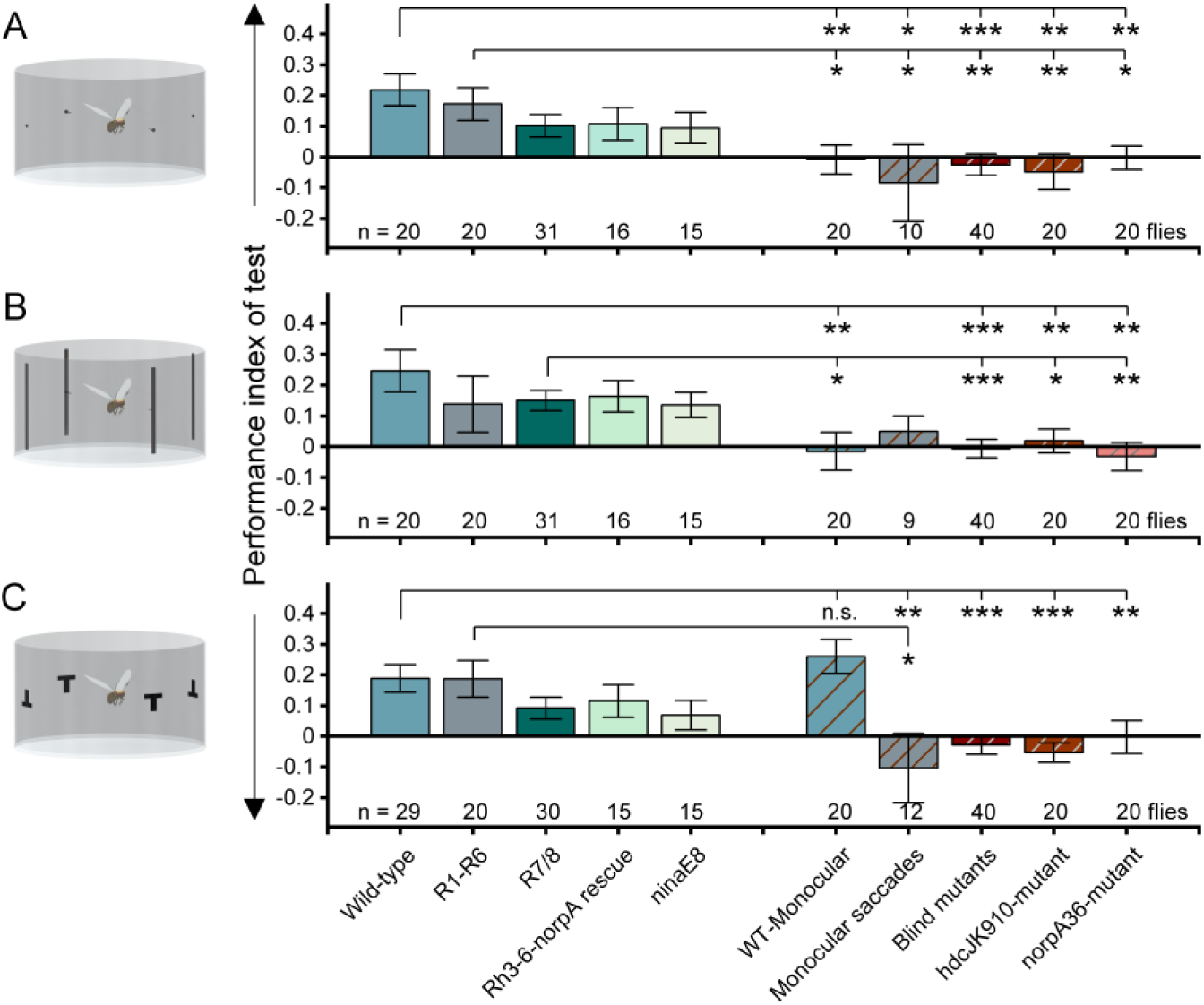
Comparing the different fly geno- and phenotypes’ stage-8 aversive performance learning indexes for the hyperacute 3D- & and large 2D-objects (T-patterns). (*A*) hyperacute pin-dots vs. hyperacute dots. (*B*) hyperacute pin- stripes vs. hyperacute stripes. (*C*) Large T- vs Ʇ- patterns.

**Table S15.**

| 3D Dots |  |  |  |  |  |  |  |  |  |  |
| --- | --- | --- | --- | --- | --- | --- | --- | --- | --- | --- |
| Table S15 | Wild-type | *norpA* Rh1 rescue | R7/8 pool | *norpA* Rh3- | *ninaE<sup>8</sup>* | WT Monocular | *norpA* Rh1 rescue | Blind pool | Blind *hdc<sup>JK910</sup>* | Blind |

|  | type | Binocular Saccades |  | 6 rescue |  | (one eye blocked) | Monocular Saccades |  |  | *norpA* |
| --- | --- | --- | --- | --- | --- | --- | --- | --- | --- | --- |
| Wild-type |  | P = 0.535 (ns) | P = 0.06 (ns) | P = 0.146 (ns) | P = 0.1 (ns) | P = 0.002 (**) | P = 0.012 (*) | P = 1.773 x 10<sup>-4</sup> (***) | P = 0.0014 (**) | P = 0.001 (**) |
| *norpA* Rh1 rescue Binocular Saccades |  |  | P = 0.261 (ns) | P = 0.407 (ns) | P = 0.313 (ns) | P = 0.016 (*) | P = 0.035 (*) | P = 0.00231 (**) | P = 0.0083 (**) | P = 0.012 (*) |
| R7/8 pooled |  |  |  |  |  | P = 0.069 (ns) | P = 0.056 (ns) | P = 0.0147 (*) | P = 0.025 (*) | P = 0.064 (ns) |
| *norpA* Rh3-6 rescue |  |  |  |  | P = 0.861 (ns) | P = 0.114 (ns) | P = 0.12 (ns) | P = 0.044 (*) | P = 0.062 (ns) | P = 0.098 (ns) |
| *ninaE*<sup>8</sup> |  |  |  |  |  | P = 0.15 (ns) | P = 0.144 (ns) | P = 0.068 (ns) | P = 0.083 (ns) | P = 0.129 (ns) |
| WT Monocular (one eye blocked) |  |  |  |  |  |  | P = 0.495 (ns) | P = 0.773 (ns) | P = 0.596 (ns) | P = 0.926 (ns) |
| *norpA* Rh1 rescue Monocular Saccades |  |  |  |  |  |  |  | P = 0.527 (ns) | P = 0.767 (ns) | P = 0.433 (ns) |
| Blind pooled |  |  |  |  |  |  |  |  |  |  |
| Blind *hdc*<sup>JK910</sup> |  |  |  |  |  |  |  |  |  | P = 0.516 (ns) |
| Blind *norpA* |  |  |  |  |  |  |  |  |  |  |

**Table S16.**

All tests were one-way ANOVA comparing two groups together.
| 3D Stripes |  |  |  |  |  |  |  |  |  |  |
| --- | --- | --- | --- | --- | --- | --- | --- | --- | --- | --- |
| Table S16 | Wild-type | *norpA* Rh1 rescue | R7/8 pooled | *norpA* Rh3-6 | *ninaE*<sup>8</sup> | WT Monocular | *norpA* Rh1 rescue | Blind pooled | Blind *hdc*<sup>JK910</sup> | Blind *norpA* |

|  |  | Binocular Saccades |  | rescue |  | (one eye blocked) | Monocular Saccades |  |  |  |
| --- | --- | --- | --- | --- | --- | --- | --- | --- | --- | --- |
| Wild-type |  | P = 0.348 (ns) | P = 0.161 (ns) | P = 0.356 (ns) | P = 0.210 (ns) | P = 0.007 (**) | P = 0.079 (ns) | P = 2.118 x 10<sup>-4</sup> (***) | P = 0.006 (**) | P = 1.690 x 10<sup>-3</sup> (**) |
| *norpA* Rh1 rescue<br>Binocular Saccades |  |  | P = 0.891 (ns) | P = 0.824 (ns) | P = 0.98 (ns) | P = 0.170 (ns) | P = 0.535 (ns) | P = 0.064 (ns) | P = 0.234 (ns) | P = 0.103 (ns) |
| R7/8 pooled |  |  |  |  |  | P = 0.013 (*) | P = 0.137 (ns) | P = 7.484 x 10<sup>-4</sup> (***) | P = 0.013 (*) | P = 1.70 x 10<sup>-3</sup> (**) |
| *norpA* Rh3-6 rescue |  |  |  |  | P = 0.676 (ns) | P = 0.038 (*) | P = 0.154 (ns) | P = 0.004 (**) | P = 0.027 (*) | P = 0.007 (**) |
| *ninaE*<sup>8</sup> |  |  |  |  |  | P = 0.068 (ns) | P = 0.205 (ns) | P = 0.013 (*) | P = 0.049 (*) | P = 0.013 (*) |
| WT Monocular (one eye blocked) |  |  |  |  |  |  | P = 0.513 (ns) | P = 0.882 (ns) | P = 0.639 (ns) | P = 0.832 (ns) |
| *norpA* Rh1 rescue<br>Monocular Saccades |  |  |  |  |  |  |  | P = 0.407 (ns) | P = 0.644 (ns) | P = 0.294 (ns) |
| Blind pooled |  |  |  |  |  |  |  |  |  |  |
| Blind *hdc*<sup>JK910</sup> |  |  |  |  |  |  |  |  |  | P = 0.400 (ns) |
| Blind *norpA* |  |  |  |  |  |  |  |  |  |  |

**Table S17.**
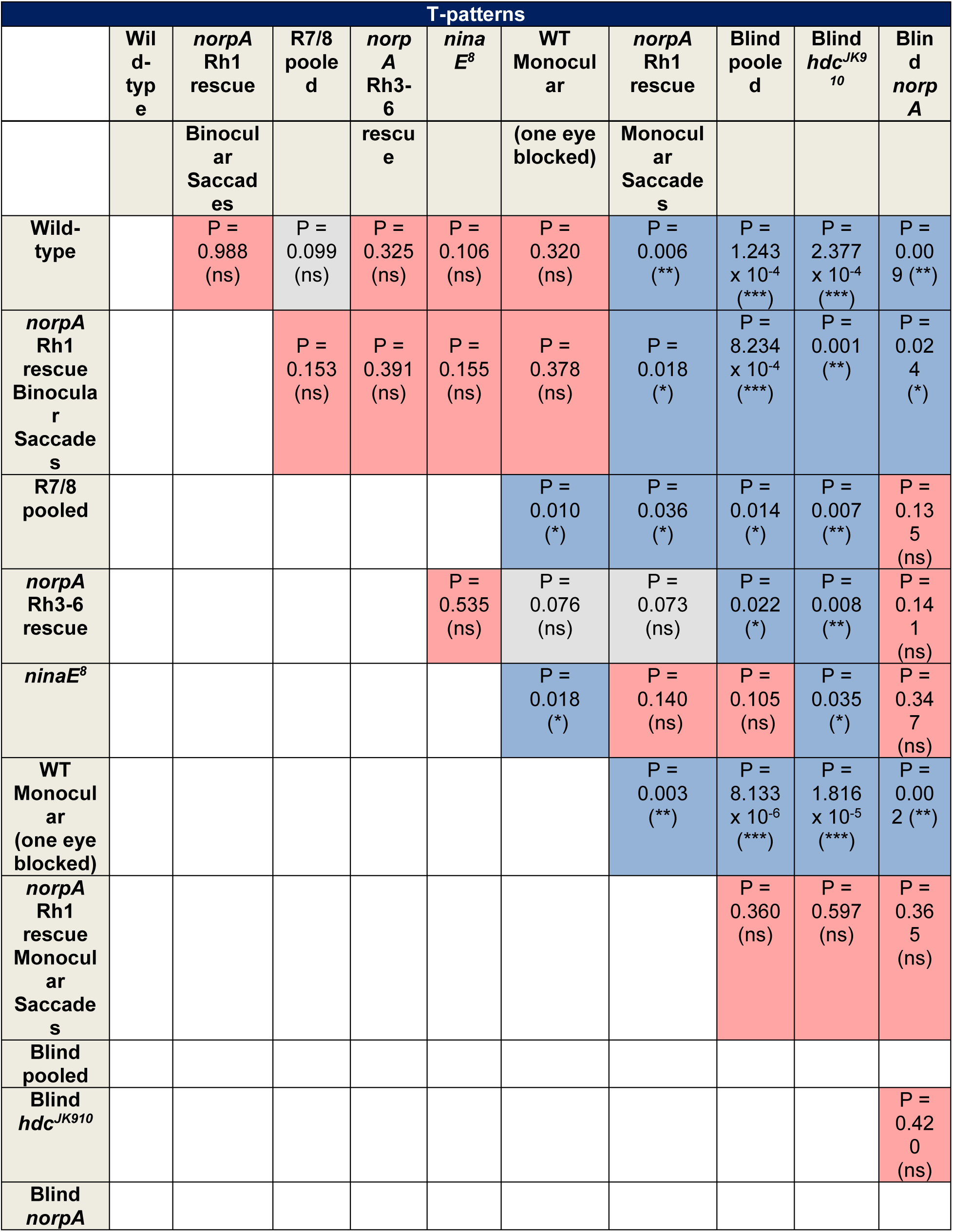

T-patterns WT Monocular had a greater PI than any other group. So, the significant differences found with R7/8 pooled and *ninaE^8^* are due to WT monocular learning better, not the other way around.

**Fig. S77.**
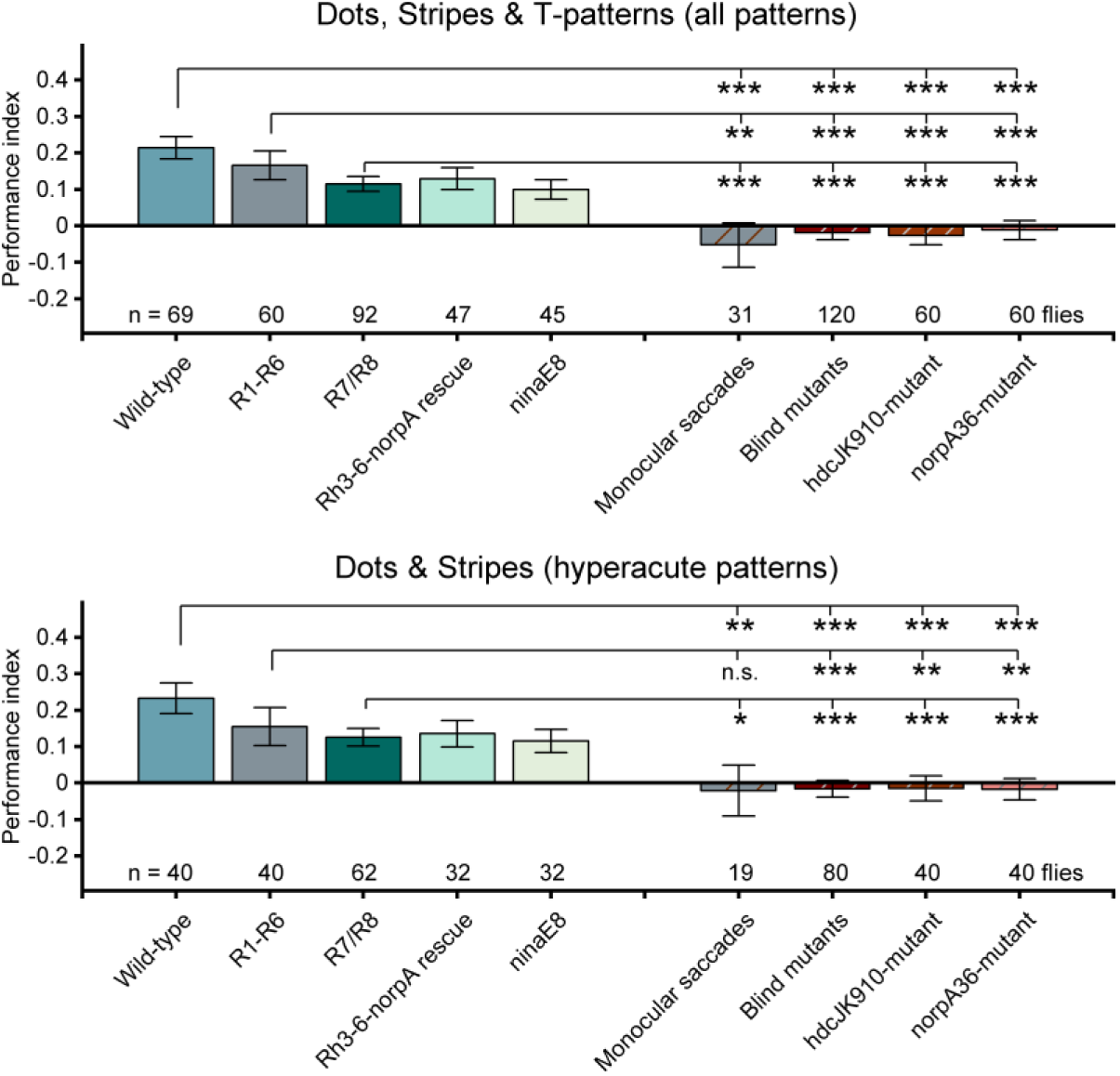
Comparing the different fly geno- and phenotypes’ stage-8 aversive performance learning indexes for the pooled 2D- and 3D-stimuli (above) and pooled hyperacute 3D- objects (below).

**Table S18.**
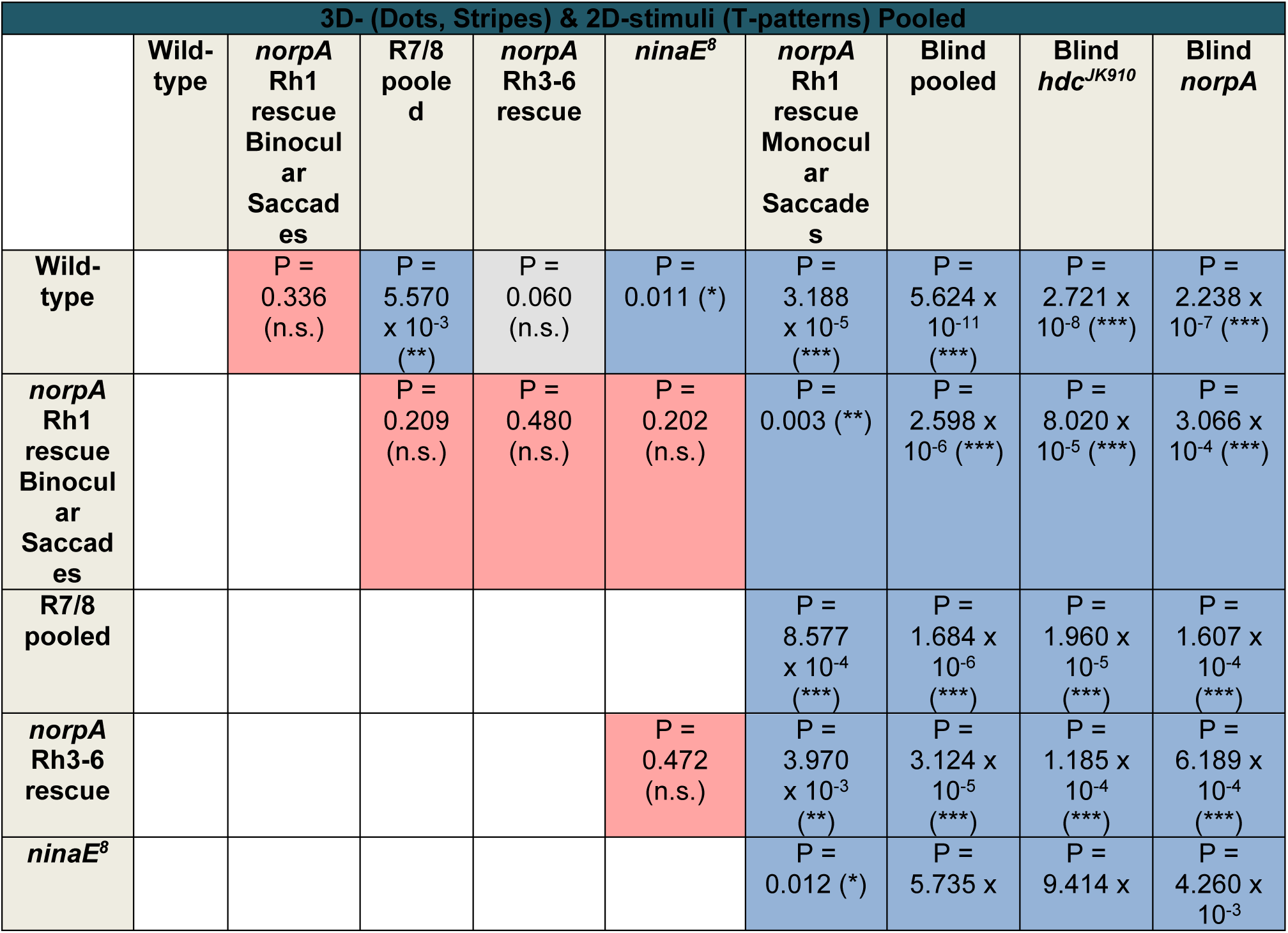
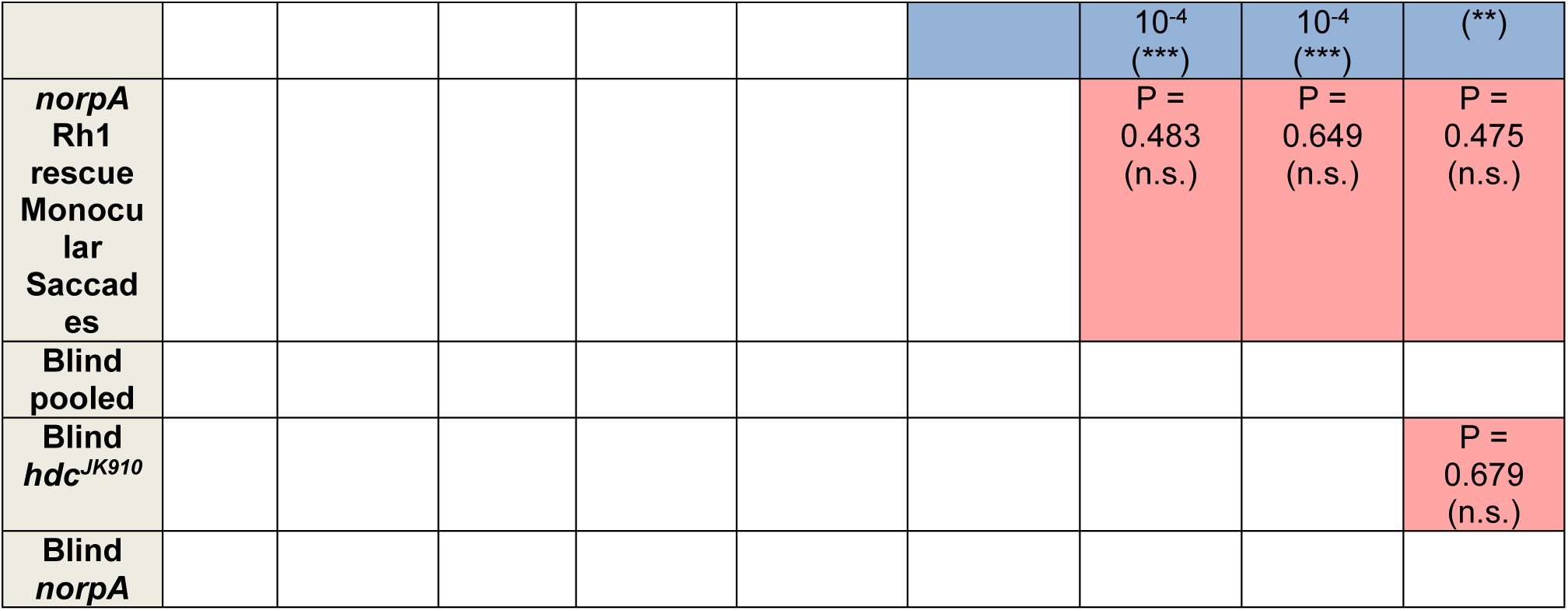

**Table S19.**
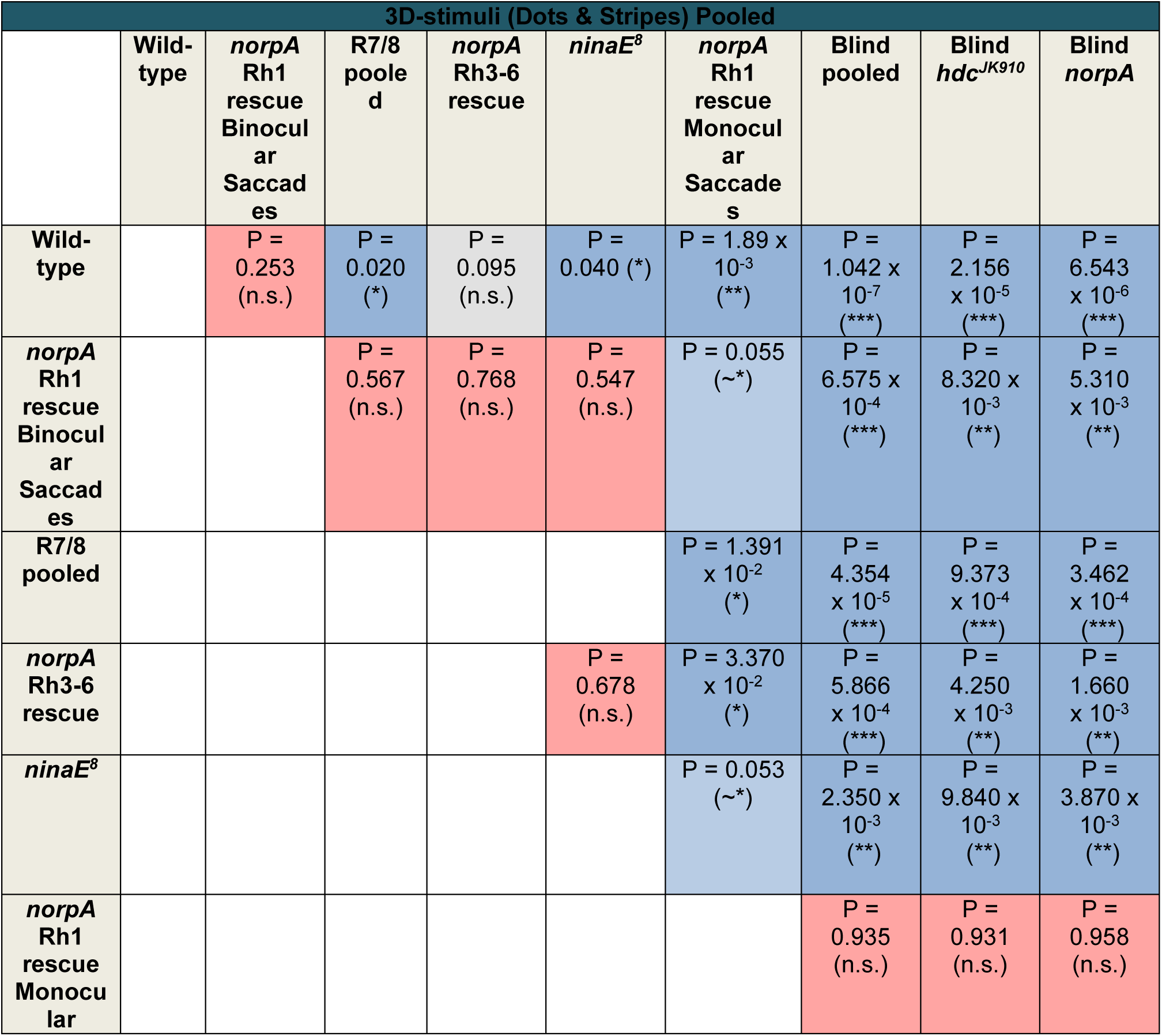
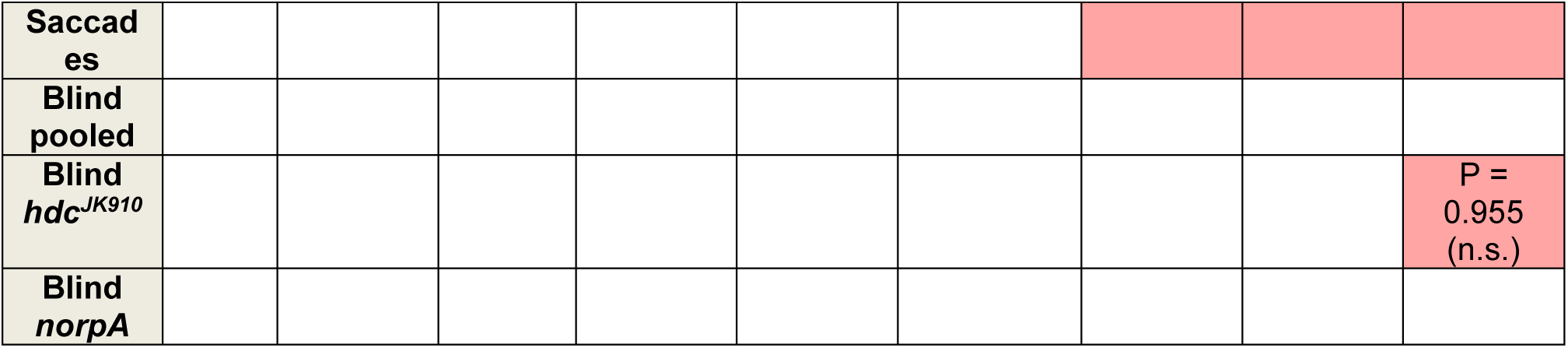

#### VIII Drosophila Genetics

Blind *hdc^JK910^ mutant flies*. *hdc^JK910^* photoreceptors have normal phototransduction but cannot synthesize their neurotransmitter, histamine. Non-functional histidine decarboxylase of *hdc^JK910^* mutants prevents neurotransmitter histamine synthesis in photoreceptors (125, 126). Therefore, their electroretinograms (ERGs) lack On- and Off-transients (125, 126), associated with synaptic light information transfer to visual interneurons, LMCs (19, 20). *hdc^JK910^* flies were received from Erich Buchner’s lab (Julius-Maximilians-Universität, Würzburg, Germany).

Blind *trp;trpl* null-mutants express normal phototransduction reactants but lack their light-gated ion channels completely. These photoreceptors cannot generate electrical responses to light, showing zero-ERG signal, but they contract photomechanically (10, 11). These dynamics are consistent with the hypothesis of the light-induced phosphatidylinositol 4,5-bisphosphate (PIP_2_) cleaving from the microvillar photoreceptor plasma membrane causing the rhabdomere contractions (10).

Blind *norpA^P24^ mutant flies*. *norpA^P24^* is a protein-null mutant of phospholipase C required for phototransduction. The mutation involves a 28-bp deletion that causes a reading frameshift, resulting in the substitution of 24 amino acids followed by a premature truncation of the protein (127). Thus, the mutants are essentially completely blind.

The UV-flies were generated using rhodopsin ninaE8, also known as Rh1, with rescued UV-rhodopsin (Rh3) insertion. The *ninaE^8^* (*ninaE^P334^*) mutation reduces the expression of the rhodopsin ninaE to 0.0004% of wild-type levels (22, 128). This particular mutation was chosen as some level of expression of ninaE is required for normal rhabdomere development (129).

The fused rhabdom line: w; *spam*^1^/*spam*^1^ Frt; sqh-GFP/Tm6B was a gift from Andrew Zelhof.

***Transgenic Rhodopsin-specific norpA rescue flies.*** Flies with functional R1-R6 were generated by crossing wild-type flies bearing a P element containing *norpA* cDNA under an Rh1 promoter (P*[Rh1+norpA]*) with a *norpA^36^* mutant (22). Rh3, Rh4, Rh5, Rh6-specific norpA rescue flies, described in (130), were used to generate flies with functional pale R7, yellow R7, pale R8, and yellow R8 by crossing with a *norpA^36^* mutant, respectively.

***Flies for 2-photon imaging.*** The UV-fly genotype used in 2-photon Ca^2+^-imaging was UAS-GCaMP6f/CyO; L2-Gal4, UV/TM6B and UAS-GCaMP6; L2-Gal4, UV/TM6B. Origins of its different parts: R1-R6 photoreceptor UV-sensitivity resulted from P(Rh1:Rh3)[4303],ninaE[8]/TM6B, see supplementary material (22). L2-Gal4 was 21D-Gal4, a gift from Martin Heisenberg (131). 21-Gal4 insertion was recombined to chromosome III together with the UV genetic set P(Rh1:Rh3)[4303],ninaE[8], using our UV- line stock and the 21D-Gal4 insertion line. The resulting lines were crossed to UAS-CD8-GFP and tested for GFP presence in L2 neurons using fluorescence microscopy. The presence of the UV genetic set was verified in positive lines by ERG testing for UV sensitivity (22). UAS-GCaMP6f was BS46747 P[20xUAS-IVS-GCaMP6f] at P40 2L. Their eyes’ structural integrity and photoreceptor microsaccade dynamics were found to be within the normal range), as tested with the goniometric deep pseudopupil imaging system (see **Section II.** above).

| Genotype | Experimental methods |  |  |  |  |  | Results |  |
| --- | --- | --- | --- | --- | --- | --- | --- | --- |
|  | X-ray imaging | Pseudopupil imaging | Direct R1-R7/8 imaging | ERG | 2-photon imaging | Flight simulator | R1-R7/8 Micro-saccades | Hyper-acute stereo vision |
| Berlin wild-type | ✓ | ✓ | ✓ | ✓ | no | ✓ | ✓ | ✓ |
| *hdc<sup>JK910</sup>* (blind) | ✓ | ✓ | ✓ | ✓ | no | ✓ | ✓ | no |
| *norpA*<sup>P24</sup><br>(blind) | ✓ | ✓ | ✓ | ✓ | no | ✓ | no | no |
| *trp;trpl* (blind) | ✓ | ✓ | ✓ | ✓ | no | ✓ | ✓ | no |
| [Rh1+*norpA*] | no | ✓ | ✓ | ✓ | no | ✓ | ✓ | ✓ |
| [Rh3+*norpA*] | no | ✓ | no | ✓ | no | no | ✓ | - |
| [Rh4+*norpA*] | no | ✓ | no | ✓ | no | no | ✓ (weak) | - |
| [Rh5+*norpA*] | no | ✓ | no | ✓ | no | no | ✓ | - |
| [Rh6+*norpA*] | no | ✓ | no | ✓ | no | no | ✓ | - |
| [Rh3-6+*norpA*] | no | ✓ | no | ✓ | no | ✓ | ✓ | ✓ |
| *ninaE*<sup>8</sup> (R1-R6<br>blind; R7/R8<br>functional) | no | ✓ | no | ✓ | no | ✓ | ✓ | ✓ |
| UV-flies<br>[Rh3(in R1-R6)<br>+*ninaE*<sup>8</sup>] | no | ✓ | ✓ | ✓ | no | no | ✓ | ✓ |
| UV-flies with<br>gCaMP6f in L2 | no | ✓ | ✓ | ✓ | ✓ | no | ✓ | ✓ |
| Canton-S with<br>gCaMP6f in L2 | no | ✓ | ✓ | ✓ | ✓ | no | ✓ | ✓ |
| *Spam* (R1-R8<br>rhabdomeres<br>fused) | ✓ | ✓ | no | ✓ | no | no | ✓ | - |
| *dSK* | no | ✓ | no | ✓ | no | no | ✓ | - |

Berlin wild-type *Drosophila* showed consistent photoreceptor microsaccade dynamics in X-ray, deep pseudopupil, and direct R1-R7/8 high-speed imaging, and hyperacute stereo vision in associative learning experiments.

## Glossary

| Parameter or abbreviation | Definition | Value | Data source or reference |
| --- | --- | --- | --- |
| ESRF | European Synchrotron Research Facility, Grenoble, France |  |  |
| DESY | Deutsches Elektronen-Synchrotron Hamburg, Germany |  |  |
| ERG | electroretinogram |  |  |
| LMCs | Large monopolar cells |  |  |
| KB | Kirkpatrick-Baez |  |  |
| $R(x, y)$ | Two-dimensional cross-correlation | | Eq. 1 |
| $T(x, y)$ | Template image | | Eq. 1 |
| $I(x, y)$ | Source image | | Eq. 1 |
| $T'(x', y')$ | Normalized template image | | Eq. 2 |
| $I'(x', y')$ | Normalized source image | | Eq. 3 |
| $D$ | Displacement | | Eq. 4 |
| R1-R8 | Photoreceptors 1-8 in an ommatidium |  |  |
| DPP | Deep pseudopupil |  |  |
| NA | Numerical aperture |  | Eq. 7 |
| $n$ | Refractive index | | Eq. 7 |
| $\theta$ | Half angle | | Eq. 7 |
| N | f-stop |  | Eq. 8 |
| $f$ | Focal length | | Eq. 8 |
| D | Lens diameter |  | Eq. 8 |
| CG | Computer graphics |  |  |
| $T(f)$ | Transfer function | | Eq. 10 |
| $c(t), C(f)$ | Contrast stimulus in time, frequency | | Eq. 10 |
| $s(t), S(f)$ | Signal in time, frequency | | Eq. 10 |
| Rh1,3,4,5,6 | Rhodopsin 1,3,4,5,6 |  |  |
| L2 | Large monopolar cell type 2 |  |  |
| F | Fluorescence |  |  |
| $F_0$ | Background fluorescence | | |
| $\Delta F$ | Fluorescence difference | | |
| R | Rayleigh criterion |  | Eq. 11 |
| $T$ | Trough amplitude | | Eq. 11 |
| $P_{min}$ | Smallest peak amplitude | | Eq. 11 |
| $P_{max}$ | Highest peak amplitude | | Eq. 11 |
| $\lambda(t)$ | Grating stimulus wavelength | | Eq. 12-13 |
| $s$ | Grating stimulus speed | | Eq. 12-13 |
| $\theta$ | Grating stimulus motion direction | | Eq. 12-13 |
| $\lambda_0$ | Grating stimulus initial wavelength | | Eq. 12-13 |
| $\lambda_1$ | Grating stimulus final wavelength | | Eq. 12-13 |
| SRA | Smallest resolved angle |  |  |
| SNR | Signal-to-noise ratio |  |  |
| $\alpha_{min}$ | Minimum inter-bar distance for SRA | | Eq. 15 |
| $\omega$ | Stimulus motion speed | | Eq. 15 |
| $f_s$ | Sampling rate | | Eq. 15 |
| ROI | Region of interest |  |  |
| $\theta$ | The incident light angle between the light point source, $p$, and the lens center axis | | |
| $p$ | Light point source | | |
| $\lambda$ | Light wavelength | 450 nm (set in simulations) | |
| $k_0$ | Wavenumber | | |
| $\mathbf{r}$ | (x,y,z) position | | Eq. 15 |
| $E_\omega(\mathbf{r})$ | Complex electrical field | | Eq. 15 |
| $n^2(\mathbf{r})$ | Refractive index | | Eq. 15 |
| $\Psi(\mathbf{r})$ | Slowly varying electrical field | | Eq. 18 |
| $n_0$ | Average refractive index | | Eq. 18 |
| $\Delta z$ | Distance step | | Eq. 20 |
| $\mathbf{M}(z)$ | Electrical field in x,y plane | | Eq. 21 |
| $k_x, k_y$ | Wavenumber in x,y-direction | | Eq. 21 |
| $\kappa$ | Material absorbance | | Eq. 22 |
| $P_{abs}$ | Total absorbed power | | Eq. 23 |
| $P_p$ | Total absorbed light flux | | Eq. 24 |
| $\Lambda$ | Optical distance | | Eq. 25 |
| $s_i$ | Ray travel distance | | Eq. 25 |
| $x', y'$ | Rays x,y position | | Eq. 25 |
| $\Delta s_l^\perp$ | Rays relative power | | Eq. 26 |
| $l$ | Ray index | | Eq. 26 |
| $\Delta s_l$ | Rays' area | | Eq. 26 |
| $\theta$ | Ray's angle compared to the z-axis | | Eq. 26 |
| $\vert E_l^{ray}\vert $ | Rays relative | | Eq. 27 |
| $\mathbf{M}_0$ | Initial electrical field strength | | Eq. 28 |
| | Ommatidial lens thickness | 8 $\mu\text{m}$ | (75) |
| | Ommatidial lens diameter | 16 $\mu\text{m}$ | (75) |
| | Ommatidial lens outer surface curvature | 11 $\mu\text{m}$ | (75) |
| | Ommatidial lens inner surface curvature | -11 $\mu\text{m}$ | (75) |
|  | Ommatidial lens refractive index | 1.45 | (75) |
| $n_{min}$ | Crystal cone refractive index and outside rhabdomere | 1.34 | (75) |
| $n_{max}$ | Rhabdomere refractive index | 1.363 | (75) |
| | Cone/pigment-cell aperture diameter | 5 $\mu\text{m}$ | (75) |
| | Cone/pigment-cell aperture thickness | 2 $\mu\text{m}$ | (75) |
|  | Cone/pigment-cell aperture total transmittance | 2.8% |  |
| QB | Quantum Bump |  |  |
| $\Gamma$ | Gamma distribution | | Eq. 29 |
| $n_g$ | Gamma distribution parameter | | Eq. 29 |
| $\tau$ | Gamma distribution parameter | | Eq. 29 |
| $Q_{10}$ | Temperature dependency | | |
| LIC | Light-induced current |  |  |
| RF | A photoreceptor's Receptive Field |  |  |
| $x_d$ | Rhabdomere displacement | | Eq. 30 |
| $act(t)$ | Light activation for rhabdomere displacement | | Eq. 30 |
| $H_{act}$ | Light activation half value | 9,000 $\text{ph}/\mu\text{m}^{1/2}$ (fitted) | Eq. 30 |
| $n_{act}$ | Light activation co-operation exponent | 2 (fitted) | Eq. 30 |
| $D_{coef}$ | Maximal positive dampener force | 0.0001 $\mu\text{m}/\text{ms}^2$ (fitted) | Eq. 30 |
| $D_{base}$ | Dampener exponent base | 2 (fitted) | Eq. 30 |
| $D_{exp}$ | Dampener exponent | 3,900 $\mu\text{m}/\text{ms}$ (fitted) | Eq. 30 |
| spring | Spring constant |  | Eq. 30-31 |
| $k_{s0}$ | Base spring constant | 0.0001/$\text{ms}^2$ (fitted) | Eq. 31 |
| $H_{coef}$ | Adjustable spring constant | 0.00115 1/$\text{ms}^2$ (fitted) | Eq. 31 |
| $H_{spring}$ | Half value of spring constant adjustment | 200 ph/ms (fitted) | Eq. 31 |
| $n_{act}$ | The exponent of spring constant adjustment | 1.3 (fitted) | Eq. 31 |
| $\Delta\varphi$ | Interommatidial angle (horizontal distance) | 4.5° (from anatomy)<br>5.1° (from geometry) | Eq. 34, (12)<br>(75) |
| $\Delta\rho_l^s$ | Optical light input RF half-width (acceptance angle) of a *static* (non-moving) rhabdomere | **R1 and R6:**<br>5.0° ± 0.1<br>**R2-R5:**<br>4.5° ± 0.1<br>**R7/R8:**<br>3.12°<br>**R1-R7/8:**<br>~2°-4° (rhabdomeres modeled *at the lens center axis*; in reality, R1-R6 are off-axis) | See **Table S6**<br>See **Table S6**<br>(75) |
| $\Delta\rho_l^d$ | Optical light input RF half-width (acceptance angle) of a *dynamic* (moving) rhabdomere | **R1 and R6:**<br>4.67°<br>**R2-R5:**<br>4.05°<br>**R7/R8:**<br>2.7°<br>**R1-R7/8** (the average)<br>3.5° (from *Drosophila* flight behavior; hence with intact microsaccades) | See **Table S6**<br>(132) |
| $\Delta\rho_v^s$ | Voltage output RF half-width (acceptance angle) of a *static* (non-moving) rhabdomere | **Dark-adapted R2-R5:**<br>6.4° ± 0.4°<br>**Dark-adapted R1 and R6:**<br>7.1° ± 0.4° | |
| $\Delta\rho_v^d$ | Voltage output RF half-width (acceptance angle) of R1-R6 *dynamic* (moving) rhabdomeres | **Dark-adapted R1-R6:**<br>9.65° ± 1.06°<br>8.23° ± 0.54°<br>**Moderately light-adapted R1-R6:**<br>7.70° ± 0.52° | (11)<br>(12)<br>(11) |
| $z$ | Depth | | Eq. 32 and Eq. 37 |
| $k$ | Eye to eye distance | 440 μm | Eq. 32 |
| $\phi$ | Photoreceptor convergence angle | | Eq. 32 and Eq. 38 |
| $z_e$ | Eye radius | 183 μm (from ommatidium lens properties) | Eq. 35 |
| $\phi_0$ | Starting photoreceptor convergence angle | 5.8° | Eq. 38 |
| $\phi_t$ | Speed-dependent exponent for photoreceptor convergence angle | 0.26565 | Eq. 38 |

## Q & A

This section provides brief answers to some common questions about this study.

**Q1.** How robust are the photoreceptor microsaccade dynamics and the given mathematical models upon parameter variations?

**A1:** *In vivo* photomechanical photoreceptor microsaccades are robust and reproducible. Every structurally intact healthy (non-damaged) wild-type fly will show them. While the microsaccade amplitudes show natural variations during repeated light-stimulation, they appear equally sensitive to light pulse stimulation in the dark- and light-adapted flies. And predictably, given their phototransduction origin, their dynamics become faster with light-adaptation, enabling reliable tracking of fast light contrast changes (Fig. S23).

The mathematical photoreceptor microsaccadic sampling models (see Section V. “Multiscale modeling the adaptive optics and photoreceptor signaling”) are robust, generating realistic stochastic response variability to light stimuli by design. In contrast to conventional hierarchical (top-down) neural processing or control architecture models (based on *ad hoc* filtering functions or generalized mathematical operators), these (bottom-up) biophysically realistic photoreceptor models have no free parameters. Instead, they are constructed to replicate a *Drosophila* photoreceptor’s ultrastructure with 30,000 compartmentalized microvilli (photon sampling units) that house phototransduction pathway, plasma-membrane electrophysiology. Consequently, the models accurately reproduce microsaccadic visual information sampling and integration of real photoreceptors, generating realistic voltage responses with realistic information transfer rates over a broad range of stimulus conditions (11, 39).

Moreover, in the current publication, the models further use the measured photoreceptor microsaccade dynamics to ray-trace the rhabdomeres’ receptive fields and the light inputs these encounter in 3D visual space. This information is then directly sampled by the models. Thus, effectively, this approach forms the foundations of a new morphodynamic active sampling theory for *Drosophila* vision. Furthermore, because the same theoretical framework is readily adaptable for other compound eyes, it provides a robust modeling platform for studying how other insects actively sample visual information, stepping away from the prevailing *static* eye assumptions.

Finally, and crucially, because their sampling and response integration processes are based on experimentally determined parameter values, in the used simulations, their outputs were not tuned to, or affected by, arbitrary parameter variations. The key fixed parameters are listed in the Glossary of this supplement. The corresponding phototransduction molecule numbers and their stochastic reaction dynamics are given in our previous publications (40, 111, 133). Please see the following publications for more details about this multiscale modeling approach (11, 39, 40, 59, 111, 133).

**Q2.** What are the functional implications of photoreceptor microsaccades for the fly vision and visual behaviors? Sensory information is obviously a key element of any behavioral control architecture. But what matters, in the end, is the question of how sensory signals are converted into adaptive motor control signals that enable reflex and goal-directed behavior. One would presume that the best spatiotemporal resolution of peripheral sensory signals does not help if along the sensorimotor pathway information transfer is constrained, for instance, due to the inertia of motor systems.

**A2: *As the fundamental information sampling bottleneck, mirror-symmetric microsaccadic photoreceptor sampling*** affects all parts of *Drosophila* vision and visual behaviors: from hyperacute small object detection and tracking, to (in)voluntary head and body movements, to optic flow processing. Following the data processing theorem, it limits optic flow processing as it limits any other (post-sampling) visual task the fly brain performs. Otherwise, as natural selection eschews wasting information, energy, or resources to maintain futile functions, the photoreceptors with a monocular view only (in the sides of the eyes) should be still, not generating microsaccades. Yet, the microsaccades happen there too (Fig. 1 to 3). To oppose these results, one can try to formulate a case that the *all-purpose* microsaccade-induced hyperacute vision is not needed, for example, for optic flow processing. From the viewpoint of a specific motion detector arrangement (*as the conventional static-eye case*), hyperacute visual information would be unnecessary or even detrimental in generating accurate optic-flow-based state estimates or motor commands. However, our results show that *Drosophila also uses such information for optic flow motor control*, as it robustly responds to hyperacute field rotations (Fig. S65 to S67). So hyperacute vision may not be necessary for the conventional static-eye state-of-the-art optic flow models, but in reality, *Drosophila* has undoubtedly evolved to use it (Fig. 6).

The prevailing general concepts and assumed (theoretical) limitations of optic flow processing depend on visual fields sampled and processed by a particular type of motion detectors (Hassenstein-Reichardt, Barlow-Levick, or combinations of these). Most (if not all) of these motion-detection models assume static-eye and processing, whereupon single neurons do only single functions. However, *Drosophila* visual behaviors (Section VII. “Flight simulator experiments”) give ample evidence that these models can only provide coarse approximations of the biological neural networks’ real functions. Such acumen is neither new nor surprising as it is well-known that none of the prevailing models can predict neural responses perfectly, especially when the stimulus conditions change dynamically.

So whilst nobody knows the flies’ real visual perception in flight, the prevailing models assume it to be limited by the compound eyes’ static optics and their photoreceptors’ (underestimated; far too slow) integration time. Yet, these models do not consider the local and global microsaccadic photoreceptor sampling that enhances fast phasic signals, in which motion-direction-sensitive dynamics are introduced in this paper. Nor do they consider stochastic refractory photon sampling that combats spatiotemporal aliasing while further accentuating and fastening the response dynamics (see, for example (11)). In contrast, these processes are integrated into the current study’s multiscale visual information sampling and integration models, predicting and replicating many experimental findings.

**Q3.** How does one know that the photoreceptor microsaccade recordings did not include, or were interfered with, eye-muscle-induced retinal movements?

**A3:** This study focused on *photomechanical photoreceptor microsaccades* on a fast 0-300 ms time-scale and how their local and global sampling dynamics enable hyperacute stereopsis. Therefore, many experiments were designed to eliminate or minimize eye-muscle-induced retinal movements. Besides the clinching *trp*/*trpl*-mutant evidence, proving the microsaccades’ phototransduction origin (Fig. 2*F*), six other consistent results from different assays further fully support this conclusion:

In the used high-speed imaging configurations (Fig. 1 to 3), with the *Drosophila* head immobilized and data collected in short 200-300 ms chunks, muscle-induced retinal movements rarely occurred. Notably, 200-300 ms is also significantly faster than the measured whole retina movement dynamics in darkness and ambient light (Fig. S34). Thus, intrinsic eye-muscle activity did not interfere with the recorded photoreceptor microsaccades.
In high-speed infra-red deep pseudo pupil (DPP) imaging, the rhabdomere tips move laterally during a photoreceptor microsaccade. But they simultaneously also move axially (11), away from the ommatidium lens and back. We measured this axial movement component directly from their DPP images (Fig. S15) as a rapid proportional intensity change (darkening increasing with the rhabdomere tip distance from the recording camera). Importantly, this fast darkening/brightening perfectly time-locks with the lateral DPP movement, meaning that these two microsaccade components are synchronous. In contrast, the DPPs imaged during eye-muscle-induced retinal movements lack this fast co-dynamic completely. Therefore, the DPP microsaccade recordings, which consistently show transiently darkening DPPs, were purely photomechanical (i.e., generated by phototransduction alone (10)).
Photoreceptor microsaccades tracked light contrast modulation with movement dynamics adapting uniquely (*yet similarly to photoreceptor voltage responses*) to temporally accelerating sinusoidal and square-wave contrast patterns (figs S23 and 24). Because these intricate movements show ultrafast light-contrast-dependent adaptive dynamics, they cannot be caused by reflex-like or light-triggered eye-muscle activity moving the whole retina.
Small targeted light-spot stimuli only evoked photoreceptor microsaccades in the ommatidia experiencing incident light (Fig. S33), whilst light-field stimulation evoked the strongest photoreceptor microsaccades in the ommatidia directly facing it (Fig. S32). If these stimuli were triggering the eye-muscles, the whole retina would have moved, not a few photoreceptors only.
The different spectral photoreceptor classes’ microsaccades summed up similarly to their ERG responses (figs S28 to S30; Table S2 to S5), validating their phototransduction origin. Thus, the microsaccades become systematically smaller when only a photoreceptor subclass (say, R7y) functions. In contrast, once triggered, reflex-like eye-muscle-induced retinal movements should be one-size-only. However, such dynamics were never seen in any of the hundreds of individual rhodopsin-rescue flies tested.
Finally, *in vivo* photoreceptor microsaccade dynamics (measured inside single ommatidia by cornea-neutralization microscopy) match the light-induced photoreceptor contractions of isolated *ex vivo* ommatidia preparations (mechanically removed from the *Drosophila* eyes and dissociated in a petri dish (10); e.g., Video 2 in (11)), lacking entirely any eye-muscles.

**Q4.** Franceschini and colleagues (e.g. (54)) have discussed hyperacuity regarding eye-muscle-induced retinal movements, first indirectly studied by Hengstenberg in housefly (*Musca domestica*) (57), to improve the fly compound eyes’ otherwise relatively poor spatial resolution. But do the eye-muscle-induced retinal movements and the photomechanical photoreceptor microsaccades in *Drosophila* differ in terms of the putative function?

**A4:** *Photomechanical photoreceptor microsaccades* sharpen local light-contrast changes near instantaneously within and between neighboring photoreceptor receptive fields (RF, “pixels”) that collectively across one eye comprise its neural image (Fig. S56). Two fundamental ultrafast-adapting optical processes narrow the photoreceptor’s RF during a photomechanical photoreceptor microsaccade, improving acuity. These are: (i) a rhabdomere’s photomechanical axial contraction (see Fig. S15 and S50) and (ii) lateral (sideways) movement (see Fig. S15 and S51). In addition, there is intrinsic “light beam clipping”, regarding the ommatidium cone/pigment-cell aperture (Fig. S52) and the angle a moving object crosses the sideways moving RF (Fig. S61), which dynamically narrow the RF even further. Interestingly, therefore, if the transiently narrowing RF and the object move in the same direction, the photoreceptor has even more time to sample finer (hyperacute) details about the object than when they move in the opposite directions (Fig. S56). These ultrafast photomechanical adaptive optics are further accentuated in time by the stochastic refractoriness of 30,000 rhabdomeric microvilli (photon sampling units; see (11, 39, 40, 59)), in which collective photon samples (quantum bumps) sum up the macroscopic voltage response (Fig. S53). Therefore, the effective photoreceptor integration time is considerably faster, and the resulting temporal resolution (of both the photoreceptor and L2-interneuron signals (Fig. 4)) is much finer, than what is currently assumed for the state-of-the-art static compound eye optics and motion-detection models. Moreover, as predicted by our new theory, L2-interneuron signals show directional hyperacute motion-sensitivity, following the microsaccade movement directions across the two eyes (Fig. 1 to 4).

Conversely, *eye-muscle-induced retinal movements* shift the whole retina and the neural image it is sampling (54) (Fig. S34). Presumably, this action dynamically refreshes the neural image, combating fast adaptation fading the perception (11). Moreover, the eye-muscle-induced retinal movement may actively (54) (and perhaps also attentively - as the vertebrate eyes do), through whole retina saccades, vergence movements, or slowly pulling and pushing retinal tissue inward and outward (Fig. S34F), improve the detection or resolution of visually interesting objects (54). While these vergence movements can happen in a coordinated way in one or the other eye or both (as shown for *Musca* (13, 57)), on top of them, the photomechanical *photoreceptor microsaccades supervene* in their hardwired directionality, leading to complex superimposed spatiotemporal (“super-saccadic”) sampling dynamics. So, whilst the photoreceptor microsaccades can enhance the neural images alone, the eye-muscle-induced whole retina movements never do so in isolation. (Meaning, each retina movement will change its photoreceptors’ light input, evoking photomechanical microsaccades; apart from the situations when a fly is in complete darkness or faces a homogeneous zero-contrast space.)

In the head-immobilized *Drosophila* recordings, photoreceptor microsaccades are fast and time-locked to light-intensity changes. Conversely, the eye-muscle-induced retinal movements happen infrequently and spontaneously, showing no clear stimulus dependency with a much slower time course. However, since adapting information sampling to behaviors must improve fitness (e.g. (11, 59)), one expects the eye-muscle-induced retinal movements to be different in the free-moving (flying or walking) flies and other insect species, as they are evolutionarily tuned to their different behavioral needs (more in A5 below).

However, owing to our experimental focus (see A2 above), we obtained *no direct recordings* to analyze *how photomechanical photoreceptor microsaccades and eye-muscle-induced retinal movements work together* to improve *Drosophila* vision. For example, whilst these two processes likely jointly occur during visual learning behaviors, we could not record photoreceptor movements during the flight simulator experiments (limited by our set-up design). Nonetheless, we performed additional long-lasting, deep pseudopupil (DPP) high-speed imaging experiments to capture eye-muscle-induced retinal movements in head-immobilized *Drosophila* to address Q4 *indirectly*. These results are shown in Section III. “High-speed optical imaging of eye-muscle-induced retina movements and antennae casting.” In brief, it was found that:

Eye-muscle-induced retinal movements happen in darkness (dark-adaptation) and ambient illumination infrequently (Fig. S34). Characteristically, they cause the observed DPPs to drift slowly with much slower temporal dynamics than the photomechanical photoreceptor microsaccades.
Antenna movements, which are sometimes apparent during the high-speed imaging experiments, can only happen 40-50 ms after the photomechanical photoreceptor microsaccades (Fig. S35).
Antenna movements do not induce DPP retinal movements (Fig. S36).

**O5.** In visual ecology, *Drosophila* is characterized as a slow-flying fly. What would one predict the photoreceptor microsaccades to be like in faster flies, such as *Musca* and *Calliphora*, or in other fast-flying insects, which instead of having open-rhabdom neural superposition eyes, often possess fused-rhabdom apposition eyes?

**A5**: As touched upon in A4, because different insects have different visual needs, it is expected that both their *photoreceptor microsaccade* and *eye-muscle-induced retinal movement* dynamics would differ from *Drosophila*’s. But in each case, one would anticipate these dynamics to have adapted to improve the acuity to see the world in motion. Based on the *Drosophila* results (see A4 above), we can try to predict how active sampling might have evolved to shape other insects’ vision, indifference to *Drosophila*.

For fast-moving insects, more ommatidia tile their compound eyes. With more pixels (in resulting neural images) and thus narrower intra-ommatidial angles and receptive field half-widths, one would expect their photoreceptor microsaccades, correspondingly, to be much smaller and faster. This prediction comes directly from the sampling theory: how to integrate the best image by moving sensors. In the case of *Drosophila*, their R1-R6’ RF half-widths are between 4.5-6°, over-completely tiling up their retinotopically mapped visual fields. In proportion, their photoreceptor microsaccades move laterally 1-1.5 µm on average, equating to about 3-4.5° RF movements in the visual space. For an analog here, consider a digital camera sampling an image. The spatial image information doubles when the camera is slightly moved, and two consecutive images are taken a 1/2-pixel part and then time-integrated for enhanced resolution. However, if the photomechanical RF movements extended more, they would eventually superimpose on neighboring RFs (if these were not exposed to light intensity changes). In that case, acuity would decrease with the resulting neural image containing fewer pixels (newly fused sampling points).

Therefore, we predict that *Musca* and *Calliphora* R1-R6 photoreceptors, with 1.0-1,5° RF half-widths, will move laterally 0.5-1.0° in the visual space. And as these flies whoosh around about 4-10-times faster than *Drosophila* and their photoreceptors show 2-3-times higher information transfer rates (12), we would predict their photoreceptor microsaccades also to be 4-10-times faster. In other words, for obtaining the best visual acuity, we would expect *Musca* photoreceptor microsaccades to move maximally ∼100-300 nm sideways with minimal delays (<∼1-2 ms), peaking within 10 ms from the light-stimulus onset.

As for the fast-flying insects with apposition eyes, we expect that their fused rhabdom’s higher structural rigidity (in relation to *Drosophila*’s flexible spatially partitioned rhabdomeres) significantly reduces the lateral photoreceptor microsaccade component. On the other hand, since their rhabdom are often much longer and more distant from the ommatidial lens (e.g., honeybee), we would predict their axial microsaccade component to be much faster and possibly larger than what we see in the relatively short *Drosophila* rhabdomeres.

Moreover, because the fast-flying insects’ photoreceptors and interneurons adapt faster than *Drosophila*’s, their eyes need powerful intrinsic mechanisms to prevent retinal images from fading. Hence, one would expect their eye-muscle-induced retinal movements (vergence sweeps) to be considerably larger and show much faster pulsatile dynamics than in *Drosophila* (*cf*. the clock-spikes in *Musca* (13, 57)).

It will be fascinating to see how these predictions fare in future studies.

## Movie legends

**Movie S1**. *In vivo* X-ray imaging reveals *Drosophila* eyes’ internal structure with the X-ray intensity modulating the retinal displacement. X-rays activate globally the right and left eye’s radially arranged string-like photoreceptors to contract rapidly and mirror-symmetrically in the back-to-front direction.

**Movie S2**. *In vivo* X-ray imaging and ERG-recording the *Drosophila* eyes’ photomechanical photoreceptor dynamics. X-rays activate phototransduction with photoreceptor contractions similar to visible light.

**Movie S3. Mapping *in vivo* the *Drosophila* eyes’ stereoscopic field of view with high-speed deep pseudopupil imaging.**

**Movie S4**. Mapping *in vivo* the photomechanical photoreceptor microsaccade movement directions across the *Drosophila* eyes.

**Movie S5**. Measuring *in vivo* the light-adapted photomechanical photoreceptor microsaccades’ movement dynamics to brief light contrast changes.

**Movie S6. The left and right eyes’ mirror-symmetrically moving photoreceptor receptive fields match a forward-flying Drosophila’s corresponding optic flow field to enhance information capture.**

**Movie S7**. **During yaw rotation, the left and right eyes’ mirror-symmetrically moving photoreceptor receptive fields enhance binocular contrast differences in the world.**

**Movie S8**. *In vivo* two-photon imaging of L2 monopolar cells’ medulla terminals reveals their hyperacute receptive field organization along with the photoreceptor microsaccade movement maps.

**Movie S9**. The corresponding left and right eye R6 photoreceptor cells’ receptive fields move with and against an object that crosses them, providing dynamic depth information to the *Drosophila* brain.

**Movie S10**. **Theory of stereoscopic information sampling by the *Drosophila* eyes**. Simulations show how the binocular left and right photoreceptor cells’ receptive fields feed dynamic depth information to the *Drosophila* brain about the distance of close-by and further away objects of the same angular size.

## Notes

### Competing Interest Statement

The authors have declared no competing interest.

## References

1. D. H. Hubel, T. N. Wiesel, Receptive fields, binocular interaction and functional architecture in the cat’s visual cortex. J Physiol 160, 106–154 (1962).

2. M. Livingstone, D. Hubel, Segregation of form, color, movement, and depth: anatomy, physiology, and perception. Science 240, 740–749 (1988).

3. V. Nityananda, C. Joubier, J. Tan, G. Tarawneh, J. C. A. Read, Motion-in-depth perception and prey capture in the praying mantis Sphodromantis lineola. J Exp Biol 222 (2019).

4. V. Nityananda, J. C. A. Read, Stereopsis in animals: evolution, function and mechanisms. J Exp Biol 220, 2502–2512 (2017).

5. G. F. Poggio, B. C. Motter, S. Squatrito, Y. Trotter, Responses of neurons in visual cortex (V1 and V2) of the alert macaque to dynamic random-dot stereograms. Vision research 25, 397–406 (1985).

6. R. Rosner, J. von Hadeln, G. Tarawneh, J. C. A. Read, A neuronal correlate of insect stereopsis. Nat Commun 10, 2845 (2019).

7. R. F. van der Willigen, B. J. Frost, H. Wagner, Stereoscopic depth perception in the owl. Neuroreport 9, 1233–1237 (1998).

8. M. F. Land, Visual acuity in insects. Annu Rev Entomol 42, 147–177 (1997).

9. S. B. Laughlin, The role of sensory adaptation in the retina. J Exp Biol 146, 39–62 (1989).

10. R. C. Hardie, K. Franze, Photomechanical responses in *Drosophila* photoreceptors. Science 338, 260–263 (2012).

11. M. Juusola et al., Microsaccadic sampling of moving image information provides *Drosophila* hyperacute vision. Elife 6 (2017).

12. P. T. Gonzalez-Bellido, T. J. Wardill, M. Juusola, Compound eyes and retinal information processing in miniature *dipteran* species match their specific ecological demands. P Natl Acad Sci USA 108, 4224–4229 (2011).

13. N. Franceschini, R. Chagneux, K. Kirschfeld, A. Mucke, “Vergence eye movements in flies” in Gottingen Neurobiology Report: Synapse - Transmission Modulation, N. Elsner, H. Penzlin, Eds. (Georg Thieme Verlag, Stuttgard, New York, 1991), pp. 1.

14. N. Franceschini, Combined optical, neuroanatomical, electrophysiological and behavioural studies on signal processing in the fly compound eye. Ser Biophys Biocyber 2, 341–361 (1997).

15. L. E. Lipetz, The mechanism of the X-ray phosphene. Radiat Res 1, 551–551 (1954).

16. L. E. Lipetz, Electrophysiology of the X-ray phosphene. Radiat Res 2, 306–329 (1955).

17. C. M. Avakjan, “Contribution to the theory of light impression on the retina under the influence of X-ray” in Electroretinographia. (University Brne, Lekarska Fakulta 1959), pp. 105–108.

18. K. D. Steidley, The radiation phosphene. Vision research 30, 1139–1143 (1990).

19. P. E. Coombe, The large monopolar cells L1 and L2 are responsible for ERG transients in *Drosophila*. J Comp Physiol A 159, 655–665 (1986).

20. A. Dau et al., Evidence for dynamic network regulation of *Drosophila* photoreceptor function from mutants lacking the neurotransmitter histamine. Front Neural Circuit 10 (2016).

21. N. Franceschini, K. Kirschfeld, Phenomena of pseudopupil in compound eye of *Drosophila*. Kybernetik 9, 159–182 (1971).

22. T. J. Wardill et al., Multiple spectral inputs improve motion discrimination in the *Drosophila* visual system. Science 336, 925–931 (2012).

23. B. Pick, Specific misalignments of rhabdomere visual axes in neural superposition eye of *dipteran* flies. Biol Cybern 26, 215–224 (1977).

24. N. Franceschini, “Pupil and pseudopupil in the compound eye of Drosophila” in Information processing in the visual systems of Anthropods R. Wehner, Ed. (Springer-Verlag, Berlin, Heidelberg, New York, 1972), pp. 75–82.

25. A. Hyvärinen, J. Hurri, P. O. Hoyer, Natural image statistics: a probabilistic approach to early computational vision (2009), vol. 39, pp. 1–448.

26. H. G. Krapp, R. Hengstenberg, Estimation of self-motion by optic flow processing in single visual interneurons. Nature 384, 463–466 (1996).

27. Y. E. Fisher, M. Silies, T. R. Clandinin, Orientation selectivity sharpens motion detection in *Drosophila*. Neuron 88, 390–402 (2015).

28. M. Silies et al., Modular use of peripheral input channels tunes motion-detecting circuitry. Neuron 79, 111–127 (2013).

29. M. Joesch, B. Schnell, S. V. Raghu, D. F. Reiff, A. Borst, ON and OFF pathways in *Drosophila* motion detection. Neuroforum 17, 30–32 (2011).

30. M. Rivera-Alba et al., Wiring economy and volume exclusion determine neuronal placement in the *Drosophila* brain. Curr Biol 22, 172–172 (2012).

31. J. C. Tuthill, A. Nern, S. L. Holtz, G. M. Rubin, M. B. Reiser, Contributions of the 12 neuron classes in the fly lamina to motion vision. Neuron 79, 128–140 (2013).

32. I. A. Meinertzhagen, S. D. O’Neil, Synaptic organization of columnar elements in the lamina of the wild-type in *Drosophila melanogaster*. J Comp Neurol 305, 232–263 (1991).

33. A. Nikolaev et al., Network adaptation improves temporal representation of naturalistic stimuli in *Drosophila* eye: II mechanisms. Plos One 4 (2009).

34. H. H. Yang et al., Subcellular imaging of voltage and calcium signals reveals neural processing *in vivo*. Cell 166, 245–257 (2016).

35. L. Zheng et al., Network adaptation improves temporal representation of naturalistic stimuli in *Drosophila* eye: I dynamics. Plos One 4 (2009).

36. L. Zheng et al., Feedback network controls photoreceptor output at the layer of first visual synapses in *Drosophila*. J Gen Physiol 127, 495–510 (2006).

37. H. J. W. M. Hoekstra, On beam propagation methods for modelling in integrated optics. Opt Quant Electron 29, 157–171 (1997).

38. S. Schuster, R. Strauss, K. G. Gotz, Virtual-reality techniques resolve the visual cues used by fruit flies to evaluate object distances. Curr Biol 12, 1591–1594 (2002).

39. Z. Song, M. Juusola, Refractory sampling links efficiency and costs of sensory encoding to stimulus statistics. J Neurosci 34, 7216–7237 (2014).

40. Z. Song et al., Stochastic, adaptive sampling of information by microvilli in fly photoreceptors. Curr Biol 22, 1371–1380 (2012).

41. M. Courgeon, C. Desplan, Coordination between stochastic and deterministic specification in the *Drosophila* visual system. Science 366, 325–336 (2019).

42. S. Tang, M. Juusola, Intrinsic activity in the fly brain gates visual information during behavioral choices. Plos One 5 (2010).

43. K. Farrow, J. Haag, A. Borst, Nonlinear, binocular interactions underlying flow field selectivity of a motion-sensitive neuron. Nat Neurosci 9, 1312–1320 (2006).

44. J. Haag, A. Borst, Electrical coupling of lobula plate tangential cells to a heterolateral motion-sensitive neuron in the fly. J Neurosci 28, 14435–14442 (2008).

45. R. Rosner, G. Tarawneh, V. Lukyanova, J. C. A. Read, Binocular responsiveness of projection neurons of the praying mantis optic lobe in the frontal visual field. J Comp Physiol A 206, 165–181 (2020).

46. T. Fujiwara, T. L. Cruz, J. P. Bohnslav, M. E. Chiappe, A faithful internal representation of walking movements in the *Drosophila* visual system. Nat Neurosci 20, 72–81 (2017).

47. N. Boeddeker, R. Kern, M. Egelhaaf, Chasing a dummy target: smooth pursuit and velocity control in male blowflies. Proc Biol Sci 270, 393–399 (2003).

48. S. N. Fry, R. Wehner, Look and turn: landmark-based goal navigation in honey bees. J Exp Biol 208, 3945–3955 (2005).

49. S. Tang, R. Wolf, S. P. Xu, M. Heisenberg, Visual pattern recognition in *Drosophila* is invariant for retinal position. Science 305, 1020–1022 (2004).

50. B. Z. Kacsoh, Z. R. Lynch, N. T. Mortimer, T. A. Schlenke, Fruit flies medicate offspring after seeing parasites. Science 339, 947–950 (2013).

51. E. Buchner, Elementary movement detectors in an insect visual-system. Biological Cybernetics 24, 85–101 (1976).

52. M. F. Land, Motion and vision: why animals move their eyes. J Comp Physiol A 185, 341–352 (1999).

53. S. Pick, R. Strauss, Goal-driven behavioral adaptations in gap-climbing *Drosophila*. Curr Biol 15, 1473–1478 (2005).

54. L. Kerhuel, S. Viollet, N. Franceschini, The VODKA sensor: a bio-inspired hyperacute optical position sensing device. IEEE Sensors J 12, 315–324 (2012).

55. V. P. Pandiyan et al., The optoretinogram reveals how human photoreceptors deform in response to light. bioRxiv https://doi.org/10.1101/2020.01.18.911339 (2020).

56. U. Bocchero et al., Mechanosensitivity is an essential component of phototransduction in vertebrate rods. Plos Biology 18 (2020).

57. R. Hengstenberg, Eye muscle system of housefly *Musca-Domestica* .1. Analysis of clock spikes and their source. Kybernetik 9, 56–77 (1971).

58. B. J. Hardcastle, H. G. Krapp, Evolution of Biological Image Stabilization. Curr Biol 26, R1010–R1021 (2016).

59. M. Juusola, Z. Song, How a fly photoreceptor samples light information in time. J Physiol-London 595, 5427–5437 (2017).

60. M. Juusola, R. C. Hardie, Light adaptation in *Drosophila* photoreceptors: I. Response dynamics and signaling efficiency at 25 degrees C. J Gen Physiol 117, 3–25 (2001).

61. R. C. Hardie, A histamine-activated chloride channel involved in neurotransmission at a photoreceptor synapse. Nature 339, 704–706 (1989).

62. A. Pantazis et al., Distinct roles for two histamine receptors (hclA and hclB) at the *Drosophila* photoreceptor synapse. J Neurosci 28, 7250–7259 (2008).

63. M. Juusola, R. O. Uusitalo, M. Weckstrom, Transfer of graded potentials at the photoreceptor interneuron synapse. J Gen Physiol 105, 117–148 (1995).

64. A. C. Zelhof, R. W. Hardy, A. Becker, C. S. Zuker, Transforming the architecture of compound eyes. Nature 443, 696–699 (2006).

65. J. Frohn et al., 3D virtual histology of human pancreatic tissue by multiscale phase-contrast X-ray tomography. J Synchrotron Radiat 27, 1707–1719 (2020).

66. T. Salditt et al., Compound focusing mirror and X-ray waveguide optics for coherent imaging and nano-diffraction. J Synchrotron Radiat 22, 867–878 (2015).

67. S. P. Krüger et al., Sub-10 nm beam confinement by X-ray waveguides: design, fabrication and characterization of optical properties. J Synchrotron Radiat 19, 227–236 (2012).

68. L. M. Lohse et al., A phase-retrieval toolbox for X-ray holography and tomography. Journal of Synchrotron Radiation 27, 852–859 (2020).

69. N. Franceschini, K. Kirschfeld, Optical study *in vivo* of photoreceptor elements in compound eye of *Drosophila*. Kybernetik 8, 1–13 (1971).

70. R. Petrowitz, H. Dahmen, M. Egelhaaf, H. G. Krapp, Arrangement of optical axes and spatial resolution in the compound eye of the female blowfly Calliphora. J Comp Physiol A 186, 737–746 (2000).

71. M. Egelhaaf et al., Neural encoding of behaviourally relevant visual-motion information in the fly. Trends Neurosci 25, 96–102 (2002).

72. J. W. Aptekar et al., Method and software for using m-sequences to characterize parallel components of higher-order visual tracking behavior in Drosophila. Front Neural Circuits 8, 130 (2014).

73. R. Wolf, M. Heisenberg, Vision in Drosophila: Genetics of Microbehavior (Springer-Verlag, Berlin; Heidelberg; New York, NY, 1984).

74. R. C. Hardie, M. Juusola, Phototransduction in *Drosophila*. Curr Opin Neurobiol 34, 37–45 (2015).

75. D. G. Stavenga, Angular and spectral sensitivity of fly photoreceptors. II. Dependence on facet lens F-number and rhabdomere type in *Drosophila*. J Comp Physiol A 189, 189–202 (2003).

76. M. Spencer, Fundamentals of light microscopy, IUPAB biophysics series (Cambridge University Press, Cambridge Cambridgeshire ; New York, 1982), pp. x, 93 p.

77. M. Juusola, A. Dau, L. Zheng, D. N. Rien, Electrophysiological method for recording intracellular voltage responses of *Drosophila* photoreceptors and interneurons to light stimuli *in vivo*. Jove-J Vis Exp ARTN e54142 10.3791/54142 (2016).

78. X. F. Li et al., Ca^2+^-activated K^+^ channels reduce network excitability, improving adaptability and energetics for transmitting and perceiving sensory information. J Neurosci 39, 7132–7154 (2019).

79. A. N. Abou Tayoun et al., The *Drosophila* SK channel (*dSK*) contributes to photoreceptor performance by mediating sensitivity control at the first visual network. J Neurosci 31, 13897–13910 (2011).

80. S. R. Henderson, H. Reuss, R. C. Hardie, Single photon responses in *Drosophila* photoreceptors and their regulation by Ca^2+^. J Physiol-London 524, 179–194 (2000).

81. P. Hochstrate, K. Hamdorf, Microvillar components of light adaptation in blowflies. J Gen Physiol 95, 891–910 (1990).

82. M. Juusola, R. C. Hardie, Light adaptation in *Drosophila* photoreceptors: II. Rising temperature increases the bandwidth of reliable signaling. J Gen Physiol 117, 27–41 (2001).

83. M. Juusola, Linear and nonlinear contrast coding in light-adapted blowfly photoreceptors. J Comp Physiol A 172, 511–521 (1993).

84. M. Juusola, G. G. De Polavieja, The rate of information transfer of naturalistic stimulation by graded potentials. J Gen Physiol 122, 191–206 (2003).

85. M. E. Fortini, G. M. Rubin, The optic lobe projection pattern of polarization-sensitive photoreceptor cells in *Drosophila melanogaster*. Cell Tissue Res 265, 185–191 (1991).

86. P. Virtanen et al., SciPy 1.0: fundamental algorithms for scientific computing in Python. Nat Methods 17, 261–272 (2020).

87. S. Seabold, J. Perktold (2010) Statsmodels: econometric and statistical modeling with Python. In Proceedings of the 9th Python in Science Conference (SCIPY 2010), pp 92–96.

88. B. Minke, The history of the prolonged depolarizing afterpotential (PDA) and Its role in genetic dissection of *Drosophila* phototransduction. J Neurogenet 26, 106–117 (2012).

89. M. E. Chiappe, J. D. Seelig, M. B. Reiser, V. Jayaraman, Walking modulates speed sensitivity in *Drosophila* motion vision. Curr Biol 20, 1470–1475 (2010).

90. J. D. Seelig, M. E. Chiappe, G. K. Lott, M. B. Reiser, V. Jayaraman, Calcium imaging in *Drosophila* during walking and flight behavior. Biophys J 100, 97–97 (2011).

91. J. H. van Hateren, Neural superposition and oscillations in the eye of the blowfly. J Comp Physiol A 161, 849–855 (1987).

92. J. B. Shi, C. Tomasi, Good features to track. 1994 Ieee Computer Society Conference on Computer Vision and Pattern Recognition, Proceedings Doi 10.1109/Cvpr.1994.323794, 593-600 (1994).

93. D. G. Stavenga, Angular and spectral sensitivity of fly photoreceptors. I. Integrated facet lens and rhabdomere optics. J Comp Physiol A 189, 1–17 (2003).

94. D. G. Stavenga, Angular and spectral sensitivity of fly photoreceptors. III. Dependence on the pupil mechanism in the blowfly *Calliphora*. J Comp Physiol A 190, 115–129 (2004).

95. J. E. Niven et al., The contribution of Shaker K^+^ channels to the information capacity of *Drosophila* photoreceptors. Nature 421, 630–634 (2003).

96. W. Wijngaard, D. G. Stavenga, Optical crosstalk between fly rhabdomeres. Biol Cybern 18, 61–67 (1975).

97. K. Kirschfeld, “Absorption properties of photo-pigments in single rods, cones and rhabdomeres” in Processing of optical data by organisms and by machines W. Reichhardt, Ed. (Academic Press, New York, 1969), pp. 116–136.

98. E. J. Warrant, D. E. Nilsson, Absorption of white light in photoreceptors. Vision research 38, 195–207 (1998).

99. U. Tepass, K. P. Harris, Adherens junctions in *Drosophila* retinal morphogenesis. Trends Cell Biol 17, 26–35 (2007).

100. M. Vahasoyrinki, J. E. Niven, R. C. Hardie, M. Weckstrom, M. Juusola, Robustness of neural coding in *Drosophila* photoreceptors in the absence of slow delayed rectifier K^+^ channels. J Neurosci 26, 2652–2660 (2006).

101. Y. C. Gu, J. Oberwinkler, M. Postma, R. C. Hardie, Mechanisms of light adaptation in *Drosophila* photoreceptors. Curr Biol 15, 1228–1234 (2005).

102. F. Wong, B. W. Knight, F. A. Dodge, Adapting bump model for ventral photoreceptors of *Limulus*. J Gen Physiol 79, 1089–1113 (1982).

103. D. G. Stavenga, Visual acuity of fly photoreceptors in natural conditions -dependence on UV sensitizing pigment and light-controlling pupil. J Exp Biol 207, 1703–1713 (2004).

104. N. Franceschini, K. Kirschfeld, Automatic-control of light flux in compound eye of *diptera* - spectral, statical, and dynamical properties of mechanism. Biol Cybern 21, 181–203 (1976).

105. S. R. Shaw, A. Frohlich, I. A. Meinertzhagen, Direct connections between the R7/8 and R1-6 photoreceptor subsystems in the dipteran visual system. Cell Tissue Res 257, 295–302 (1989).

106. G. J. Taylor et al., Bumblebee visual allometry results in locally improved resolution and globally improved sensitivity. Elife 8 (2019).

107. A. Riehle, N. Franceschini, Motion detection in flies: parametric control over ON-OFF pathways. Exp Brain Res 54, 390–394 (1984).

108. V. Nityananda et al., Insect stereopsis demonstrated using a 3D insect cinema. Sci Rep-Uk 6 (2016).

109. B. Hassenstein, W. Reichardt, Systemtheoretische Analyse der Zeit-, Reihenfolgen- und Vorzeichenauswertung bei der Bewegungsperzeption des Rüsselkäfers Chlorophanus. Z. Naturforsch, 513–524 (1956).

110. G. G. de Polavieja, Neuronal algorithms that detect the temporal order of events. Neural Comput 18, 2102–2121 (2006).

111. M. Juusola, Z. Song, R. C. Hardie, “Phototransduction biophysics” in Encyclopedia of Computational Neuroscience, D. Jaeger, R. Jung, Eds. (Springer New York, New York, NY, 2015), 10.1007/978-1-4614-6675-8_333, pp. 2359–2376.

112. F. Galton, Vox Populi. Nature 75, 450–451 (1907).

113. H. Otsuna, K. Ito, Systematic analysis of the visual projection neurons of *Drosophila melanogaster*. Lobula-specific pathways. J Comp Neurol 497, 928–958 (2006).

114. M. Wu et al., Visual projection neurons in the *Drosophila* lobula link feature detection to distinct behavioral programs. Elife 5 (2016).

115. G. W. Meissner et al., An image resource of subdivided *Drosophila* GAL4-driver expression patterns for neuron-level searches. http://biorxiv.org/lookup/doi/10.1101/2020.05.29.080473 (2020) doi:10.1101/2020.05.29.080473 (2020).

116. K. Shinomiya et al., The organization of the second optic chiasm of the *Drosophila* optic lobe. Front Neural Circuits 13, 65 (2019).

117. M. A. Z. Dippé, E. H. Wold, Antialiasing through stochastic sampling. ACM SIGGRAPH Computer Graphics 19, 69–78 (1985).

118. J. I. Yellott, Spectral-analysis of spatial sampling by photoreceptors - topological disorder prevents aliasing. Vision research 22, 1205–1210 (1982).

119. J. I. Yellott, Spectral consequences of photoreceptor sampling in the *Rhesus* retina. Science 221, 382–385 (1983).

120. W. Salem, B. Cellini, M. A. Frye, J. M. Mongeau, Fly eyes are not still: a motion illusion in *Drosophila* flight supports parallel visual processing. J Exp Biol 223 (2020).

121. M. V. Srinivasan, S. W. Zhang, Visual motor computations in insects. Annu Rev Neurosci 27, 679–696 (2004).

122. P. G. Clarke, I. M. Donaldson, D. Whitteridge, Binocular visual mechanisms in cortical areas I and II of the sheep. J Physiol 256, 509–526 (1976).

123. K. Y. Cheng, R. A. Colbath, M. A. Frye, Olfactory and neuromodulatory signals reverse visual object avoidance to approach in *Drosophila*. Curr Biol 29, 2058–2065 (2019).

124. L. Liu, R. Wolf, R. Ernst, M. Heisenberg, Context generalization in *Drosophila* visual learning requires the mushroom bodies. Nature 400, 753–756 (1999).

125. M. G. Burg, P. V. Sarthy, G. Koliantz, W. L. Pak, Genetic and molecular-identification of a *Drosophila* histidine-decarboxylase gene required in photoreceptor transmitter synthesis. Embo J 12, 911–919 (1993).

126. J. Melzig et al., Genetic depletion of histamine from the nervous system of *Drosophila* eliminates specific visual and mechanosensory behavior. J Comp Physiol A 179, 763–773 (1996).

127. M. T. Pearn, L. L. Randall, R. D. Shortridge, M. G. Burg, W. L. Pak, Molecular, biochemical, and electrophysiological characterization of Drosophila norpA mutants. J Biol Chem 271, 4937–4945 (1996).

128. T. Washburn, J. E. O’Tousa, Molecular defects in *Drosophila* rhodopsin mutants. J Biol Chem 264, 15464–15466 (1989).

129. J. P. Kumar, D. F. Ready, Rhodopsin plays an essential structural role in *Drosophila* photoreceptor development. Development 121, 4359–4370 (1995).

130. T. Wang, X. Wang, Q. Xie, C. Montell, The SOCS box protein STOPS is required for phototransduction through its effects on phospholipase C. Neuron 57, 56–68 (2008).

131. J. Rister et al., Dissection of the peripheral motion channel in the visual system of *Drosophila melanogaster*. Neuron 56, 155–170 (2007).

132. K. G. Götz, Optomotory investigation of the visual-system of some eye mutations of the *Drosophila* fruit-fly. Kybernetik 2, 77–91 (1964).

133. Z. Song, Y. Zhou, J. Feng, M. Juusola, Multiscale ’whole-cell’ models to study neural information processing - New insights from fly photoreceptor studies. J Neurosci Methods 357, 109156 (2021).

